# A Unique Cellular Organization of Human Distal Airways and Its Disarray in Chronic Obstructive Pulmonary Disease

**DOI:** 10.1101/2022.03.16.484543

**Authors:** Samir Rustam, Yang Hu, Seyed Babak Mahjour, Scott H. Randell, Andre F. Rendeiro, Hiranmayi Ravichandran, Andreacarola Urso, Frank D’Ovidio, Fernando J. Martinez, Bradley Richmond, Vasiliy Polosukhin, Jonathan A. Kropski, Timothy S. Blackwell, Olivier Elemento, Renat Shaykhiev

## Abstract

In the human lung, terminal bronchioles (TBs), the most distal conducting airways, open to respiratory bronchioles (RBs) that lead to the alveolar region where gas exchange takes place. This transition occurs in pulmonary lobules, lung tissue units supplied by pre-TBs, which give rise to TBs. Accumulating evidence suggests that remodeling and loss of pre-TBs and TBs underlies progressive airflow limitation in chronic obstructive pulmonary disease (COPD), the third leading cause of death worldwide. Understanding the nature of these changes at the single-cell level has so far been limited by poor accessibility of pre-TBs and TBs. Here, we introduce a novel method of region-precise airway dissection, which enables capture of the entire anatomical continuum of peripheral airways, from pre-TBs to RBs, and the associated alveolar region within the lobule. This approach allowed us to identify terminal airway-enriched secretory cells (TASCs), a unique epithelial cell population of distal airways expressing secretoglobin 3A2 (SCGB3A2) and/or surfactant protein B (SFTPB). TASCs were enriched in TBs, particularly, in areas of TB-RB transition and exhibited an intermediate, broncho-alveolar molecular pattern. TASC frequency was markedly decreased in pre-TBs and TBs of COPD patients compared to those in non-diseased lungs, accompanied by changes in cellular composition of vascular and immune microenvironments. *In vitro* regeneration assays identified basal cells (BCs) of pre-TBs and TBs as a cellular origin of TASCs in the human lung. Generation of TASCs by these region-specific progenitors was suppressed by IFN-γ signaling that was augmented in distal airways of COPD patients. Thus, altered maintenance of region-specific cellular organization of pre-TBs and TBs represents a key component of distal airway pathology in COPD.

## Introduction

Distal airways, also known as “small airways” or “bronchioles” because of their small size (diameter <2 mm), constitute peripheral segments of the tracheobronchial tree. Terminal bronchioles (TBs), the most distal conducting airways, transition to respiratory bronchioles (RBs), which mark the beginning of the respiratory zone and lead to the alveolar region, where gas exchange takes place^1^. This transition occurs in secondary pulmonary lobules (“lobules”), lung units, each supplied by an individual secondary lobular bronchiole, or pre-TB, which bifurcates inside the lobule to give rise to TBs^2,3^. Accumulating evidence generated in computed tomography imaging studies indicates that remodeling and loss of distal airways, including pre-TBs and TBs, underlie progressive and irreversible airflow limitation in chronic obstructive pulmonary disease (COPD)^4–11^, a major smoking-related lung disorder and the 3^rd^ leading cause of death worldwide^12^. The nature of these structural alterations remains poorly understood at the cellular level, in part, due to poor accessibility of pre-TBs and TBs and limited knowledge about specific aspects of their cellular organization and homeostasis.

In this study, we established the method of a region-precise dissection of human distal airways at the resolution of pre-TBs and lobules and characterized their cellular organization at the single-cell level. This method enabled us to identify terminal airway-enriched secretory cells (TASCs), a unique epithelial cell state of distal airways. TASCs were marked by expression of secretoglobin 3A2 (SCGB3A2) and/or surfactant protein B (SFTPB), exhibited a molecular pattern associated with the transition of conducting airways to the respiratory zone, and were maintained by basal cells residing in pre-TBs and TBs. Loss of TASCs was observed in pre-TBs and TBs of COPD patients paralleled by changes in the cellular composition of region-specific vascular and immune microenvironments. These data suggest that altered maintenance of region-specific cellular heterogeneity of pre-TBs and TBs underlies distal airway remodeling in COPD.

## Results

### Anatomical Landmarking of Human Distal Airways

Based on the anatomical definition of a pre-TB as the last macroscopically recognizable segment along the bronchial pathway that leads to secondary pulmonary lobule^2,3^, we developed the method of a region-precise distal airway dissection, which regards pre-TBs as the boundary between pre-terminal (pre-T) and terminal (T) airways **(Fig. 1A)**. The pre-T-region included pre-TBs and 3-4 generations of airways proximal to pre-TBs, the T-region included macroscopically inseparable segments distal to pre-TBs (TBs, RBs, alveolar ducts and alveoli) within the lobule, whereas cartilaginous bronchi represented proximal (P) airways **(Fig. 1B** and **Fig. S1)**. The natural distribution of the landmark tissue elements was preserved in isolated samples: a) airway surface epithelium (ASE) with basal, ciliated and secretory cells; b) stroma with smooth muscle and vessels; c) submucosal glands (SMGs) and cartilage in P-airways; d) alveolar attachments in pre-T- and T-regions, and e) TBs, RBs, alveolar ducts and alveoli in T-region **(Fig. 1C-1E)**.

**Figure 1.**
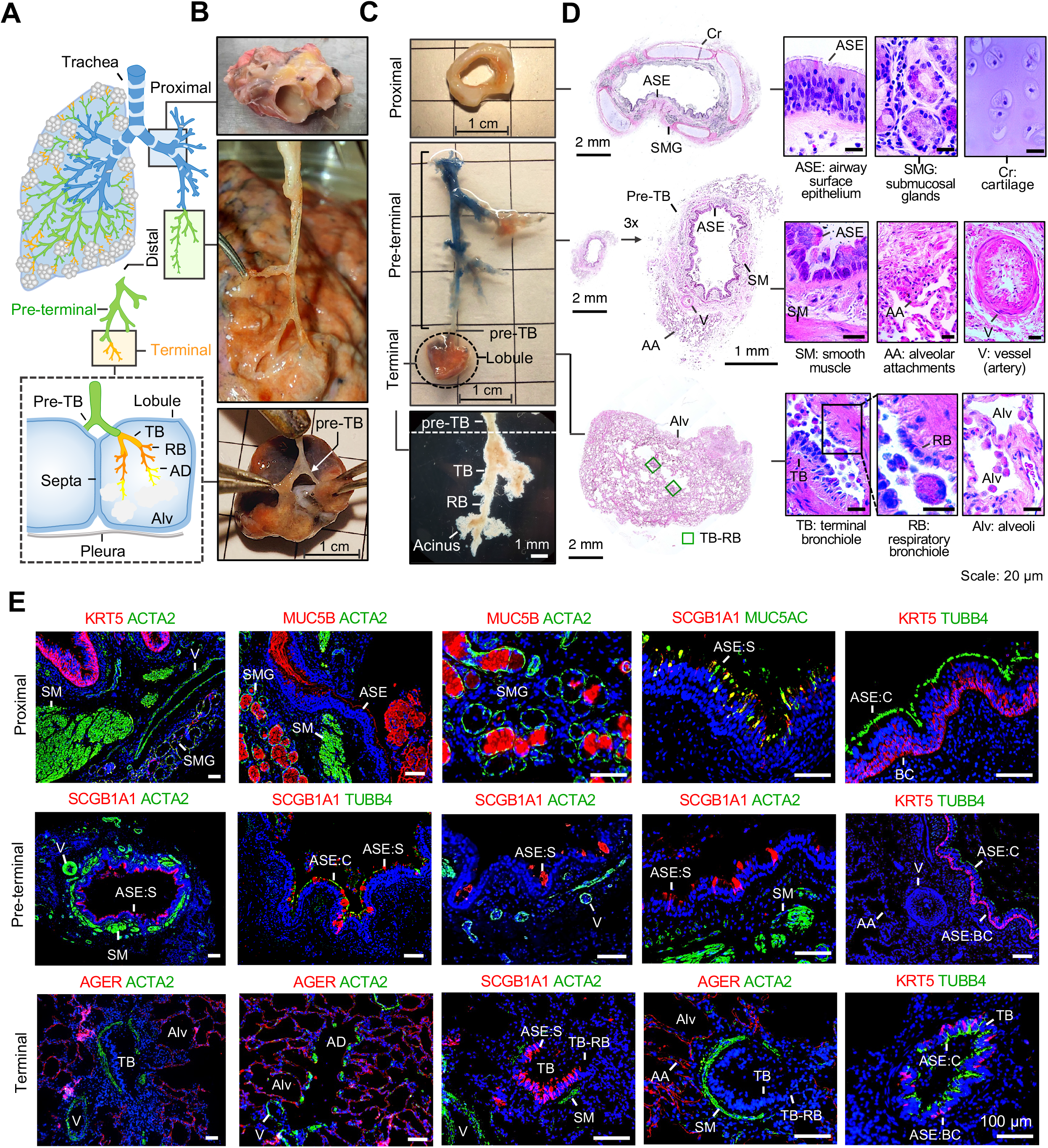
Anatomical landmarking and region-precise dissection of human distal airways. **A.** The human respiratory tree is composed of proximal (P; blue; trachea and cartilaginous bronchi) and distal (bronchioles, diameter < 2 mm) airways. The distal airway dissection method in our study regards pre-terminal bronchioles (pre-TBs), the last macroscopically visible segments in the bronchial pathway, that lead to secondary pulmonary lobules (“lobules”), as a boundary between pre-terminal (pre-T; green) and terminal (T; orange) airway regions. Pre-T region included pre-lobular bronchioles (pre-TBs and 3-4 generations proximal to pre-TBs). T-region included intralobular segments distal to visible parts of pre-TBs (terminal bronchioles [TBs], respiratory bronchioles [RBs], alveolar ducts [ADs] and alveoli [Alv]). **B.** Examples of samples used for dissection of P **(upper)**, pre-T **(middle)**, and T (**lower**; pre-TBs leading to lobules are indicated) regions. **C.** Example of dissected regions: P **(upper)**, pre-T and T **(middle**; T-region, represented by lobule, is outlined by dashed line; pre-TB marking the boundary between pre-T and T regions is indicated); **lower:** T-region inspected under tissue dissection microscope (shows intralobular segments distal to pre-TBs: TBs, RBs and alveolar tissue descending from RBs that constitute pulmonary acinus). **D.** Hematoxylin and eosin staining of representative dissected P, pre-T and T regions; major histological landmarks are shown. **E**. Immunofluorescence staining of dissected P, pre-T and T regions shown in **D**; histological landmarks are labeled as described in **D**; antibodies used to mark specific cell types: KRT5 (keratin 5; basal cells [BCs]), SCGB1A1 (secretoglobin 1A1; secretory cells [S]), mucins MUC5AC (mucus-producing S-cells of airway surface epithelium [ASE]) and MUC5B (mucinous S-cells of submucosal glands [SMG] and ASE), TUBB4 (tubulin, β4; cilia/ciliated [C] cells of ASE); ACTA2 (α-smooth muscle actin; smooth muscle [SM]); AGER (alveolar type 1 [AT1] cells; alveolar epithelium).

### Cellular Diversity along the Bronchoalveolar Axis

To systematically assess cellular heterogeneity along the P-T airway axis, transcriptomes of 111,412 single cells isolated from airway regions of 17 donors (11-67 yrs old; **Table S1**), including P-airways of 5 donors without lung disease (“normal lung”), pre-T-region of 12 normal lung donors and 5 donors with COPD, and T-region of 7 normal lung donors, were analyzed by single-cell RNA-sequencing (scRNA-seq). By performing UMAP clustering and inspecting marker genes in identified clusters, 48 cell types and subtypes were identified **(Fig. 2A** and **Fig. S2; Table S2)**. Hierarchical clustering classified these cell groups into three superfamilies: 1) epithelial, including ASE, SMG and alveolar (AT) cell families, 2) structural, including stromal and endothelial (En) cell families, and 3) immune, including myeloid and lymphoid cell families **(Fig. 2B)**.

**Figure 2.**
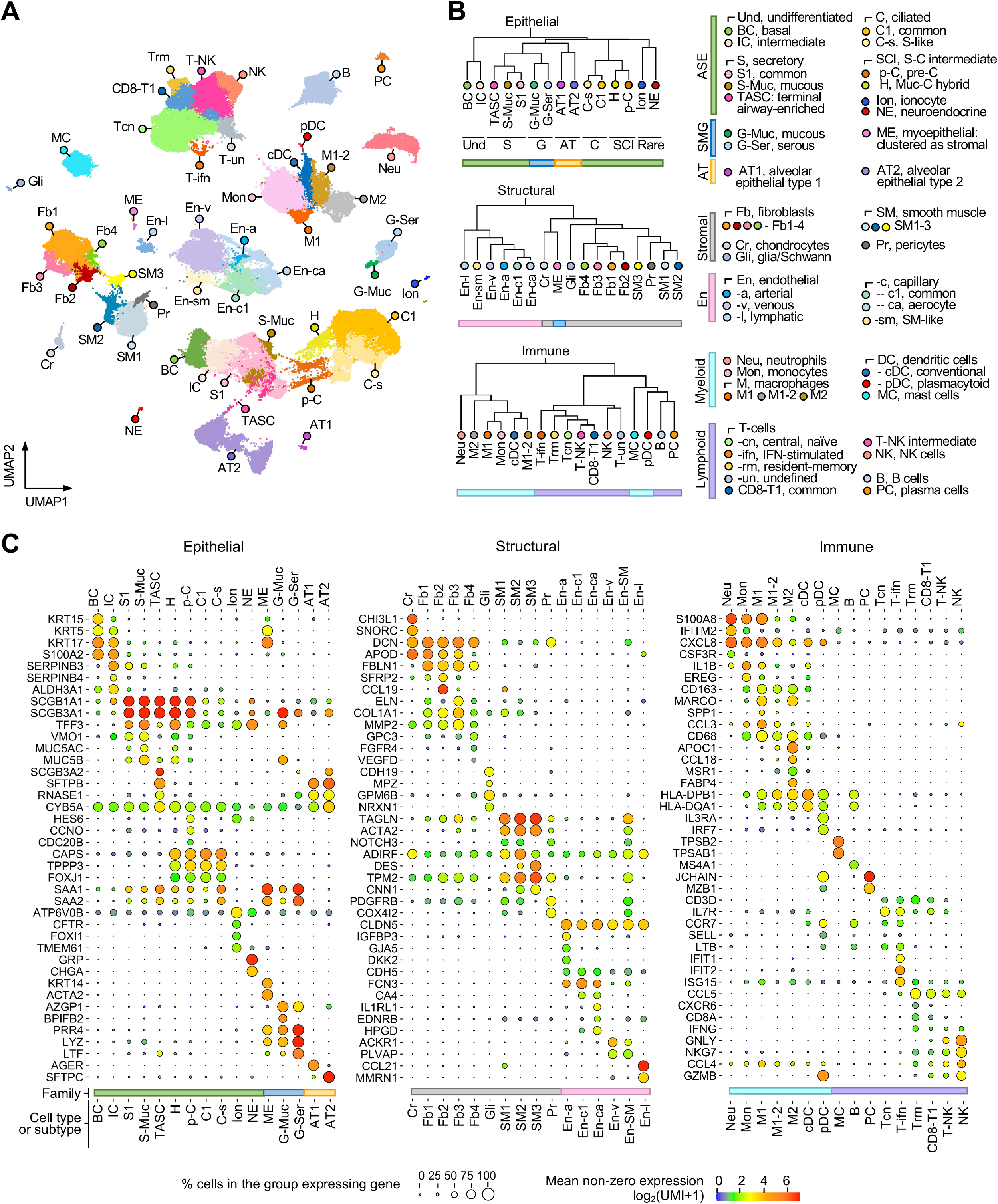
Single-cell RNA-seq analysis of cellular diversity along the bronchoalveolar axis. **A.** Uniform manifold approximation and projection (UMAP) clustering of 111,744 single cells isolated from proximal (P; n=5), pre-terminal (pre-T; n=13) and terminal regions (T; n=7) of normal lungs and pre-T airways of COPD subjects (n=5). **B.** Dendrograms showing hierarchical clustering of cell groups identified in **A** based on top 2000 variable genes using Spearman correlation and complete linkage algorithm: three large clusters representing epithelial, non-epithelial structural (“structural”) and hematopoietic/immune (“immune”) cell superfamilies are shown as separate dendrograms; cell groups are listed. **C.** Dot-plots showing expression of selected marker genes in indicated cell groups; dot size reflects percentage of cells within a group expressing given gene; dot color shows mean expression level as indicated.

#### Epithelial cells

ASE was the most diverse family of epithelial cells. It included a spectrum of cell states reflecting a trajectory of airway epithelial differentiation from undifferentiated cells *(S100A2^+^ KRT17^+^),* including basal cells (BCs; *KRT5^hi^ KRT15^hi^*), stem cells of ASE^13,14^, and intermediate cells (ICs, *SERPINB3^hi^ ALDH3A1^hi^*), to two major differentiated ASE cell types, i.e., secretory (S) and ciliated (C) cells **(Fig. 2B, C)**. S-cells *(SCGB1A1^hi^ SCGB3A1^hi^)* included common S-cells (S1), mucus-producing cells (S-Muc, *TFF3^hi^ MUC5AC^hi^ MUC5B^hi^ VMO1^hi^*), and a distinct subset of cells expressing *SCGB3A2* and *SFTPB,* which formed a “bridge” cluster between S- and type 2 alveolar epithelial (AT2) cells **(Fig. 2A, C)**. Cells in this intermediate cluster were present almost exclusively in pre-T and T-regions, being enriched in the latter (see below), and, therefore, were named “terminal airway-enriched secretory cells” (TASCs).

C-cells *(CAPS^+^ FOXJ1^+^)* included major (C1) and secretory-like (C-s; *SAA1^+^ SAA2^+^)* subsets. Two clusters were found between C- and S-cells (S-C-intermediate; SCI cells; **Fig. 2A, B)**. One of these SCI clusters expressed markers of deuterosomal precursors of C-cells *(CCNO, CDC20B, HES6*)^15^ and was termed “pre-ciliated” (p-C) cells. Another co-expressed C, S1 and S-Muc marker genes, and, consistent with recently reported findings^15–17^, was named “S-C hybrid” (H) cells. Our analysis also captured rare ASE cell types - ionocytes (Ion, *ATP6V0B^+^ FOXI1^+^ TMEM61^+^ CFTR^+^*)^18,19^ and neuroendocrine (NE) cells *(CHGA^+^ GRP^+^)* (**Fig. 2C**).

The SMG family included glandular (G; *AZGP1^hi^)* mucous (G-Muc; *MUC5B^+^ BPIFB2^+^*) and serous (G-Ser; *LYZ^hi^ LTF^hi^ PRR4^hi^)* cells^20^, and myoepithelial (ME) cells *(KRT5^+^ KRT14^+^ ACTA2^+^,* SMG stem cells^21,22^) **(Fig. 2B, C)**. The AT family included type 1 (AT1; *AGER^+^*) and type 2 (AT2; *SFTPC^+^*) alveolar epithelial cells **(Fig. 2C)**. Consistent with their intermediate clustering between S- and AT2 cells, TASCs, in addition to common S-cell genes, expressed a set of genes enriched in AT2 cells *(SFTPB, RNASE1, CYB5A*), but, in contrast to the latter, did not express *SFTPC*, the obligatory AT2 cell marker **(Fig. 2C**; and below).

#### Structural cells

The stromal family included chondrocytes (Cr; *SNORC^+^ CHI3LI^+^*), fibroblasts (Fb; *FBLN1^+^ DCN^+^*), smooth muscle cells (SM; *ACTA2^+^ TAGLN^+^)* and pericytes *(PDGFRB^+^ COX4I2^+^)* **(Fig. 2C)**. Fb included four subsets: common (Fb1), *CCL19^+^* (Fb2; included *SFRP2^+^* adventitial Fb^23^), matrix (Fb3, *ELN^hi^ COL1A1^hi^ MMP2^hi^*; contained *ACTA2^+^* myoFbs^24,25^), and Fb4 (contained *GPC3^+^ FGFR4^+^ VEGFD^+^* alveolar Fbs^23^). SM included SM1 and SM2 clusters expressing markers of vascular SM *(ADIRF, NOTCH3, PDGFRB*)^23^ and SM3 *(DES^+^ TPM2^+^ CNN1^+^,* airway SM markers^23^) **(Fig. 2C)**. A distinct group of cells clustered among stromal cells and expressed markers of neuronal and glial/Schwann cells *(CDH19, GPM6B, MPZ, NRXN1)* and, therefore, was annotated as glial/Schwann cells (Gli; **Fig. 2B, C**). The endothelial (En) cell family *(CLDN5^hi^ CDH5^+^)* included arterial (En-a; *IGFBP3^+^ GJA5^+^ DKK2^+^*), common capillary (En-c1; *FCN3^+^ CA4^+^*), a distinct cluster of En-capillary cells expressing markers of recently described alveolar capillary aerocytes^23^ (En-ca; *HPGD*, *EDNRB*, *IL1RL1*), venous (En-v; included cells expressing markers of the fenestrated endothelium *ACKR1* and *PLVAP*^23^), En-sm co-expressing En- and SM markers, and lymphatic (En-1; *CCL21^+^MMRN^+^)* **(Fig. 2C)**.

#### Immune cells

Myeloid cells included neutrophils (Neu; *S100A8^hi^ IFITM2^hi^ CXCL8^hi^ CSF3R^+^*)^23^, mast cells (MC; *TPSB2^+^ TPSAB1^+^*), a spectrum of cellular states representing the mononuclear phagocyte system - from monocytes (Mon; *IL1B^+^ EREG^+^)* to macrophages (M; *CD163^hi^ CD68^hi^)* and dendritic cells (DCs). The latter included conventional, or myeloid, DCs (cDCs) marked by high expression of the *HLA* family genes, and plasmacytoid (pDCs; *IL3RA^+^IRF7^+^)* **(Fig. 2B, C)**. Three M-subsets were identified: inflammatory (M1; *SPP1^+^ CCL3^+^*), non-inflammatory (M2; *MSR1^+^ FABP4^+^ CCL18^+^*), which resembled classically (M1) and alternatively (M2) activated macrophage polarization states^26^, both expressing *MARCO,* a marker of embryonically derived lung-resident macrophages^27,28^, and intermediate (M1-2) state **(Fig. 2C)**.

The lymphoid family included B (*MS4A1^+^*), plasma (PC; *JCHAIN^+^ MZB1^+^*), T (*CD3D^+^*) and NK *(GNLY^hi^ NKG7^hi^)* cells **(Fig. 2C)**. Multiple T-cell subsets were identified, including those expressing markers of central memory and naïve T cells (Tcn; *CCR7^+^IL7R^+^ LTB^+^ SELL^+^*), T-cells expressing interferon (IFN)-response genes (T-ifn, *IFIT1^+^IFIT2^+^ISG15^+^*), two CD8^+^ T cell-enriched subsets - resident-memory (Trm; *CXCR6^+^ CCL5^+^)* and “common” CD8^+^ T cells (CD8-T1), and T-cells co-expressing NK cell markers (T-NK; *GNLY^+^ NKG7^+^*)^29–31^ **(Fig. 2C)**.

### Regional Heterogeneity of Epithelial Cells along the Bronchoalveolar Axis

SMG cells were uniquely enriched in P-airways, whereas AT cells were found only in pre-T and T-regions **(Fig. 3A)**, consistent with unique histological organization of these regions **(Fig. 1D; Fig. S3)**. Undifferentiated ASE cells (BCs and ICs) were most abundant in P-airways (43% of ASE cells *vs* <10% in pre-T and T; p<0.004), largely due to enrichment of ICs in P-airways **(Fig. 3A, B; Table S3)**, where *SERPINB3^+^* ICs contributed to the pseudostratified ASE pattern typical for P-airways **(Fig. S3A)**. The overall frequency of secretory-ciliated intermediate (p-C and H) cells was higher in pre-T and T-airways (8-10% of ASE cells *vs* 1.4% in P; p<0.02; **Fig. 3B)**. The overall frequency of S- and common S1-cells was similar across the regions **(Fig. 3A, B)**. However, TASCs, a novel S-cell population identified in this study, were found almost exclusively in distal (pre-T and T-regions) with significant enrichment in the T-region (p<0.02 *vs* pre-T; **Fig. 3B; Table S3**), the basis for their designation as T-airway-enriched S-cells.

**Figure 3.**
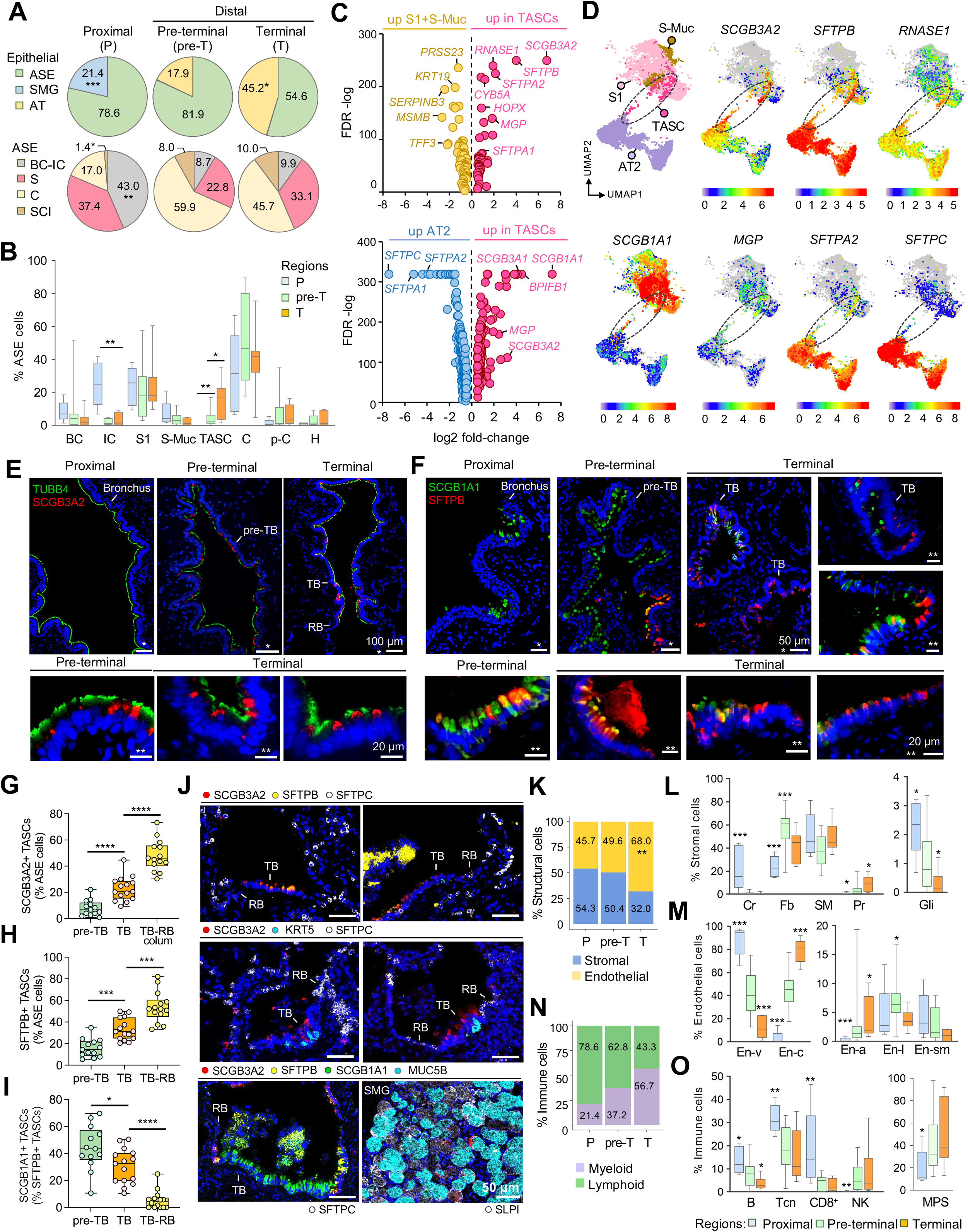
Region-specific cellular heterogeneity along the bronchoalveolar axis. **A.** Proportions (%) of cells belonging to indicated families based on scRNA-seq data described in **Fig. 2** (see details in **Table S3**) within epithelial superfamily (**upper;** ASE; airway surface epithelium; SMG; submucosal gland; AT; alveolar epithelium): average % of cells per sample in indicated regions of non-diseased lungs: 5 proximal, 13 pre-terminal, 7 terminal samples); and in ASE family (**lower;** BC-IC, undifferentiated [basal and intermediate], S, secretory [S1 common S, S-Muc, mucous; TASCs, terminal airway-enriched S], C, ciliated [C1, common, and C-s, S-like], SCI, S-C intermediate [H, hybrid; p-C, pre-ciliated] cells; **B.** Average % of cells belonging to indicated cell types (per sample; as described in **A**) within ASE family in different regions; **C.** Volcano plots (based on scRNA-seq data shown in **Fig. 2**): differentially expressed genes (DEGs) identified by comparing all: **(upper)** TASCs and other S (S1 and S-Muc) cells; **(lower)** TASCs and AT type 2 (AT2) cells; shown DEGs with FDR <0.01 and log2FC >0.5 or < −0.5. **D.** Portion of the UMAP graph showed in **Fig. 2A** including only S1, S-Muc, TASC and AT2 cells. A clustering region, where TASCs are enriched and form a bridge between S- and AT2 cells, is demarcated (dashed line). Expression of selected DEGs identified in the analysis shown in **C** is color-mapped (see scale bar for expression levels). **E, F.** Immunofluorescence (IF) images showing expression of **E:** secretoglobin 3A2 (SCGB3A2) and tubulin beta 4 (TUBB4; cilia marker); **F:** surfactant protein B (SFTPB) and SCGB1A1 (S1-cell marker) in indicated regions; **G**-**I.** % of **G:** SCGB3A2^+^ TASCs; **H:** SFTPB^+^ TASCs among ASE cells, and **I**: SCGB1A1^+^ cells among SFTPB^+^ TASCs in indicated regions; TB-RB: simple epithelium (colum, columnar) in TB-RB transition areas. **J.** Representative CyTOF images showing expression of indicated markers in different regions; **K.** Proportions of structural cell families (average % cells per sample) in different regions; **L, M.** Average % of indicated cell types (per sample; as described in **A**; cell type names as described in **Fig. 2B**) within stromal **(L)** and endothelial (En; **M**) cell families; **N.** Proportions of cell representing immune families (average % cells per sample) in different regions; **O.** Average proportions of indicated cell types (per sample; as described in **A**; cell type names as described in **Fig. 2B**) in different regions. In **A, B, G-I, K-O**: *p<0.05, **p<0.01; ***p<0.005; ****p<0.0001 (two-tailed Mann-Whitney test) *vs* other or indicated regions.

### Terminal Airway-enriched Secretory Cells (TASCs)

To comprehensively characterize the molecular phenotype of TASCs, we compared their transcriptomes to those of their closest clustering neighbors - other ASE S- (S1 and S-Muc) cells and AT2 cells **(Fig. 3C, D)**. Consistent with their intermediate clustering position, between S-and AT2 cells, TASCs exhibited a transitional molecular pattern integrating features of these cell types yet were distinct from each of them **(Fig. 3C)**. Compared to other ASE S-cells, TASCs had significantly higher expression of TASC markers *SCGB3A2* and *SFTPB,* and a distinct set of genes highly expressed in AT cells, including *RNASE1, SFTPA2, CYB5A, HOPX*and *SFTPA1* **(Fig. 3C, D)**. Compared to AT2 cells, TASCs had significantly higher expression of airway S-cell markers *SCGB1A1, SCGB3A1* and *BPIFB1,* but lower expression of AT2 marker genes *SFTPC, SFTPA1* and *SFTPA2.* Notably, *SCGB3A2* and *MGP* were expressed in TASCs higher than in both S- and AT2 cells **(Fig. 3C, D; Table S4)**. TASCs were *bona fide* single cells, as determined by a rigorous doublet detection protocol^32^ (see **Supplemental Methods)**.

Consistent with scRNA-seq data, immunofluorescence (IF) analysis of normal lung tissue samples confirmed that TASCs, identified as ASE cells containing secretoglobin SCGB3A2 and surfactant protein SFTPB, are unique to distal airways **(Fig. 3E, F; Fig. S4)**. IF analysis of 24 normal lung tissue samples (>80,000 ASE cells; **Table S5)** determined that SCGB3A2^+^ TASCs constitute 8% and 23% of ASE cells in pre-TBs and TBs, respectively (p<0.0002 TB *vs* pre-TB) and >40% in the transitioning simple columnar-to-cuboidal epithelium at TB-RB junctions (p<10^-4^ *vs* TB overall; **Fig. 3G**). SFTPB marked a broader subset of TASCs (~16% of ASE cells in pre-TBs; ~33% in TBs; p<0.0003, TB *vs* pre-TB; ~54% in TB-RB junction areas; p=0.0002 *vs* TB overall; **Fig. 3H)**. Compared to a dome-shaped appearance of TASCs in the pseudostratified epithelium of pre-TBs and proximal aspects of TBs, in TB-RB junctional areas, where the epithelium transitioned to a simple columnar-to-cuboidal pattern, TASCs acquired a cuboidal shape typical for the transitional cuboidal epithelium of RBs^33^ **(Fig. S4A-B)**.

The molecular pattern of TASCs changed along the pre-TB-RB axis. In pre-TBs, 44.2% of TASCs co-expressed SCGB1A1, a marker of common S-cells, compared to 30.5% of TASCs in TBs (p<0.03) and only 5.5% in TB-RB junctional areas (p<10^-5^ *vs* TB overall; **Fig. 3I**). At the single-cell mRNA level, TASCs in the T-region showed significantly higher expression of *SFTPA2* compared to those in pre-T airways **(Fig. S5)**, but in contrast to AT2 cells, which also expressed this gene, TASCs were largely negative for *SFTPC* **(Fig. 3J)**. As shown earlier^34^, a subset of SMG cells were SCGB3A2^+^, but they did not express *SFTPB* or *SCGB1A1* **(Fig. 3J; Fig. S3B and S4)** and, based on their transcriptome profiles, clustered separately from TASCs **(Fig. 2A)**. Thus, TASCs represent a distinct subset of S-cells unique to distal airways and enriched in areas of TB-RB transition.

### Regional Heterogeneity of Structural and Immune Cells

Distribution of structural cells showed differences along the P-T axis, with significantly higher frequency of endothelial (En)-cells in the T-region **(Fig. 3K)**. Among stromal cells, chondrocytes, as expected, were found in cartilaginous P-airways, fibroblasts were most frequent in pre-T and less frequent in P-airways **(Fig. 3L)**. Among fibroblasts, Fb2 (*CCL19^+^*) were most frequent in the P-region, whereas proportions of Fb3 and Fb4 increased along the P-T axis **(Table S3)**. Pericytes also increased in frequency along this axis **(Fig. 3L**). By contrast, the frequency of Gli-cells declined along the P-T axis from 2.4% of stromal cells in P-airways to 1.1% in pre-T (p<0.05 *vs* P) and 0.3% in T-region (p=0.0057 *vs* P+pre-T; **Fig. 3L)**. Among endothelial cells, capillary (En-cl and En-ca) cells were significantly enriched in the T-region (p<0.0003 *vs* other regions), whereas the proportion of En-v cells, representing venous and fenestrated endothelium, was highest in P-airways (p=0.00125 *vs* other regions) **(Fig. 3M)**. The frequency of endothelial lymphatic (En-1) cells was highest in pre-T-region **(Fig. 3M)**.

Within the immune cell superfamily, the distribution of myeloid and lymphoid cells showed opposite trends along the P-T axis, with the trend of a higher proportion of lymphoid cells in P-airways **(Fig. 3N)**. Among lymphoid cell types, B, Tcn, CD8^+^ (Trm + CD8^+^T1) showed a higher frequency in P-airways, whereas NK cells were more frequent in pre-T and T-regions **(Fig. 3O)**. The total frequency of cells of the mononuclear phagocyte system (Mon, M, DCs) in pre-T and T regions was higher compared P-airways **(Fig. 3O; Table S3)**.

### Altered Region-specific Cellular Composition of Distal Airways in COPD

Pre-T airways isolated by our method included pre-TBs and more proximal bronchioles with a luminal diameter <2 mm **(Fig. 1A)** and, thus, represented small airways known to be affected in COPD^4–8^. To determine cellular basis of disease-related pathologic changes in these airways at the single-cell level, we compared the cellular composition of pre-T samples isolated from 5 subjects with COPD and 12 donors without lung disease (“normal” lungs) using scRNA-seq analysis. Whereas the overall distribution of major cell types was comparable in both groups **(Fig. 4A)**, disease-related changes preferentially involved cell populations, which contributed to physiological cellular heterogeneity along the airway axis. This included the loss of TASCs and En-c cells normally enriched in the distal airway region, and increased proportions of Gli and CD8-T1 cells normally enriched in P-airways. There was also a dramatic, 4.7-fold increase in the frequency of MCs in pre-T airways of COPD patients compared to normal. Proportions of these cells were significantly different in pre-T airways of COPD patients compared to normal lung donors among all captured cells **(Fig. 4A)** and within corresponding superfamilies **(Fig. 4B)**.

**Figure 4.**
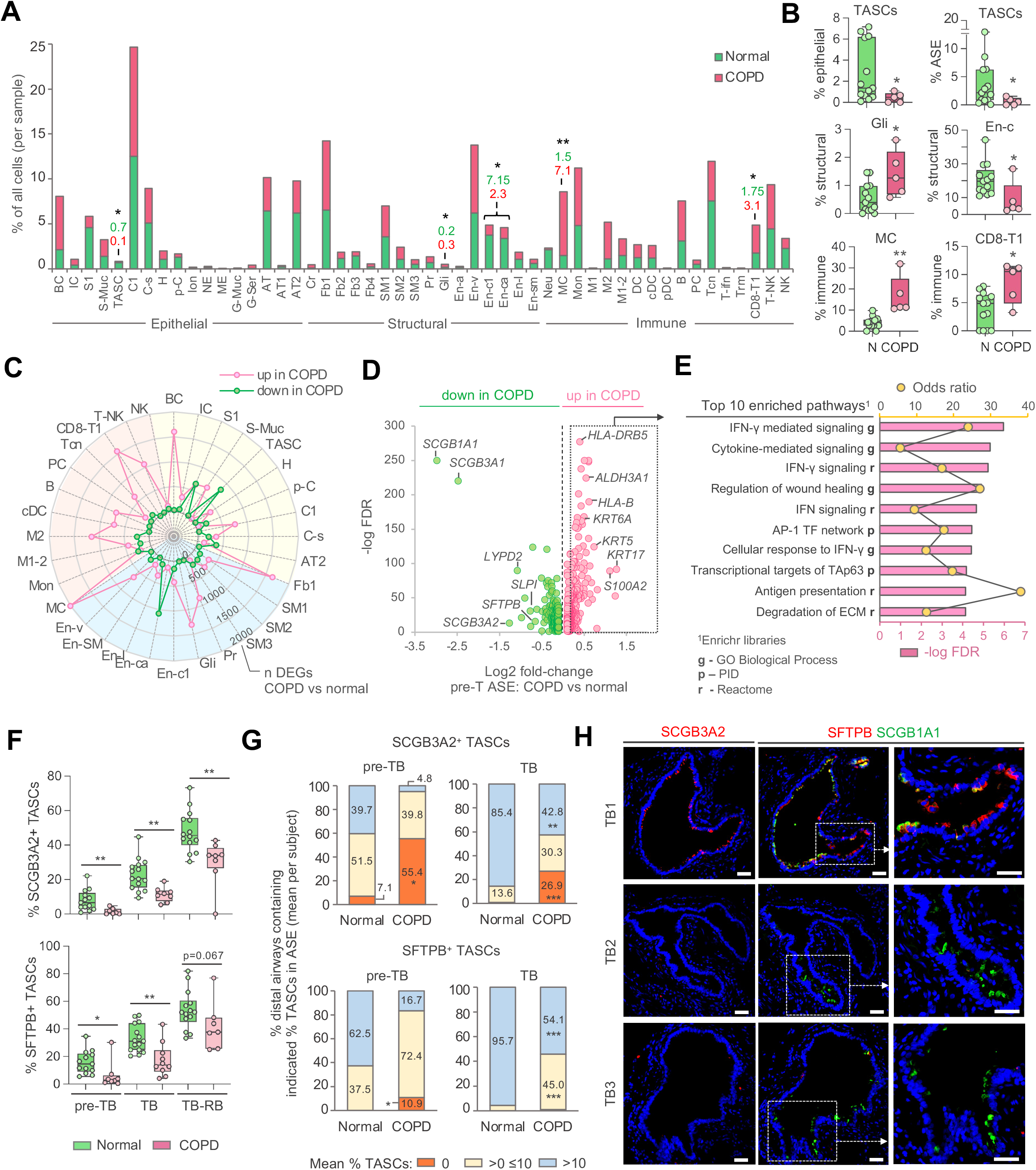
Changes in cellular organization of distal airways in COPD. **A, B.** scRNA-seq analysis-based proportions of indicated cell types and subtypes (abbreviations as described in **Fig. 2B**) in pre-terminal (pre-T) airway samples of donors without lung disease (“normal”; n=13) and those from subjects with COPD (n=5): **A.** Mean percentage (%) of cells in indicated cell groups among all single cells per sample (values are shown for cell groups with significant differences between the groups; green, normal; pink, COPD); **B.** Mean % of cells in indicated cell groups among all single cells per sample within corresponding superfamilies (dots: individual samples). **C.** Radar plot showing numbers of differentially expressed genes (DEGs) identified by comparing scRNA-seq average gene expression profiles of indicated cell types/subtypes in pre-T samples (COPD *vs* normal) described in **A-B**. Criteria for cell type/subtype inclusion: >10 cells per group (normal; COPD) and at least 3 cells in >50% samples in each group. Colored areas - epithelial (yellow), structural (blue), and immune (red) cell superfamilies. Dots show numbers of DEGs up- (pink) or down-regulated (green) in COPD *vs* normal samples (Mann-Whitney p<0.05). **D.** Volcano plot showing DEGs (FDR <0.05) identified by comparison of scRNA-seq average expression profiles of airway surface epithelial (ASE) cells in COPD *vs* normal pre-T samples (described in **A-B**). **E.** Top 10 annotation categories (pathways) among Gene Ontology (GO) Biological Process; Reactome and NCI-Nature Pathway Interaction database (PID) libraries identified as enriched in DEGs up-regulated in ASE of COPD pre-T *vs* ASE of normal pre-T samples (log2 fold-change >0.2; dashed box in **D**); ranked by -log FDR of enrichment; dots - odds ratio of enrichment based on Enrichr analysis. **F-H.** IF analysis: **F.** mean % of SCGB3A2^+^ **(upper)** and SFTPB^+^ **(lower)** TASCs per subject among ASE cells in pre-TB, TB, and TB-RB (defined as in **Fig. 3G-I**) of normal lung donors (green) and COPD subjects (pink); dots represent individual subjects (see **Table S5** for details). **G.** Distribution of distal airways (pre-TBs and TBs) in normal and COPD lung tissue samples (described in **F**) based on average % of SCGB3A2^+^ **(upper)** and SFTPB^+^ **(lower)** cells in ASE of these airways (per subject); in **A**, **B**, **F** and **G**: *p<0.05, **p<0.01; ***p<0.005 (two-tailed Mann-Whitney test) comparing COPD vs normal. **H.** Representative IF images showing SCGB3A2^+^ TASCs (left), SFTPB^+^ TASCs and SCGB1A1 expression (right) in different TBs of the same COPD patient’s lung; scale bars: 50 μm.

To identify disease-related molecular changes in specific cell types and subtypes, we compared gene expression profiles of specific cell populations captured in scRNA-seq analysis of pre-T airways of COPD patients and normal lung donors **(Fig. 4C)**. The highest numbers of up-regulated genes in COPD *vs* non-diseased pre-T samples were observed in MCs (2716 genes), BCs (1605 genes), Fb1 (1641 gene), CD8-T1 (1560 genes), Gli (1302 genes), M2 (1151 gene) and T-NK (1074 genes). The highest numbers of suppressed genes in COPD compared to normal pre-T samples were found in En-ca subset of capillary En-cells (1072 genes), TASCs (831 gene) and common S (S1) cells (648 genes) **(Fig. 4C; Table S6)**. Thus, altered cellular composition of pre-T airways in COPD subjects is paralleled by changes in transcriptional states of a distinct set of cell populations, including those contributing to the region-specific pattern of distal airways.

To determine most prevalent molecular changes occurring within the ASE compartment of COPD pre-T airways, we performed pseudobulk analysis of transcriptional profiles of pre-T ASE cells of COPD subjects compared to those from normal lung donors using scRNA-seq data described above. Among top down-regulated genes were common S (S1) cell markers, including *SCGB1A1* and *SCGB3A1,* and markers of TASCs *SCGB3A2* and *SFTPB* **(Fig. 4D)**. Thus, loss of secretory differentiation features, both general and region-specific, represents a major disease-related transcriptional phenotype of ASE in COPD pre-T airways. Among the genes up-regulated in ASE of COPD pre-T airways were those related to the *HLA* family *(HLA-DRB5* and *HLA-B)* and markers of undifferentiated ASE cells, BCs and ICs, *ALDH3A1, KRT6A, KRT5, KRT17* and *S100A2}* **(Fig. 4D; Table S7)**. Immune signaling pathways, including those related to IFN-γ signaling, were identified as the top biological categories enriched among genes up-regulated in ASE of COPD *vs* normal pre-T-airways **(Fig. 4E; Table S8)**.

In agreement with scRNA-seq data, quantitative IF analysis, in which >50,000 ASE cells lining pre-TB/TBs in lung tissue samples of 11 COPD subjects were assessed comparatively to non-diseased samples described above, revealed significantly reduced frequency of SCGB3A2^+^ and SFTPB^+^ TASCs in pre-TBs and TBs of COPD subjects **(Fig. 4F; Table S5)**. Particularly dramatic loss of SCGB3A2^+^ TASCs in the lungs of COPD patients was in pre-TBs (on average, 1.5% of ASE cells *vs* 8.0% in non-diseased donors; p<0.005). The average frequency of SFTPB^+^ TASCs in the ASE of COPD pre-TBs was 6.3% *vs* 15.9% in normal lung donors (p<0.007) **(Fig. 4F)**. Whereas in non-diseased lungs, TASCs were detected in all inspected pre-TBs (based on SFTPB positivity) and SCGB3A2^+^ TASCs were found in >92% of them, no single TASC could be detected in 55.4% and 10.9% of pre-TBs in COPD subjects, based on SCGB3A2 and SFTPB positivity, respectively (p<0.05 *vs* non-diseased donors; **Fig. 4G**).

The average frequency of TASCs in TBs of COPD subjects was about two times lower compared to that in non-diseased lungs based on both SCGB3A2 and SFTPB staining (p<0.003 for both markers; **Fig. 4F**). Whereas >99% of inspected TBs in non-diseased lungs contained SCGB3A2^+^ TASCs, this subset of TASCs was not detected in 26.9% of TBs of COPD subjects **(Fig. 4G)**. There was also significantly lower proportion of TBs containing TASCs at frequency of >10% of ASE cells in COPD compared to non-diseased lungs, based on both SCGB3A2 and SFTPB expression **(Fig. 4G)**. In TB-RB junction areas, where TASCs were most abundant in both groups, differences in their frequency between COPD and normal lungs were less dramatic, but still significant for SCGB3A2^+^ TASCs **(Fig. 4F; Table S5)**. A remarkable heterogeneity was noted with regard to the extent of TASC loss in the lungs of different COPD patients and/or in different pre-TB/TBs regions of the same patient lungs **(Fig. 4F-H)**. In some COPD samples, TASCs were preserved in a subset of pre-TB/TBs but undetectable in other distal airways of the same anatomical category **(Fig. 4H; Fig. S6A-B)**.

Notably, common SCGB1A1^+^ S-cells could still be detected in a subset of pre-TBs/TBs of COPD patients, where TASCs were absent **(Fig. 4H; Fig. S6B)**, indicating that loss of TASCs in COPD occurs earlier and is independent from changes in common S (S1) cell differentiation in these airways. In support of the latter possibility, based on quantitative IF analysis, there was no significant change in the frequency of SCGB1A1^+^ S1-cells in pre-TBs and TBs of COPD subjects, and no significant correlation was observed between loss of TASCs and changes in the overall SCGB1A1^+^ S1-cell frequency in pre-TBs and TBs **(Fig. S7)**. Further, no significant change was detected in the frequency of TASCs in pre-TBs and TBs of smokers without lung disease compared to nonsmokers **(Fig. S8)**, suggesting that loss of TASCs in distal airways of COPD patients represents a disease-related phenotype rather than a common response of distal airways to cigarette smoking, the major risk factor for COPD^12^.

### Distal Airway BCs are Cellular Origin of TASCs

To determine the biological basis of regional heterogeneity of epithelial cells along the proximal-distal airway axis, we focused on airway BCs, stem cells of ASE^13,35^. We hypothesized that, as local epithelial progenitors^36^, BCs maintain a regional memory, an intrinsic capacity to regenerate epithelium with cellular composition specific to the region of their residence. To test this hypothesis, we isolated BCs from proximal (P) and distal (D: pre-T) airways of 6 donors without lung disease **(Table S10)** and compared their differentiation potential in the air-liquid interface (ALI) model. In this model, BCs after establishing a confluent monolayer generate differentiated airway epithelium within few weeks upon apical exposure to air^37^ **(Fig. 5A)**. Paired comparative analysis of transcriptomic profiles of epithelia generated by P- and D-airway BCs after 18 days of ALI culture identified 2,005 DEGs, including 888 and 1117 genes with significantly higher expression in epithelia produced by P- and D-airway BCs, respectively **(Table S11)**. Among the genes uniquely up-regulated in epithelia generated by D-airway BCs were *SFTPB, SCGB3A2, SFTA2, RNASE1* and *SFTPA2* **(Fig. 5B)**, i.e., genes, which as described above, were uniquely expressed or enriched in TASCs *in vivo.* Indeed, 21.3% of genes most robustly up-regulated in D-*vs* P-airway BC-derived epithelia (log2FC >1) were genes that distinguished TASCs from other ASE cells *in vivo* **(Fig. 5C)**. Consistent with these data, IF analysis identified SCGB3A2^+^ cells in differentiated epithelia derived from D-airway BCs, but not P-airway BCs **(Fig. 5D)**. By contrast, MUC5AC^+^ mucus-producing S-cells were found more frequently in epithelia derived from P-airway BCs **(Fig. 5D)**, although this was not a consistent region-specific feature of P-airway epithelium at the gene expression level. Together, these data suggest that the ability to generate TASCs is a unique, region-specific feature of D-airway BCs, and even after isolation of BCs from their *in vivo* microenvironment, these progenitors continue to maintain an intrinsic memory of region-specific differentiation.

**Figure 5.**
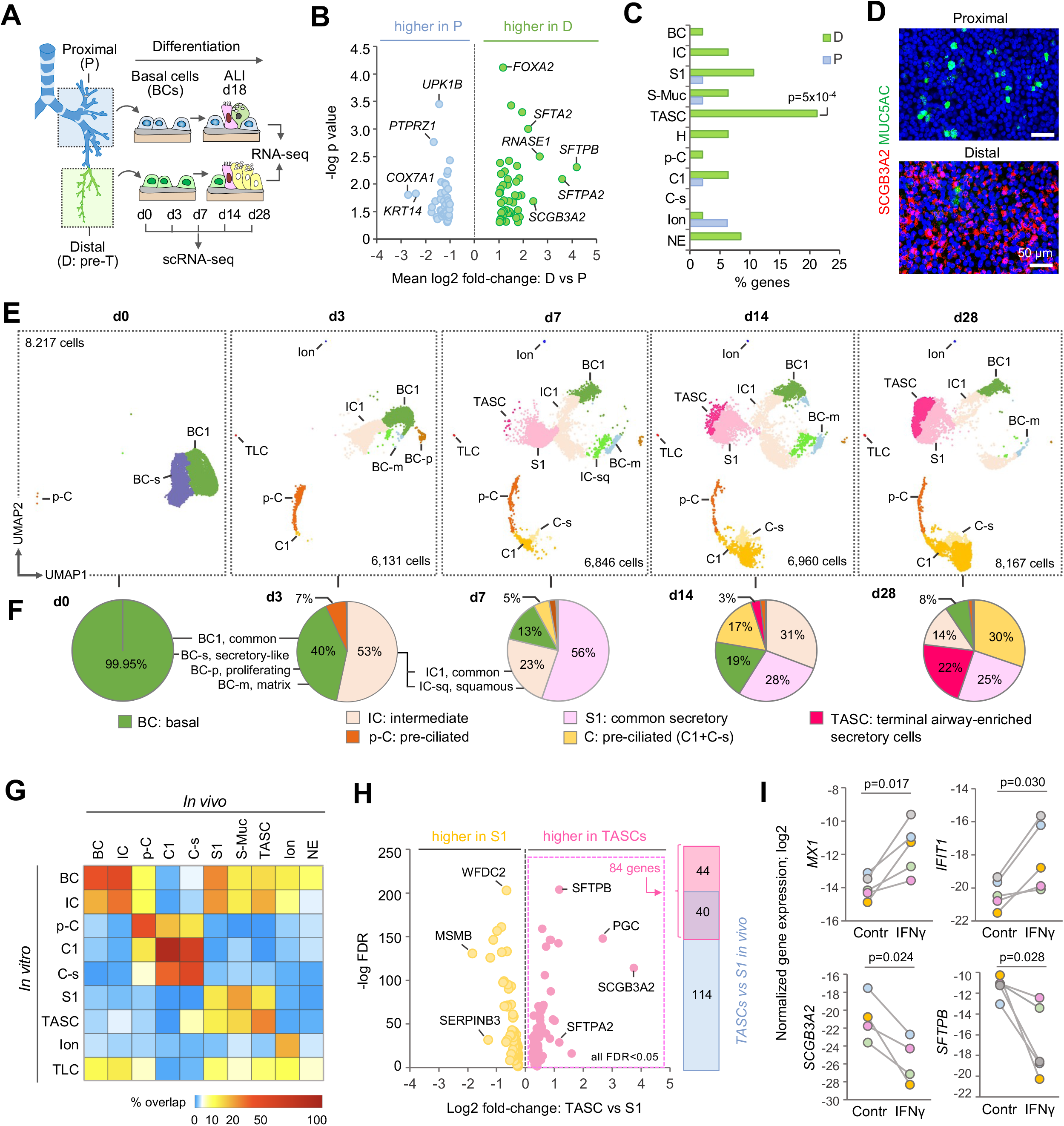
Distal airway basal cells (BCs) as the cellular origin of terminal airway-enriched secretory cells (TASCs). **A.** Design of *in vitro* studies. BCs isolated from proximal (P) and distal (D; pre-terminal [pre-T] region) were cultured under the same conditions in the transwell system. After BCs established confluent layer, apical media was removed to establish air-liquid interface (ALI) and media was supplied only via the basolateral side. At indicated time-points of ALI culture representing different stages of BC differentiation samples were analyzed. **B.** Volcano plot showing differentially expressed genes (DEGs) identified by comparison of transcriptomes of ALI-day 18 epithelia generated by BCs isolated from P- and D-airways of 6 donors without lung disease (shown DEGs with log2 FC >1, 2-tailed paired t-test p<0.05). **C.** Proportions (%) of DEGs up-regulated in ALI-day 18 ASE generated by BCs from matched P (blue) and D (green) airways (described in **B**) overlapping with marker genes of indicated pre-T ASE cell types based on scRNA-seq analysis (criteria: log2FC >1; FDR<0.05). **D.** Immunofluorescence staining of ALI-day 18 membranes showing apical surface of epithelia generated by BCs from P- (upper) and D- (lower) airways; red: secretoglobin SCGB3A2, green: mucin MUC5AC; blue: DAPI. **E.** UMAP clustering of cells at indicated time-points of ALI culture of the same distal airway BC sample; total number of single cells captured in each time-point is indicated. **F**. % of cell types at ALI time-points shown in **E**. TLCs, tuft-like cells. **G.** Heatmap: % DEGs of indicated ASE cell types *in vivo* overlapping with those identified for cell types in ALI-day 28 epithelia derived from D-airway BCs shown in **E** using the same criteria: FDR<0.05; log2 fold-change >0.5 *vs* other ASE cells. **H.** Volcano plot: DEGs (FDR <0.05) identified by comparison of TASCs and S1-cells in ALI-day 28 epithelia generated by D-airway BCs (shown in **E**); right side: overlap between DEGs up-regulated in TASCs *vs* S1 *in vitro* and those identified in the same comparison *in vivo*. **I.** log2 normalized expression of indicated genes in ALI-day 18-40 epithelia generated by distal airway BCs in the presence or absence of 5 ng/ml IFN-γ; dot colors mark independent samples; p values based on 2-tailed paired t-test; see details in **Fig. S9D**.

To evaluate the dynamics of distal airway epithelial differentiation at the single-cell level, epithelia generated at days 0, 3, 7, 14 and 28 of ALI culture of the same D-airway BC sample were profiled using scRNA-seq. A total of 36,321 single cells were captured in this analysis **(Table S12)**. As expected, cellular diversity progressively increased during BC differentiation **(Fig. 5E, F)**. At ALI-day 0, >99% of cells were BCs, which included two subsets: BC1, which expressed typical BC markers, and S-like BCs (BC-s), which differed from BC1 by expression of S-cell-related genes, e.g., *SCGB1A1* and *MUC5B* (see markers of all cell types in **Table S13**). At ALI-day 3, in addition to BCs, which at this stage included 3 subsets: BC1, proliferating (BC-p), and matrix (BC-m) showing high expression of extracellular matrix-related genes (e.g., *MMP1, LAMC2, MMP10* and *LAMB3;* **Table S13)**, several other cell types emerged. They included IC1 *(SERPINB3^high^*) and squamous ICs (IC-sq; *KRT6A^+^ SPRR1B^+^*), which together constituted 53% of cells at ALI-day 3; pre-ciliated (p-C) cells *(CCNO^+^ HES6^+^;* ~7% of cells at this time-point), initial ciliated cells (C1), Ion, and rare tuft-like cells (TLCs) co-expressing markers of Ion, NE and brush, or tuft cells^18,19^ **(Fig. 5E, F** and **Fig. S9)**. At ALI-day 7, proportions of BCs and ICs reduced, while S1-cells *(SCGB1A1^high^*), which emerged at this time-point as a cluster extending further from ICs on the BC differentiation trajectory, constituted ~56% of all cells. Also at day 7, TASCs emerged as an extension of the S-cell cluster further distant from BCs, and the frequency of C-cells, containing C1 and C-s subsets, increased to 5% **(Fig. 5E, F; Table S12)**. Frequencies of C-cells and TASCs increased to 17% and 3%, respectively, by ALI-day 14 and further to 30% and 22% by ALI-day 28 **(Fig. 5F)**. The overall cellular composition of epithelium produced by D-airway BCs after 28 days of differentiation on ALI resembled that observed in D-airways *in vivo,* with major differentiated cell types (C- and S-, including TASCs) constituting together >80% of ASE cells **(Fig. 5E-F; Tables S3 and S12)**.

Remarkable similarity of marker gene profiles of cell types generated by D-airway BCs *in vitro* to those *in vivo,* measured for each ASE cell type as an overlap of DEGs identified for the same ASE cell type *in vitro* and *in vivo* using identical criteria (FDR<0.05; log2 fold-change >0.5 *vs* other ASE cells), was noted. The highest degree of similarity in this regard was observed for C1-cells, whose *in vitro* markers overlapped with 91.3% of DEGs distinguishing these cells from other ASE cells *in vivo,* followed by C-s (56.4%), p-C (48.8%), BC (47.0%), IC (38.9%), TASC (35.9%) and Ion (28%) **(Fig. 5G; Table S14)**. Compared to common secretory (S1) cells, TASCs generated in the same ALI culture displayed significantly higher expression of 84 genes, 40 of which were among those identified in a similar comparison of TASCs and S1 cells *in vivo,* including *SCGB3A2* and *SFTPB* **(Fig. 5H; Table S15)**. Together, these data indicate that distal airway BCs are region-specific epithelial progenitors capable for regenerating TASCs in this anatomical region.

### IFN-γ Inhibits TASC Regeneration by Distal Airway BCs

Based on our *in vitro* data showing the role of distal airway BCs as the cellular origin of region-specific ASE including TASCs, and our *in vivo* scRNA-seq data showing the enrichment of IFN-γ signaling pathway in COPD distal ASE **(Fig. 4E)**, we hypothesized that loss of TASCs in COPD is mediated by augmented IFN-γ signaling in BCs residing in distal airways. Consistent with earlier studies^4,38,39^, the frequency of CD8^+^ T cells, one of the major *IFNG*-expressing cell populations in the human lung **(Fig. 2C and Fig. S10A)**, was significantly greater in distal (pre-T) airways of COPD patients compared to those in normal lung donors **(Fig. 4B)**. Imaging CyTOF analysis mapped CD8^+^ cells to sub- and intraepithelial domains of pre-TBs and TBs in close proximity to BCs and ICs **(Fig. S10B)**, which, based on scRNA-seq data, expressed IFN-γ receptor genes *IFNGR1* and *IFNGR2 in vivo* and *in vitro* **(Fig. S10C)** and, thus, represent potential target cell populations for IFN-γ in distal airways.

We tested this hypothesis by evaluating the differentiation of BCs isolated from distal airways in the presence or absence of IFN-γ added from the basolateral side of ALI culture to enable the access to BCs during at least 10 days between ALI days 0 and 40 **(Fig. S10D)**. In 5 independent experiments, stimulation with IFN-γ induced IFN-γ-response genes *MX1*, *IFIT1* and *CXCL10* in BC-derived epithelia **(Fig. 5I** and **S10D)**, suggesting that BCs can operate as IFN-γ-sensors and responders in human airways. This was accompanied by down-regulation of TASC marker genes *SCGB3A2* and *SFTPB* in epithelia derived from IFN-γ-treated distal airway BCs **(Fig. 5I)**. Although there was a trend of reduced expression of a common S-cell gene *SCGB1A1* in IFN-γ-treated group, there was no consistent change in expression of the mucus-related gene *MUC5B* and the ciliated cell marker *FOXJ1* in response to IFN-γ **(Fig. S10D)**. Thus, by acting on distal airway BCs, IFN-γ inhibits the regeneration of non-mucous S-cells, predominantly TASCs, by these progenitors, inducing the altered epithelial differentiation phenotype observed in pre-T airways of COPD patients.

## Discussion

In this study, using a novel, region-precise method of airway dissection, we assembled an atlas of cellular heterogeneity along the proximal-distal axis of the human respiratory tree. Our study captured 48 cell types and subtypes representing all histological elements along this axis. A novel, anatomy-guided approach to distal airway dissection in our study, which regards secondary lobular bronchioles, pre-TBs, the last macroscopically distinguishable segments along the bronchial pathway^2^, as anatomical landmarks for region-precise isolation of distal airways, allowed us to characterize cellular organization of peripheral bronchioles at the single-cell level. In our approach, pre-TBs served as the boundary between pre-terminal (pre-T) airways and the terminal (T) bronchoalveolar region. While the former included pre-TBs and bronchioles leading to them, the T-region harbored the entire continuum of microscopic anatomical segments, from TBs to alveolar sacs, within secondary pulmonary lobules, known to be the functional and structural lung units, each supplied by an individual pre-TB^2,3^. Such an approach allowed us to systematically characterize, at single-cell resolution, gradual changes in cellular composition occurring along the bronchoalveolar continuum of human lung.

One of the major findings of our study enabled by this region-precise dissection method was the identification of terminal airway-enriched secretory cells (TASCs), a unique population of epithelial cells specific to distal airways that expressed secretoglobin 3A2 *(SCGB3A2)* and/or surfactant protein B *(SFTPB).* As their name implies, TASCs were enriched in terminal airway segments, TBs, and, particularly, in TB-RB junction areas, where conducting airways transition to the respiratory zone^1^. Consistent with their enrichment in this transitional region, TASCs displayed an intermediate molecular pattern that integrated a distinct set of airway secretory and alveolar type 2 cell (AT2) features and gradually changed as airways approached the respiratory zone. In pre-TBs, >40% of TASCs expressed SCGB1A1, a marker of airway secretory (S)-cells, and, similar to the latter, appeared as dome-shaped non-ciliated bronchiolar, or club, cells^40^. Yet, compared to other S-cells, TASCs expressed *SCGB3A2* and *SFTPB* and a set of genes, including *SFTPA2*, *HOPX* and *SFTPA1,* characteristic for AT2 cells^23^. In TB-RB junction areas, however, >90% of TASCs did not express SCGB1A1 protein and were morphologically identical to cuboidal cells of RBs^33,41^. Consistent with this, TASCs in T-airways up-regulated *SFTPA2,* a gene coding for surfactant protein A, known to mark cuboidal cells of transitional epithelium of RBs^33^. TASCs did not typically express *SFTPC,* the obligate marker of AT2 cells. Thus, TASCs comprise a spectrum of epithelial states in the peripheral region of the respiratory tree gradually changing as distal conducting airways transition to the respiratory zone, culminating in cuboidal cells of RBs^33,41^, which remained poorly characterized prior to this study.

A progressive increase in TASC abundance along the proximal-distal axis of the airway tree was paralleled by changes in the distribution of several non-epithelial cell types contributing to region-specific tissue microenvironments. Endothelial capillary cells, including recently described alveolar capillary aerocytes^23^, progressively increased in frequency as airways became smaller and transitioned to the respiratory zone. By contrast, venous endothelial cells, including those with features of the fenestrated endothelium, were most abundant in P-airways, likely representing post-capillary venules of the bronchial vasculature serving as ports of leukocyte entry into the bronchial mucosa or lymph nodes^42^. Glia/Schwann cells were most abundant in P-airways and decreased in frequency along the proximal-distal axis, mirroring the density of cholinergic innervation that regulates bronchoconstriction and mucus secretion from submucosal glands in this anatomical region^20,43,44^.

Our study also identified regional compartmentalization of immune cell niches along the bronchoalveolar axis with reciprocal enrichment of lymphoid and myeloid cells in proximal (P) and distal airway regions, respectively. B-cells and various T-cell subsets, such as central and naïve, resident memory and common CD8^+^-enriched T-cells, were more abundant in P-airways, while cells of the mononuclear phagocyte family were overall more frequent in distal regions. Such heterogeneity may govern region-specific host defense strategies in the respiratory tract, with higher baseline adaptive immune potential, capable of post-infection memory, in P-airways, more frequently exposed to pathogens^45^, and reliance on innate immune potential of monocytes and resident macrophages as a homeostatic defense strategy in distal airways and alveoli^28,46^.

Many key aspects of region-specific cellular organization of distal airways identified in our study showed an altered distribution in distal airways of patients with COPD, a disease, in which progressive airflow obstruction develops due to remodeling of peripheral bronchioles, from pre-TBs to RBs^4–8^, i.e., those captured by our airway dissection method. Although various aspects of this remodeling, including airway wall thickening, lumen narrowing, inflammation and loss of small airways, have been extensively studied using micro-/computed tomography and histologically^4–8^, the cellular basis of these changes remains poorly understood. The region-precise approach to peripheral airway analysis in our study allowed us to systematically identify changes in cellular organization of distal airways in COPD at single-cell resolution.

Among the changes identified in COPD distal airways in our analysis was loss of TASCs, which, as detailed above, represent a unique epithelial cell population of distal airways. In COPD lungs, TASC frequency was reduced in all distal airway segments, from pre-TBs to TB-RB junctions, with particularly dramatic loss in pre-TBs, where TASCs were on average about five times less frequent than in non-diseased lungs. Loss of TASCs in COPD was not paralleled by altered distribution of other airway surface epithelial cells in this anatomical region, including common S- and ciliated cells, known to be affected by cigarette smoking, the major risk factor for COPD^47^. Further, no significant difference in the frequency of TASCs was noted in distal airways of healthy smokers compared to nonsmokers, suggesting that loss of TASCs in COPD represents a disease-associated phenotype rather than merely a response to cigarette smoking.

In our previous study^48^, down-regulation of *SCGB3A2* and *SFTPB,* markers of TASCs, paralleled by acquisition of P-airway-like molecular pattern, was found in samples obtained by bronchoscopy from the 10^th^-12^th^ generation bronchi of COPD subjects and healthy smokers who had a lower forced expiratory volume in one sec/forced vital capacity (FEV1/FVC) ratio, a spirometric measure of airway obstruction, compared to other smokers. Thus, it is possible that molecular changes in more proximal, bronchoscopically accessible small airways may precede and predict loss of TASCs in pre-TBs and TBs (generations 15-16)^49^ as COPD develops. Future studies are needed to determine whether loss of TASCs in pe-TBs and TBs correlates with or precedes loss of these airways known to occur relatively early during disease progression^4–8^.

Loss of TASCs in distal airways in COPD was paralleled by changes in the composition of non-epithelial cell populations contributing to the region-specific microenvironment in these airways, including a dramatic reduction of capillary endothelial cells, particularly those with features of capillary aerocytes^23^, normally enriched along the proximal-distal airway axis. This was reciprocated by an increased frequency of glial/Schwann cells and CD8^+^-enriched T-cell subsets, which in normal lungs were more abundant in proximal airways. Although the role of Schwann cells in COPD is unknown, it is possible that higher abundance of these cells in distal airways reflects increased density of sensory neurons potentially relevant to pathogenesis of chronic cough and mucus hypersecretion in this disease^50,51^. The increased number of CD8^+^ T cells in distal airways of COPD patients has been observed in earlier studies^39^. Consistent with higher expression of *INFG* in these CD8^+^ T cells and earlier findings of augmented cytotoxic IFN-γ-secreting T-cell responses in COPD^38^, IFN-γ signaling was identified in our analysis as the top biological pathway enriched in the distal airway epithelium of COPD patients.

Using an *in vitro* model of airway epithelial differentiation, our study identified distal airway BCs as region-specific progenitors harboring “regional memory”, an intrinsic program that enables these cells to regenerate region-specific epithelium even after their separation from their *in vivo* tissue microenvironment. Consistent with the concept that, as tissue-resident stem cells, airway BCs regenerate epithelium specific to a region of their residence^13,14,36,52^, our study provides evidence for regional memory of BCs in pre-TBs/TBs, the most distal airway segments. BCs isolated from these bronchioles, but not those from proximal airways, were able to generate epithelia with unique cellular composition observed in these distal bronchioles *in vivo*, including the presence of TASCs. This suggests that homeostatic maintenance and repair of the unique epithelial phenotype in pre-TBs/TBs relies on a region-specific regenerative function of resident BCs determined by their regional memory and regulated by their microenvironment.

This led us to hypothesize that loss of TASCs, a region-specific differentiation feature of distal airway epithelium in COPD, is due to altered regenerative function of resident BCs. Earlier studies have shown the altered self-renewal and differentiation of BCs from proximal airways of COPD patients^53^, and BC clones expanded from COPD lungs can form metaplastic lesions in xenograft models^54^. Based on scRNA-seq data indicating increased CD8^+^ T cell-driven IFN-γ signaling in COPD distal airway epithelium and expression of IFN-γ receptor genes in BCs, we reasoned that, as a part of the disease-modified niche, IFN-γ may impact the stem cell function of BCs sensing this cytokine. Indeed, we found that IFN-γ, when included in the BC “niche” during their differentiation *in vitro,* suppressed their ability to regenerate TASCs mimicking the altered differentiation pattern observed in distal airway epithelium of COPD patients *in vivo.* This is the first evidence linking T-cell inflammation to altered distal airway regeneration in COPD. The pathophysiological role of other changes in COPD distal airways identified in our study, such as an increased number of mast cells and reduced frequency of capillary endothelial cells, as elements of the disease-modified distal airway microenvironment, requires further investigation.

Although functional studies are needed to establish the impact of TASC loss in the lungs of COPD patients on disease evolution, our study suggests that loss of these cells may contribute to altered distal airway homeostasis in this disease. As the major source of surfactant protein expression in this transitional region, where AT2 cells are absent and the airflow pattern shifts from bulk flow characteristic for conducting airways to diffusion in the respiratory zone^55^, loss of TASCs may lead to increased surface tension in this region, collapse of peripheral bronchioles during expiration and, thus, contribute to airflow limitation that defines COPD. Further, loss of SCGB3A2, a protein with multiple immunomodulatory and tissue protective properties^56^, for which TASCs serve the key source, may facilitate inflammation and contribute to the altered host defense in distal airways in COPD^57^, making these airways vulnerable to chronic injury.

Suppression of regenerative potential of distal airway BCs due to disease-related changes in their niche may serve as a mechanism of remodeling and loss of distal airways in COPD^4–8^. If so, COPD may represent a disease driven by regenerative failure of distal airways. In support of this concept, previous studies have shown that BCs expanded from small airways or distal lung region of COPD patients fail to establish and maintain mechanically stable, differentiated airway epithelial barrier^58^ but instead produce metaplastic pro-inflammatory lesions^54^. In this regard, the pathologic regenerative pattern in COPD distal airways appears remarkably distinct from that observed in idiopathic pulmonary fibrosis (IPF), a disease, in which fibrosis of lung parenchyma is accompanied by the emergence of ectopic bronchiolar epithelium containing BC-like cells^24,59–61^ and S-cells, including those expressing TASC markers SCGB3A2 and SFTPB^25^. This suggests the possibility that, similar to distal airway progenitors and bronchoalveolar stem cells in murine lungs^36,62–64^, BCs residing in TBs of the human lung, via generation of TASCs, contribute to structural homeostasis in transitional bronchoalveolar segments, RBs, which do not contain BCs or AT2 cells, epithelial progenitors residing in conducting airways and alveoli, respectively^65^. Pathologically augmented repair of these segments by distal airway BCs may lead to fibrosis^66^, while suppression of this regenerative mechanism may compromise structural integrity of this transitional region and lead to centrilobular emphysema characteristic of COPD^6^.

Several key aspects of cellular organization of distal airways characterized in this study, including those affected in COPD, are unique to the human lung. Distal airway BCs, epithelial progenitors in this anatomic region in humans, and RBs, transitional bronchoalveolar segments in the human lung, are minimal or absent in commonly used laboratory animals, including the mouse^35,67,68^. A region-precise approach to study cellular organization and regeneration of human distal airways established in our study will facilitate the discovery and pre-clinical targeting of patient-specific mechanisms of distal airway pathology in COPD and other lung diseases.

## Authors’ contributions

RS: conceived, supervised and supported study

SR, SBM: established the methodology of lung tissue dissection, processing and cell isolation

SR: performed tissue dissection, immunofluorescence staining and analysis, and *in vitro* studies

SR, AFR, RS: performed analysis of immunofluorescence data

AFR: performed analysis of imaging mass cytometry (CyTOF) data

YH: performed analysis of RNA-seq and scRNA-seq data

HR: performed imaging mass cytometry (CyTOF) analysis

OE: supervised analysis of RNA-seq, scRNA-seq and imaging CyTOF data

SHR, BR, VP, JAK, TSB, FDO: provided lung tissue samples

AU: assisted with lung tissue procurement

SHR, VP and TSB: provided histological samples for imaging studies

FJM: assisted with clinical expertise and resources

SR, YH, OE, RS: performed integrated data analysis and interpretation of data

RS: wrote the manuscript

All authors reviewed and edited the final version of the manuscript.

## Acknowledgements

These studies were supported by the National Institutes of Health grants U01 HL145561, R01 HL123544 and R01 HL127393 to RS. We thank personnel of the Marsico Lung Institute Tissue Procurement and Cell Culture Core (supported by Cystic Fibrosis Foundation Grant BOUCHE19R0 and NIH Grant DK065988) for providing lung tissue samples. SBM was supported by the American Lung Association Senior Research Training Fellowship and Weill Cornell Clinical and Translational Science Center TL1 Training Award. AFR is supported by the NCI T32 CA203702 grant. BR was supported by the VA career development grant IK2BX003841. TSB was supported by the VA Merit Review Grant 2 101 BX002378 to TSB.

