## Supplemental Information for "A Unique Cellular Organization of Human Distal Airways and Its Disarray in Chronic Obstructive Pulmonary Disease"

### Methods

*Human Lung Tissue Samples*

Fresh lung tissue samples were obtained from the University of North Carolina (UNC) Tissue and Cell Culture Procurement Core, the Lung Transplant Program at Columbia University Medical Center (CUMC) and the Lung Transplant Center at Vanderbilt University Medical Center (VUMC) following the protocols approved by the UNC, CUMC and VUMC Institutional Review Boards. Normal lung samples represented donor lungs unused for transplant from subjects without a history of prior chronic lung disease, who had been maintained on mechanical ventilation <7 days, which were not eligible for transplant due to physical mismatching or other reasons not related to the presence of lung disease or pathology[1,2]. COPD lung tissue samples were from patients undergoing lung transplantation due to very severe COPD (GOLD stage IV; forced expiratory flow in 1 second [FEV1] <30%)[3,4] or from lung donors with documented history of COPD. Lung tissue samples from 12 normal lung donors and 5 subjects with COPD were used for single-cell RNA-sequencing (scRNA-seq) analysis. Demographic data, lung health information, smoking status and history, causes of death (for lung donors) and source of samples are detailed in **Supplemental Table I**. The mean age of normal lung donors and COPD subjects was 39 years (median: 38 years; range 11-67 years) and 55 years (median: 57 years; range 41-65 years), respectively, with no significant difference between the groups (p=0.246). Both groups included males and females: 4 of 12 (33.3%) of normal lung donors and 3 of 5 (60%) of COPD subjects were females. All five COPD subjects were smokers compared to 5 of 12 (41.7%) in the normal lung group (p=0.044 *vs* COPD group), consistent with nature of COPD is a smoking-related disease[3]. Smoking exposure among the normal lung donor group with known smoking history was on average 20.8 pack-years compared to 37.5 pack-years in COPD group (p=0.16). These data are presented in **Supplemental Table I**.

*Lung Tissue Dissection*

Three anatomical regions, proximal airways (P), distal pre-terminal airways (pre-T) and terminal bronchoalveolar units (T), were dissected from fresh lung tissue samples for scRNA-seq analysis. P-airways were represented by cartilaginous bronchi (generations 1-4) **(Supplemental Fig. 1A)**. Distal airways were identified based on their dimension (luminal diameter <2 mm, i.e., definition of small airways[5]). Air was injected through candidate airways to inflate and locate the associated alveolar region **(Supplemental Fig. 1B)**. Distal airways were dissected by following them toward lung periphery containing associated alveolar region until the last macroscopically distinguishable airway, representing the secondary lobular, or pre-terminal bronchiole (pre-TB)[6] **(Supplemental Fig. 1C)**. Each pre-TB supplies a separate secondary pulmonary lobule (further referred to as "lobule"), a structural unit of lung[6,7] **(Supplemental Fig. 1D)**. In our method, pre-TBs served as anatomical coordinates for annotation of distal airways based on their branching order relative to a given pre-TB (i.e., pre-TB -1, an airway one generation proximal to pre-TB)[8,9] and for locating proximal borders of lobules **(Supplemental Fig. 1E)**. Within the lobule, pre-TB gives rise to TBs, which at their distal ends open to respiratory bronchioles (RBs) that lead to alveolar region[6,7,10]. In our method, pre-TBs were regarded as a boundary between pre-T-airways (pre-TB and 3-4 generations of bronchioles proximal to pre-TB) and associated T-region (TBs and respiratory segments, from RBs to alveoli, belonging to lobule supplied by the same pre-TB; **Supplemental Fig. 1E**). Airway-containing portions of T-regions, representing centriacinar or centrilobular domains[7], were excised from surrounding tissue by pulling them out *en bloc* with associated pre-T-airways and then separated from the latter by cutting immediately below the last macroscopically visible part of pre-TB **(Supplemental Fig. 1E)**. Selected dissected samples were evaluated under dissection microscope **(Supplemental Fig. 1E$_2$)** and by hematoxylin-eosin staining **(Supplemental Fig. 1F)** to evaluate their anatomical and histological organization.

To enable intra-individual analysis of region-specific cellular heterogeneity, all 3 regions (P, pre-T and T) were isolated from normal lung samples of 5 donors. To enable comprehensive analysis of distal airways, pre-T airways were dissected from 6 additional normal donor lungs, from 2 of which matched T-regions were also isolated. For one normal donor lung, 2 pre-T samples were included to assess intra-subject variability. To enable the analysis of disease-related changes in cellular composition of distal airways in COPD, pre-T-airways were dissected from lungs of 5 subjects with COPD. All these samples are described in **Supplemental Table I**.

*Tissue Dissociation for scRNA-seq Analysis*

Dissected tissues were minced into small pieces (< 3 mm) and digested in DMEM/F-12 (Thermo Fisher Scientific) containing 2 mg/ml collagenase I (Millipore), 1 mg/ml dispase II (Sigma Aldrich) and 0.3 mg/ml DNase I (Stemcell Technologies) at 37°C with rotation for 2 hrs. Digested samples were filtered through a 100-µm cell strainer (Corning), washed in DMEM/F-12 and pelleted by centrifugation (170g, 5 min, 4°C). Red blood cell lysis was performed using RBC lysis solution (Miltenyi Biotec) according to the manufacturer's instructions, after which cells were washed again with DMEM/F-12 and cell count was performed.

*scRNA-seq Library Preparation and Sequencing*

Single-cell suspensions generated from fresh lung tissue samples used for scRNA-seq analysis **(Supplemental Table I)** showed viability >70% and cilia beating detectable under the microscope. Single cell libraries were prepared using 10x Genomics Single Cell 3' v2 Reagent Kits following the manufacturer's instructions targeting 10,000 cells per sample. Libraries were pooled and sequenced on Illumina HiSeq 2500, HiSeq 4000 or Novaseq 6000 instruments. Reads were aligned to GRCh38 (GENCODE v32/Ensembl 98) human reference genome using Cell Ranger (version 6.03.0, 10x Genomics). Library preparation, sequencing and initial processing of sequencing data was performed at the Weill Cornell Epigenomics Core Facility.

*Dimensionality Reduction, Cell Clustering, and Doublet Removal*

Output files for individual samples from Cell Ranger Count were read into Seurat v4 [ref11] to generate a unique molecular identifier (UMI) count matrix that was used to create a Seurat object containing a count matrix and analysis. All Seurat objects were combined into a merged dataset, and a percentage of mitochondrial genes were calculated for each sample in the merged object. Raw expression data from 30 fresh lung tissue samples contained a total of 147,381 cells. Cells with fewer than 200 genes or 1,000 UMIs identified or more than 25% of reads mapped to mitochondrial genes were excluded. Gene expression counts were normalized, log-transformed and scaled by SCTransform using Seurat v4. Principal component analysis was performed on the top 2000 most variable genes, and the first 100 principal components were used for Uniform Manifold Approximation and Projection (UMAP) clustering. A shared nearest neighbor graph was constructed based on UMAP using 'FindNeighbors' function. Cell clusters were detected using the Louvain clustering algorithm. Different clustering resolutions were used to segregate cell groups representing cell types and subtypes identified based on markers as described below. 'SubsetData' function and iteratively 'FindClusters' function were applied to major cell types until no differentially expressed gene (DEG) could be identified between subclusters. Once clustering was completed, 4,388 possible doublets were identified by DoubletFinder[12] and removed. A total of 111,412 high quality singlets remained after quality control and filtering steps described above were used for further analysis. This included a total of 20,333 single cells representing P-airways (4,067 cells per sample; range 1,606-5,672) of normal donor lungs; 66,498 single cells from pre-T (3,912 per sample; range 1,071-7,902), including 48,089 cells from pre-T airways of normal donors lungs (4,007 cells per sample) and 18,409 cells from pre-T airways of COPD subjects (3,690.5 cells per sample), and 24,581 single cells from T-airways of normal donors lungs (3,512 cells per sample; range 772-6,038 cells) as described in **Supplemental Table I**.

*Cell Group Annotation*

To identify markers of individual cell groups, we performed differential expression analysis between each cluster against all other clusters, using the 'MAST' statistical framework[13] implemented in the Seurat's 'FindMarkers' function, adjusted by cellular detection rate. Genes with Bonferroni-corrected q-value < 0.05 were considered as DEGs **(Supplemental Table II)**. Using these DEGs, individual cell types and cell subtypes were identified as described below. Lower resolutions (res) of UMAP clustering, starting with res 0.0001, were used to identify cell families, e.g., epithelial (*KRT19*$^+$), stromal (*COL3A1*$^+$), endothelial (*PECAM1*$^+$), leukocytes / immune (*PTPRC*$^+$) (**Supplemental Fig. 2A$_1$**). Resolutions were gradually increased to 5.0 to determine clustering patterns (**Supplemental Fig. 2A$_2$-A$_3$**). DEGs identified for clusters at res 0.8-5.0 were used to annotate cell types and subtypes. Initial labeling was performed based on clustering and identified markers generated at res 0.8 as the baseline annotation with subsequent identification of potential cell subtypes or subsets at res 1.0-5.0. Each cluster representing a cell type or subtype was inspected to identify topological outliers, i.e., cells located outside the main clustering area and spatially overlapped with a distant, i.e., non-adjacent cell type/cluster. To ensure high specificity of cell annotation, these outlier cells labeled as "Unknown and ambiguous cells" (Un) and excluded from the subsequent analysis. Cell types were examined at res 1.0-5.0 to determine potential subtypes (cell states or subsets), their topological boundaries and markers. Expression of identified cell subtype markers was inspected in adjacent clusters representing the same family to locate and label cells representing a given cell subtype within these clusters.

*Epithelial cells.* Majority of epithelial cells were found in cluster 1 at res 0.0001 (**Supplemental Fig. 2A$_1$**). Basal cells (BCs), stem/progenitor cells of airway surface epithelium (ASE)[14,15], were identified in cluster 3 at res 0.8 (**Supplemental Fig. 2A$_3$**) by expression of *KRT5*, a canonical BC marker[16,17], and other BC markers identified in recent scRNA-seq studies, including *KRT15*, *MIR205HG*, *DLK2* and *F3*[18–22] **(Supplemental Fig. 2B)**. Intermediate cells (ICs), also known as indeterminate, para-BCs, supra-BCs, differentiating or secretory-primed BCs[18,23–27], were found in clusters 26 and 39 at res 2.0 between BCs and secretory (S) cells **(Supplemental Fig. 2C$_1$)**. ICs expressed a number of BC markers **(Supplemental Table II)**, including *KRT5*, but compared to BCs, had lower expression of *KRT15* and other BC-specific genes **(Supplemental Fig. 2B)** but higher expression of *SERPINB3* and *KRT7*, differentiating BC markers[18], also expressed by S-cells **(Supplemental Fig. 2C$_1$)**. Further in line with their intermediate nature, as previously described[27,28], a subset of ICs expressed *SCGB1A1*, a marker of S-cells, but at a lower level, compared to the latter, as well as *ALDH3A1*, which was also detected in ciliated (C) and S-cells **(Supplemental Fig. 2C$_1$)**. A subset of ICs expressed markers of squamous differentiation, including *KRT6A* and *SPRR1B*[ref29–31], and cell proliferation-related genes, such as *MKI67* and *UBE2C* **(Supplemental Fig. 2C$_2$)**. This is in agreement with earlier studies showing that ICs (para-BCs) are a source of proliferating and squamous cells in the human ASE[21,24,29,30]. These proliferating ICs represent recently described proliferating and "cycling" BCs[18,32]. BCs and ICs showed high expression of *S100A2* and *KRT17* **(Supplemental Fig. 2C$_3$)**, which together with *KRT5* represent common markers of undifferentiated ASE cells (BCs and ICs combined; cluster 6 at res 0.1; **Supplemental Fig. 2A$_2$**).

Secretory (S)-cells, identified based on expression of *SCGB1A1* and *SCGB3A1*, S-cell markers[18,32–36], were found in clusters 10, 29, 36, 38 and 49 at res 0.8, adjacent to ICs, further distant from BCs **(Supplemental Fig. 2D)**. A subset of S-cells (clusters 86, 103 and 111 at res 2.0) exhibited high expression of mucous (goblet) cell markers, including *MUC5AC*, *MUC5B, TFF3,* and *VMO1*, and were annotated as S-mucous (S-Muc) cells **(Supplemental Fig. 2D)**. Another subset of S-cells enriched in clusters 77 and 122 at res 2.0 expressed *SCGB3A2* and

*SFTPB*, genes identified in our previous study as distal airway epithelial signature genes[37]. Consistent with that study, these cells almost exclusively represented pre-T and T-regions, with significant enrichment in the latter **(Supplemental Table III)**, and, therefore, were annotated as "terminal airway-enriched secretory cells" (TASCs). Based on their transcriptomes, TASCs formed a "bridge" between ASE (S-cells) and alveolar epithelium (AT2 cells) **(Supplemental Fig. 2E$_{1\text{-}2}$)**. The latter were identified in clusters 4, 18 and 42 at res 0.8 by expression of *SFTPC* and other well-established AT2 cell markers, including *SFTPD* and *NAPSA* **(Supplemental Fig. 2F$_1$)**. By contrast to AT2 cells, 93% of TASCs expressed *SCGB1A1*, a marker of airway S-cells, <20% of TASCs were *SFTPC*$^+$ and <10% of TASCs were *SFTPD*$^+$ or *NAPSA*$^+$ **(Supplemental Fig. 2E$_1$ and 2F$_1$)**. Apart from *SCGB3A2* and *SFTPB*, TASCs expressed a number of genes also detected in AT2 cells, such as *RNASE1*, *CYB5A, HOPX,* and *SFTPA2* **(Supplemental Fig. 2E$_2$)**. Only <2% of TASCs expressed *AGER*, a marker of AT1 cells[18,20]. AT cells clustered near AT2 cells (cluster 74 at res 0.8) and showed high expression of *CLDN18* and *EMP2* **(Supplemental Fig. 2F$_2$)**. Since TASCs shared a number of common features with other S- and AT2 cells, the latter were inspected to find cells expressing *SCGB3A2* or *SFTPB* (expression level >1 for each) among remaining S-cells and *SCGB3A2* and *SCGB1A1* (expression level >1 for both) among *SFTPC*-negative cells within AT2 cell cluster were annotated as TASCs. After labeling S-Muc and TASCs as described above, the remaining S-cells were labeled as common S (S1) cells.

Several groups of cells expressing ciliated (C) cell markers, including *CAPS*, *FOXJ1*, *TPPP3* and *TMEM190*[ref18,21,38], were identified (enriched within cluster 1 at res 0.01), together representing further extension of the BC-IC-S "differentiation" trajectory within the ASE family (cluster 0 at res 0.001) **(Supplemental Fig. 2A$_2$ and 2G$_1$)**. Topologically, these groups of cells differed from each other based on their proximity to S-cell clusters, the most distant cells (cluster 0 at res 0.1) marked almost exclusively by C-cell genes, and intermediate groups of cells, located between S- and C-cells (clusters 16 and 18 at res 0.1), which expressed both S- and C-cell genes. The first group included two subsets of C-cells, i.e., common C-cells (C1), expressing typical C-cell markers (clusters 0, 7, 34, 50, 51 at res 0.8) and those, expressing in addition to these C-cell genes a set of genes associated with serous S-cell phenotype, including *SAA*-family genes, such as *SAA1*, *SAA2* and others (designated serous secretory-like ciliated cells, C-s cells; clusters 8, 52 and 56 at res 0.8) **(Supplemental Fig. 2$G_1$ and 2$G_2$)**. The second group, designated "secretory-ciliated intermediate" (CSI) cells included 2 subgroups of cells: 1) those expressing markers of recently described deuterosomal precursors of ciliated cells[21,39–41] *CCNO, HES6* and *CDC20B*, designated by us as pre-ciliated (p-C) cells **(Supplemental Fig. 2$G_3$)** enriched in cluster 61 at res 2.0, and 2) cells not expressing these markers, but exhibiting high expression of S-, including S-Muc-related genes **(Supplemental Fig. 2$G_4$)**. The latter (clusters 58 and 59 at res 0.8) resembled recently described mucous-C-hybrid cells[32], and were designated as "S-Muc-C hybrid" (H) cells.

Clusters representing rare ASE cell types, ionocytes (Ion) and neuroendocrine (NE) cells, were identified already at the lowest resolutions (clusters 14 and 13, respectively, at res 0.0001). Ion (cluster 77 at res 0.8) were identified based on expression of *ATP6V0B*, *FOXI1*, *TMEM61*, *CFTR* **(Supplemental Fig. 2H)** and other markers described for this recently characterized cell type[18,32,42,43]. NE cells (cluster 75 at res 0.8) were found based on expression of *CHGA*, *GRP*, *CALCA, SEC11C* **(Supplemental Fig. 2I)** and other known NE marker genes[18,20,42,43].

All major cell types of airway submucosal glands (SMG) were identified. They included myoepithelial (ME) cells and two subtypes of glandular S-cells, i.e., serous (G-Ser) and mucous (G-Muc) cells. ME cells are found in cluster 15 at res 0.0001 (corresponding to cluster 78 at res 0.8) based on co-expression of the common BC marker *KRT5,* BC subpopulation marker *KRT14* and smooth muscle (SM) cell-associated actin *ACTA2* **(Supplemental Fig. 2$J_1$)**, known to be co-expressed in these cells[44,45]. Due to this dual gene expression profile, ME cells clustered distant from other epithelial cells. Glandular S-cells, marked by expression of *AZGP1* **(Supplemental Fig. 2J$_2$)**, as previously noted[32], clustered (cluster 9 at res 0.001) separately from, but adjacent to ASE cell supercluster (cluster 0 res 0.001). SMG mucous (G-Muc) cells (cluster 155 at res 3.0) were detected by expression of *MUC5B*, a known marker of these cells[2], and *BPIFB2*[ref46] **(Supplemental Fig. 2J$_2$)**, and distinguished from ASE mucous (S-Muc) cells by the absence of *MUC5AC* expression **(Supplemental Fig. 2D)**. G-Ser cells (cluster 57 res 0.8) were detected by expression of *LYZ*, *LTF* and *PRR4* **(Supplemental Fig. 2J$_2$)** known to mark these cells[18,21,32,47]. Some G-Ser cells expressed *SCGB3A2*, but, compared to TASCs, they did not express *SFTPB* and had low/no expression of the ASE S-cell marker *SCGB1A1* **(Supplemental Fig. 2E$_1$)**.

*Stromal cells.* Majority of stromal cells were found in cluster 3 at res 0.0001, except for chondrocytes (cartilage cells; Cr; cluster 9 at res 0.0001; cluster 66 at res 0.8) identified by expression of *SNORC*, *COL2A1* and ACAN, Cr-specific genes[ref48,49] **(Supplemental Fig. 2K)**. The remaining stromal cells at res 0.01 formed 2 groups, corresponding to 2 major stromal cell types, fibroblasts (Fb; cluster 5) and smooth muscle (SM) cells (cluster 6) **(Supplemental Fig. 2A$_2$)**. Fbs were identified based on high expression of *FBLN1* and *DCN* **(Supplemental Fig. 2L$_1$)** consistent with previous reports[21] and included at least 4 subsets (Fb1-Fb4), defined based on markers of Fb clusters at res 0.8 (subclusters of the initial Fb cluster identified at res 0.01). One of the Fb subsets (Fb1; cluster 13 res 0.08) expressed Fb markers **(Supplemental Table II)** shared by other subsets and, therefore, was named "common Fb subset". The remaining Fb subsets exhibited some features distinguishing them from other Fbs. Fb2 subset (cluster 57 at res 1.0 and partially in cluster 155 at res 5.0) displayed high expression of *CCL19*, previously shown to be expressed in perivascular Fbs[50], and *SFRP2*, a recently described marker of lung adventitial Fbs[18] **(Supplemental Fig. 2L$_2$)**. A third Fb subset (Fb3; cluster 65 at res 0.08, and clusters 82 and 98 at res 2.0) showed high expression of extracellular matrix (ECM)-related genes, including *ELN*, *COL1A1*, *MMP2* and others **(Supplemental Table II)** and *ACTA2*, a marker of SM cells **(Supplemental Fig. 2L$_3$)**. For this reason, F3 subset was designated "matrix Fbs", which included myofibroblasts (myoFbs). The fourth Fb subset (Fb4; cluster 155 at res 5.0) was marked by expression of *GPC3*, *FGFR4*, *VEGFD* and *WNT2* **(Supplemental Fig. 2L$_4$)** recently shown to be markers of alveolar Fbs[18,51].

Three SM subsets (SM1-SM3) were identified. SM1 (clusters 23 and 35 at res 0.8) and SM2 (cluster 45 at res 0.8) shared a number of marker genes, which were particularly highly expressed in pericytes (Pr, described below), including *NOTCH3* and *PDGFRB* **(Supplemental Fig. 2M$_1$)** previously shown to mark vascular SM cells[18]. Compared to other SM cells, SM2 subset showed particularly high expression of *ADIRF* **(Supplemental Fig. 2M$_1$)**, which was earlier found among markers of both airway and vascular SM cells[18], suggesting that this subset may represent a biological state of both SM subtypes. The third SM subset (SM3) displayed high expression of *DES*, *TPP2*, *CNN1* and other airway SM markers[18,21], as well as *BCHE* and *PRUNE* **(Supplemental Fig. 2M$_3$)**, markers of recently described fibromyocytes[18]. Pericytes (Pr) clustered (cluster 64 at res 0.8) near SM cells and were identified based on high expression of *COX4I2*, *KCNK3*, *PDGFRB* **(Supplemental Fig. 2M$_4$)** and other markers previously shown to mark this cell type[18,20,21].

Glial (Schwann) cells (cluster 11 at res 0.0001, corresponding to cluster 73 at res 0.8) were identified based on expression of *CDH19, GPM6B, MPZ, NRXN1* **(Supplemental Fig. 2N)** and other genes known to be expressed in these cells[52–55]. Whereas this cell type has not been captured in previous lung scRNA-seq studies, analysis of scRNA-seq data sets available in the Azimuth[56] confirmed the identity of this cell population as "Schwann cells" based on similarity to Schwann cells identified in the human pancreas (Azimuth data is detailed separately below).

*Endothelial cells.* Endothelial (En; *PECAM1*$^+$ *CLDN5*$^+$ *CDH5*$^+$*)* cells were identified in cluster 2 at res 0.0001 (major group of En-cells), except for lymphatic cells (En-l; defined below), which formed a separate cluster (cluster 8 at res 0.0001) **(Supplemental Fig. 2A$_1$, 2A$_3$ and 2O$_1$)**. The following En-cell subsets were identified. Arterial En (En-a) cells (enriched in cluster 186 at res 4.0 and partially dispersed in clusters representing capillary En-cells; described below) were identified based on expression of *IGFBP3, GJA5* and *DKK2* **(Supplemental Fig. 2O$_2$)**, known markers of En-a cells confirmed in recent scRNA-seq studies[18,57]. Capillary En (En-c) cells, identified based on known lung En-c markers[18,57] including *FCN3* and *CA4* **(Supplemental Fig. 2O$_3$)** included 2 subsets, which were clustered separately from each other at res 0.1 (clusters 7 and 24) **(Supplemental Fig. 2A$_2$)** and at later resolutions. One of these subsets (En-c1; cluster 7 at res 0.1, corresponding to clusters 15 and 48 at res 0.08) expressed common En-c markers and, therefore, was designated as "common En-c" subset. The second cluster (En-ca; cluster 24 at res 0.1, corresponding to clusters 33, 46 and 61 at res 0.8, and enriched in cluster 100 at res 2.0) included cells exhibiting high expression of markers of recently described pulmonary capillary aerocytes[18], including *HPGD*, *EDNRB* and *IL1RL1* **(Supplemental Fig. 2O$_4$)**. Venous En (En-v) cells (cluster 4 at res 0.1, corresponding to clusters 12, 24, 40 and 53 at res 0.8) included cells expressing markers known to be expressed in En-v cells and in the fenestrated endothelium[18,57], including *ACKR1*, *PLVAP*, *SELE* and *POSTN* **(Supplemental Fig. 2O$_5$)**. A subset of En-cells, which co-expressed markers of En- and SM cells and formed a cluster (cluster 68 at res 0.8) within the major En-c group oriented toward Pr and SM clusters **(Supplemental Fig. 2O$_6$)**. Lymphatic En (En-l) cells (clusters 60 and 76 at res 0.8) were identified based on expression of *CCL21*, *MMRN1*, *LYVE1*, *PROX1* **(Supplemental Fig. 2O$_7$)** and *TFF3* **(Supplemental Fig. 2D)**, known markers of En-l cells[18,32,57].

*Immune cells.* Different groups of myeloid cells (Im-m), identified based on markers described below, were found in cluster 3 at res 0.001, except for neutrophils (Neu) and mast cells (MC), which clustered separately already at very low resolutions (clusters 5 and 6 at res 0.0001; clusters 7 and 8 at res 0.001) **(Supplemental Fig. 2A$_{1\text{-}3}$ and 2P-Q)**. Neu (clusters 6 and 39 at res 0.8) were identified based on high expression of *CSF3R*, *S100A8*, *FCGR3B*, *IFITM2* **(Supplemental Fig. 2P)** and others shown to mark this cell type in previous scRNA-seq studies[18,32]. The overall gene expression profile of Neu was remarkably similar to cells described in recent scRNA-seq studies as a subset of monocytes (see Azimuth analysis data described below). MCs (cluster 5 at res 0.8) were identified based on expression of the classical MC markers *TPSB2*, *TPSAB1*, *CPA3* and *KIT* **(Supplemental Fig. 2Q)** consistent with previous studies[18,32].

Cells of the mononuclear phagocyte system, including monocytes (Mon), macrophages (M) and dendritic cells (DCs), were found in the cluster 3 at res 0.001. Monocytes (Mon; cluster 5 at res 0.1; corresponding to clusters 2, 25 and 32 at res 0.8) were identified based on expression of *VCAN* and *FCN1*, found to mark these cells in previous scRNA-seq studies[18,58], and *CD163*, which distinguished Mon and other mononuclear phagocytes from Neu **(Supplemental Fig. 2R)**. *IL1B* and *EREG* were among the top Mon markers **(Supplemental Fig. 2R)**. Macrophages (M) were identified based on high expression of the classical M-markers *CD68*, *CD163* and *MARCO* **(Supplemental Fig. 2S$_1$)** and included 3 subsets. The M1 subset (cluster 47 res 0.8) showed high expression of *SPP1* and genes known to mark M1-polarized, inflammatory, classically activated macrophages[59,60], including *CCL3*, *CXCL9* and *CXCL10* **(Supplemental Fig. 2S$_2$)**, and clustered near Mon. The M2-subset (clusters 22 and 70 at res 0.8; enriched in clusters 105, 112 and 114 at res 2.0) showed high expression of *APOE*, *CCL18*, *MSR1*, *MRC1*, and *FABP4* **(Supplemental Fig. 2S$_3$)** known to be expressed in tissue-resident, non-inflammatory, or alternatively activated macrophages[38,59,60], as well as higher expression of common M-genes **(Supplemental Fig. 2S$_1$)**.

The third M-subset (M1-2) was clustered between M1 and M2 subsets (cluster 19 at res 0.8) and showed expression of markers of both subsets **(Supplemental Fig. 2$S_2$ and 2$S_3$)**.

Two types of DCs were identified, conventional (myeloid) (c-DC) and plasmacytoid (p-DCs), which clustered next to each other within the mononuclear phagocyte family. Compared to other mononuclear phagocytes, both DC types showed high expression of *CCR7* **(Supplemental Fig. 2$T_1$)**. c-DCs (cluster 31 at res 0.8) were distinguished by particularly high expression of the HLA family genes, e.g., *HLA-DPB1* and others **(Supplemental Table II)**, and contained cells with high expression of known DC markers *FCER1A*, *CD86*, *CD1C* and *CD1E* **(Supplemental Fig. 2$T_1$)** consistent with previous scRNA-seq studies[18,32,58]. p-DCs (cluster 79 at res 0.8) were distinguished based on expression of *IL3A*, *IRF4*, *IRF7* and *GZMB* **(Supplemental Fig. 2$T_2$)** known to mark this cell type[18,58].

Lymphoid (Im-l) cells included a heterogeneous family of T cells (cluster 1 at res 0.001), B cells and plasma cells (PCs), which clustered separately from T cells and each other already at lower resolutions (clusters 4 and 10, respectively, res 0.0001) **(Supplemental Fig. 2$A_1$)**. B cells (clusters 16, 41 and 55 at res 0.8) were distinguished by high expression of well-known markers of these cells, including *MS4A1*, *BANK1*, *CD19* and *CD79A* **(Supplemental Fig. 2$U_1$)**. The latter two genes were also detected in PCs (cluster 67 at res 0.8), which were identified based on high expression of PC markers, such as *JCHAIN*, *MZB1*, *IGHA1* and *IGKC* **(Supplemental Fig. 2$U_2$)** consistent with previous scRNA-seq studies[18,32].

T cell supercluster included cells expressing T cell markers *CD3D* and *CD3E* on one side of the supercluster, *NK* cell markers *GNLY* and *NKG7* on the opposite side of the supercluster, as well as intermediate cells expressing both markers (labeled as “T-NK” cells) clustered between the two **(Supplemental Fig. 2$V_1$)**. At the T-cell “end” of this supercluster were identified cells expressing markers of central memory and/or naïve T cells (Tcn; enriched in cluster 3 at res 0.1, corresponding to clusters 1, 9, 11, 28, 37, 69 at res 0.8), including *LEF1*, *CCR7*, *SELL* and *IL7R* **(Supplemental Fig. 2V$_2$)**, shown mark these subsets of T cells in previous studies[32,61–63]. Next to Tcn cells was identified a cluster of T cells (cluster 29 at res 0.1, corresponding to cluster 72 at res 0.8) expressing interferon (IFN)-response genes, including *IFIT1*, *IFIT2*, *OASL*, *ISG15* and *HERC5* **(Supplemental Fig. 2V$_3$)**. Similar to previously described activated T cells expressing "IFN response" genes[63], we designated this T cell subset as IFN-responsive T cells (T-ifn).

Cells expressing *CD8A* and *CD8B*, markers of cytotoxic T cells, were enriched in clusters located between Tcn and NK cells (clusters 1 and 15 at res 0.1, corresponding to clusters 17, 26 and 43 at res 0.8, including clusters 91 and 92 at res 2.0) **(Supplemental Fig. 2V$_4$)**. Collectively, cells in this group of clusters were labeled as CD8$^+$ T-cell enriched. A subset of cells within this group (cluster 92 at res 2.0) showed high expression of genes shown to mark T resident memory cells (Trm)[32,61–63], such as *PDCD1*, *ITGA1*, *CXCR6* and *CD101* **(Supplemental Fig. 2V$_4$)**. After labeling cells in this cluster as Trm cells, the remaining cells in the CD8$^+$ T-cell enriched group were labeled as a common CD8$^+$ T-cell enriched subtype (CD8-T1). NK cells (cluster 14 at res 0.8) were found, as mentioned above, at the opposite, relative to Tcn, end of T cell supercluster based on high expression of *GNLY* and *NKG7* **(Supplemental Fig. 2V$_1$)** as well as *GZMB*, *KLRD1* and *PRF* **(Supplemental Fig. 2V$_5$)**, known markers of NK cells[64] or recently described terminally differentiated effector T cells[63]. These cells were annotated as NK cells. Clusters of cells located between T cells and NK cells included cells co-expressing markers of T cells and intermediate levels of NK cells (clusters 21 and 30 at res 0.8) and were designated as "T-NK cells" **(Supplemental Fig. 2V$_5$)**. We found that cells expressing *CCL5* and *IFNG* were enriched in Trm, T-NK and NK cell clusters **(Supplemental Fig. 2V$_5$)**. Complete lists of markers of each of the identified cell type and subtype are provided in **Supplemental Table II**. Distribution of each cell type and subtype in different airway regions is described in **Supplemental Table III**.

*Determination of Consistency of Cell Type Annotations with Reference Data Sets*

To establish the consistency of cell types and cell subtype annotations in our scRNA-seq analysis to those captured in previous scRNA-seq studies, we compared cell type/subtype annotations in our analysis to those in publicly available scRNA-seq datasets in Azimuth[56] (https://azimuth.hubmapconsortium.org/references/), and the scRNA-seq human lung dataset of Travaglini et al[18]. We used three different analytical tools to cross-validate our cell type/subtype annotations. First, we re-annotated cells based on cell type markers in Azimuth datasets using the SCINA algorithm[65]. Second, we used Enrich[66] and cell type annotations and markers in Azimuth datasets to determine the enrichment score and odds ratio of enrichment of Azimuth cell annotation categories in our scRNA-seq data for each cell type and subtype to evaluate the degree of similarity of our annotations to those in Azimuth datasets. Categories with enrichment FDR<0.05 and odds ratio >100 were considered significantly enriched. Third, we used SingleR[67] to annotate cells in our dataset using the dataset of Travaglini et al[18] as reference.

Epithelial cell types and subtypes identified in our analysis were remarkably similar to those in the human lung reference data set[18] **(Supplemental Fig. 2W$_{1,2}$)**. Using this data set as a reference, 96.2% of BCs in our study were labeled as “basal” (BCs) or proximal BCs. Consistent with their intermediate nature, ICs annotated as such in our study were similar to differentiating BCs (34.9%), BCs and proximal BCs (cumulatively 38%), proliferating BCs (11.8%), and goblet cells (12.4%) in this reference data set. Majority (78.6%) of S1 cells in our study overlapped with goblet (68.1%) or mucous (10.5%) cells in the reference data set, and >85% of S-Muc cells were similar to goblet cells. Since TASCs were not described as a separate S-cell category in the reference study, they overlapped, as expected, with common S- (mucous, 69%; and club, 21.5%) cells in the reference data set. H and p-C cells, consistent with their S-C intermediate nature, overlapped with C- (>70%) and goblet cells in the lung reference data set. More than 90% of C-cells (both C1 and C-s subsets) in our study were similar to C-cells in the lung reference data set. Likewise, majority of Ion (96.2%) and NE cells (88.8%) overlapped with their corresponding counterparts in the lung reference study. Consistent with their dual nature, ME cells identified in our study resembled airway SM (34.1%), BCs (31.7%) and fibromyocytes (29.3%) in the lung reference data set. The two SMG secretory cell types, G-Muc and G-ser, exhibited similarity to corresponding S-cell subsets in the lung reference study, such as mucous and goblet cells (collectively, 73.4% of G-Muc) and serous (74.7% of G-Ser). The AT cell types, AT1 and AT2 captured in our study, also showed similarity (>80% overlap for AT1 cells and >90% of AT2 cells) to corresponding cell types in the lung reference data set. Similarity of transcriptional profiles of described epithelial cell types and subtypes in our study to those in the lung reference data sets as described above was confirmed statistically **(Supplemental Fig. 2X$_1$)**.

Among stromal cells, different degrees of similarity to cell types and subtypes in the lung reference data set were observed **(Supplemental Fig. 2W$_3$)**. Since Cr (chondrocytes) were not captured in previous reference studies, only their similarity to described stromal cell types, including Fbs, (adventitial Fbs and myofibroblasts) could be detected. Fb1, Fb2 and Fb3 exhibited similarity to adventitial Fbs (>80%) in the lung reference data set. Notably, 7% of Fb3 in our study resembled myofibroblasts in the lung reference data set, consistent with our annotation as myofibroblast-enriched Fb subset. Also, in line with our annotation, 58.2% of Fb4 (alveolar Fb-enriched) in our study showed similarity to alveolar fibroblasts in the lung reference data set. Likewise, SM cells in our study were annotated in general consistent with the lung reference data set. SM1 subset resembled vascular (57%) and airway (20%) SM, whereas 87.8% of SM2 cells were similar to airway SM. The latter distinction was not that remarkable in our study. The vast majority (93.1%) of SM3 cells in our study, as estimated based on enriched marker genes, were similar to fibromyocytes in the lung reference study. Likewise, 78.6% of Pr in our study showed similarity to cells representing the same cell type in the lung reference data set. Gli (Gli-like / Schwann) cells identified in our study were not described in previous lung scRNA-seq studies, including the Azimuth lung reference data set[18]. Consistent with their annotation as Gli-cells, comparison to reference data sets representing other organs and tissues, available in Azimuth, determined similarity of these cells to both neurons (Vip+ GABAergic neuron 2, 24.4%) and Schwann cells (21.3%) in the pancreas data set **(Supplemental Fig. 2W$_3$)**.

Among endothelial (En) cells **(Supplemental Fig. 2W$_4$)**, lymphatic (En-l) and arterial (En-a) cell subtypes had the highest degree of similarity (99.4% and 85.7%, respectively) to the corresponding cell groups in the lung reference data set. The two subsets of capillary (En-c1 and En-ca) cells identified in our study, consistent with showed the highest degree of similarity to capillary En-cells in the lung reference data set. Whereas 45% of En-c1 cells showed similarity to capillary intermediate cells (45%), 27.7% of cells in this group resembled bronchial vessel cells, and 17.2% arterial En-cells in the lung reference data set, suggesting that, consistent with their annotation in our study as "common En-capillary" cells, they may be shared with different types of vasculature in different regions of the lung. 70.2% of En-ca cells in our study resembled capillary cells, and 13.6% of them were similar to capillary aerocytes in the lung reference data set, consistent with their annotation as capillary aerocyte-enriched subset of En-capillary cells. Notably, majority of En-v and En-SM cells in our study (89% and 76.9%, respectively) showed similarity to bronchial vessel cells described in the lung reference data set. This is consistent with enrichment of En-v cells in the proximal airways observed in our study **(Supplemental Table III)**. Consistent with their annotation as "venous" En-cells, comparison to cell types in all organs and tissues available in Azimuth, determined "vein" cell as the most enriched reference cell type annotation for these cells. Enrichment of the reference cell type annotations for stromal and En-cell groups described above was evaluated statistically by Enrichr **(Supplemental Figure 2X$_2$)**.

Among immune cell types **(Supplemental Fig. 2W$_{5,6}$)**, mast cells (MCs) showed the highest degree of annotation consistency (cons): 98.2% of MCs in our study were annotated as "basophil/mast" cells based on the Azimuth lung reference data set), followed by pDCs (cons 95.9%), PCs (cons 92.8%), Mon (cons 77.1%), NK (cons 76.4%) and B cells (cons 72.6%). Notably, 88.9% cells labeled as Neu in our study were annotated as "OLR1+ classical monocytes" in the lung reference data set. In our study, these cells clustered separately from Mon and from mononuclear phagocyte family, and shared a number of classical Neu markers (described in the section above, including *CSF3R*, *FCGR3B*, *S100A8*, *S100A9* and *IFITM2*) identified as top Neu markers in the study that formed the lung reference data set[18]. Therefore, we kept the Neu annotation label for this cluster of cells. Consistent with their clustering adjacent to Mon and pro-inflammatory profile, M1 macrophages showed similarity to monocytes of the lung reference data set. By contrast, the majority (83.3%) of M2 macrophages resembled cells labeled as "macrophages" in the lung reference data set. Notably, mononuclear phagocyte cell population sharing M1-M2 markers and labeled in our study as M1-2 macrophages resembled various subsets of DCs in the lung reference data set. The majority (92.4%) of conventional DC (cDC) subset in our study overlapped with various subsets of DCs in the lung reference data sets.

Determination of consistency of T cell annotations **(Supplemental Fig. 2W$_6$)** between our and reference data sets was complicated due to lack of standardized nomenclature of diverse T cell subsets. Consistent with our annotation strategy, 79.9% of Tcn (central memory and naïve) cells showed similarity to CD4$^+$ memory/effector (49%) or CD4$^+$ naïve (30.9%) T cells in the lung reference data set. T-ifn cells found in our study were not found as a separate cell group in the reference data sets, but, in line with their IFN-activated immune gene expression pattern, a subset of Tifn cells showed similarity to proliferating NK/T cells in the lung reference data set. Also, consistent with their annotation in our study, Tcn and Tifn exhibited similarity to CD4$^+$ T cells in the reference data set, and CD8-T1 cells showed the highest degree of overlap, compared to other T cell subsets, with $CD8^+$ memory effector T cells (26.1%) in the lung reference data set. Trm cells categorized as such in our study based on markers identified in recent studies[61–63], as described in the section above, were not defined as a separate cell group in the reference data sets in Azimuth. Consistent with their overall annotation as $CD8^+$ T cell-enriched cell group, major reference cell type categories enriched among Trm cells represented different subsets of T cells (cumulatively 32.5%). When reference data sets were extended to include HuBMAP ASCT plus B category data sets available in Enrichr (https://maayanlab.cloud/Enrichr/#libraries), category "Resident Memory CD8 T Cell" was enriched in Trm cell group compared to other T cell groups identified in our study **(Supplemental Fig. 2$W_7$)**. Consistent with their annotation in our study as a T cell subset, >60% of T-NK cells showed similarity to different subtypes of T cells, and 6.4% resembled NK T cells, as compared to NK cells, 76.4% of which, as described above, overlapped with the NK cell category in the lung reference data set **(Supplemental Fig. 2$W_6$)**. Enrichment of Azimuth reference cell type annotations for immune cell groups described above was evaluated statistically by Enrichr **(Supplemental Figure 2$X_3$)**.

*scRNA-seq Analysis of Changes in Pre-T Airways of Subjects with COPD*

Disease-related changes in cellular composition of pre-T airways in COPD subjects were identified by comparing frequencies of cells belonging to individual cell groups (cell types and cell subtypes defined as described above) in pre-T samples isolated from subjects with COPD and those from donors without lung disease ("normal") based on scRNA-seq data generated as described above (samples described in **Supplemental Table I**). Frequencies of individual cell groups (cell types/subtypes) relative to all single cells captured in each sample as well as relative to all cells within corresponding cell superfamily or family were determined for each group of samples and presented in **Supplemental Table III**. Differences in frequencies of specific cell types/subtypes between individual sample groups (e.g., COPD *vs* normal) were determined by comparing average values in each group per sample and evaluated statistically using two-tailed Mann-Whitney test; data for all comparisons is presented in **Supplemental Table III**.

Differentially expressed genes (DEGs) between COPD and normal pre-T samples were identified by comparing average gene expression in individual cell types/subtypes or families between these study groups per sample (method A; data shown in **Fig. 4C** for cell sub-/types) or per group (method B; data shown in **Fig. 4D** for ASE cell family). In method A, criteria for cell type/subtype inclusion were: >10 cells in group, and at least 3 cells per sample in >50% samples in each group. If no cells representing a specific cell type/subtype included in the analysis were captured in a sample, "0" was assigned as a value representing gene expression level; statistical significance of difference in gene expression between groups was determined by two-tailed Mann-Whitney considering average expression values for individual genes in each sample. In method B, DEGs were determined as described for markers of cell types/subtypes. Criteria for DEGs were p value <0.05, based on two-tailed Mann-Whitney test in method A and adjusted p value (FDR) in method B. DEGs identified using method A for cell types/subtypes in COPD *vs* normal pre-T samples are listed in **Supplemental Table VI**. DEGs identified using method B for ASE cells in COPD *vs* normal pre-T samples are listed in **Supplemental Table VII**.

DEGs identified as up-regulated in ASE cells of COPD *vs* normal pre-T samples using method B and having log2 fold-change >0.2 were analyzed for enrichment of biological process or pathway annotation categories of Gene Ontology (GO) Biological Process 2021, Reactome 2016 and NCI-Nature Pathway Interaction Database 2016 using Enrichr. Significantly enriched categories (adjusted p value [FDR] <0.05) and odds ratios of their enrichment were identified using Enrichr. Significantly enriched categories were ranked by adjusted p value **(Supplemental Table VIII)**, top 10 ranked categories are shown in **Fig. 4E**.

*Immunofluorescence (IF) Analysis*

IF analysis was performed for detection of specific markers in lung tissue samples from donors without lung disease and subjects with COPD, and for quantification of cells expressing specific markers of interest (quantitative IF). Standard histological procedures were used for preparation of tissue samples for this analysis as previously described[2,30,37]. Primary antibodies used for IF staining are described in **Supplemental Table IX**. IF imaging was performed using EVOS FL Auto Imaging System (Invitrogen, US). Quantitative IF analysis was performed for lung tissue samples from donors without COPD and COPD subjects described in **Supplemental Table V** to determine the proportions of cells expressing SCGB3A2, SFTPB and/or SCGB1A1 in ASE of pre-TBs and TBs, including areas of TB-RB junctions. TASCs were defined as cells occupying the luminal compartment of ASE and positive for SCGB3A2 and/or SFTPB as shown in **Fig. 3E-F** and **Supplemental Fig. 4**. Co-staining of SFTPB and SCGB1A1 was performed to distinguish 2 subsets of TASCs ($SFTPB^{+}$ $SCGB1A1^{+}$ and $SFTPB^{+}$ $SCGB1A1^{-}$) from common S (S1)-cells ($SCGB1A1^{+}$ $SFTPB^{-}$).

In quantitative IF analysis, airway segments were classified based on standard anatomical definitions[9,10,68–70]. TBs were defined as airways opening to RBs or containing areas of transition to simple cuboidal epithelium, a precursor of TB-RB transition. RBs were identified as segments immediately distal to TBs lined by simple cuboidal epithelium lacking features of epithelium of conducting airways (cilia and $SCGB1A1^{+}$ S-cells) and interrupted by alveolar out-pockets. TB-RB junction areas were identified as distal regions of TBs lined by single columnar epithelium transitioning to single cuboidal epithelium of RBs. Because it is difficult to precisely annotate pre-TBs in lung tissue sections, in this analysis, pre-TBs were defined more broadly as airways with luminal diameter <2 mm, lined by pseudostratified airway epithelium, lacking features of simple columnar or cuboidal epithelium typical for TBs and not opening directly to RBs.

A total of 130,476 distal ASE cells were evaluated in quantitative IF analysis, including 45,800 and 34,304 ASE cells lining pre-TB and TB regions, respectively, in lung tissue samples of 24 donors without lung disease, and 22,221 and 28,151 ASE cells lining these corresponding regions in lung tissue samples representing 11 subjects with COPD **(Supplemental Table V)**. Description of demographic and general characteristics of samples used in quantitative analysis of SCGB3A2$^+$, SFTPB$^+$ and SCGB1A1$^+$ cells in pre-TBs, TBs and TB-RB junction regions of samples in COPD and non-diseased study groups is provided in **Supplemental Table V**. Data for different airways or airway areas representing each anatomical region (pre-TB, TB including TB-RB junction, and TB-RB junctions separately) in each subject were averaged to determine the average frequency of SCGB3A2$^+$ TASCs, SFTPB$^+$ TASCs, SCGB1A1$^+$ SFTPB$^+$ TASCs and SCGB1A1$^+$ SFTPB$^-$ common S- (S1) cells in pre-TBs, TBs and TB-RB junction in each subject. For each marker and anatomical region, quantification data determined for each sample was used for comparison of groups, and statistical significance of differences between the groups was determined using two-tailed Mann-Whitney test. Statistical summary of quantitative IF analysis for each comparison is provided in **Supplemental Table V**.

*Imaging CyTOF analysis*

Imaging CyTOF analysis was performed for selected samples **(Supplemental Table V)** for simultaneous visualization of multiple markers. Antibodies used in this analysis are listed in **Supplemental Table IX**. Antibody clones were validated by IF staining of lung tissue samples as described above prior to their usage for imaging CyTOF analysis. After validation, 100 µg of purified antibody was procured from respective vendors for custom conjugation **(Supplemental Table IX)**. MaxPar antibody labeling kit (Fluidigm) was used to conjugate to appropriate metal choice following manufacturer’s protocol. Metal-labeled antibodies were re-validated by staining control lung tissue samples. Formalin fixed paraffin embedded slides containing lung tissue samples were stained with selected antibodies following manufacturer's protocol as previously described[71]. Imaging mass cytometry (IMC) ablation was performed using Hyperion (Fluidigm) at the Englander Institute for Precision Medicine (Weill Cornell). MCD files acquired with the Hyperion instrument were analyzed using MCD Viewer (Fluidigm) to visualize IMC data for selected markers. Representative images generated using this method are shown in **Fig. 3J** and **Supplemental Fig. 3**.

*Cell Culture Studies*

For *in vitro* studies, single cell suspensions were prepared from samples representing proximal (P) and distal (D) airways, the latter representing pre-T airway region, following the protocol of tissue dissociation for scRNA-seq analysis as described above. Samples used in cell culture studies are described in **Supplemental Table X**. Dissociated cells were seeded into cell culture flasks (Corning) in PneumaCult-Ex media (Stemcell Technologies), which supports expansion of basal cells (BCs; https://www.stemcell.com/pneumacult-ex.html)[72], supplemented with 0.25 µg/ml amphotericin B and 50 µg/ml gentamicin (both: Sigma) and incubated at 37°C; 5% $CO_2$. Within 12 hr of seeding, cells were washed with PBS and fresh media was added. After reaching ~80% of confluency (typically after 1-2 wk of propagation), the cells were dissociated using accutase (Innovative Cell Technologies) and used in *in vitro* studies described below.

Air-liquid interface (ALI) culture system was used to study BC differentiation *in vitro* following standard protocols[73] with some modifications to media composition specified below. Suspension of BCs ($8x10^4$ cells/well) expanded as described above were seeded onto 6.5-mm transwell inserts (Corning) pre-coated with type IV collagen (Sigma, C7521) as described[73] and cultured in PneumaCult-Ex Plus media (Stemcell Technologies) supplied from both apical and basolateral sides. Once cells established a confluent layer, ALI was established (ALI-day 0) by aspirating media from the apical chamber, and differentiation (ALI) media was added from the basolateral side. ALI media contained Small Airway Epithelial Cell Basal Medium (PromoCell), supplemented with 2 μg/ml insulin, 0.2 μg/ml hydrocortisone, 0.2 μg/ml epinephrine, 2.7 ng/ml triiodothyronine, 4 ng/ml transferrin, 1 mg/ml bovine serum albumin, 1.6 μl/ml bovine pituitary extract (all PromoCell), $10^{-7}$ M retinoic acid, 0.88 mM $CaCl_2$, 0.25 μg/ml amphotericin B and 50 μg/ml gentamicin (all Sigma). Barrier integrity as well as emergence and persistence of beating cilia (a hallmark of differentiation) were monitored every other day[73]. Samples were harvested at different time-points of ALI (from day 0 to 28) to assess differentiation. To evaluate the impact of IFN-γ on BC differentiation, in selected experiments, recombinant human IFN-γ (5 ng/ml; R&D Systems) was applied together with media every other day from the basolateral ALI side (to enable direct access to BCs) at different time-periods of ALI culture (from days 0-29 to days 18-40; specified in **Supplemental Table X**) to capture the effect on various differentiation states of BC-derived epithelia.

Differentiation phenotype of epithelia generated in ALI model was assessed by analysis of gene expression using 3 methods: 1) bulk RNA-seq - to compare transcriptional profiles of epithelia generated by matched P- and D-airway BCs isolated from 6 donors and cultured in ALI during 18 days under identical conditions; 2) scRNA-seq analysis - to characterize dynamics of BC differentiation isolated from D-airways of donor without lung disease by profiling epithelia generated by D-airway BCs at ALI-days 0, 3, 7, 14 and 28 at single-cell level; and 3) targeted analysis of genes indicative of specific aspects of airway epithelial differentiation using TaqMan real-time PCR - to evaluate the effect of IFN-γ on D-airway BC differentiation. Samples used in each of these analyses are described in **Supplemental Table X**. For selected samples, IF analysis of ALI membranes was performed to detect cells expressing SCGB3A2 and MUC5AC using antibodies described in **Supplemental Table IX** and established methods summarized above.

For bulk RNA-seq and TaqMan real-time PCR, cultured cells were lysed using Trizol (Thermo Fisher Scientific), and RNA was extracted from cell lysates using RNeasy MinElute Cleanup Kit (Qiagen) following manufacturer's instructions. Description of bulk and scRNA-seq analyses of cultured samples is provided below. For quantitative TaqMan real-time PCR, cDNA was reverse transcribed from total RNA using cDNA synthesis kit (Thermo Fisher Scientific) as per manufacturer's instructions. Quantitative real-time PCR was performed using TaqMan Fast Advanced Master Mix (Thermo Fisher Scientific) on QuantStudio 5 Real-Time PCR System (Applied Biosystems). The following TaqMan gene expression assays (Thermo Fisher Scientific) were utilized: IFN-γ response genes *MX1* (Hs00895608_m1) and *IFIT1* (Hs01675197_m1), differentiation-related genes *SCGB3A2* (Hs00944206_g1), *SFTPB* (Hs01090667_m1), *SCGB1A1* (Hs00171092_m1), *MUC5B* (Hs00861588_m1) and *FOXJ1* (Hs00230964_m1). 18S rRNA (Thermo Fisher Scientific, 4310893E) was used as an endogenous control for relative gene expression quantification as previously described[37].

*Bulk RNA-seq Analysis of Epithelial Samples Generated in ALI*

A total of 12 epithelial samples generated in ALI culture (ALI-day 18) from matched P- and D-airway BCs isolated from 6 donors without lung disease **(Supplemental Table X)** were processed for bulk RNA-seq analysis following standard protocols. Briefly, RNA isolated from these samples as described above was reverse transcribed to cDNA, amplified, and prepared as sequencing libraries. Libraries were sequenced on a MiSeq Illumina instrument to obtain 2x150 base pair paired-end reads. The reads were mapped using STAR[74], gene symbols were converted to GRCh38. DEGs between ALI samples generated by D- and P-airway BCs were identified by two-tailed paired t-test ($p<0.05$). Mean log2 fold-change (log2 FC) was determined based on log2 (TPM+1) fold-change values calculated for all six P-D pairs. The complete list of DEGs identified in this analysis is presented in **Supplemental Table XI**. Then proportions of DEGs up-regulated in ALI-day 18 ASE generated by D- and P-airway BCs overlapping with marker genes of pre-T ASE cell types identified based on scRNA-seq analysis of in vivo pre-T samples (criteria: log2FC >1; FDR<0.05) were determined.

*scRNA-seq Analysis of Distal Airway BC Differentiation In Vitro*

To evaluate the dynamics of distal airway BC differentiation at single-cell level, epithelia generated at days 0, 3, 7, 14, 21 and 28 of ALI culture by the same initial D-airway BC sample from a donor without lung disease (see **Supplemental Table X**) were profiled using scRNA-seq. At each ALI time-point indicated above, cells were dissociated using accutase and processed for scRNA-seq library preparation and analysis as described above (see section *scRNA-seq Library Preparation and Sequencing*). scRNA-seq data generated for these samples included dimension reduction, cell clustering, and doublet calling step following the protocols mentioned above (see section *Dimensionality Reduction, Cell Clustering, and Doublet Removal*). A total of 43,854 single cells were captured in this analysis **(Supplemental Table XII)**. Data obtained for ALI-day 21 sample was comparable to ALI-day 28 data in many critical aspects including the overlapped UMAP clustering pattern **(Supplemental Fig. 9$A_1$)**. Thus, whereas this time-point was included in all steps of scRNA-seq data analysis, as detailed below, to avoid redundancy and due to space limitations, data for this time-point was not presented in the main part of this manuscript. The latter included data for a total of 36,321 single cells captured for samples representing ALI-days 0, 3, 7, 14 and 28 **(Supplemental Table XII)**.

Cell type/subtype annotation was performed based UMAP clustering data generated for cells captured for all time-points, using clustering resolution (res) 0.01, 0.09 and 0.6 and markers of identified clusters using methods described in *Cell Group Annotation* section above. Basal cells (*KRT5*$^+$) were identified at res 0.01 within clusters 2 (unique to ALI-day 0), 1 (emerged and dominant at ALI-day 3) and 4 (containing cells from later time-points) **(Supplemental Fig. 9A**

**and B)**. This indicates heterogeneity of BCs related to the differentiation state of the epithelium, which is progressively achieved in this model. Several BC subsets were identified within these clusters. First, secretory-like BCs (BC-s; *SCGB1A1*$^{+}$ *MUC5B*$^{+}$) were uniquely present at ALI-day 0. They clustered together with other cells (BCs) at this time-point (cluster 2; res 0.01) but formed a separate cluster (cluster 5) at res 0.09 **(Supplemental Fig. 9B)**. Notably, BC-s showed relatively low expression of BC marker genes, including *KRT5*, suggesting that this subset may represent the earliest intermediate cell (IC) precursors primed toward S-cell differentiation. To distinguish these cells from separately clustered ICs at later ALI time-points, and based on their co-clustering with BCs at ALI-day 0, BC-s were annotated as a BC subset. Second, proliferating BCs (BC-p; *MKI67*$^{+}$) were identified as a separate cluster 14 at res 0.09 corresponding to cluster 37 at res 0.6 **(Supplemental Fig. 9B)**. A unique subset of BCs (clusters 12 and 13 at res 0.09) showed high expression of extracellular matrix-related genes *(MMP1, LAMC2, FN1, MMP10, LAMB3,* see all marker genes in **Supplemental Table XIII**) and, because of this unique gene expression pattern, was termed matrix BCs (BC-m) **(Supplemental Fig. 9B)**. Remaining cells in clusters 2 and 4 at res 0.01 and cluster 9 at res 0.09 were annotated as common BCs (BC1).

Intermediate cells (IC) were identified within clusters 6 and 7 at res 0.09 and clusters 23 and 25 at res 0.6 based on high expression of *SERPINB3* and intermediate clustering position, between BCs and secretory (S)-cells (described below). A unique subset of ICs expressing genes associated with squamous differentiation, including *KRT6A*, *SPRR1B* and other genes coding for SPRR-family proteins **(Supplemental Table XIII)**, were found in cluster 32 at res 0.6 and were designated "squamous ICs" (IC-sq) cells **(Supplemental Fig. 9C)**. Thus, ICs included a minor, IC-sq subset, whereas remaining cells constituted common ICs (IC1). Secretory (S) cells were identified in cluster 0 at res 0.01 based on high expression of *SCGB1A1*, *BPIFA1* **(Supplemental Fig. 9D)** and other S-cell markers **(Supplemental Table XIII)**. S-cells emerged at ALI-day 7 as a cluster extending further away from IC1 on the differentiation trajectory of BCs. TASCs were identified within S-cell cluster based on expression of *SCGB3A2*, *SFTPB* (expressed broadly by S-cells generated by D-airway BCs in ALI model) and *PGC,* and constituted clusters 0 and 30 at res 0.6 **(Supplemental Fig. 9D)**. TASCs emerged at ALI-day 7 as an extension of S-cell cluster further away from BCs and ICs, and progressively increased in number by ALI-day 28 **(Fig. 5F)**.

Ciliated (C) cells were identified in cluster 3 at res 0.01 by high expression of *FOXJ1* **(Supplemental Fig. 9E)** and other C-cell marker genes **(Supplemental Table XIII)**. Similar to *in vivo* airway tissue samples, C-s subset of C-cells was identified (cluster 9 res 0.6) based on high expression of *SAA* family genes, including *SAA1* **(Supplemental Fig. 9E)**, *SAA2* and *SAA4* **(Supplemental Table XIII)**. Thus, C-cells included C-s subset and remaining common C (C1) - cells. In addition to mature C-cells, pre-ciliated (p-C) cells were identified (cluster 1 at res 0.09) based on high expression of *CCNO* and *HES6* **(Supplemental Fig. 9E)**, markers of p-C cells in *in vivo* airway tissue samples. In support of their "pre-ciliated" nature, pre-C cells emerged early during BC differentiation (ALI-days 0-3), prior to emergence or expansion of mature C-cells.

The D-airway BC-derived epithelia also contained ionocytes (Ion; cluster 6 at res 0.01) found based on expression of *FOXI1*, *HEPACAM2*, *ATP6V0C*, *ATP6V0B* **(Supplemental Fig. 9F)** and other genes known to mark Ion **(Supplemental Table XIII)**. Ion emerged at ALI-day 3 and reached highest frequency (0.28%) by ALI-day 7 and maintained at similar level until ALI-day 28 **(Supplemental Table XIV)**. Another group of rare cells (<0.5% cells; cluster 5 res 0.01) emerged, similar to Ion, at ALI-day 3 (0.1%). Their frequency increased at ALI-days 21 and 28 to ~0.3-0.4% **(Supplemental Table XII)**. These cells expressed some molecular features of Ion (*FOXI1, HEPACAM2)* and NE cells (*TFF3, HES6, CRYM, GRASP, TUBB3*), but differed from the latter by expression of markers of brush, or tuft (*DCLK1*, *ASCL2*, *POU2F3, TRPM5, SOX4, NKX3-1*)[42,43] cells **(Supplemental Fig. 9G)** and, thus, were annotated as "tuft-like" cells (TLCs).

### Supplemental Figure Legends

**Supplemental Figure 1.** Region-precise dissection of human lung tissue. **A.** Proximal airways. **$A_1$.** Representative images of human lung tissue samples; proximal airways (bronchi, Br) and arteries (Ar) are labeled; **$A_2$.** Examples of dissected proximal airways (cartilaginous bronchi, 1-4 generation). **B.** Examples of distal portions of lung tissue samples from: **$B_1$** - donors without lung disease, and **$B_2$** - subjects with chronic obstructive pulmonary disease (COPD); age of subjects (years, yr) are indicated; magnified images show distal airways (Aw) and vessels (Vs) identified in these samples upon macroscopic inspection (labeled); arrows: air trapping in emphysematous alveolar region in the lungs of COPD subjects. **$B_3$.** Examples of lung tissue samples from a donor without lung disease ("normal lung") and subject with COPD ("COPD lung") before and after inflation performed by injection of air through distal airway to visualize supplied alveolar region. **C.** Representative images showing dissection of distal airways until the last macroscopically distinguishable segments (secondary lobular bronchioles, or pre-terminal bronchioles, pre-TBs). **D.** Characterization of pre-TBs and secondary pulmonary lobules as anatomical landmarks of distal lung region: **(i)** initial lung tissue sample; highlighted area contains region shown in **ii-viii**; **(ii)** cross section of lung tissue containing lobule (yellow dashed area) with surrounding septa, veins; centrilobular region contains artery and pre-TB labeled; **(iii)** lung region containing lobule shown in **ii** was excised to facilitate access to alveolar tissue supplied by distal airway shown in white dashed area in **ii** and **iv** and through which air **(v)** and subsequently ink (trypan blue; **vi**) were injected to visualize terminal (lobular, alveolar) region supplied by this airway (by inflation and trypan blue staining, respectively); after which **(vii)** this airway was dissected toward stained supplied alveolar region, containing lobules, which were dissected together with supplying pre-TBs **(viii)** with subsequent separation from the latter (**ix**, shown: two lobules separated by septa). **E.** Examples of dissected distal airway units **($E_1$)**, containing pre-terminal (pre-T) airways (pre-

TBs and bronchioles proximal to pre-TBs, annotated based on their branching order relative to pre-TBs) and terminal (T) region, containing anatomical segments distal to pre-TBs (contributing to lobules each supplied by an individual pre-TB). Pre-TBs served the boundary between pre-T- and T-regions. Dashed area in the middle panel in **E1** was inspected under dissection microscope **(**$\mathbf{E_2}$ **left**; showing indicated airway segments; air bubbles traveling along these airways following inflation; and alveolar attachments (AA) to bronchiolar walls); region in the white dashed area was additionally examined under the light microscope **(E2 right)**. Pre-T and T-regions of sample shown in $\mathbf{E_1}$ **middle** separated from each other using pre-TB as a boundary **(**$\mathbf{E_3}$**; left)**; T-region was examined under the light microscope ($\mathbf{E_3}$ **right**; note: lack of airways and the presence of air bubbles reaching alveolar region after inflation). $\mathbf{E_4}$**.** Examples of pre-T airways dissected from lungs of donors without lung disease ("normal") and lungs of COPD subjects. **F.** Hematoxylin and eosin staining of representative dissected proximal, pre-T and T regions; major histological landmarks are shown and described.

**Supplemental Figure 2.** Annotation of cell types and cell subtypes based on scRNA-seq data. $\mathbf{A_1}$**.** Identification of clusters representing major cell families based on expression of indicated markers at clustering resolution (res) 0.0001. $\mathbf{A_2}$. Clusters at res 0.001, 0.1, 0.1. $\mathbf{A_3}$. Clusters at res 0.8, 1.0, 2.0, 3.0, 4.0 and 5.0. **B-V.** Annotation of **(B-J)** epithelial, **(K-O)** structural, **(M-V)** immune cell types and subtypes using indicated markers (see description in the corresponding Supplemental Information - Methods section). $\mathbf{W_{1\text{-}6}}$**.** Distribution of Azimuth-based cell type annotations of cells within each cell type and cell subtype identified in scRNA-seq analysis of our study. In all panels, annotation was performed based on the human lung reference scRNA-seq data set (Travaglini et al.[18]) in Azimuth; numbers represent % of a specific cell type/subtype reference annotations within an individual cell type/subtype identified in our study. For Gli and En-v, annotations using reference data sets for all tissues available in Azimuth were determined using SCINA algorithm, in addition to human lung reference data set. Categories with frequency of distribution >5% within each indicated cell group are labeled. **$W_7$.** Enrichment of "Resident Memory CD8 T Cell (id:540)" HuBMAP ASCT plus B category determined using Enrichr in indicated T cell subsets identified in our study; dots: enrichment % (% genes overlapped with the reference gene set; bars: -log FDR of enrichment. **X.** Bubble-plots: distribution of Azimuth annotations determined using SCINA for epithelial **($X_1$)**, structural **($X_2$)** and immune **($X_3$)** cell types. Categories with Odds ratio >100 and FDR<0.05 are included.

**Supplemental Figure 3.** Cell mapping to lung tissue domains using imaging CyTOF. **A** and **B.** Representative examples of CyTOF images showing expression of indicated markers in different airway regions – proximal airways (bronchus) and distal lung region containing distal airways, including terminal bronchioles (TBs) and respiratory bronchioles (RBs). Images shown on each page show staining of the same markers (different colors) unless indicated otherwise.

**Supplemental Figure 4.** Morphological identification and characterization of terminal airway-enriched secretory cells (TASCs) in the human lung. Representative images of proximal airway (bronchus) and lung tissue sections of donors without lung disease (from samples described in **Supplemental Table V**) showing immunofluorescence (IF) staining of: **$A_{1\text{-}13}$.** secretoglobin 3A2 (SCGB3A2, red), a TASC marker; and tubulin, beta 4 (TUBB4, green), a marker of cilia (ciliated cells); and **$B_{1\text{-}12}$.** surfactant protein B (SFTPB, red), a marker of TASCs; and secretoglobin 1A1 (SCGB1A1, green), a marker of secretory (S) cells, including common S cells (S1 cells).

**Supplemental Figure 5.** Comparison of transcriptional profiles of terminal airway-enriched secretory cells (TASCs) in terminal (T) *vs* pre-terminal (pre-T) airway regions based on scRNA-seq analysis. **A.** Volcano plot showing differentially expressed genes (adjusted p value [FDR] <0.05) identified by comparing TASCs in T and pre-T airway regions; **B.** Box plot showing expression of the surfactant protein A2 *(SFTPA2)* gene in TASCs of pre-T and T-regions.

**Supplemental Figure 6.** Representative images showing immunofluorescence (IF) staining of human lung tissue samples from indicated groups for: **A**. secretoglobin 3A2 (SCGB3A2, red), a TASC marker and, in selected images, tubulin, beta 4; TUBB4, green, a marker of cilia; and **B.** surfactant protein B (SFTPB, red), a marker of TASCs; and secretoglobin 1A1 (SCGB1A1, green), a marker of secretory (S) cells; shown examples of pre-terminal bronchioles (pre-TBs) and terminal bronchioles (TBs) in lung tissue samples from donors without lung disease and subjects with chronic obstructive pulmonary disease (COPD) used in quantitative IF analysis; sample ID – see sample characteristics in **Supplemental Table V**.

**Supplemental Figure 7.** Immunofluorescence (IF) analysis of secretoglobin (SCGB1A1)$^+$ common secretory (S1) cells (distinguished from SCGB1A1$^+$ TASCs by SFTPB negativity) in pre-terminal (pre-TB) and terminal (TB) bronchioles in lung tissue samples of donors without lung disease (Normal; N) and COPD subjects (described in **Supplemental Table V**). **A.** Data of quantitative IF analysis showing percentage (%) of SCGB1A1$^+$ S1-cells among airway surface epithelial (ASE) cells in pre-TBs and TBs of indicated groups; **B.** Correlation plots showing relationships between frequencies of SFTPB$^+$ TASCs (x axis) and SCGB1A1$^+$ (SFTPB-negative) S1-cells in pre-TBs and TBs of indicated groups; Spearman correlation coefficient (Rho) and p values of correlation (two-tailed) are shown. Each dot represents an individual subject.

**Supplemental Figure 8.** Quantitative immunofluorescence (IF) analysis of terminal airway-enriched secretory cells (TASCs) in pre-terminal (pre-TB) and terminal bronchioles (TBs) in lung tissues of nonsmokers (NS) and smokers (S) without lung disease (each dot represents an individual subject). Shown frequencies (% of all airway surface epithelial [ASE] cells) of: **A.** SCGB3A2$^+$ TASCs and **B.** SFTPB$^+$ TASCs in indicated groups; p-values shown are based on two-tailed Mann-Whitney test.

**Supplemental Figure 9.** Annotation of cell types and cell subtypes based on scRNA-seq data of epithelia generated by distal (D)-airway basal cells (BCs) during their differentiation in air-liquid interface (ALI) culture. **$A_1$. Left:** UMAP clustering plot showing distribution of 43,854 single cells captured at indicated time-points of ALI culture. **Right:** Cell type/subtype annotation (see detailed description in **Supplemental information**, principles of annotation are demonstrated in panels **B-G** as described below. **$A_2$**. UMAP clusters at resolutions (res) 0.01, 0.09 and 0.6 used for identification and annotation of cell types and subtypes. **B-G.** Annotation of undifferentiated cells: **B:** BCs; **C:** intermediate cells (ICs), **D:** secretory (S) cells, including common S-cells (S1) and terminal airway-enriched S-cells (TASCs); **E:** ciliated (C) cells, including common C-cells (C1) and serum amyloid A (SAA)$^+$ secretory-like C (C-s) cells; and pre-ciliated (p-C) cells; **F:** ionocytes (Ion), and **G:** tuft-like cells (TLCs). Distribution of individual cell types/subtypes in each time-point of ALI culture is provided in **Supplemental Table XII**; see complete list of cell type/subtype markers in **Supplemental Table XIII**.

**Supplemental Figure 10.** Characterization of IFN-gamma signaling components in the human lung. **A.** Mean expression levels of *IFNG*, *CD8A* and *CD8B* genes across the cell types/subtypes captured in our scRNA-seq data (data for all samples and regions as described in **Supplemental Table I**). These data show that CD8$^+$ T cells are among the major *IFNG*-expressing cells in the human lung/airways. **B.** Images representing imaging CyTOF analysis of lung tissue samples of subjects with COPD **($B_1$-$B_3$)** and donors without lung disease **($B_4$-$B_6$)** using indicated markers. Note, CD8A$^+$ T cells are distributed in subepithelial and intraepithelial compartments of distal (pre-terminal [pre-TB] and/or terminal bronchioles [TB]) airways in close proximity to airway basal cells (BCs; keratin 5 [KRT5]$^+$). **C.** scRNA-seq data showing expression of IFN-$\gamma$ receptor genes *IFNGR1* and *IFNGR2* in different cell types/subtypes of airway surface epithelium (ASE) in *in vivo* airway tissue samples **($C_1$)** (all samples described in **Supplemental Table I**) and those in the epithelium generated by distal airway BCs *in vitro* in air-liquid interface (ALI) culture **($C_2$)** (combined data for all time-points as described in **Supplemental Table X**). **D.** Analysis of the effect of IFN-γ on airway epithelial phenotype in air-liquid interface (ALI) culture. **$D_1$.** Study design. BCs isolated from distal (pre-terminal, pre-T) airways dissected from indicated samples (see **Supplemental Table X** for sample characteristics) were cultured in ALI model, in which BCs differentiate into mucociliary airway epithelium over 3-4 weeks of culture, in the presence or absence of 5 ng/ml of recombinant human IFN-γ added from basolateral side every other day during indicated time-periods of ALI culture. Once treatment was completed, epithelia derived from BCs at indicated time-points of ALI was analyzed using TaqMan real-time PCR (data shown in **$D_2$**). **$D_2$.** log2 normalized expression of indicated genes in ALI-day 18-40 epithelia generated by distal airway BCs in the presence or absence of 5 ng/ml IFN-γ; dot colors mark independent samples shown in **$D_1$**; p values based on 2-tailed paired t-test.
