## Supplementary material for "A Unique Cellular Organization of Human Distal Airways and Its Disarray in Chronic Obstructive Pulmonary Disease": Fig. S1

**A₁** Inspection of proximal lung region

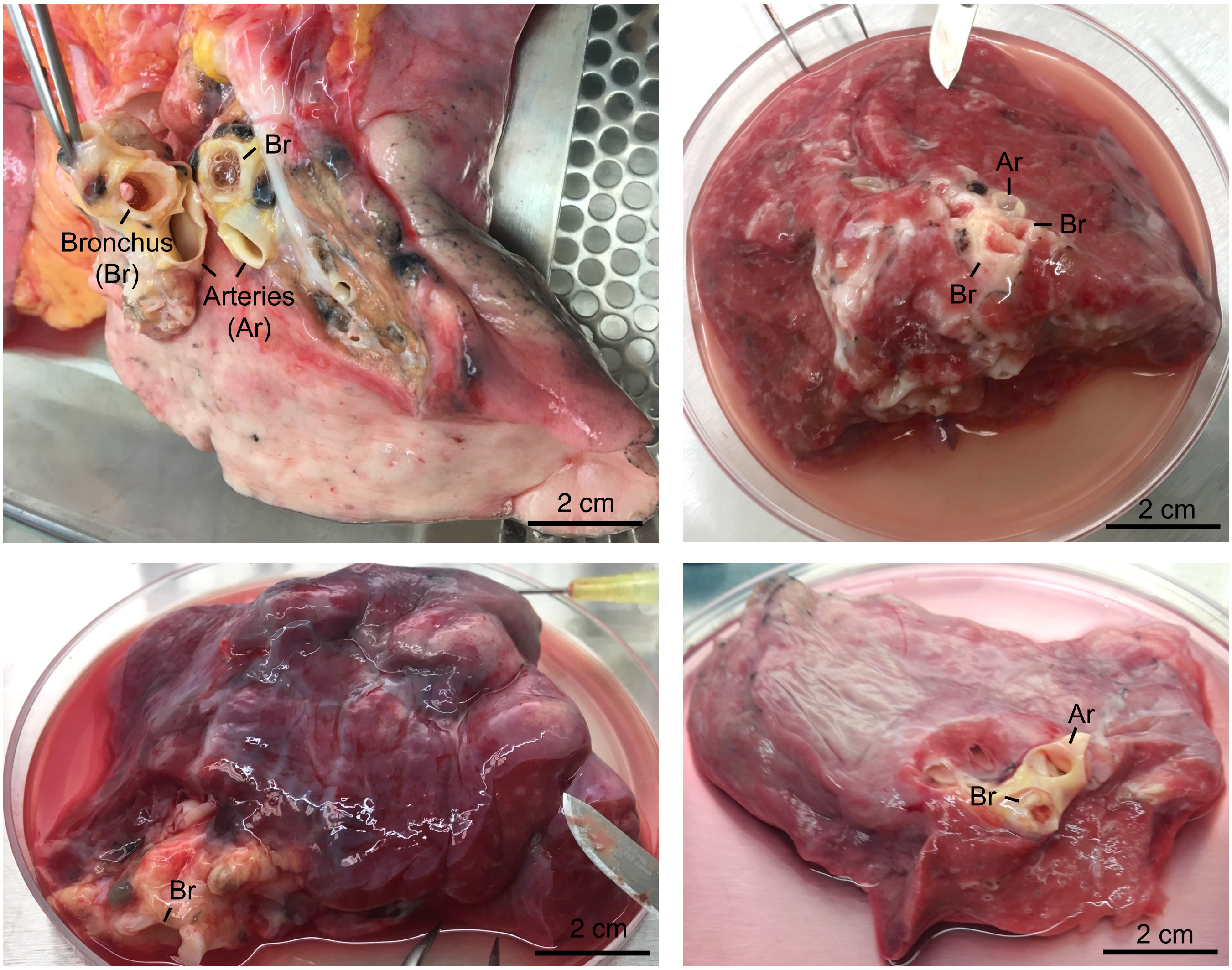


**A₂** Examples of dissected proximal airways (bronchi)

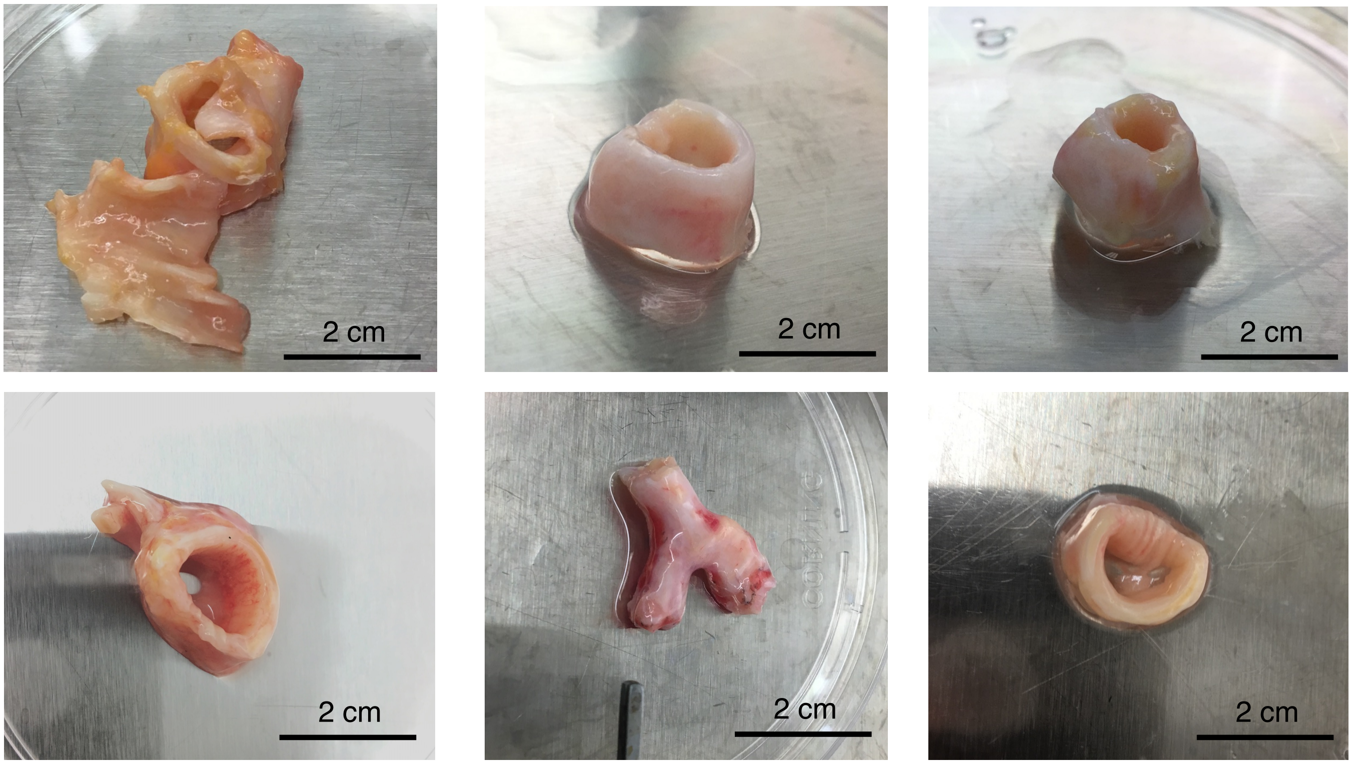

**Supplemental Figure 1 (continued)**

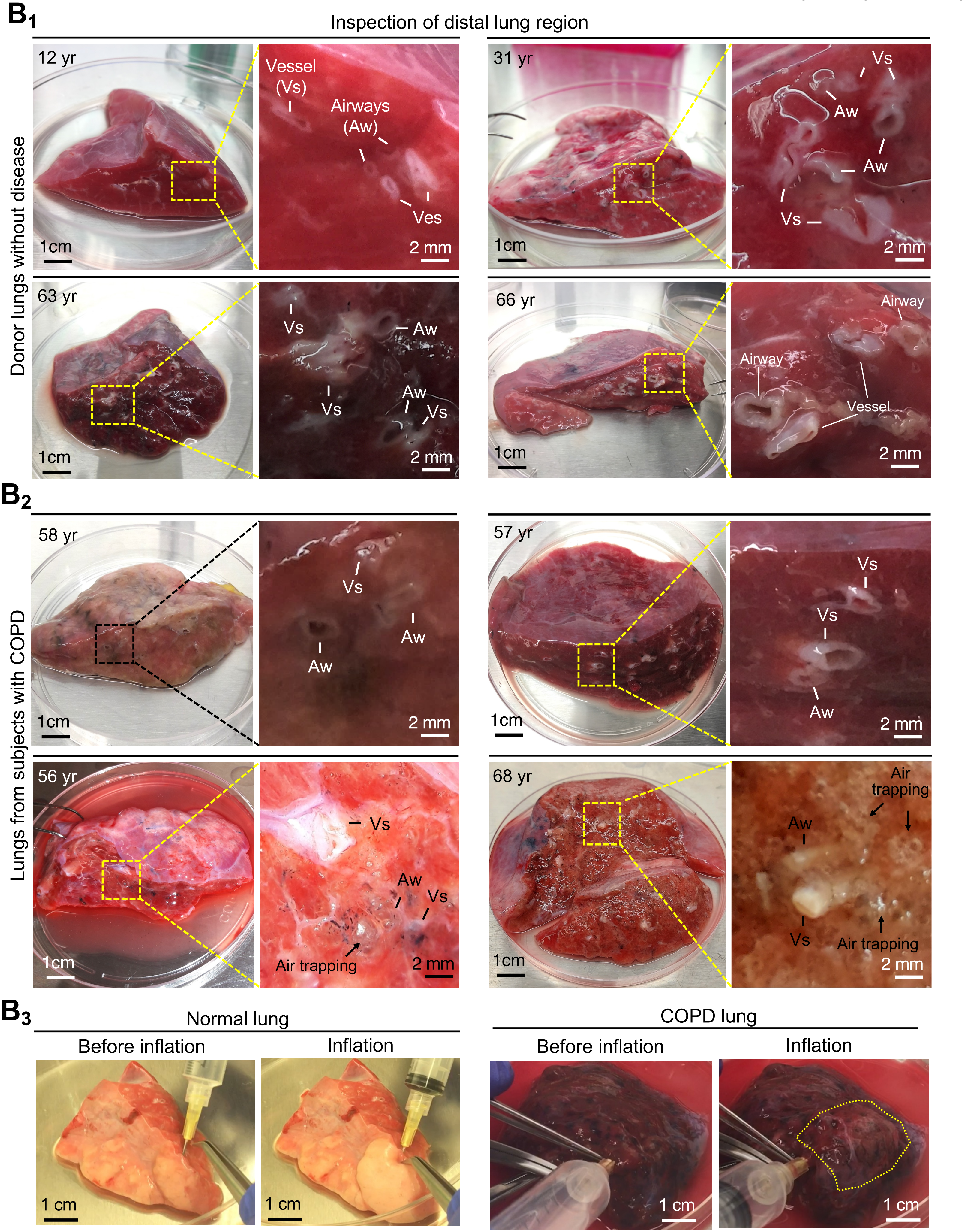

**C**

Dissection of distal airways

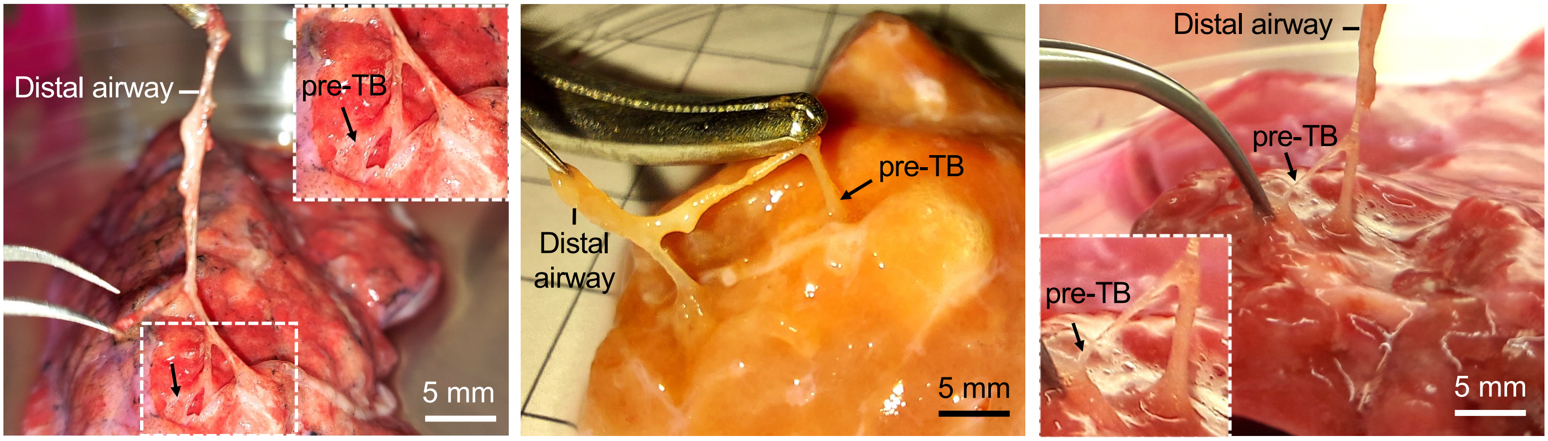


**D**

Characterization of pre-TB and lobules

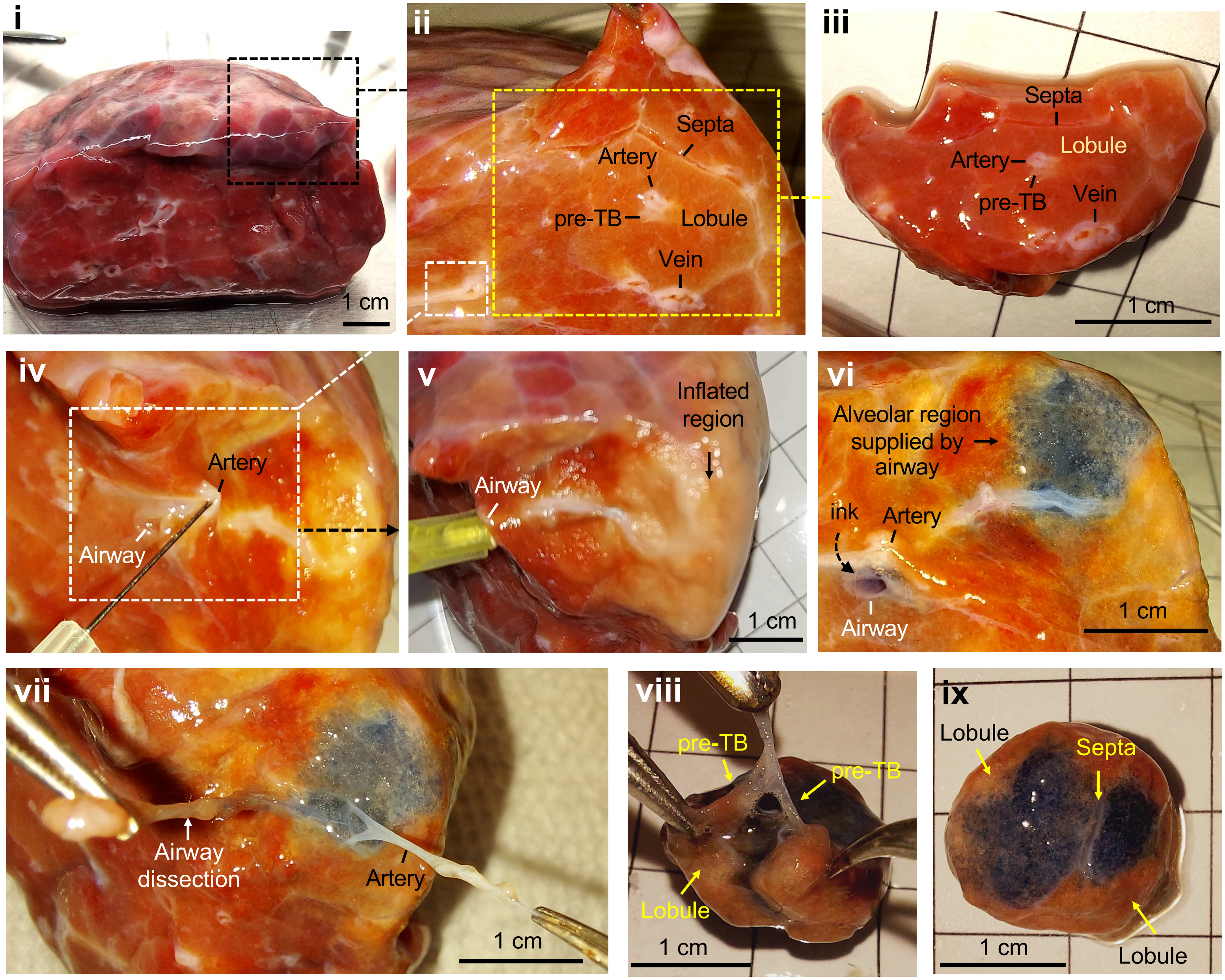

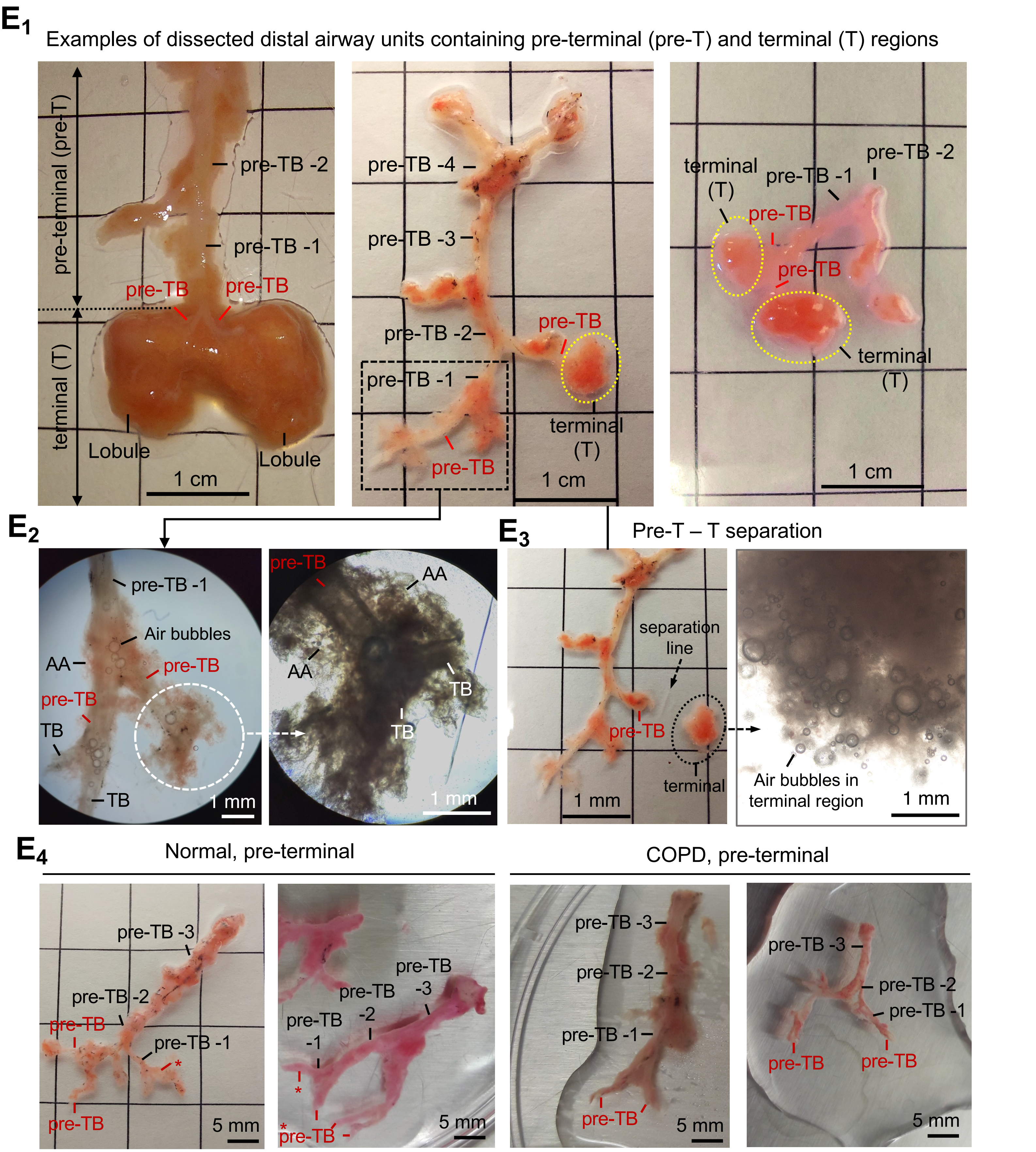
E1
Examples of dissected distal airway units containing pre-terminal (pre-T) and terminal (T) regions
pre-terminal (pre-T)
terminal (T)
pre-TB -2
pre-TB -1
pre-TB
pre-TB
Lobule
Lobule
1 cm
pre-TB -4
pre-TB -3
pre-TB -2
pre-TB -1
pre-TB
pre-TB
terminal (T)
1 cm
terminal (T)
pre-TB -1
pre-TB -2
pre-TB
pre-TB
terminal (T)
1 cm
E2
pre-TB -1
Air bubbles
AA
pre-TB
pre-TB
TB
TB
1 mm
pre-TB
AA
AA
TB
TB
1 mm
E3
Pre-T – T separation
separation line
pre-TB
terminal
1 mm
Air bubbles in terminal region
1 mm
E4
Normal, pre-terminal
COPD, pre-terminal
pre-TB -3
pre-TB -2
pre-TB
pre-TB -1
*
pre-TB
5 mm
pre-TB -3
pre-TB -2
pre-TB -1
*
*
pre-TB
5 mm
pre-TB -3
pre-TB -2
pre-TB -1
pre-TB
5 mm
pre-TB -3
pre-TB -2
pre-TB -1
pre-TB
pre-TB
5 mm

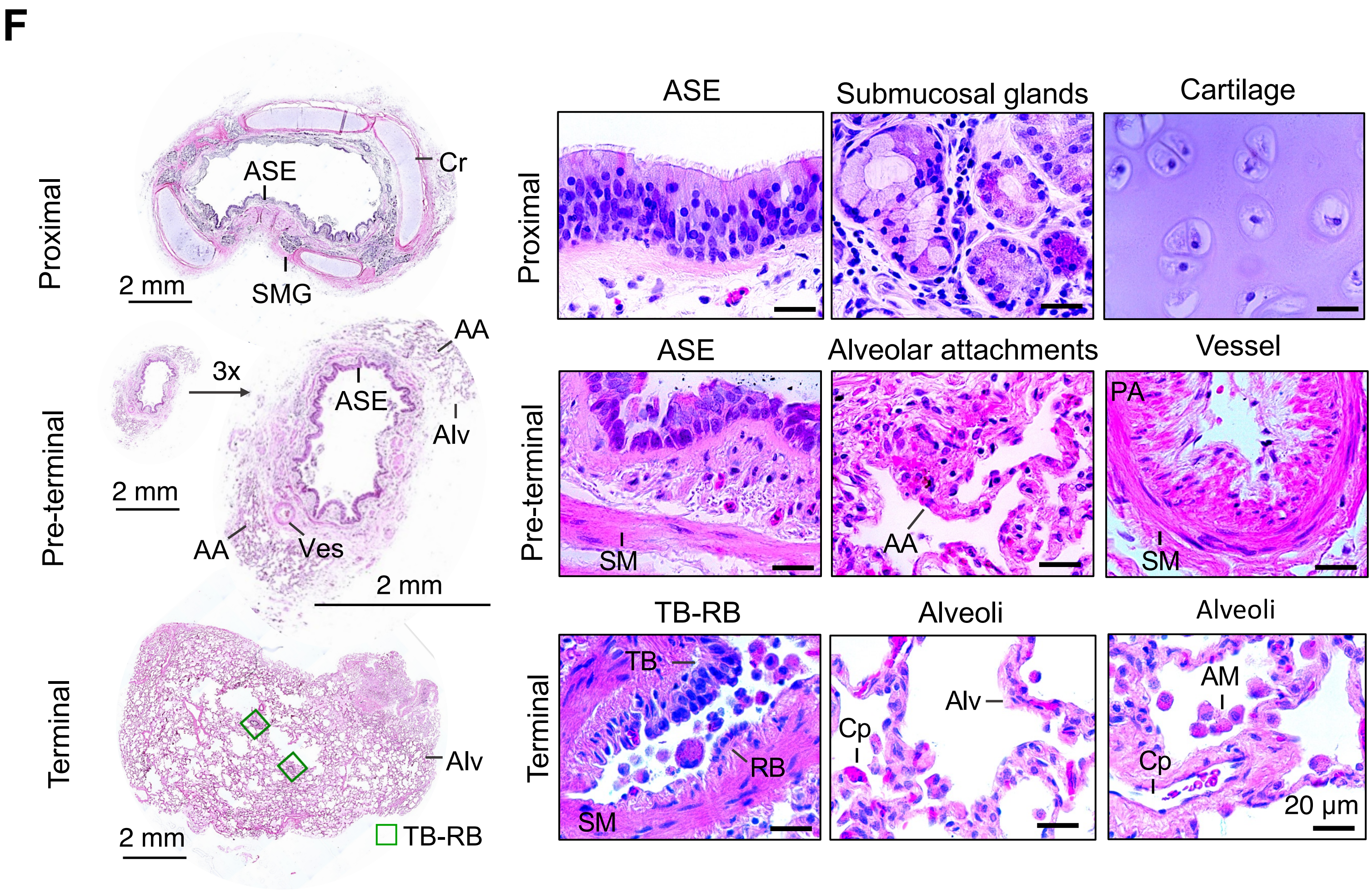


**ASE**, airway surface epithelium
**AA**, alveolar attachments
**SMG**, submucosal glands
**Cr**, cartilage
**Alv**, alveoli
**Ves**, vessels
**TB**, terminal bronchiole
**RB**, respiratory bronchiole
**SM**, smooth muscle
**PA**, pulmonary artery
**Cp**, capillary
**AM**, alveolar macrophage
