## Supplementary material for "A Unique Cellular Organization of Human Distal Airways and Its Disarray in Chronic Obstructive Pulmonary Disease": Fig. S2

### Supplemental Figure 2

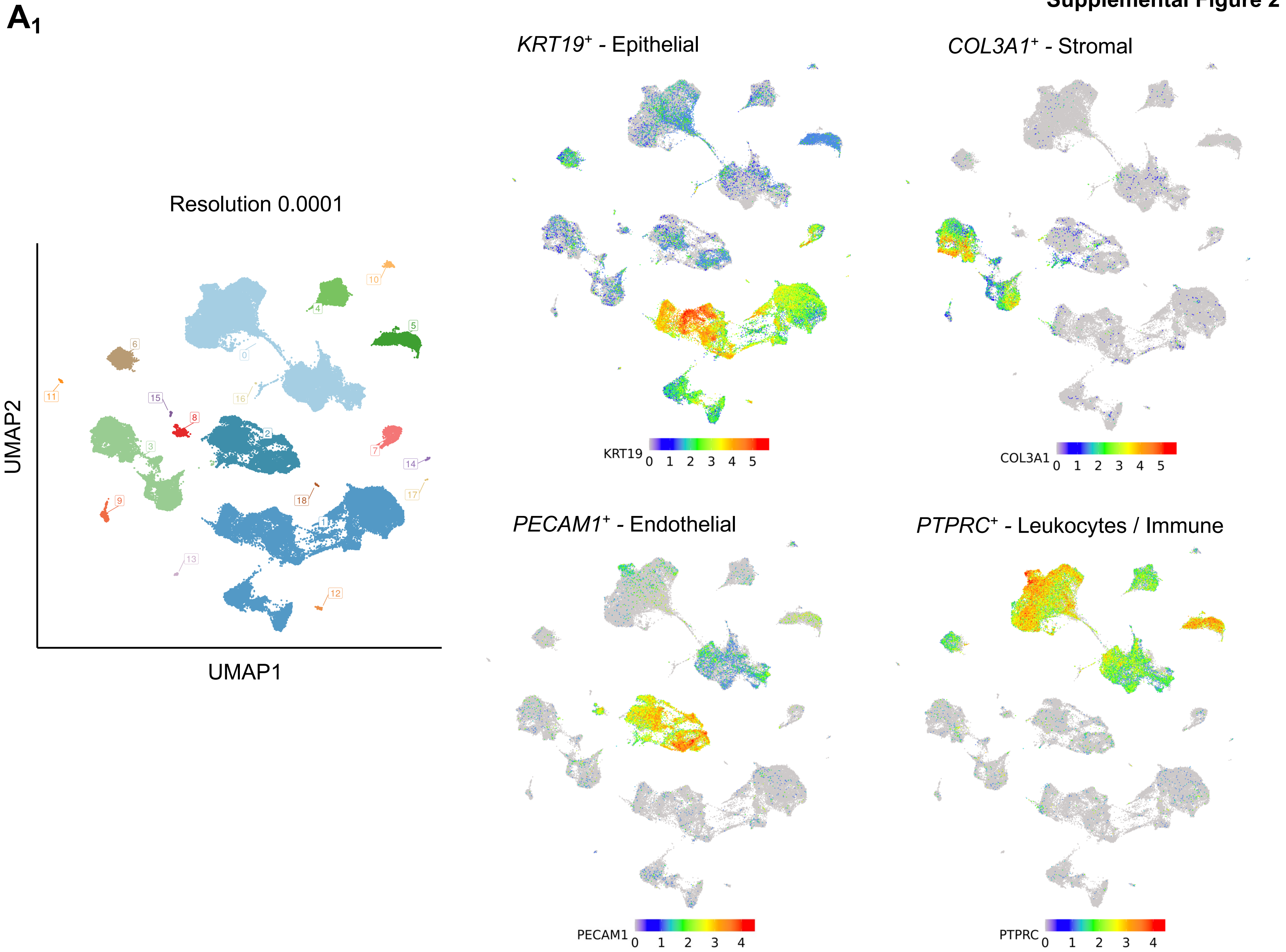

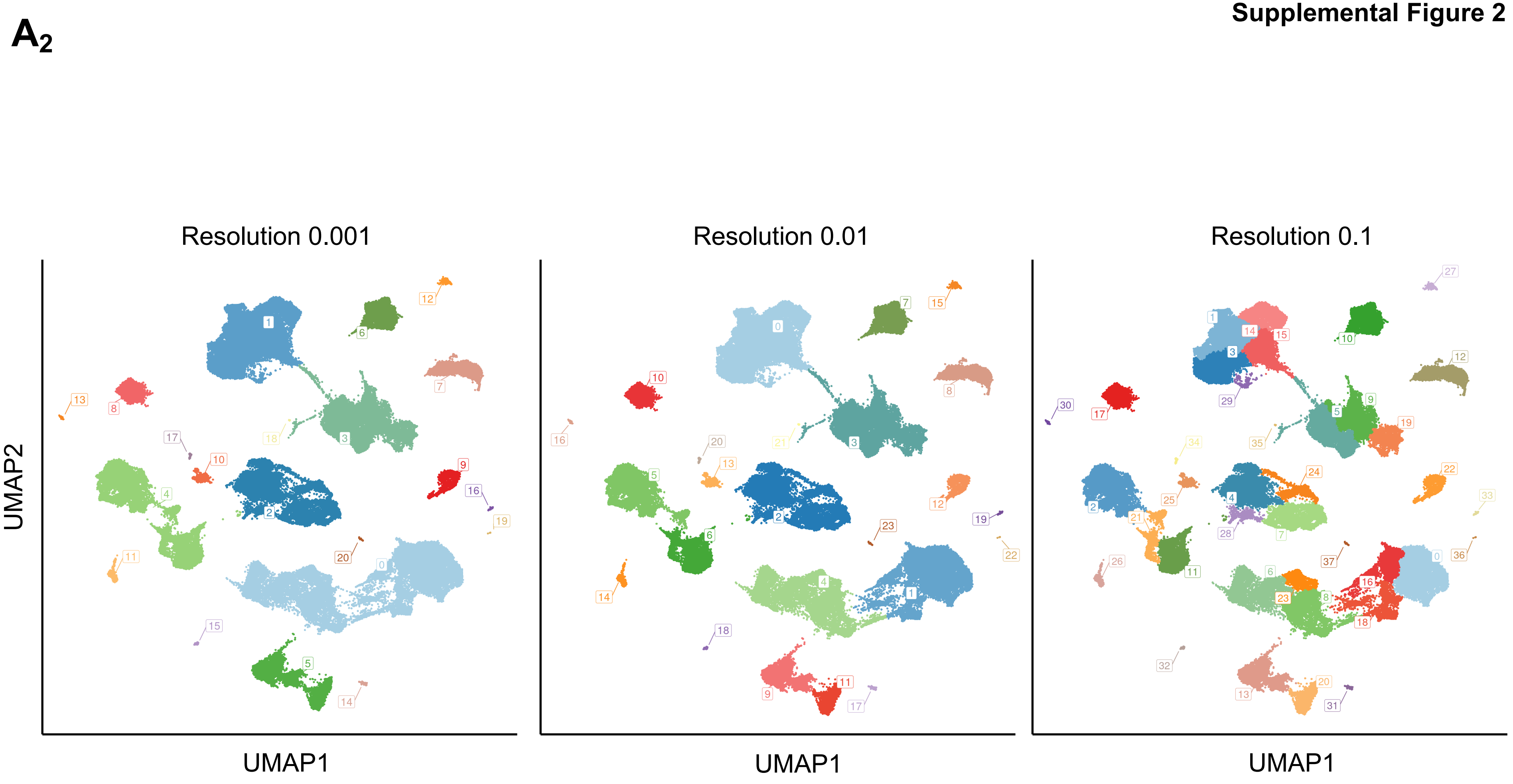
A2
Resolution 0.001
Resolution 0.01
Resolution 0.1
UMAP2
UMAP1
UMAP1
UMAP1

**Supplemental Figure 2**

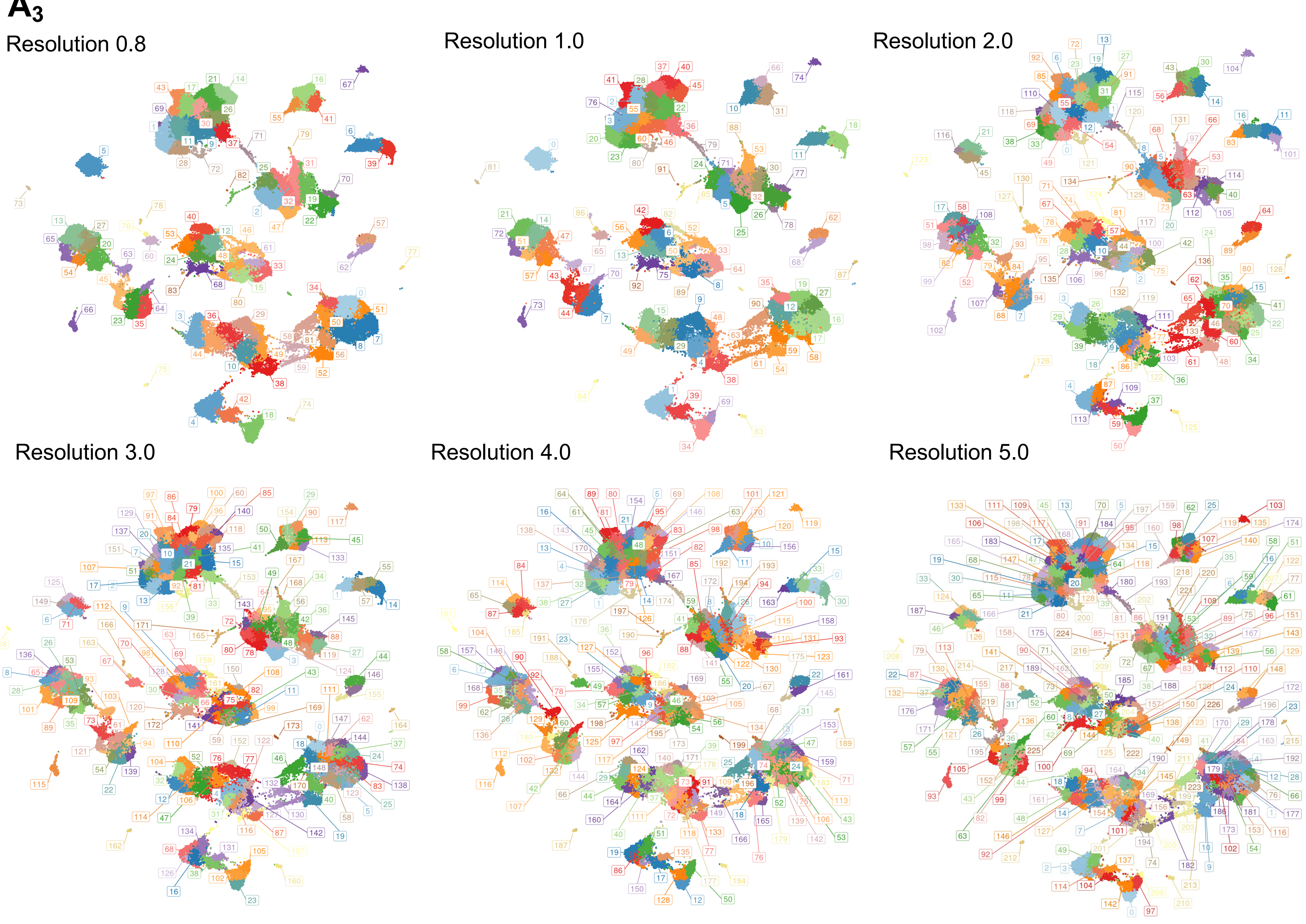

Supplemental Figure 2

**B**

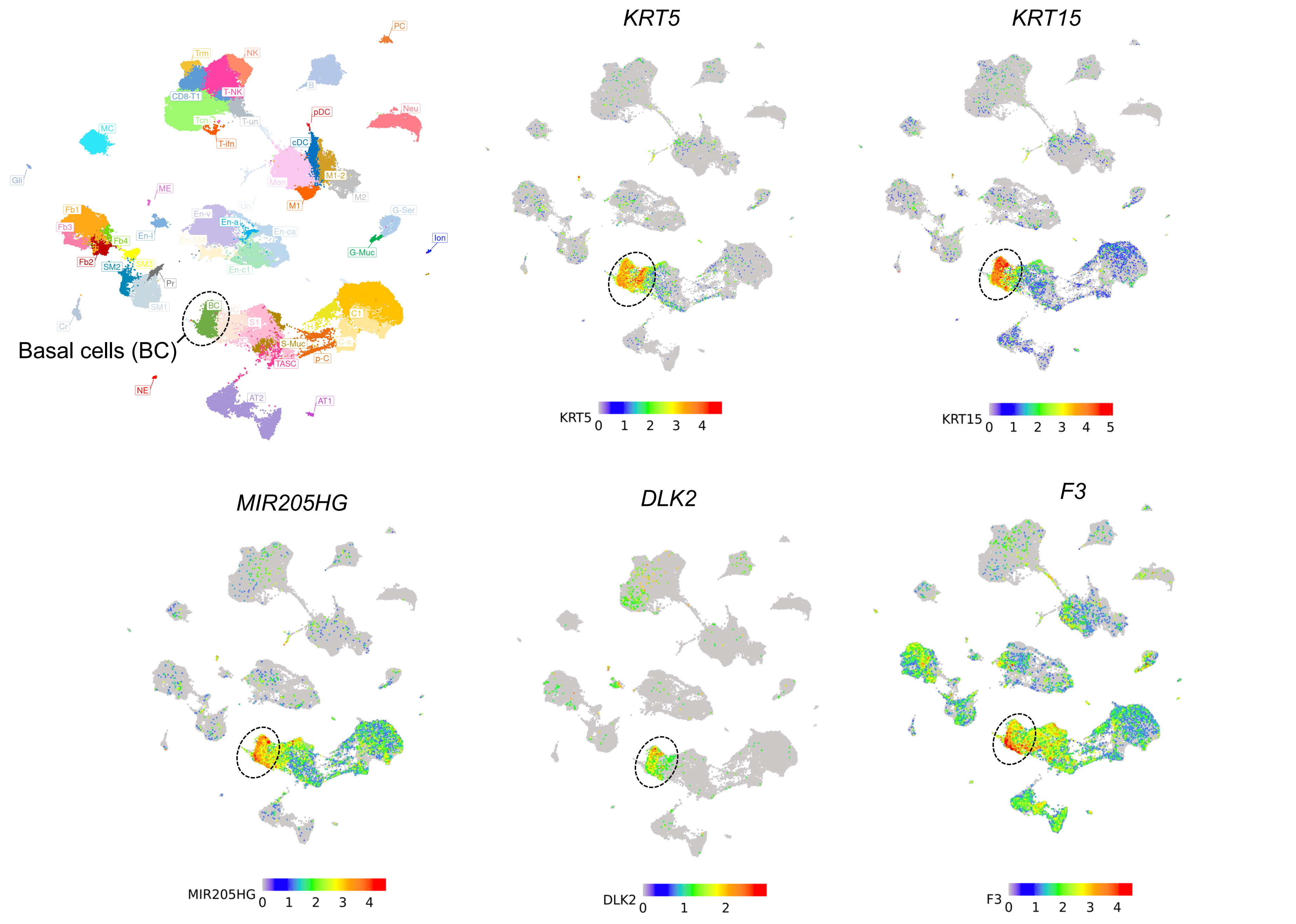

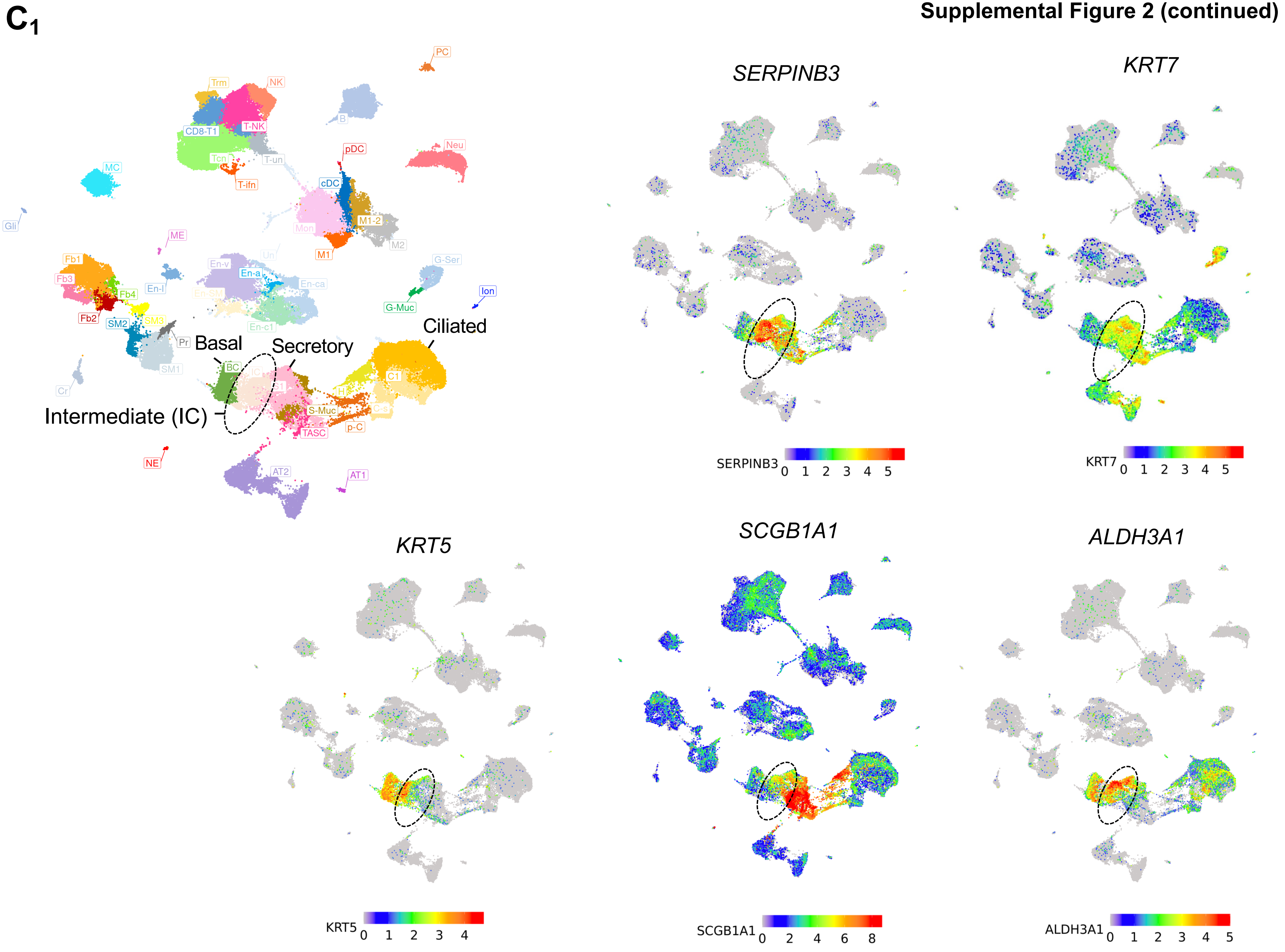
C1
PC
Trm
NK
B
CD8-T1
T-NK
Neu
pDC
Tcn
T-un
T-ifn
MC
cDC
Gli
M1-2
Mon
M2
ME
M1
Un
En-v
G-Ser
Fb1
Fb3
En-a
En-ca
En-l
Fb4
Ion
G-Muc
Fb2
SM2
SM3
En-c1
Ciliated
Pr
Basal
Secretory
BC
IC
SM1
C1
Cr
H
Intermediate (IC)
S-Muc
C-s
TASC
p-C
NE
AT2
AT1
SERPINB3
KRT7
KRT5
SCGB1A1
ALDH3A1

**Supplemental Figure 2 (continued)**

**$C_2$**

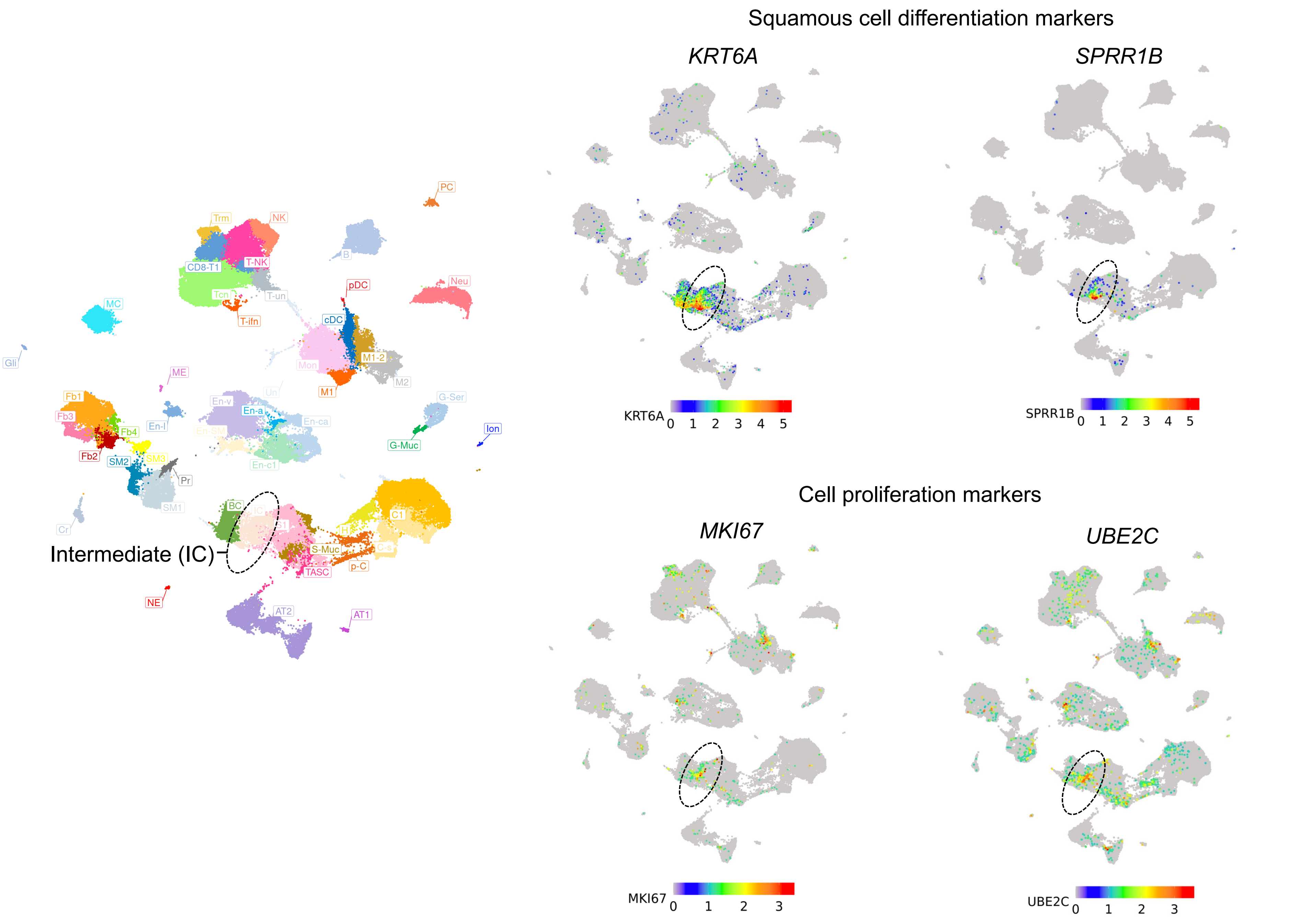

# C$_3$

Undifferentiated airway surface epithelial cells -
Basal cells (BC) + Intermediate cells (IC)

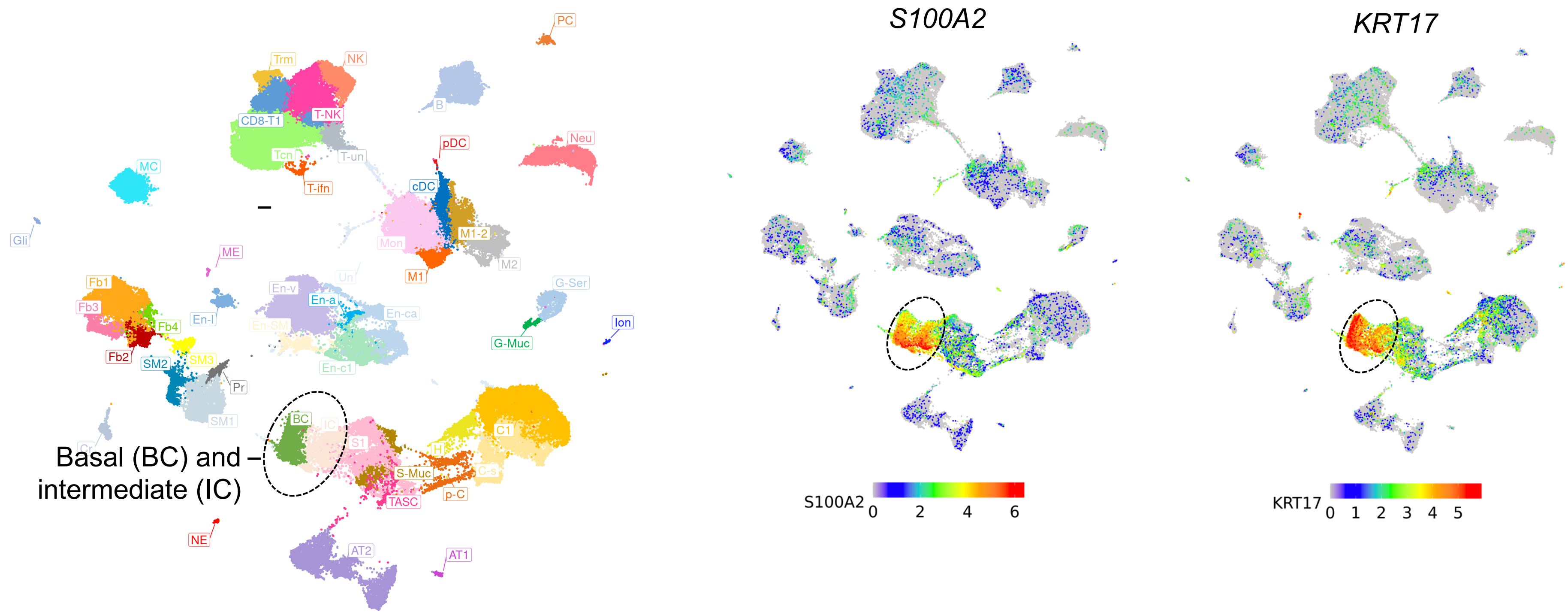

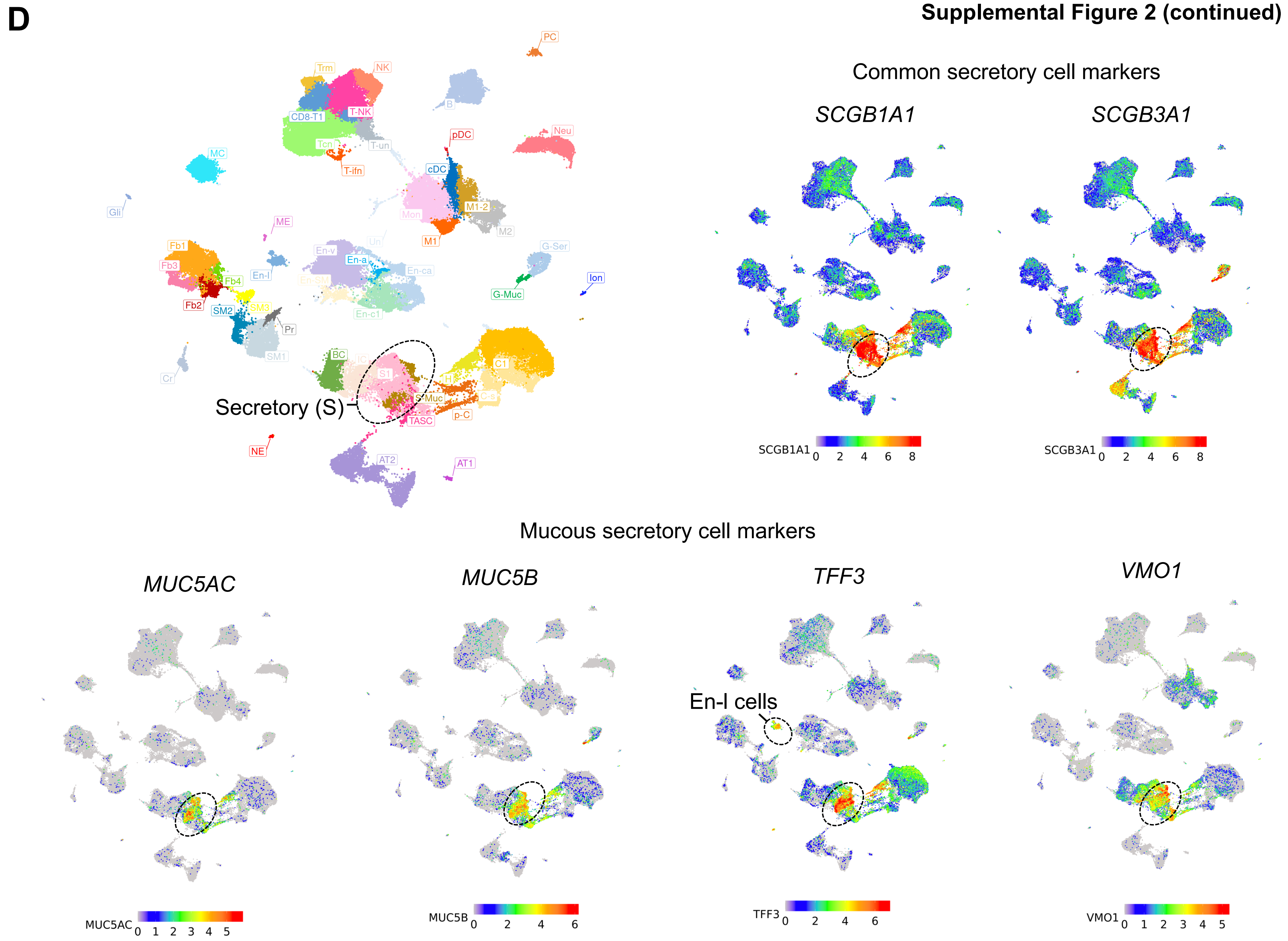
D
Common secretory cell markers
SCGB1A1
SCGB3A1
Secretory (S)
Mucous secretory cell markers
MUC5AC
MUC5B
TFF3
VMO1
En-l cells
PC
Trm
NK
CD8-T1
T-NK
B
Neu
pDC
T-un
Tcn
T-ifn
MC
cDC
Gli
Mon
M1-2
M2
M1
ME
Fb1
Fb3
Fb4
Fb2
En-l
En-v
En-a
En-ca
En-SM
En-c1
G-Ser
G-Muc
Ion
SM2
SM3
SM1
Pr
Cr
BC
S1
C1
H
S-Muc
C-s
TASC
p-C
NE
AT2
AT1
SCGB1A1 0 2 4 6 8
SCGB3A1 0 2 4 6 8
MUC5AC 0 1 2 3 4 5
MUC5B 0 2 4 6
TFF3 0 2 4 6
VMO1 0 1 2 3 4 5

**Supplemental Figure 2 (continued)**

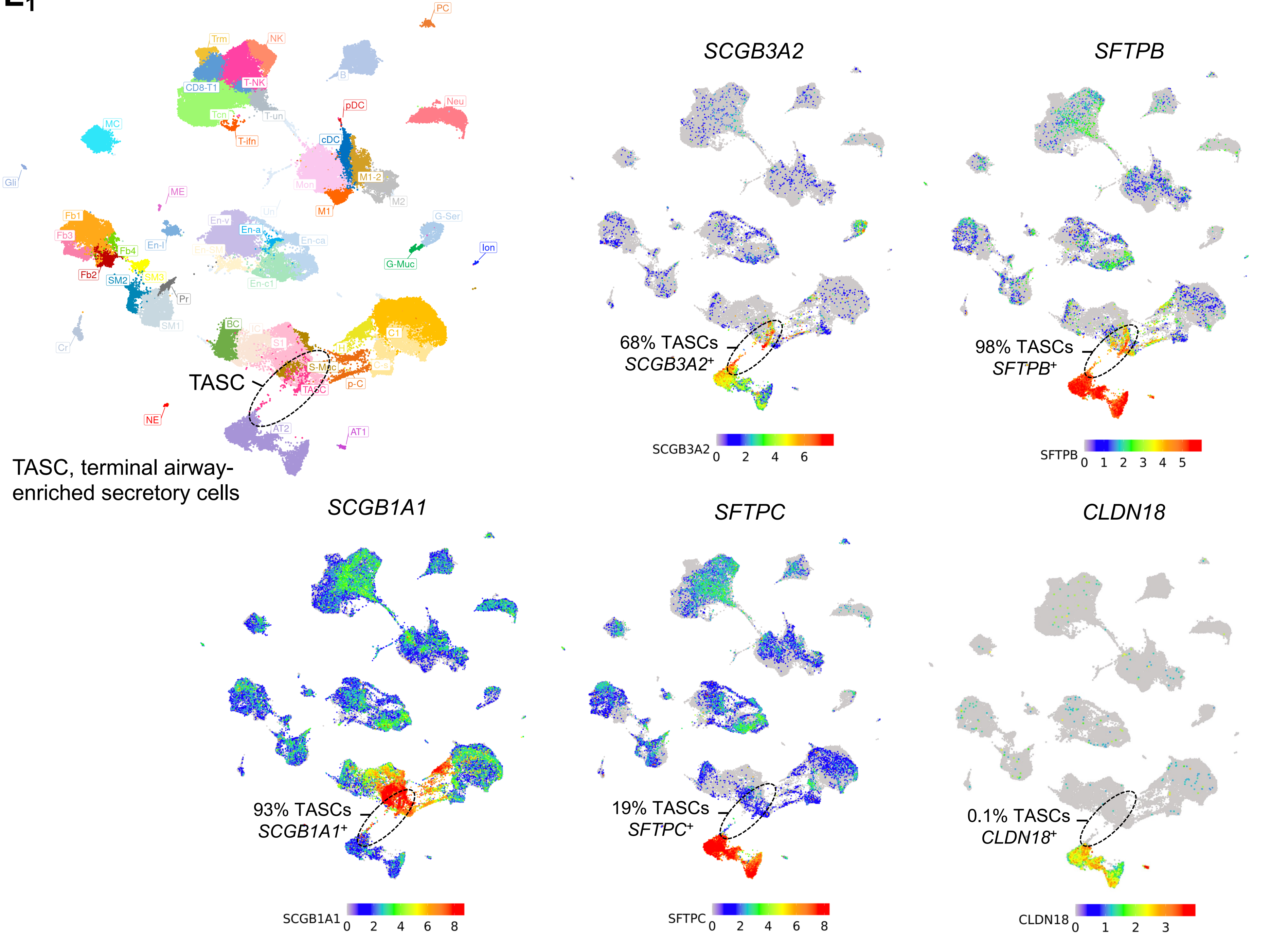

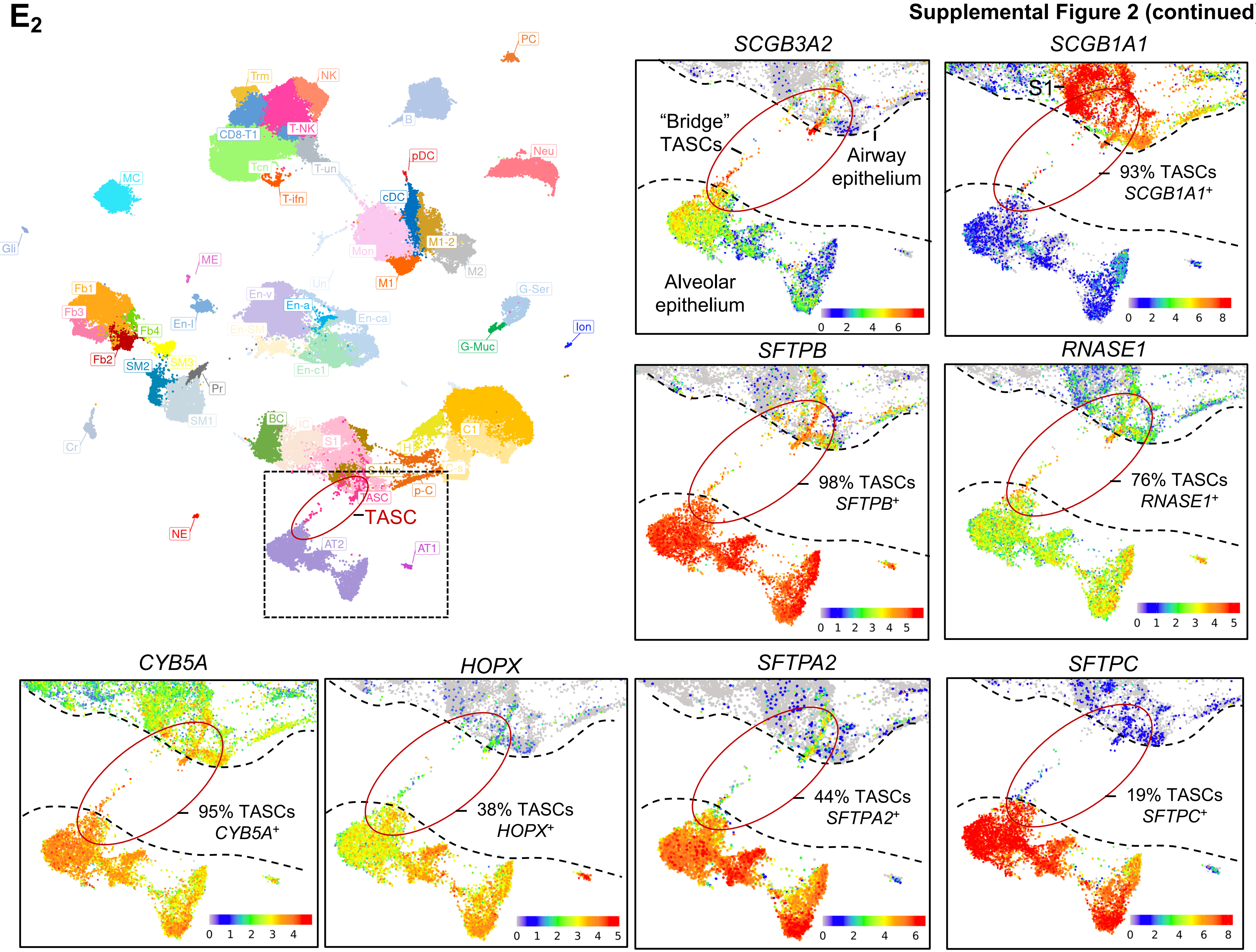
E2
PC
Trm
NK
B
CD8-T1
T-NK
Neu
pDC
Tcn
T-un
MC
T-ifn
cDC
M1-2
Gli
Mon
M2
ME
M1
Un
G-Ser
Fb1
En-v
En-a
En-ca
Fb3
En-l
Ion
Fb4
En-SM
G-Muc
SM3
Fb2
SM2
En-c1
Pr
SM1
BC
IC
C1
Cr
S1
H
S-Muc
C-s
p-C
TASC
TASC
NE
AT2
AT1
SCGB3A2
"Bridge" TASCs
Airway epithelium
Alveolar epithelium
SCGB1A1
S1
93% TASCs SCGB1A1+
SFTPB
98% TASCs SFTPB+
RNASE1
76% TASCs RNASE1+
CYB5A
95% TASCs CYB5A+
HOPX
38% TASCs HOPX+
SFTPA2
44% TASCs SFTPA2+
SFTPC
19% TASCs SFTPC+

**$F_1$**



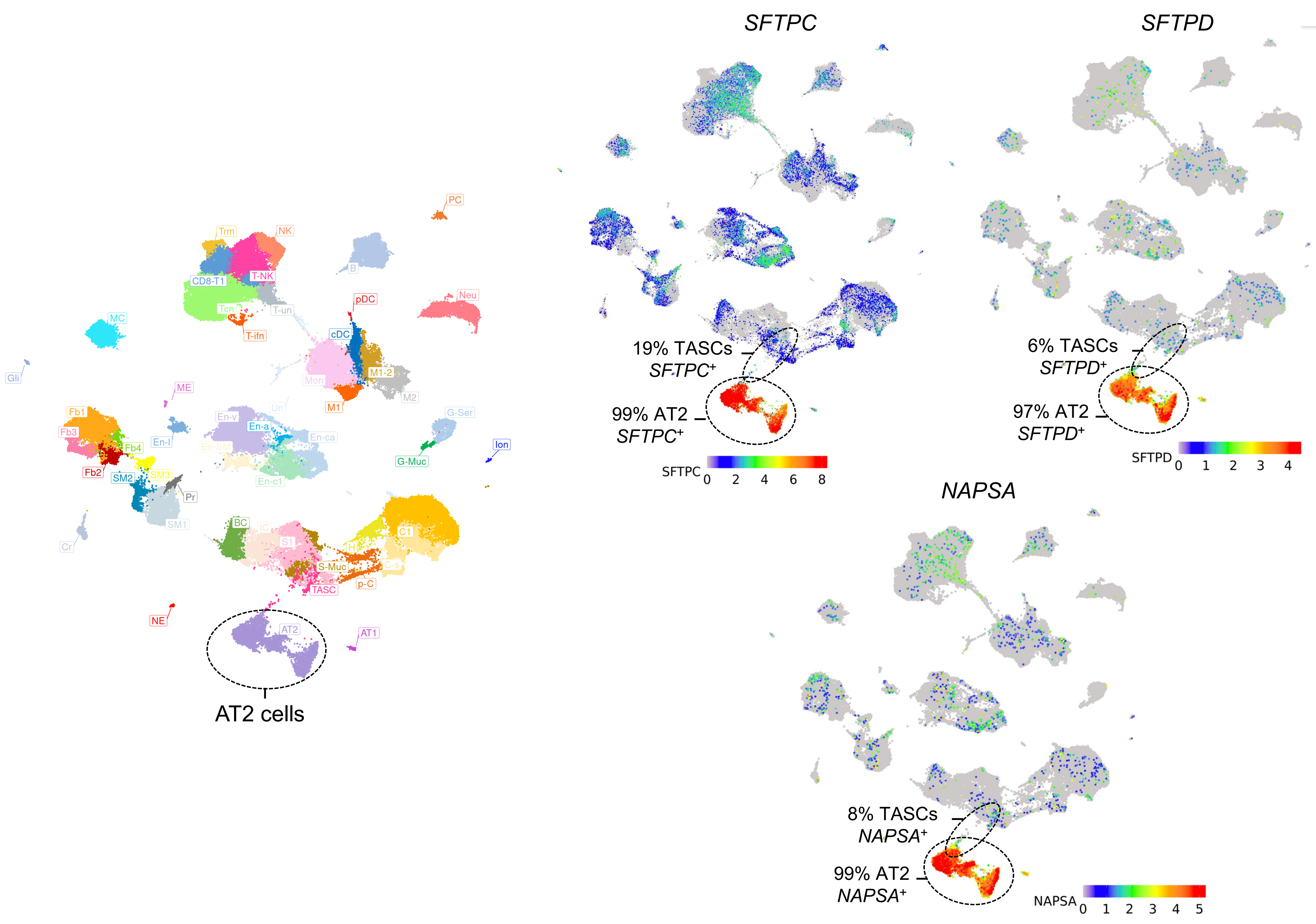
PC
Trm
NK
B
CD8-T1
T-NK
pDC
Neu
MC
Tcn
T-un
T-ifn
cDC
Gli
ME
Mon
M1-2
M2
M1
Un
En-v
Fb1
Fb3
En-a
En-ca
G-Ser
En-l
En-SM
Fb4
Ion
G-Muc
Fb2
SM2
SM3
Pr
En-c1
BC
IC
C1
S1
H
S-Muc
C-s
TASC
p-C
NE
AT2
AT1
Cr
SM1
AT2 cells
SFTPC
19% TASCs SFTPC+
99% AT2 SFTPC+
SFTPC 0 2 4 6 8
SFTPD
6% TASCs SFTPD+
97% AT2 SFTPD+
SFTPD 0 1 2 3 4
NAPSA
8% TASCs NAPSA+
99% AT2 NAPSA+
NAPSA 0 1 2 3 4 5

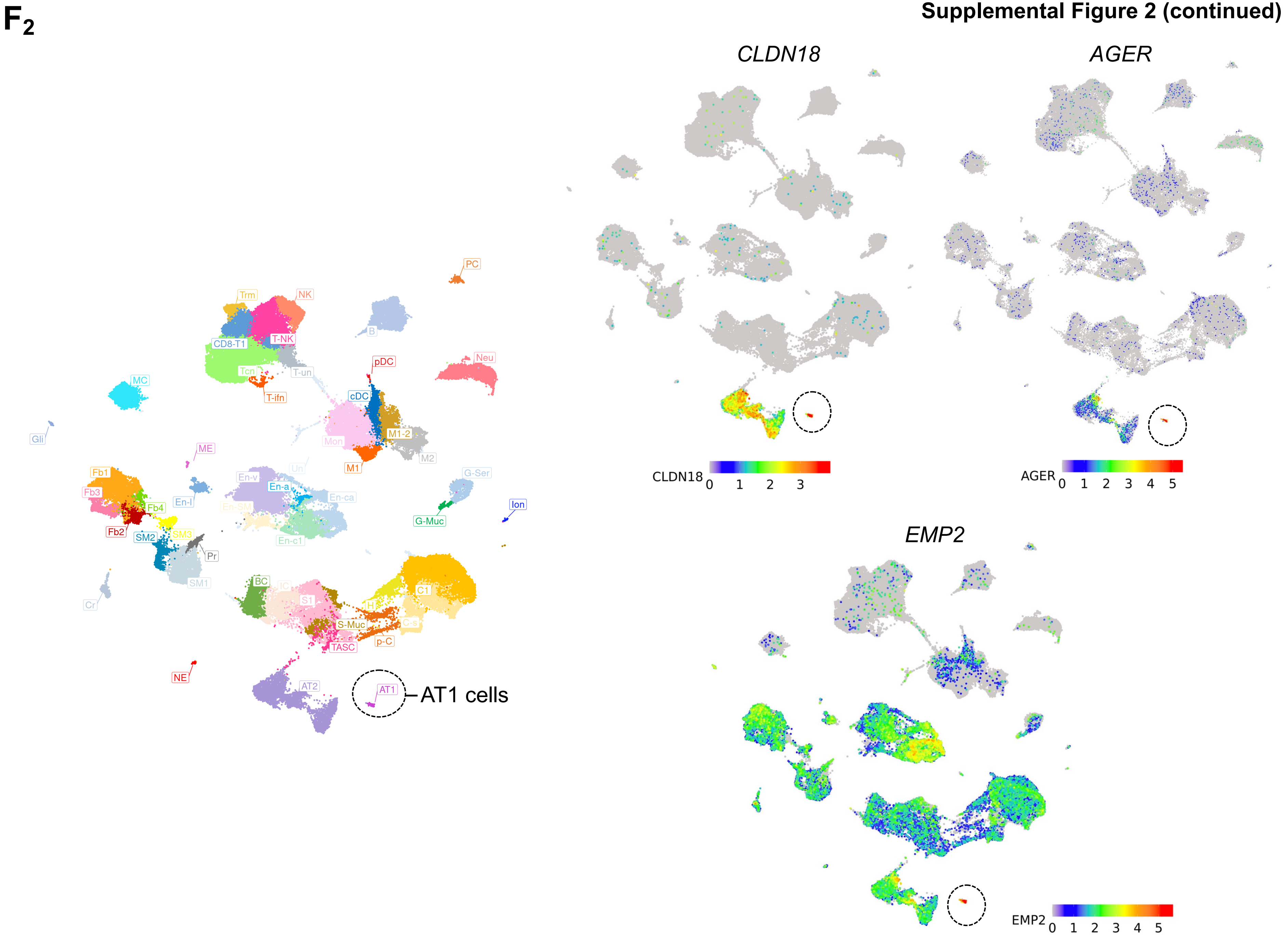
F2
CLDN18
AGER
EMP2
AT1 cells
PC
Trm
NK
B
CD8-T1
T-NK
pDC
Neu
MC
Tcn
T-un
T-ifn
cDC
Gli
M1-2
Mon
ME
M2
M1
Un
Fb1
En-v
G-Ser
En-a
Fb3
En-ca
En-l
Ion
Fb4
En-SM
G-Muc
Fb2
SM2
SM3
En-c1
Pr
BC
IC
SM1
C1
S1
Cr
H
C-s
S-Muc
p-C
TASC
NE
AT2
AT1
CLDN18 0 1 2 3
AGER 0 1 2 3 4 5
EMP2 0 1 2 3 4 5

**Supplemental Figure 2 (continued)**

**$G_1$**

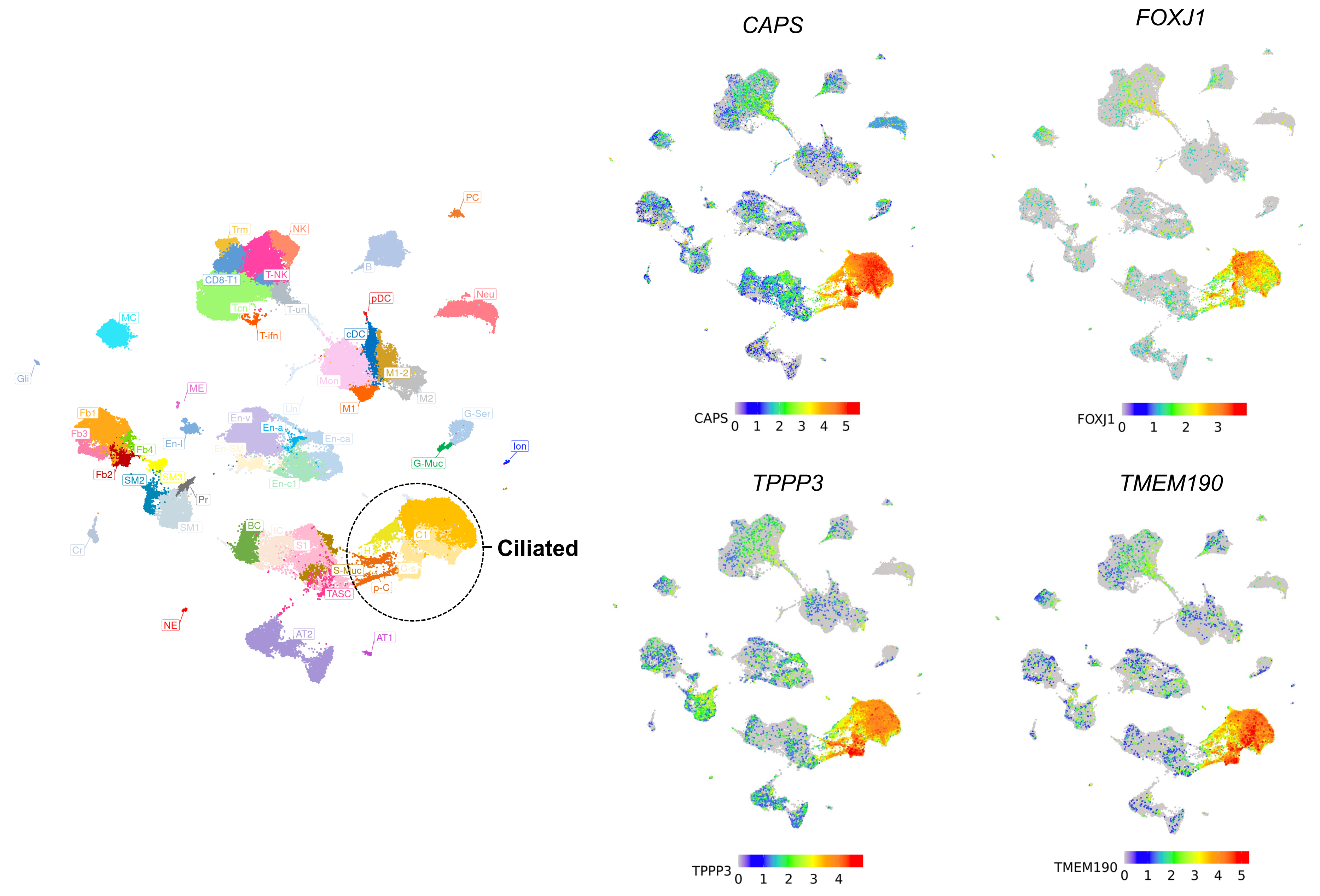

G$_2$

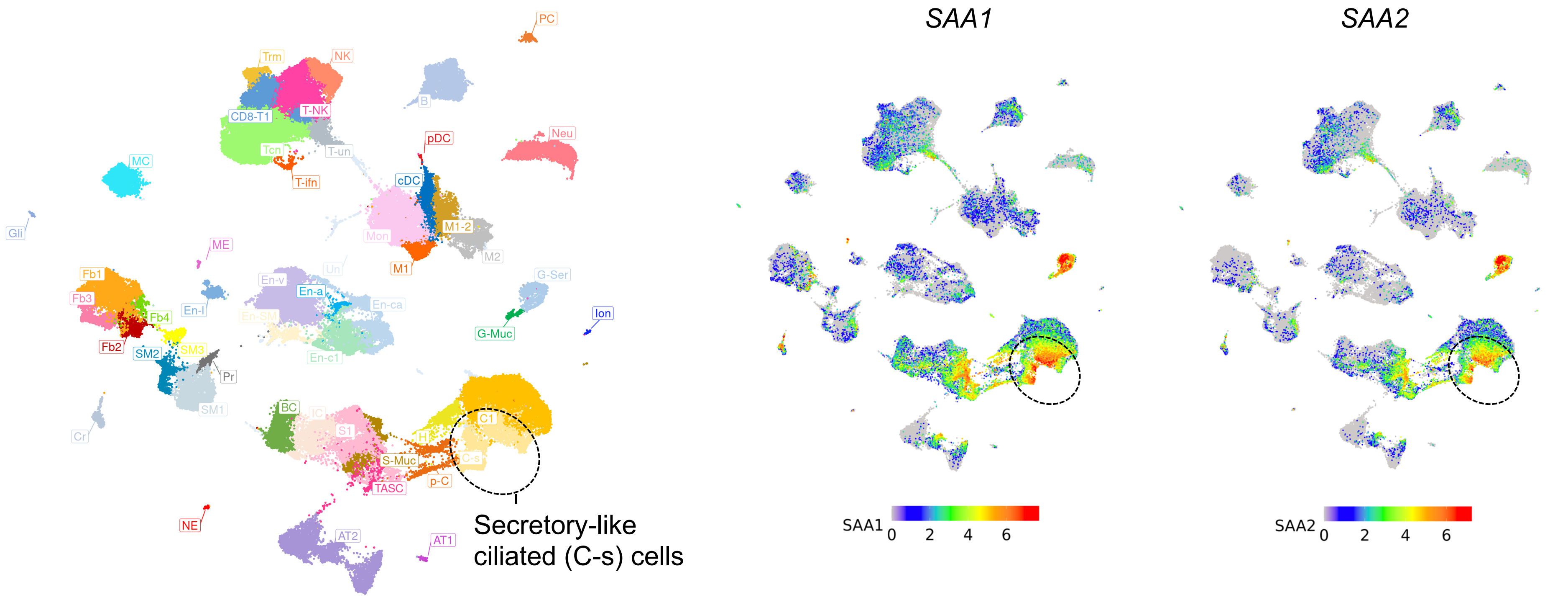
PC
Trm
NK
B
CD8-T1
T-NK
Neu
pDC
Tcn
T-un
MC
T-ifn
cDC
Gli
M1-2
Mon
ME
M2
M1
Un
Fb1
En-v
G-Ser
En-a
Fb3
En-ca
Ion
En-l
Fb4
En-SM
G-Muc
Fb2
SM3
SM2
En-c1
Pr
SM1
BC
IC
C1
Cr
S1
H
C-s
S-Muc
p-C
TASC
NE
Secretory-like
ciliated (C-s) cells
AT2
AT1
SAA1
SAA2
SAA1
0
2
4
6
SAA2
0
2
4
6

**Supplemental Figure 2 (continued)**

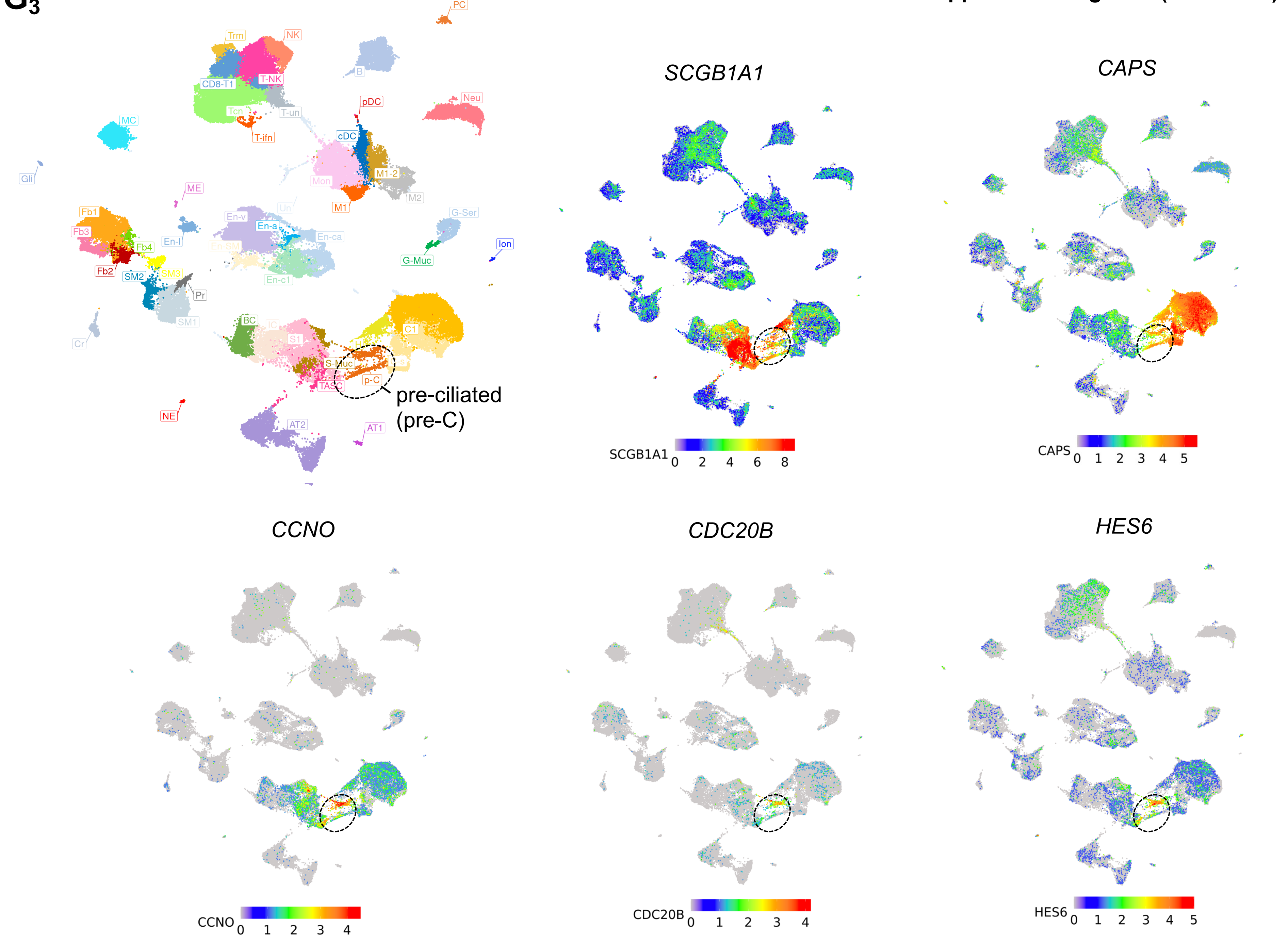

**$G_4$**

**S-C hybrid (H) cells:**
single cells co-expressing C1, S1, and S-Muc markers
C1 markers: *CAPS, FOXJ1*
S1 markers: *SCGB1A1, SCGB3A1*
S-Muc markers: *TFF3, MUC5AC*

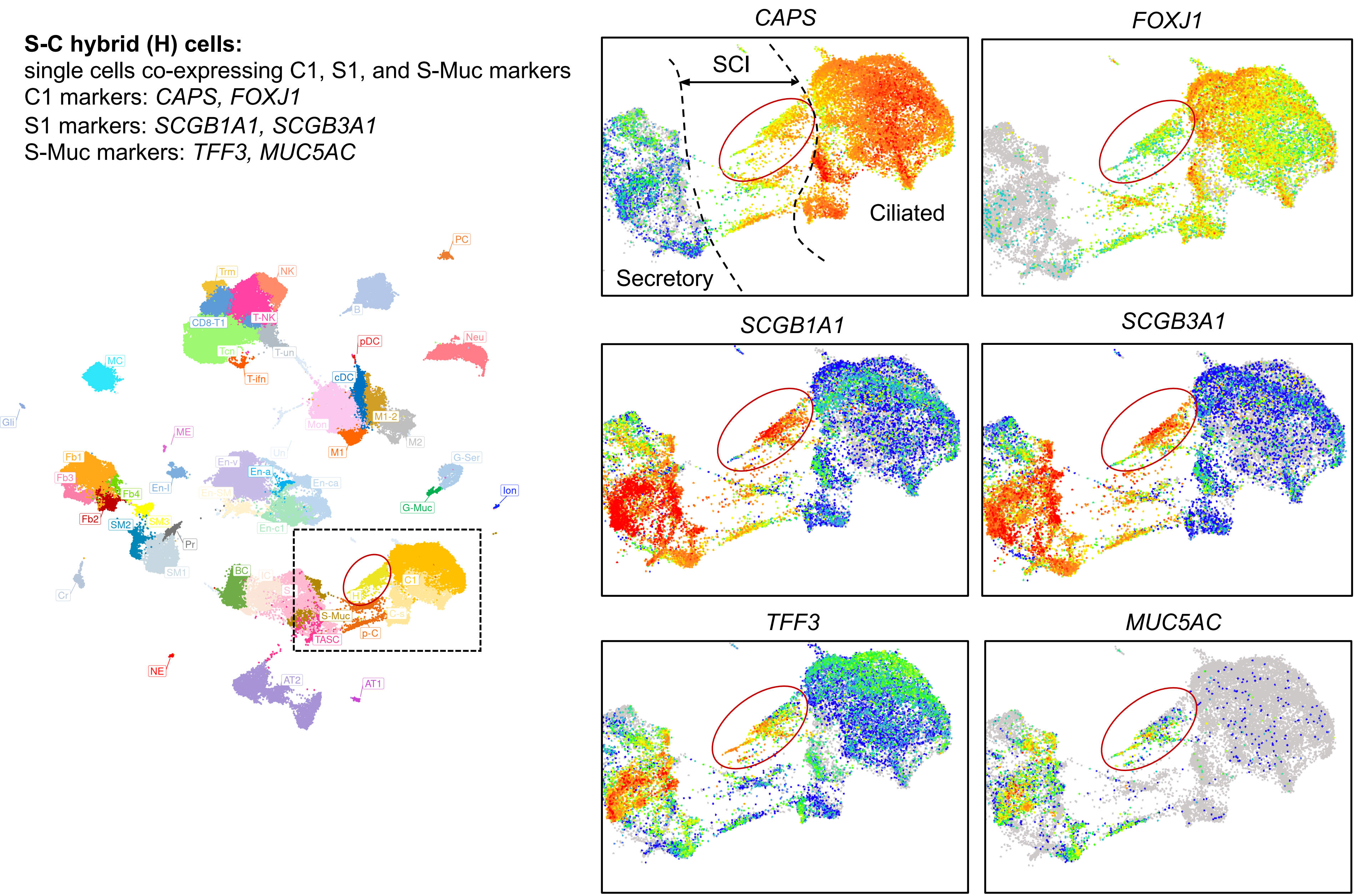

H

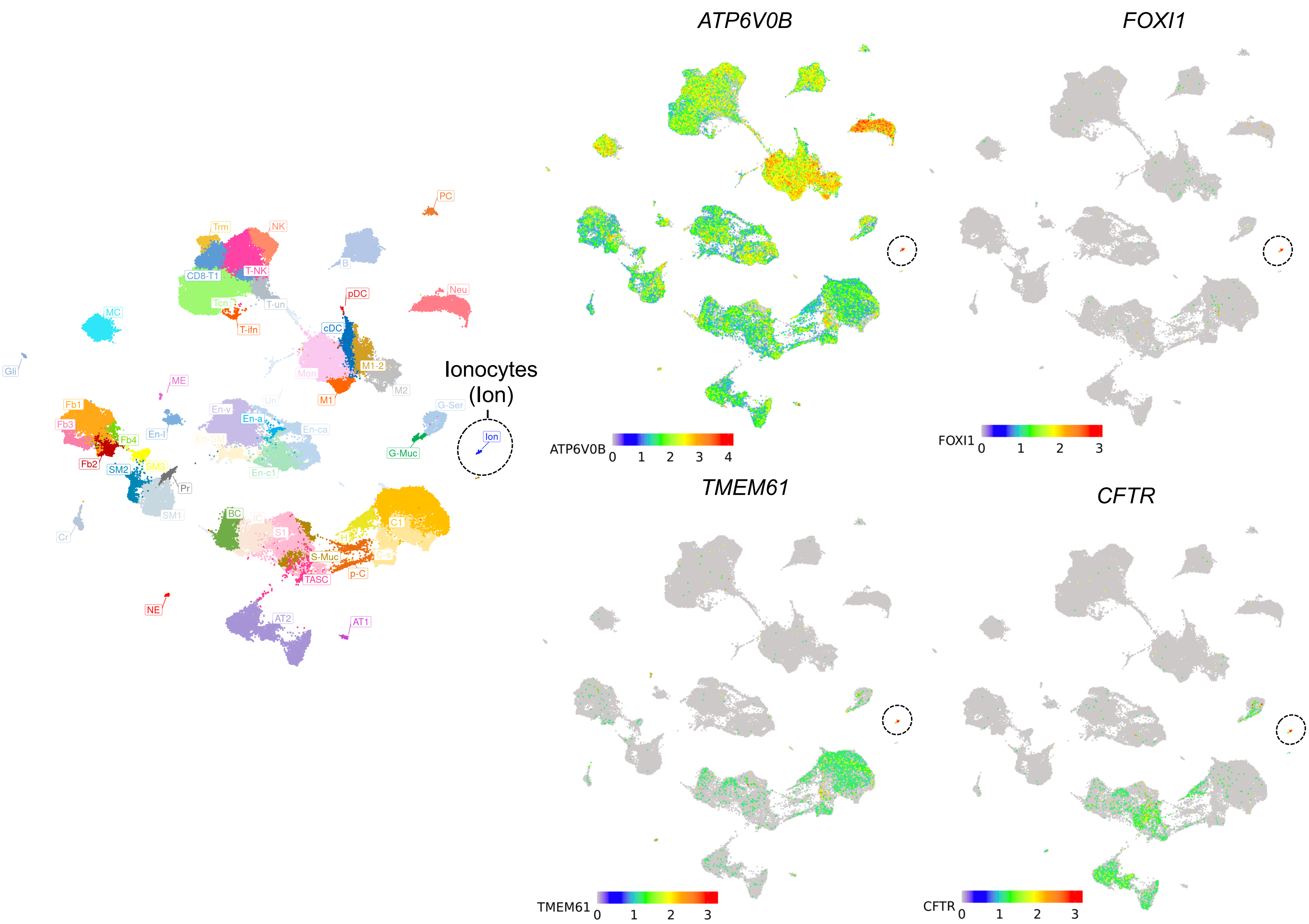

PC
Trm
NK
B
CD8-T1
T-NK
Neu
pDC
MC
Tcn
T-un
T-ifn
cDC
Gli
ME
Mon
M1-2
M2
M1
Ionocytes
(Ion)
Fb1
Fb3
Fb4
En-l
En-v
En-a
Un
En-ca
En-SM
En-c1
G-Ser
G-Muc
Ion
Fb2
SM2
SM3
Pr
SM1
Cr
BC
IC
S1
C1
H
S-Muc
C-s
p-C
TASC
NE
AT2
AT1
ATP6V0B
FOXI1
TMEM61
CFTR
0
1
2
3
4

Supplemental Figure 2 (continued)

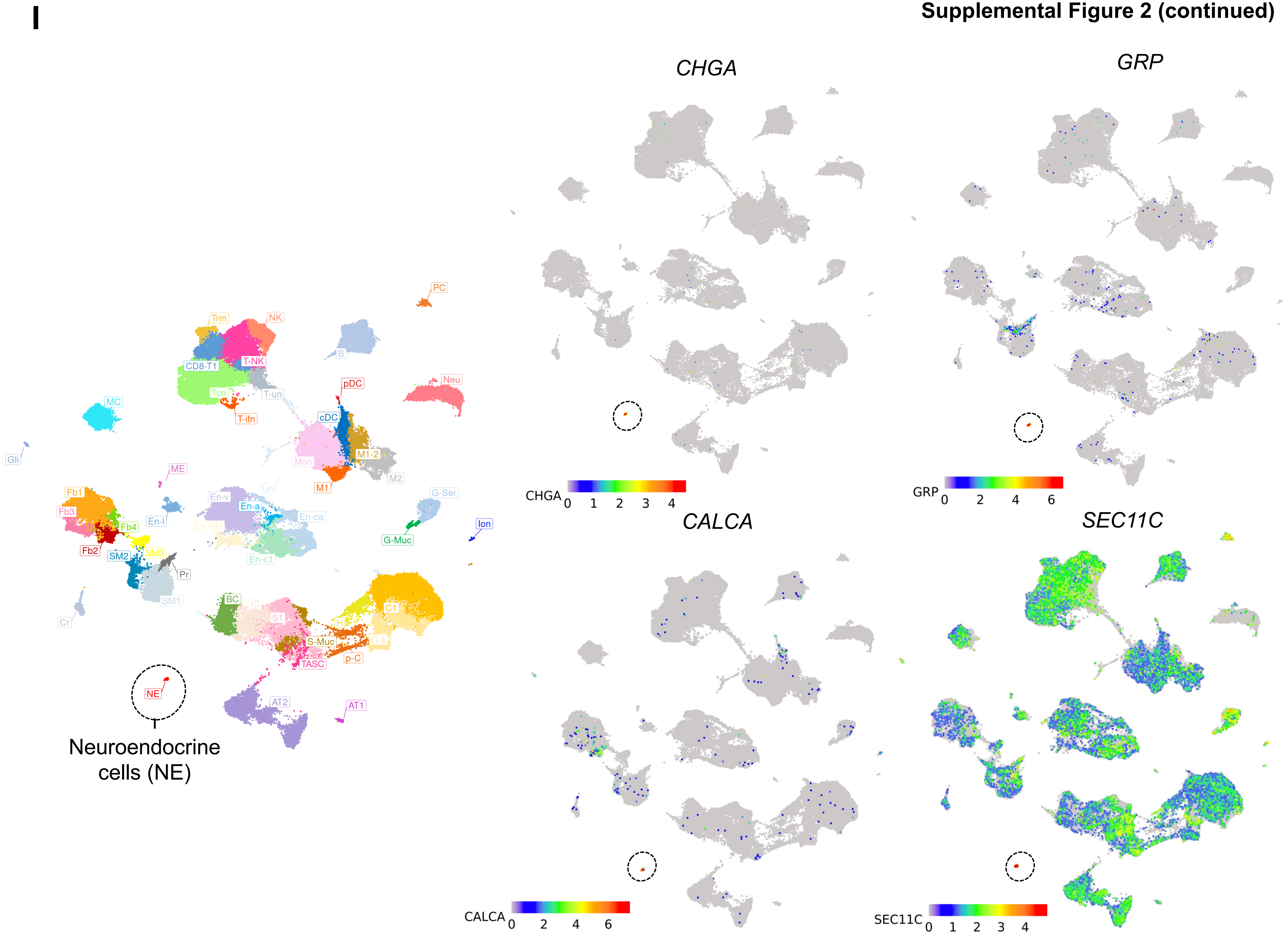

**J$_1$**

**Submucosal gland (SMG) cells**

- **Myoepithelial cells (ME)**
  *KRT5⁺ KRT14⁺ ACTA2⁺*
- **Glandular secretory:** see next page

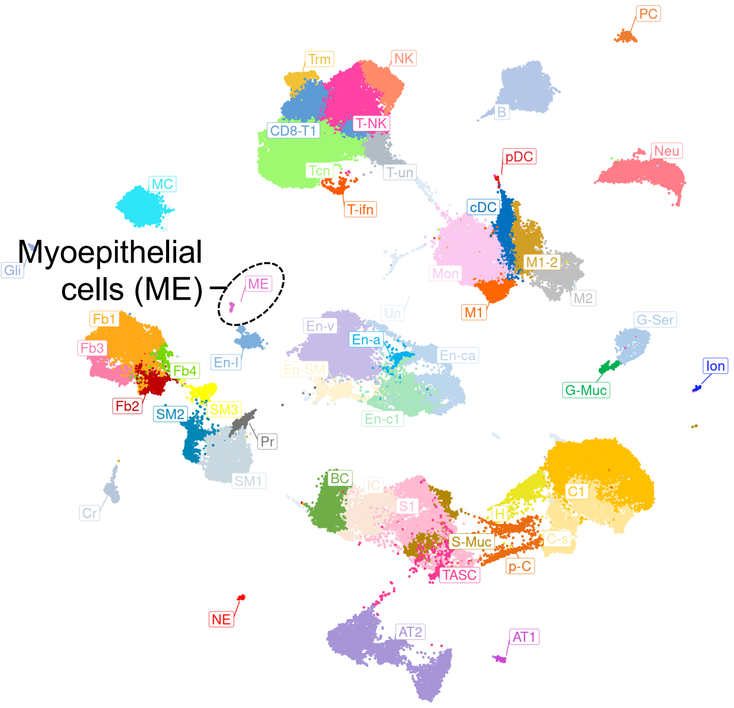


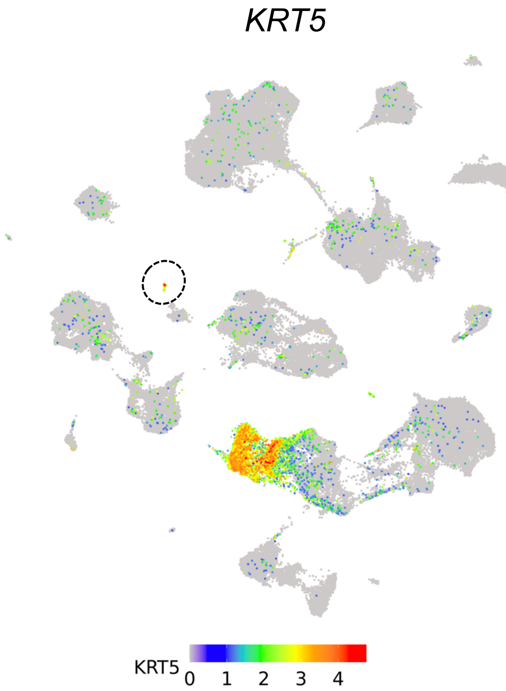


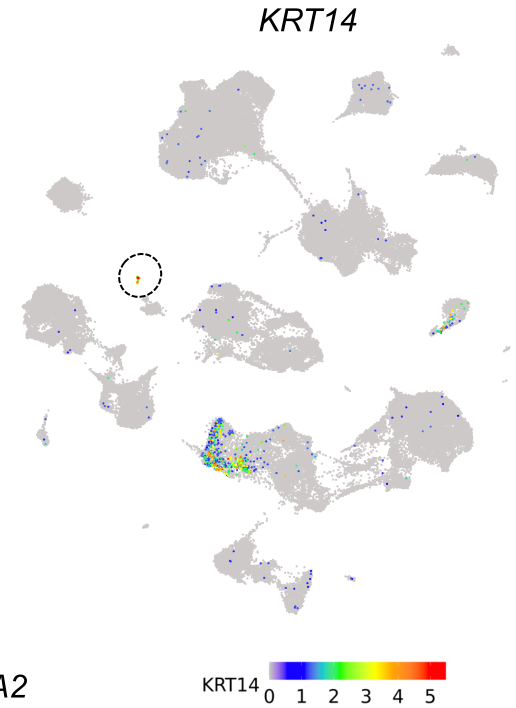


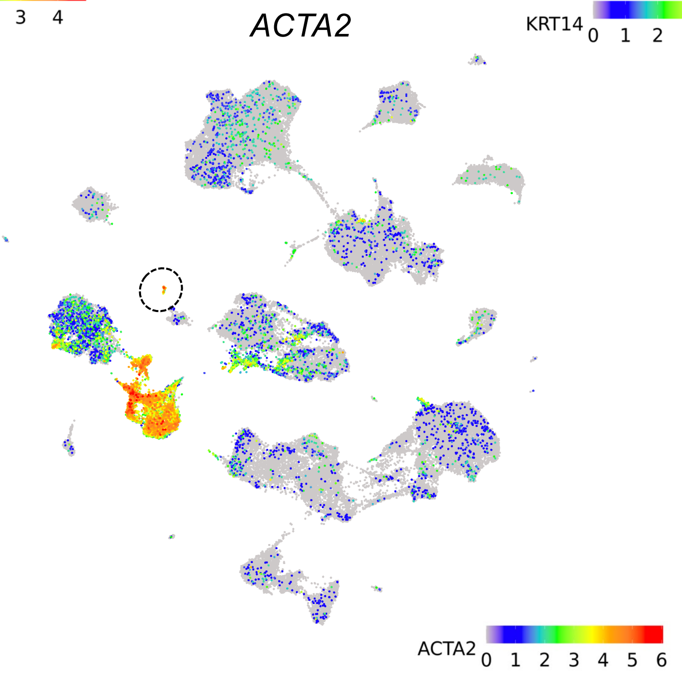

**J$_2$**

**Submucosal gland (SMG) cells**

- **Myoepithelial cells (ME)**
  see previous page
- **Glandular secretory:** *AZGP1*$^+$
  - Mucous (g-Muc): *MUC5B*$^{hi}$ *BPIFB2*$^{hi}$
  - Serous (g-Ser): *LYZ*$^{hi}$ *LTF*$^{hi}$ *PRR4*$^{hi}$

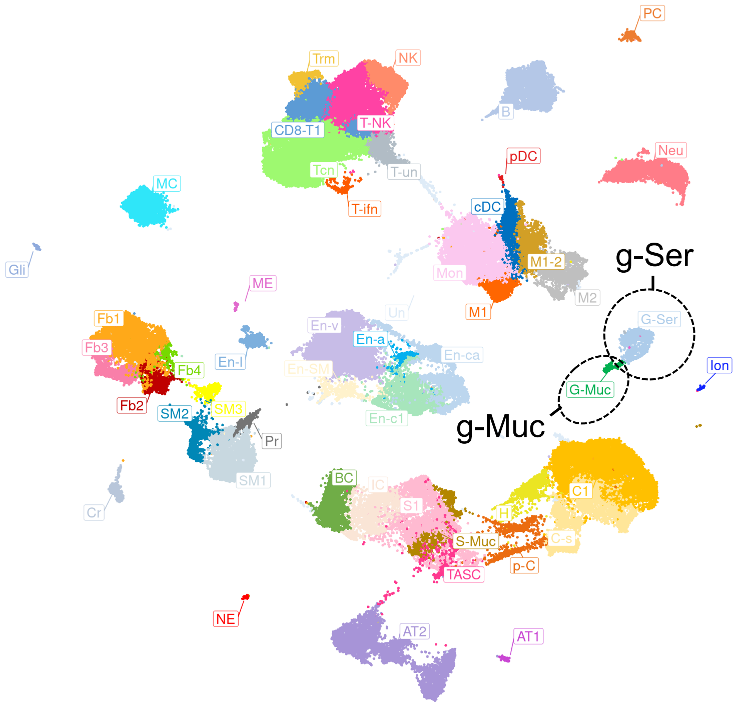


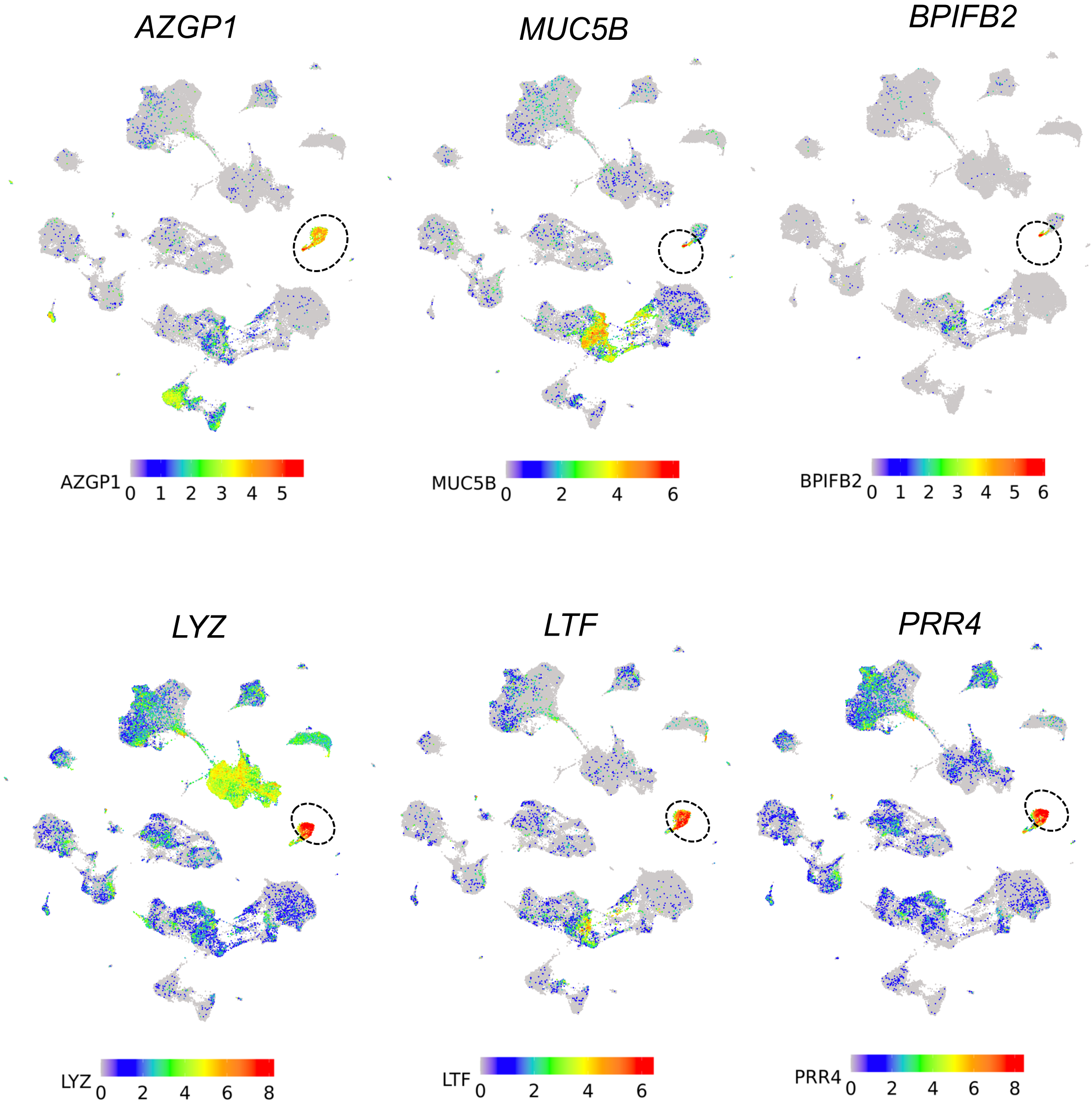

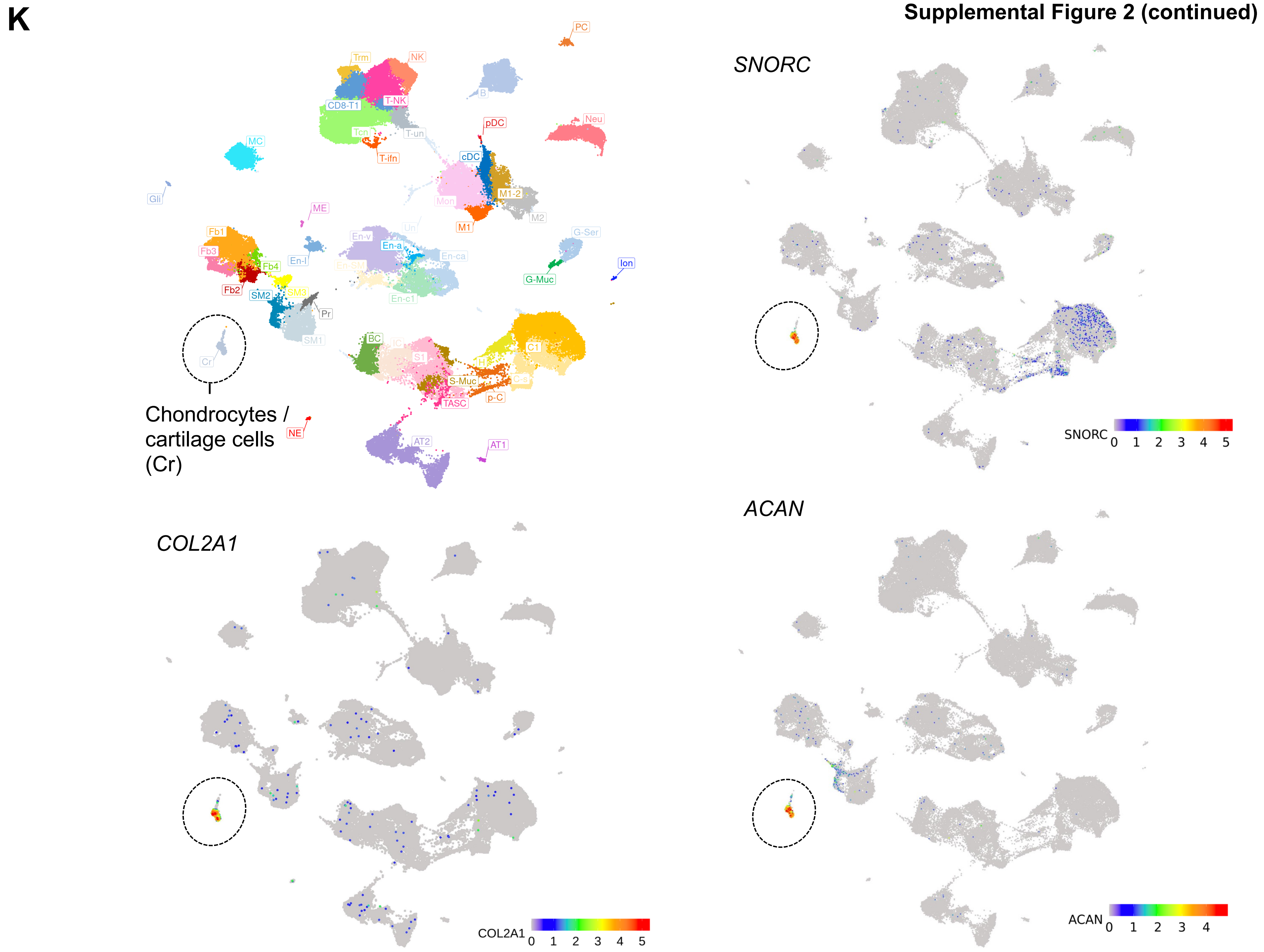
K
PC
Trm
NK
B
T-NK
CD8-T1
T-un
Tcn
pDC
Neu
MC
T-ifn
cDC
Gli
M1-2
Mon
M2
ME
M1
Un
G-Ser
Fb1
En-v
En-a
Fb3
En-ca
Ion
Fb4
En-l
En-SM
G-Muc
SM3
Fb2
SM2
En-c1
Pr
SM1
BC
IC
C1
S1
Cr
H
S-Muc
C-s
TASC
p-C
Chondrocytes / cartilage cells (Cr)
NE
AT2
AT1
SNORC
SNORC 0 1 2 3 4 5
COL2A1
COL2A1 0 1 2 3 4 5
ACAN
ACAN 0 1 2 3 4

**L1**

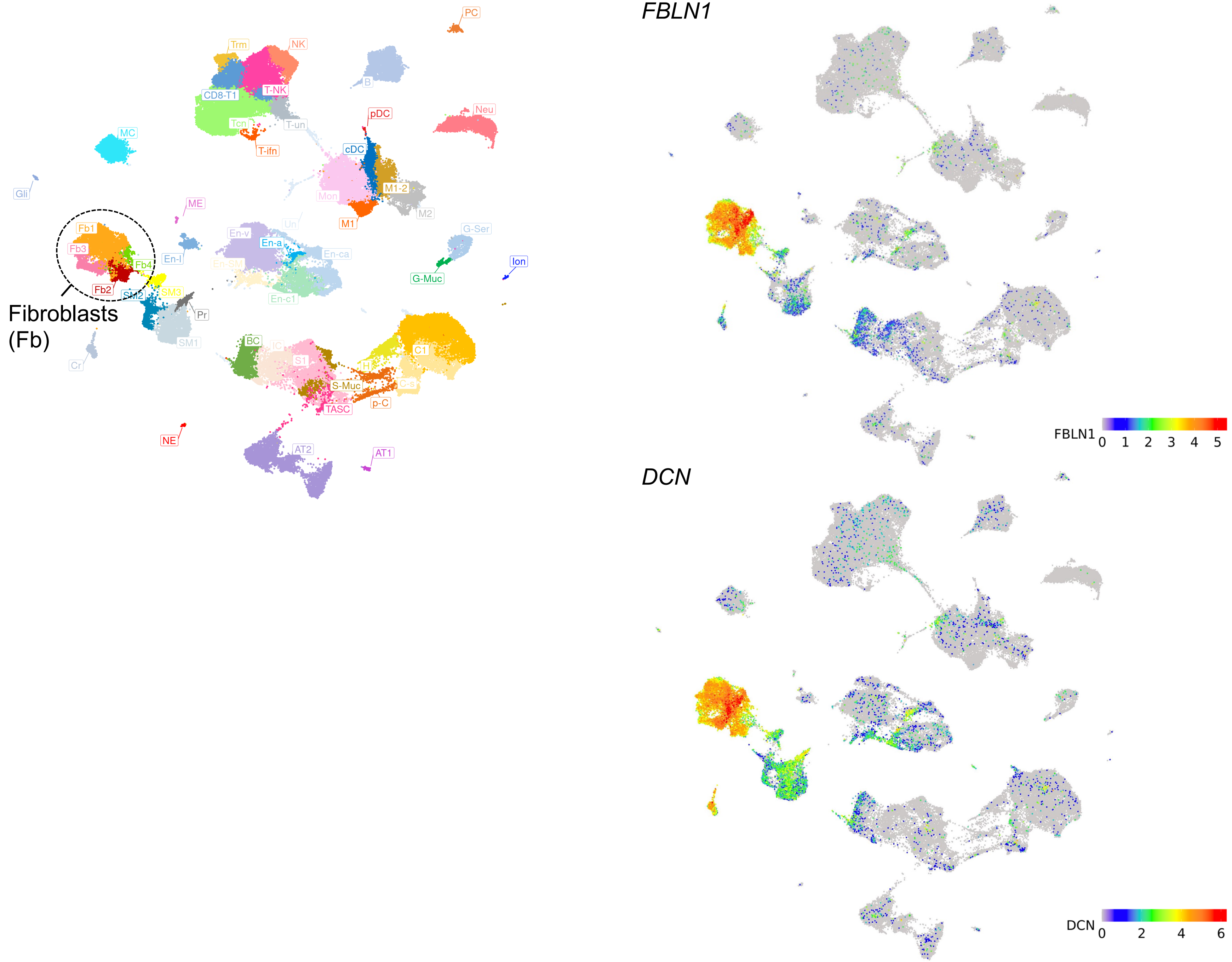
PC
Trm
NK
B
CD8-T1
T-NK
pDC
Neu
Tcn
T-un
T-ifn
MC
cDC
M1-2
Mon
M2
Gli
ME
M1
Un
G-Ser
Fb1
En-v
En-a
En-ca
Fb3
En-I
Fb4
En-SM
G-Muc
Ion
Fb2
SM2
SM3
En-c1
Pr
Fibroblasts
(Fb)
SM1
BC
IC
Cr
S1
C1
H
S-Muc
C-s
TASC
p-C
NE
AT2
AT1
FBLN1
FBLN1
0 1 2 3 4 5
DCN
DCN
0 2 4 6

**L$_2$**



Fibroblasts (Fb) subsets

Fb1: common
Fb2: *CCL19*$^{+}$, contains *SFRP2*$^{+}$

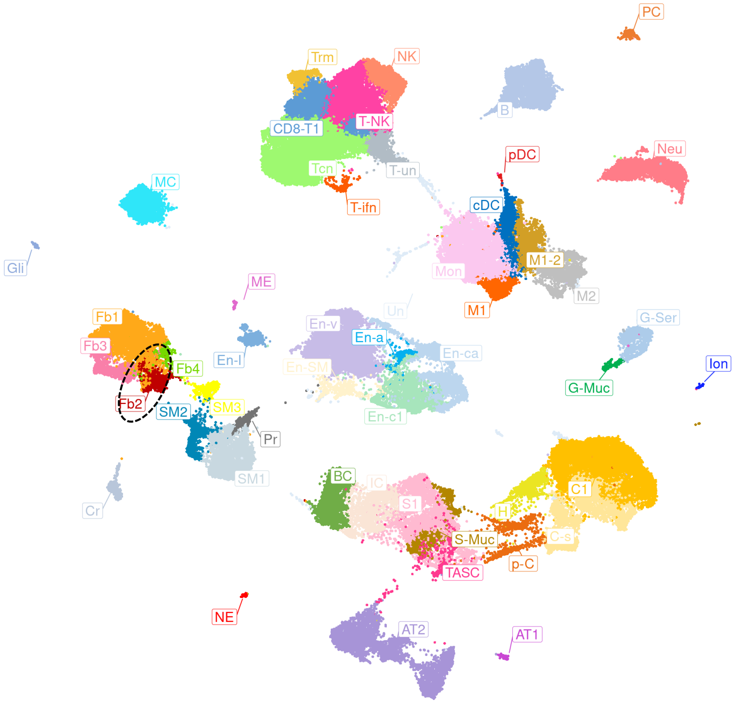


*CCL19*

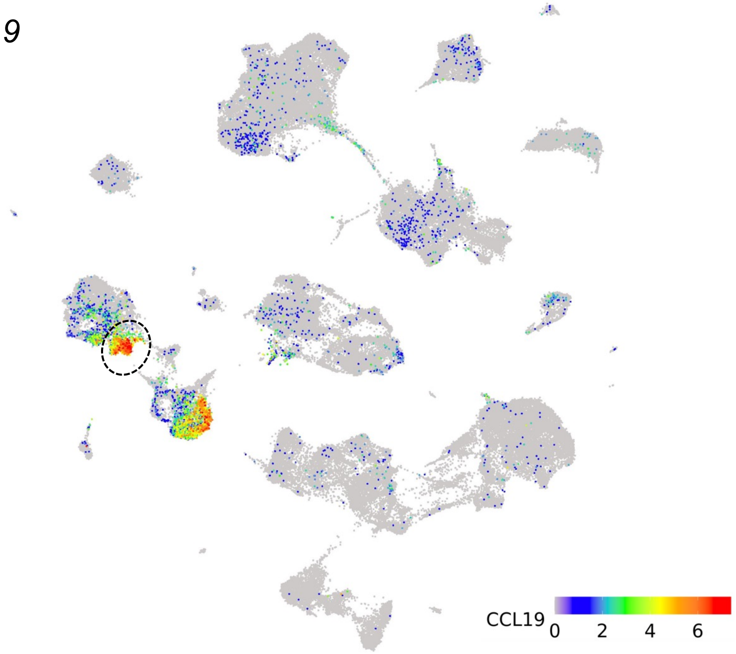


*SFRP2*

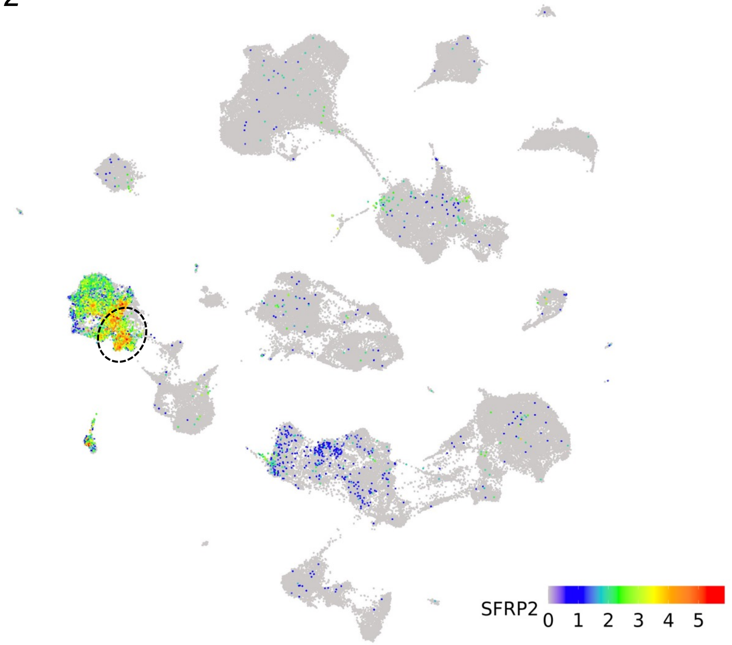

# L$_3$

Fibroblasts (Fb) subsets

Fb3 (matrix Fbs; contains myoFbs): contains *ELN*$^{+}$ *COL1A1*$^{+}$ *MMP2*$^{+}$ *ACTA2*$^{+}$

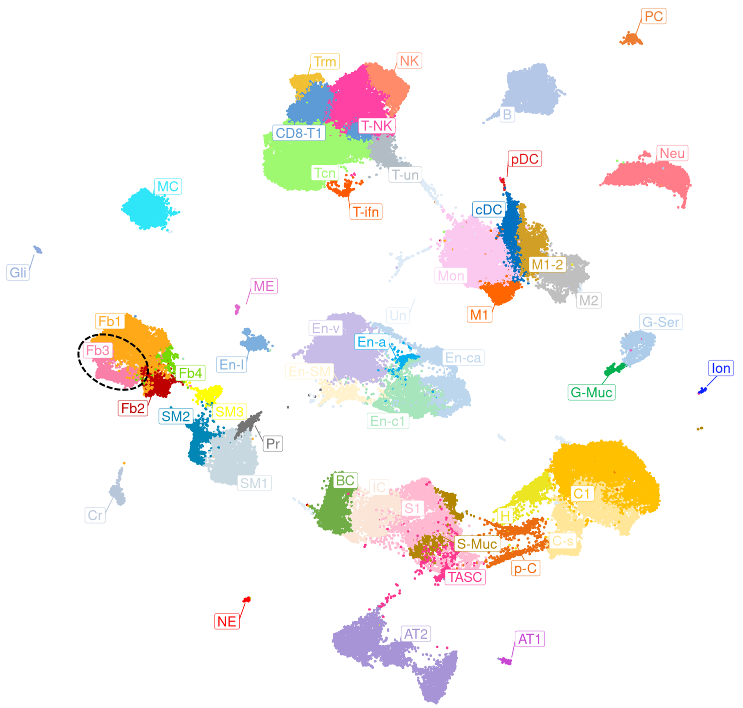


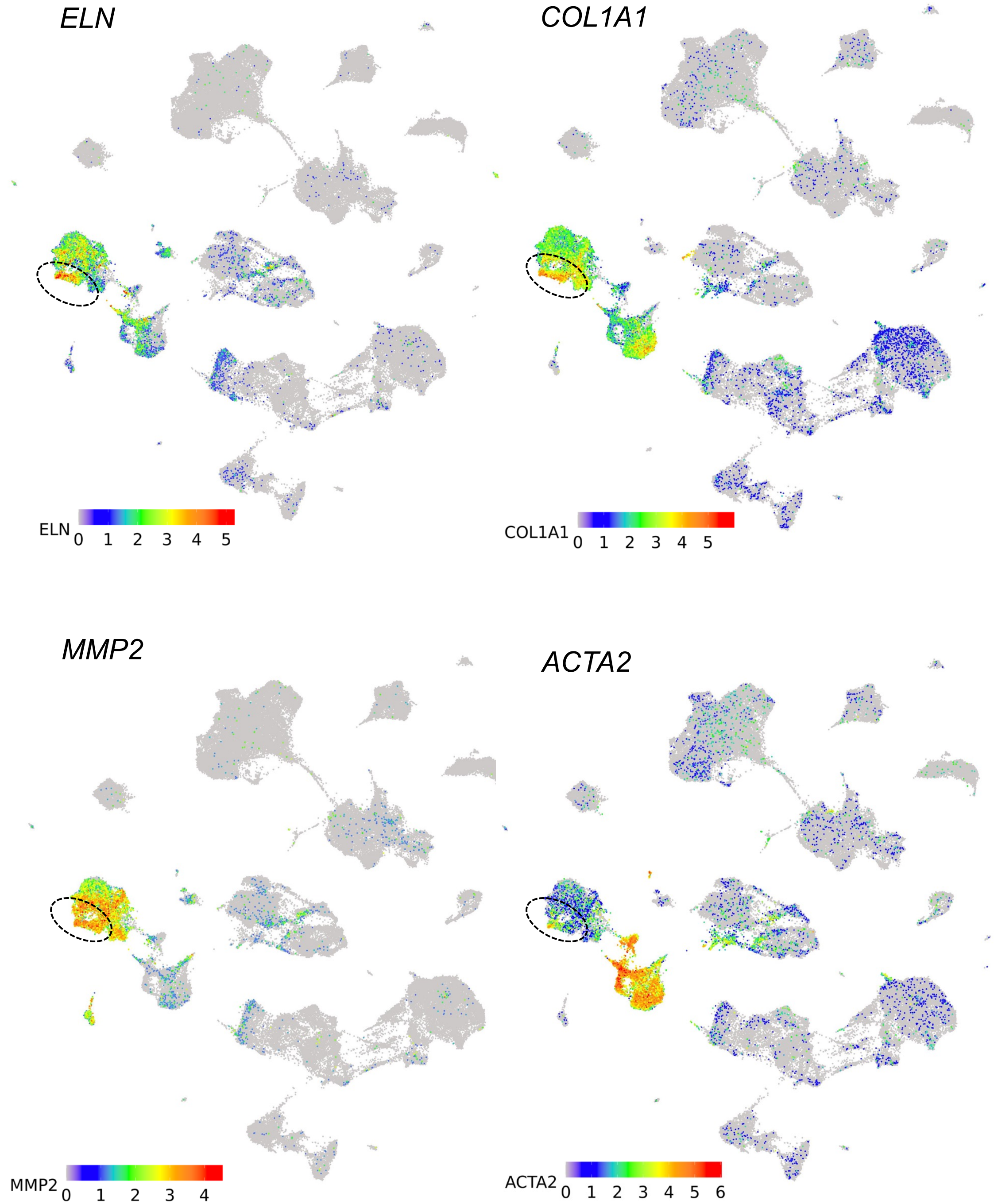

**$L_4$**

Fibroblasts (Fb) subsets

Fb4
contains *GPC3*⁺ *FGFR4*⁺ *VEGFD*⁺ *WNT2*⁺

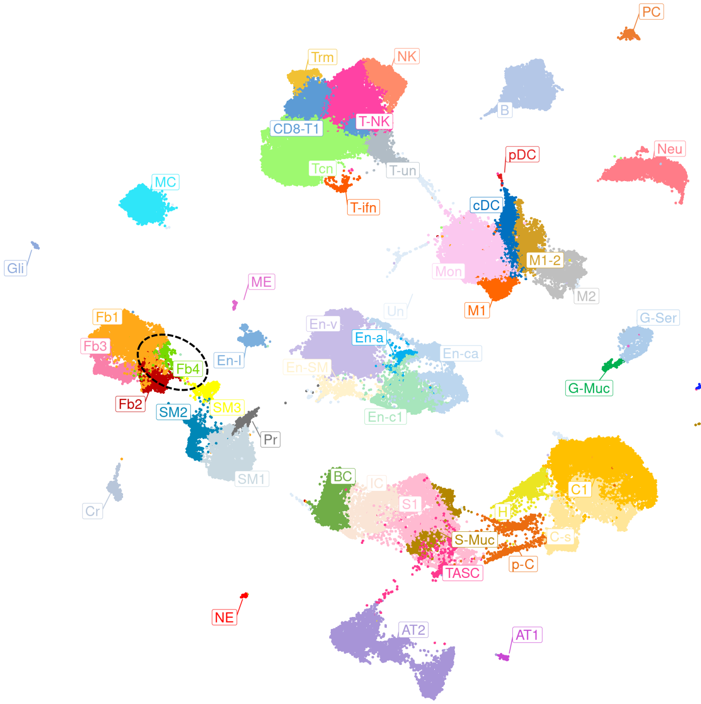


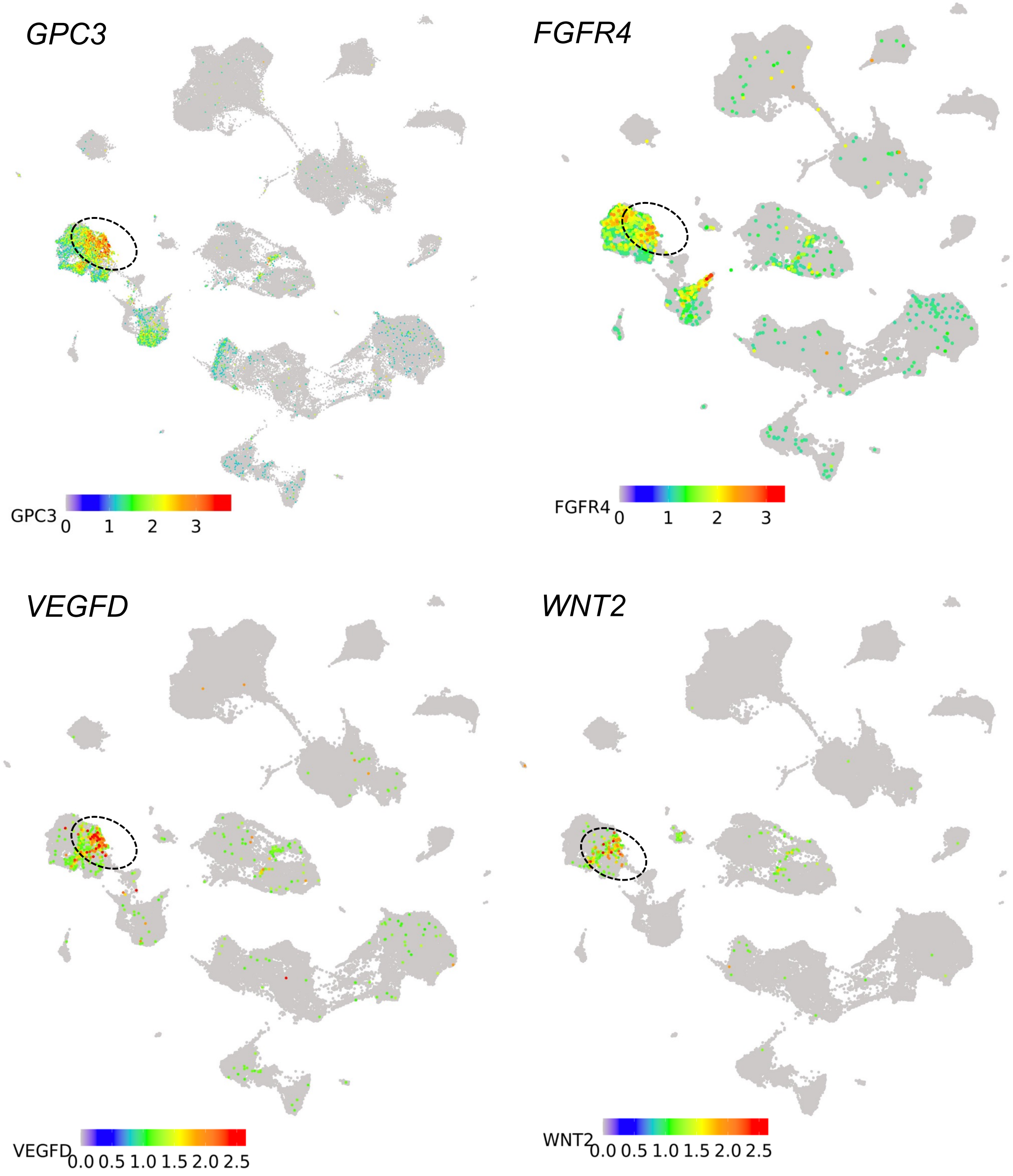

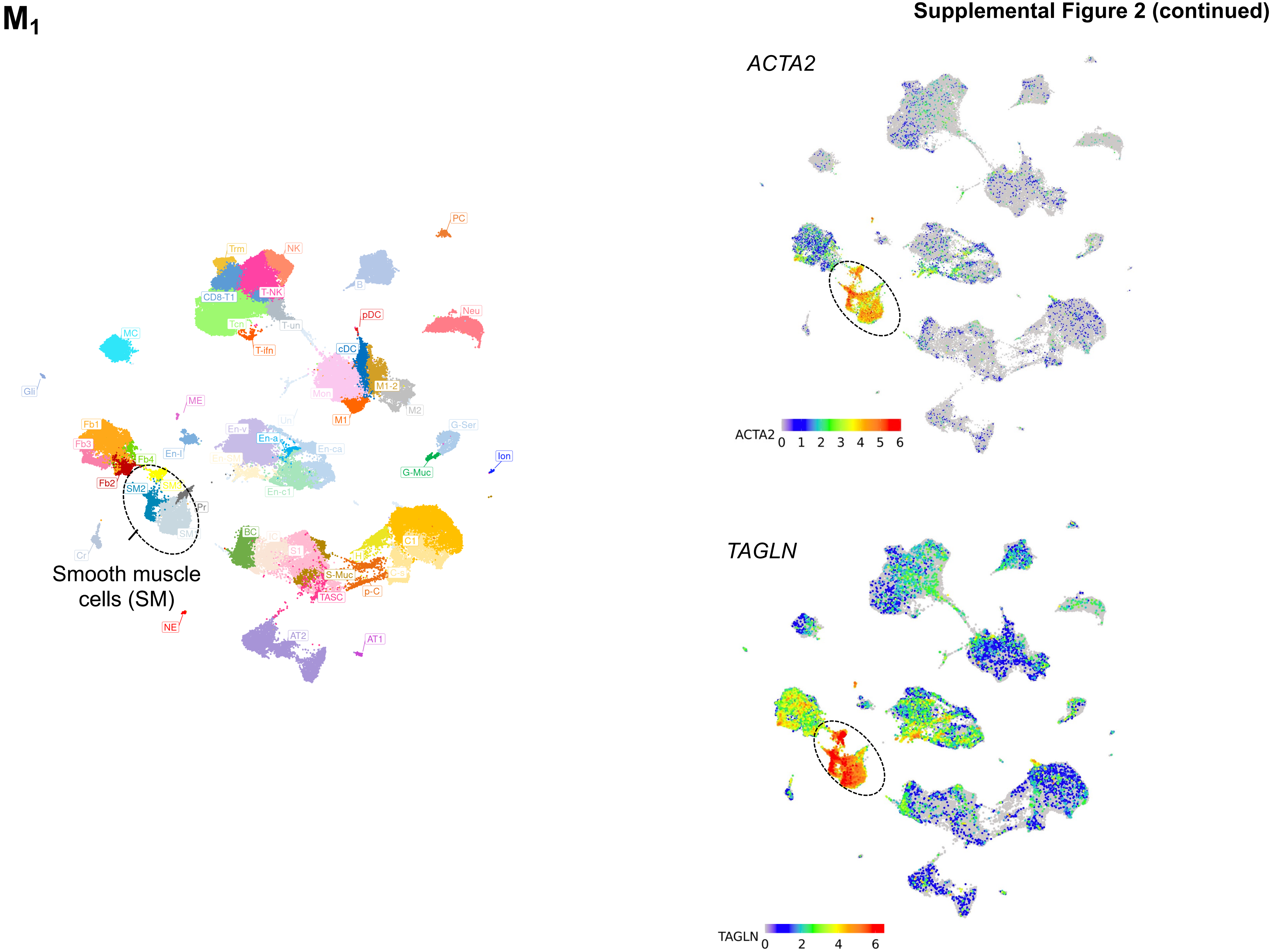
M1
PC
Trm
NK
B
CD8-T1
T-NK
pDC
Neu
Tcn
T-un
MC
T-ifn
cDC
Gli
Mon
M1-2
ME
M2
M1
Un
G-Ser
Fb1
En-v
En-a
Fb3
En-ca
En-l
Fb4
Ion
G-Muc
Fb2
SM2
SM3
En-c1
Pr
BC
IC
C1
SM
S1
H
Cr
C-s
S-Muc
Smooth muscle
cells (SM)
p-C
TASC
NE
AT2
AT1
ACTA2
ACTA2
0 1 2 3 4 5 6
TAGLN
TAGLN
0 2 4 6

Supplemental Figure 2 (continued)

$M_2$

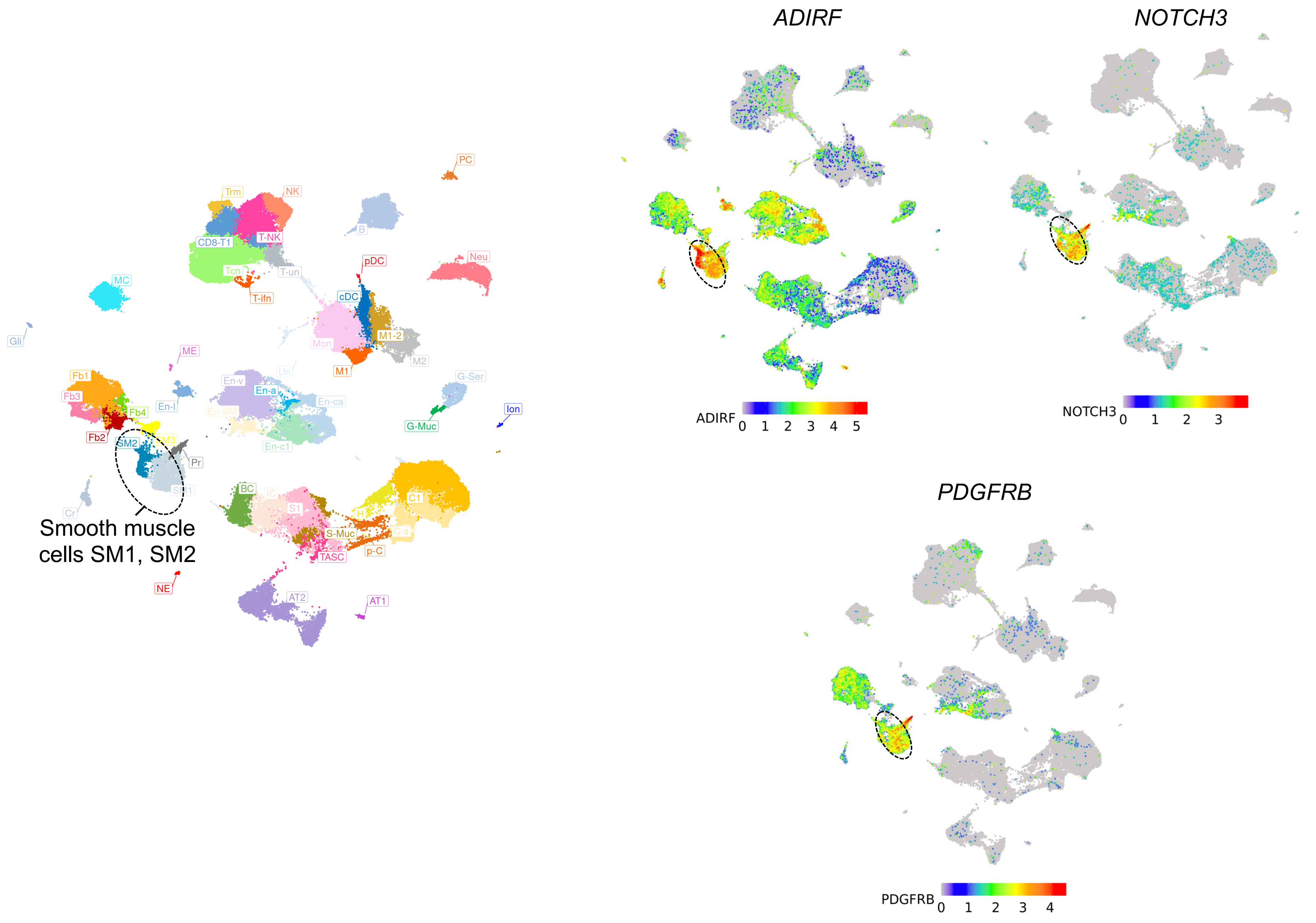

**Supplemental Figure 2 (continued)**

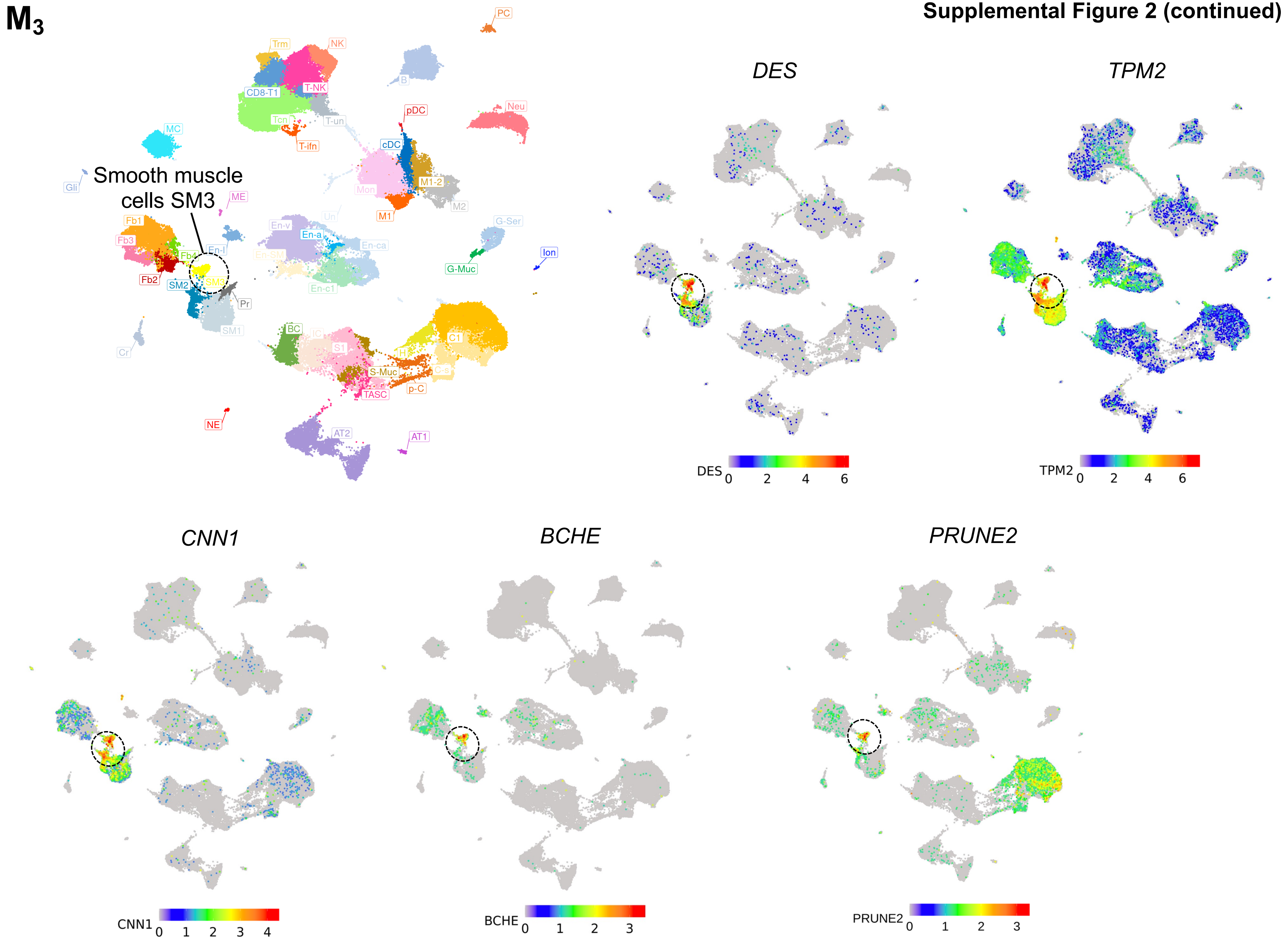

Supplemental Figure 2 (continued)

$M_4$

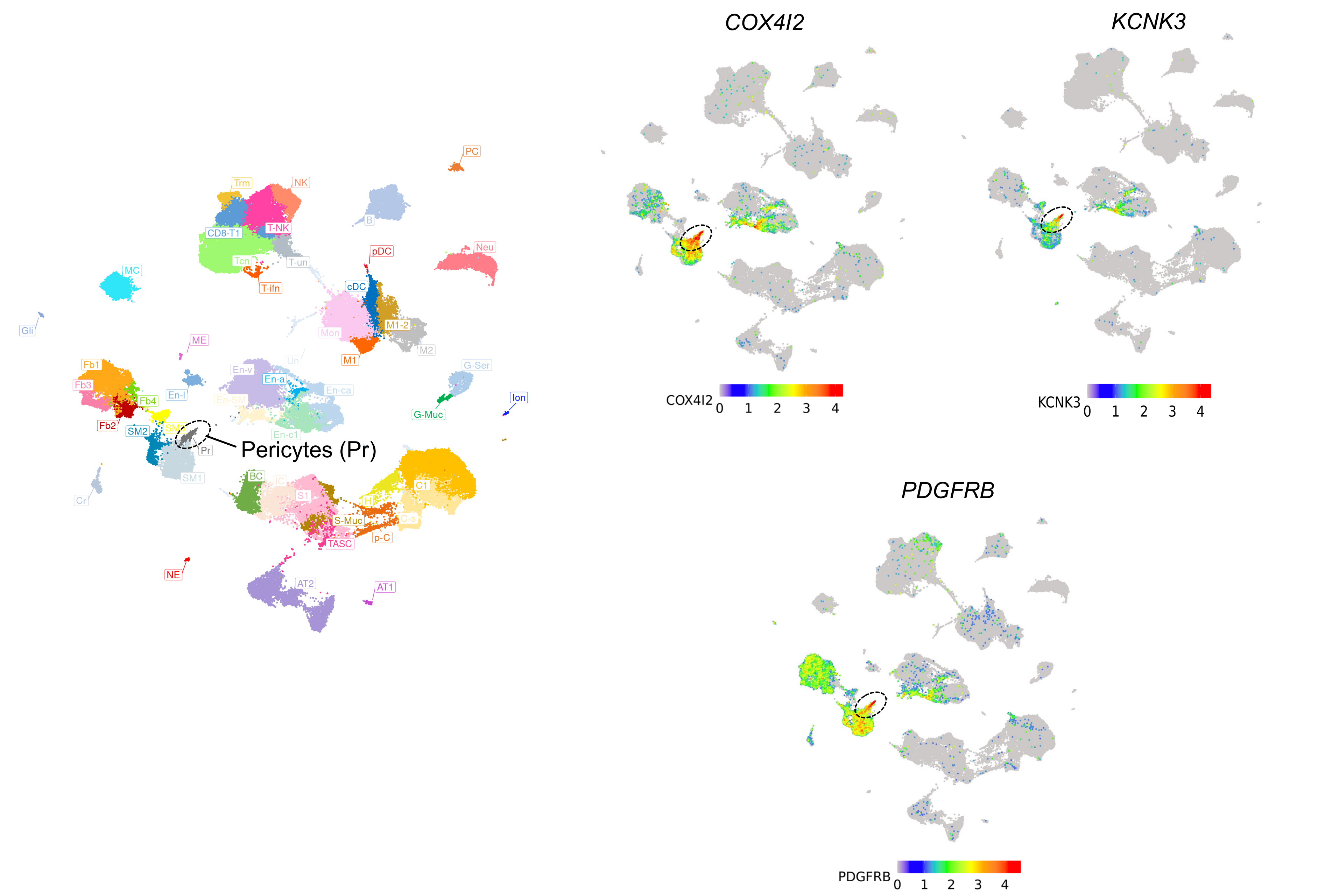

Supplemental Figure 2 (continued)

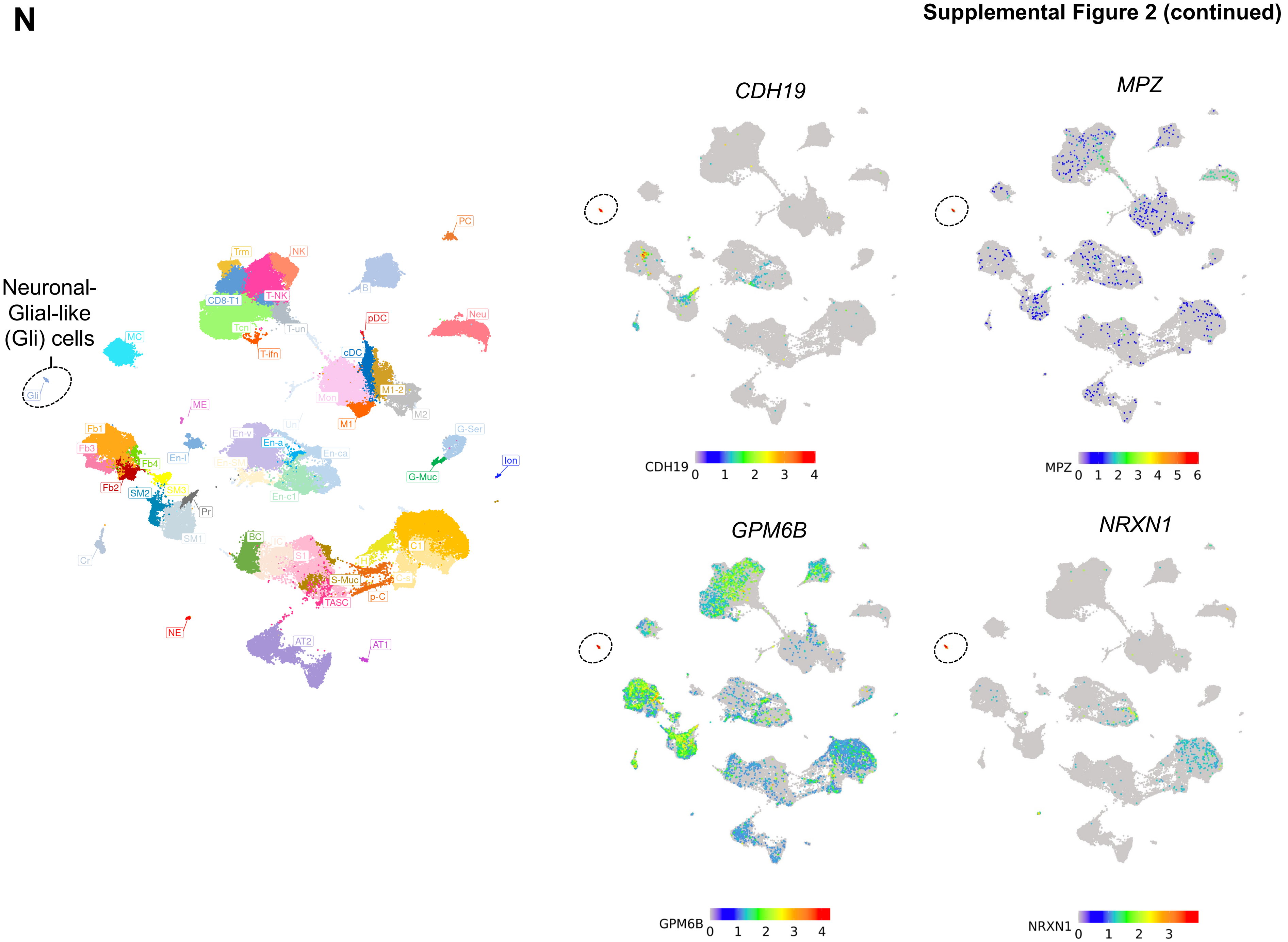

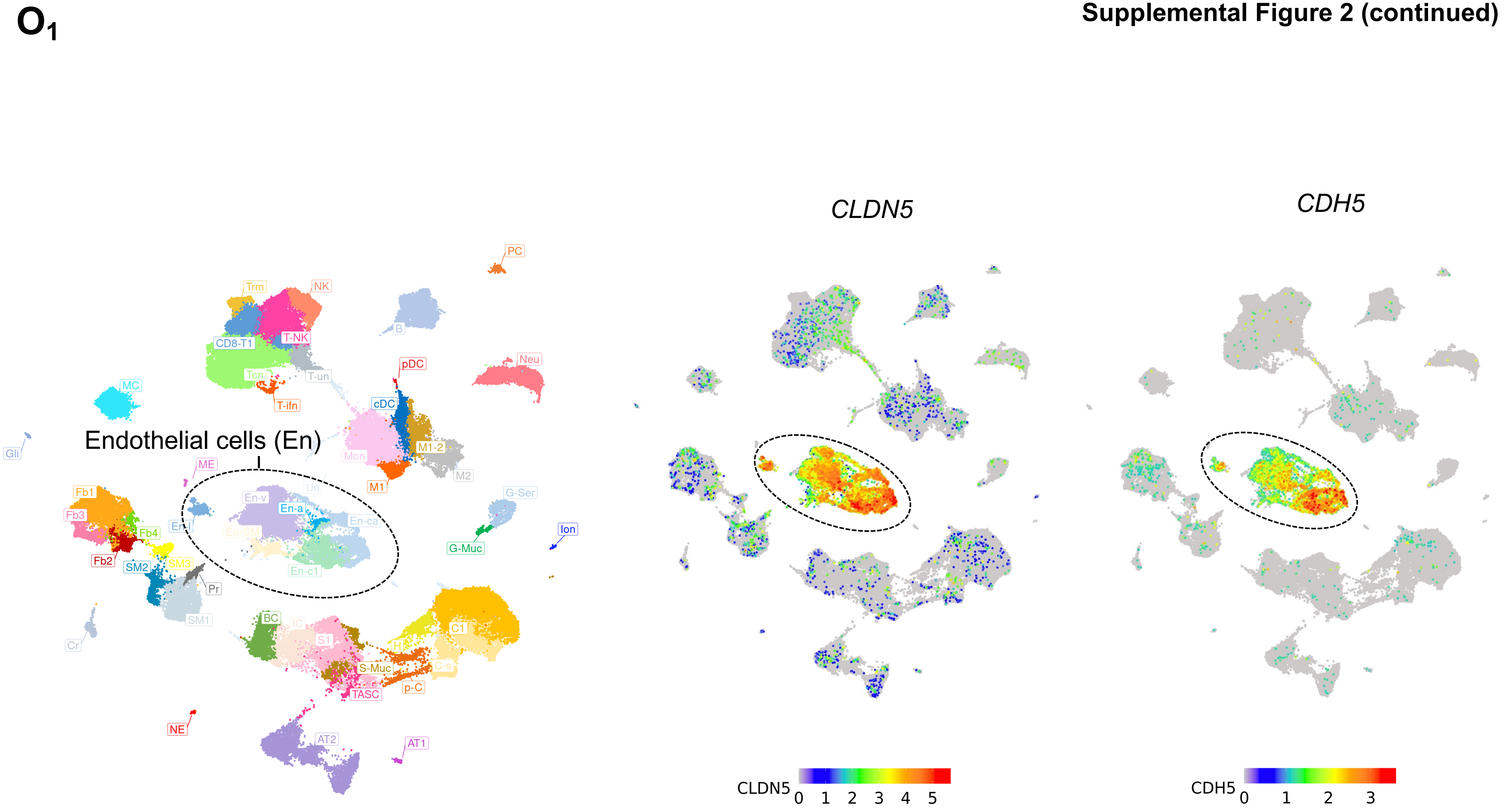
O1
Endothelial cells (En)
PC
Trm
NK
B
CD8-T1
T-NK
T-un
Tcn
pDC
Neu
MC
T-ifn
cDC
Gli
M1-2
Mon
ME
M2
M1
En-v
Un
Fb1
En-a
G-Ser
Fb3
En-l
En-ca
Ion
Fb4
En-SM
G-Muc
Fb2
SM2
SM3
En-c1
Pr
SM1
BC
IC
C1
S1
Cr
H
S-Muc
C-s
p-C
TASC
NE
AT2
AT1
CLDN5
CDH5
CLDN5
0 1 2 3 4 5
CDH5
0 1 2 3

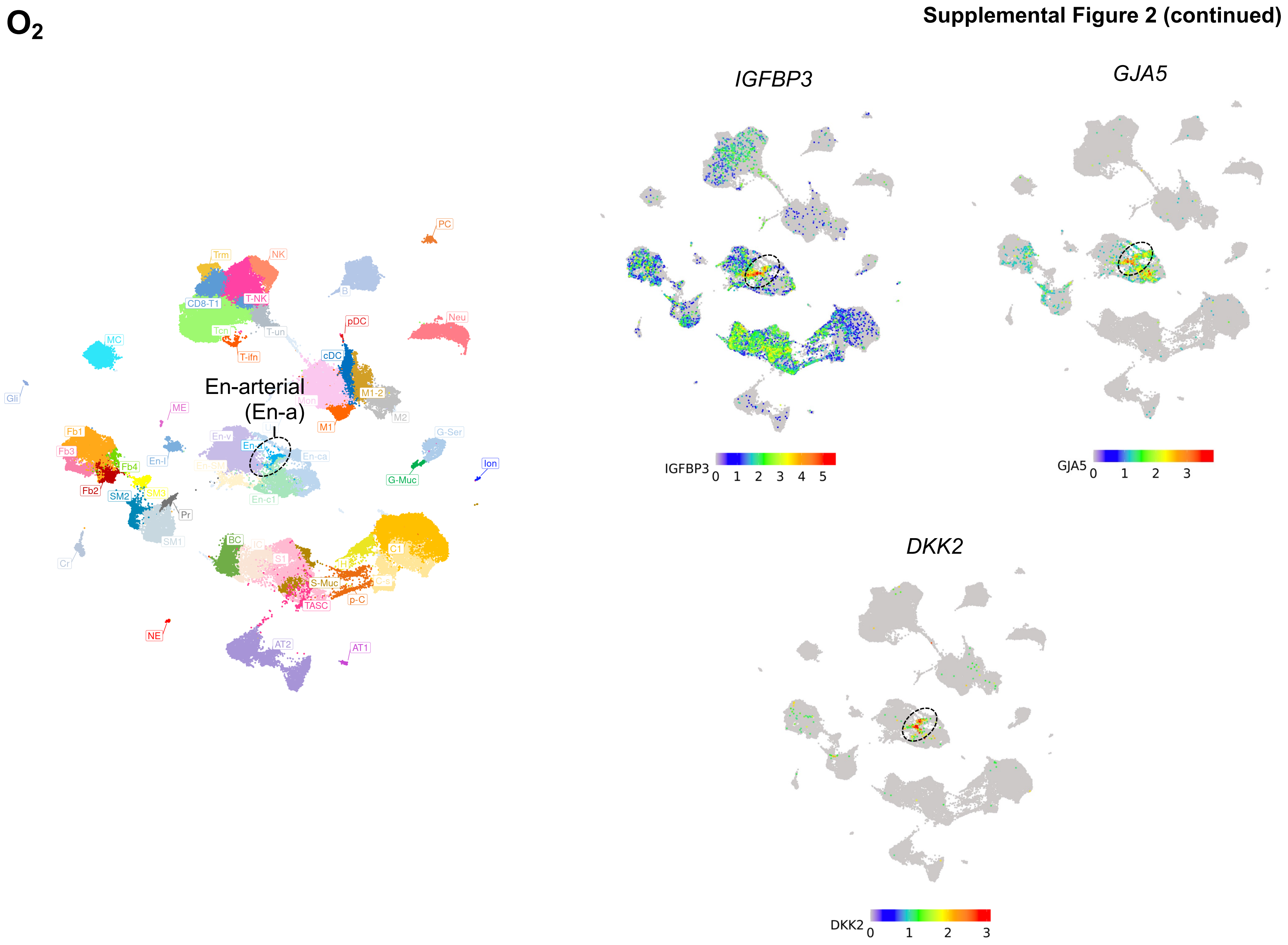
O2
En-arterial
(En-a)
PC
Trm
NK
B
CD8-T1
T-NK
Neu
pDC
Tcn
T-un
MC
T-ifn
cDC
Gli
M1-2
Mon
M2
ME
M1
Fb1
En-v
Fb3
En-l
En-ca
G-Ser
Fb4
Ion
Fb2
SM2
SM3
En-c1
G-Muc
Pr
SM1
BC
C1
Cr
S1
S-Muc
C-s
TASC
p-C
NE
AT2
AT1
IGFBP3
GJA5
DKK2
IGFBP3 0 1 2 3 4 5
GJA5 0 1 2 3
DKK2 0 1 2 3

# O$_3$



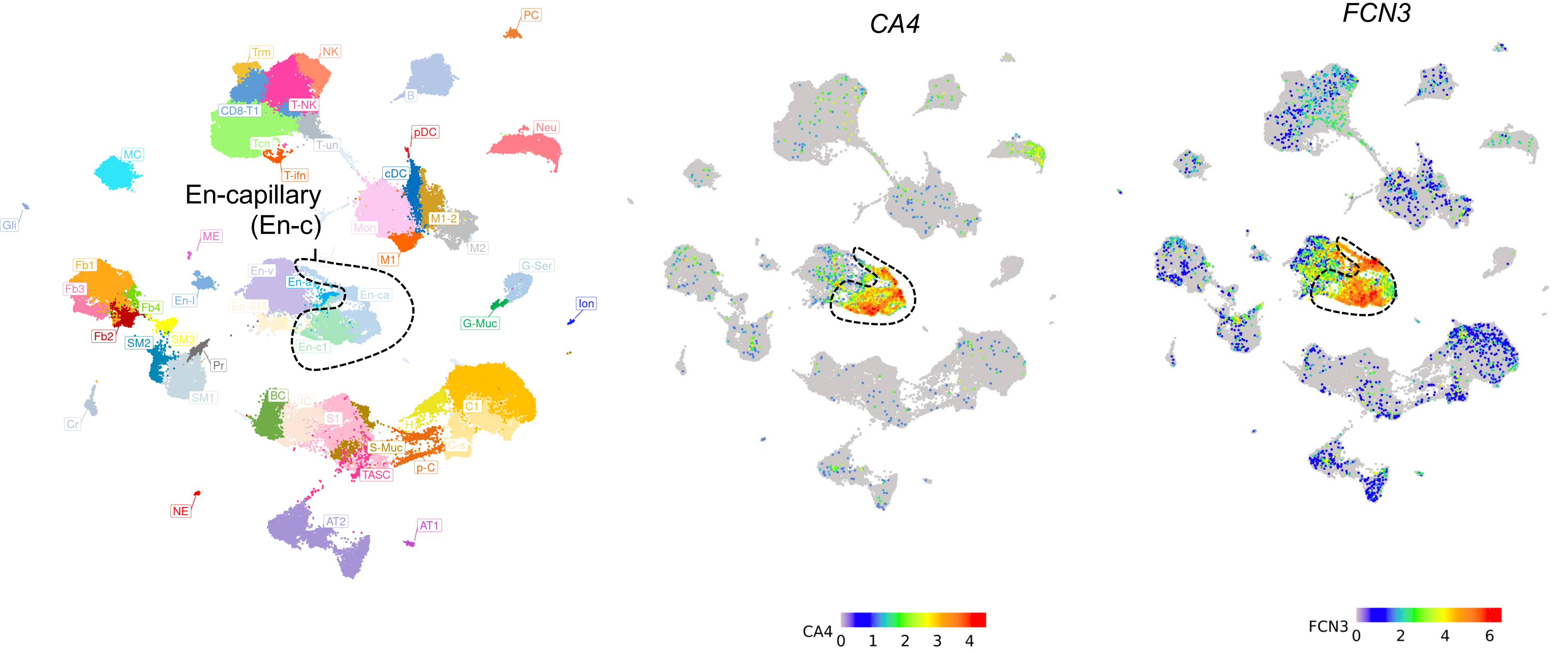
PC
Trm
NK
B
CD8-T1
T-NK
Tcn
T-un
pDC
Neu
MC
T-ifn
cDC
En-capillary
(En-c)
Gli
Mon
M1-2
ME
M2
M1
Fb1
En-v
G-Ser
En-a
En-ca
Fb3
Fb4
En-l
En-SM
Ion
Fb2
SM2
SM3
G-Muc
En-c1
Pr
BC
C1
SM1
IC
S1
Cr
H
S-Muc
C-s
p-C
TASC
NE
AT2
AT1
CA4
FCN3
CA4 0 1 2 3 4
FCN3 0 2 4 6

O4

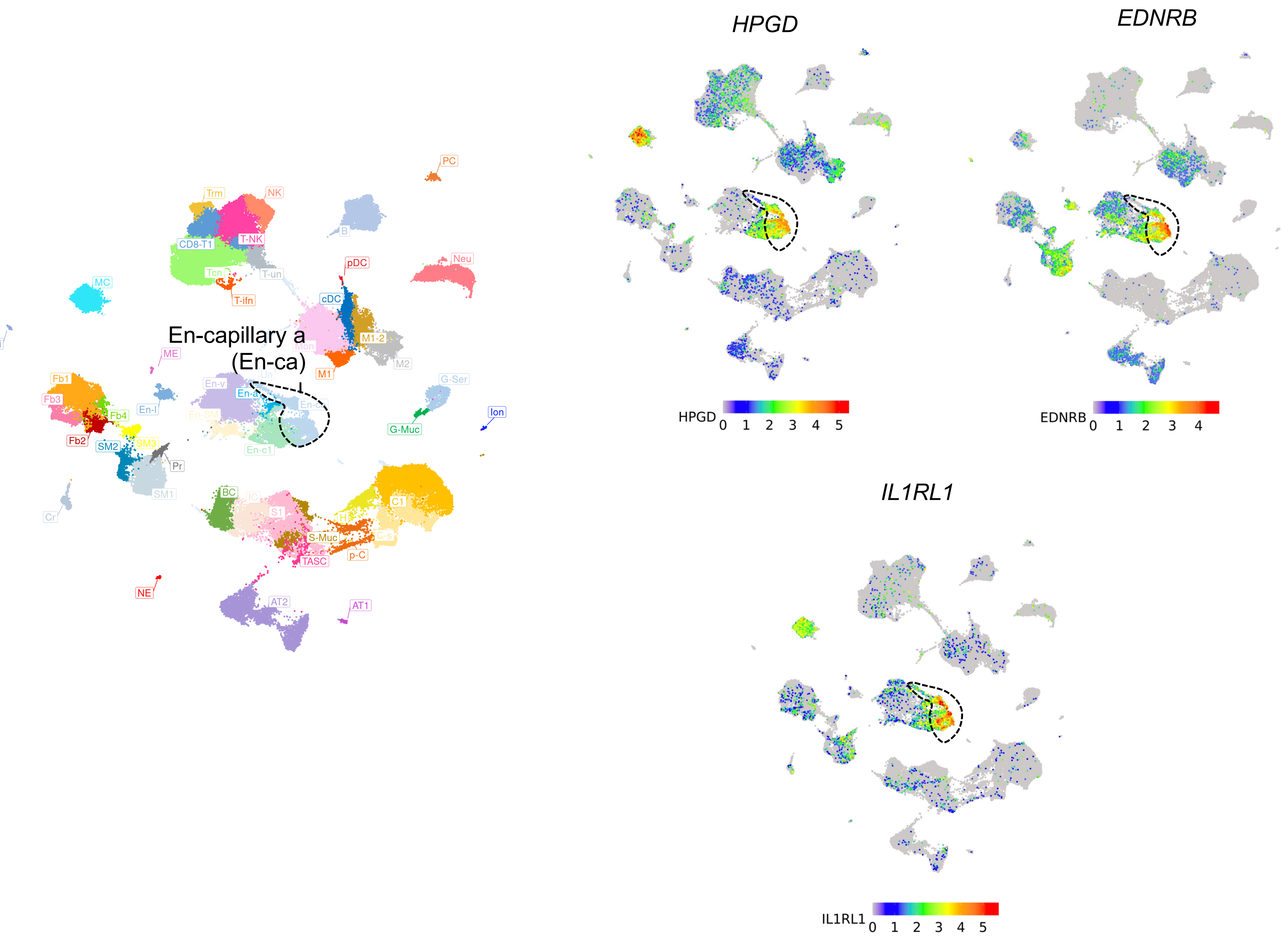
En-capillary a
(En-ca)
PC
Trm
NK
CD8-T1
T-NK
B
Tcn
T-un
T-ifn
pDC
Neu
MC
cDC
Gli
ME
Mon
M1-2
M2
M1
Un
En-v
En-a
En-ca
En-SM
En-c1
En-l
Fb1
Fb3
Fb4
Fb2
SM2
SM3
Pr
SM1
G-Ser
G-Muc
Ion
Cr
BC
IC
S1
C1
H
C-s
S-Muc
TASC
p-C
NE
AT2
AT1
HPGD
EDNRB
IL1RL1
HPGD 0 1 2 3 4 5
EDNRB 0 1 2 3 4
IL1RL1 0 1 2 3 4 5

$O_5$

Supplemental Figure 2 (continued)

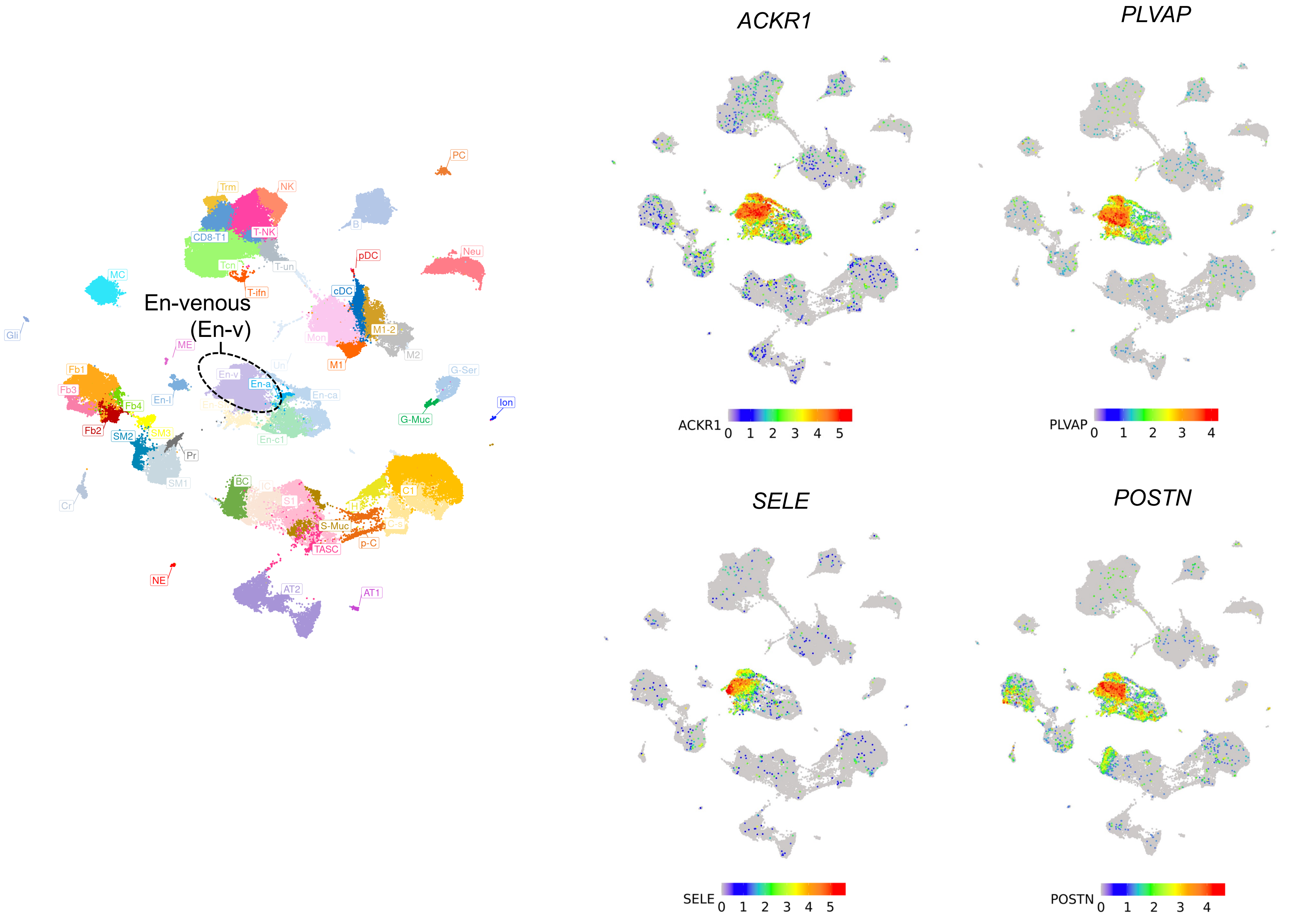

O6

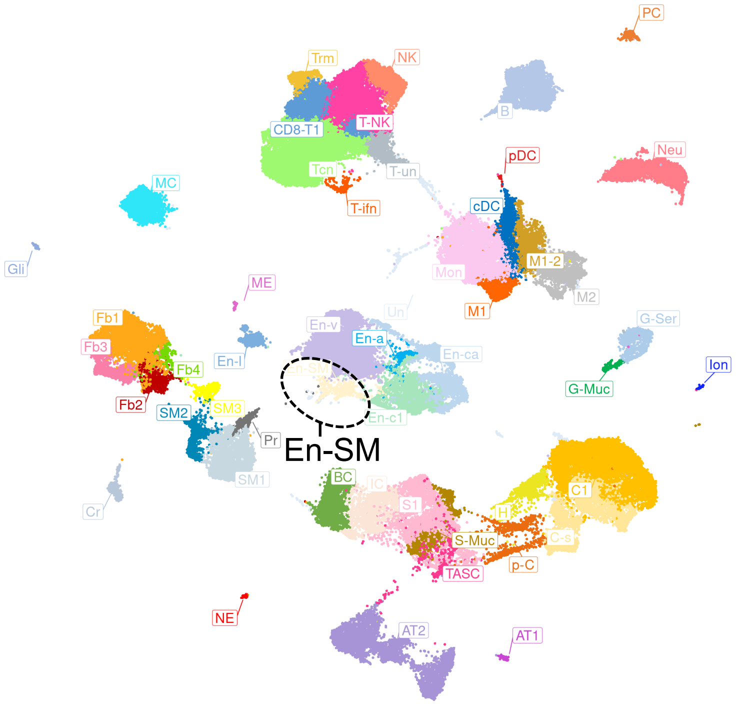
PC
Trm
NK
B
CD8-T1
T-NK
pDC
Neu
T-un
Tcn
T-ifn
MC
cDC
Gli
Mon
M1-2
ME
M1
M2
Un
En-v
En-a
G-Ser
Fb1
Fb3
En-l
En-ca
Fb4
Ion
Fb2
G-Muc
SM2
SM3
En-c1
Pr
En-SM
SM1
BC
C1
S1
Cr
H
S-Muc
C-s
TASC
p-C
NE
AT2
AT1

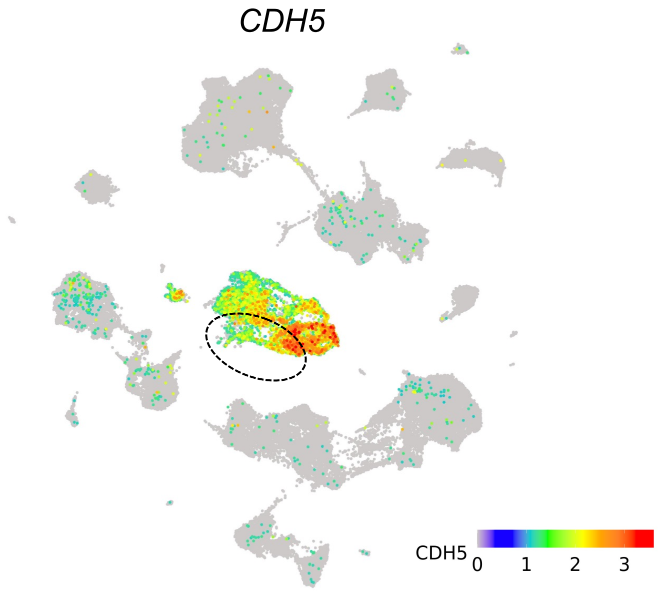
CDH5
CDH5
0 1 2 3

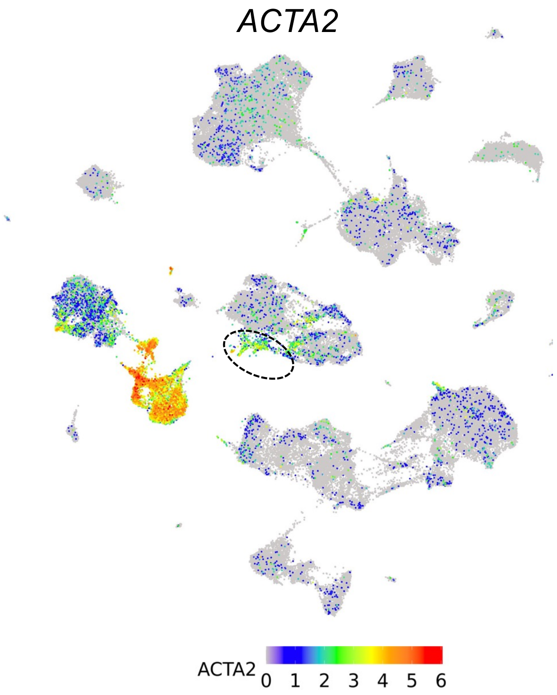
ACTA2
ACTA2
0 1 2 3 4 5 6

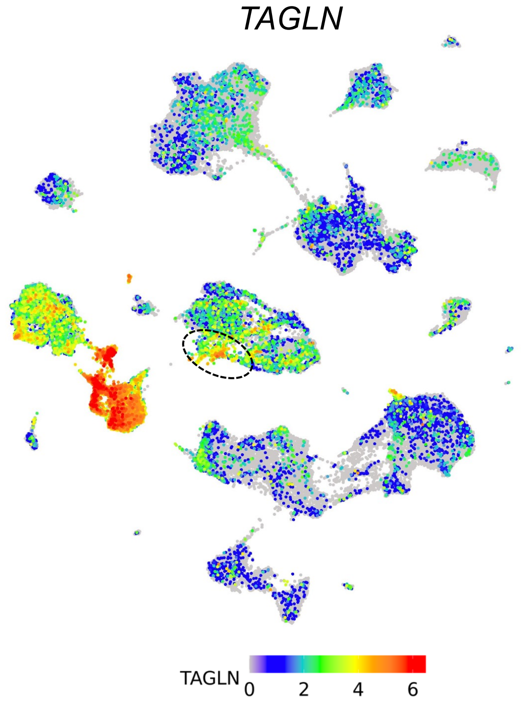
TAGLN
TAGLN
0 2 4 6

**Supplemental Figure 2 (continued)**

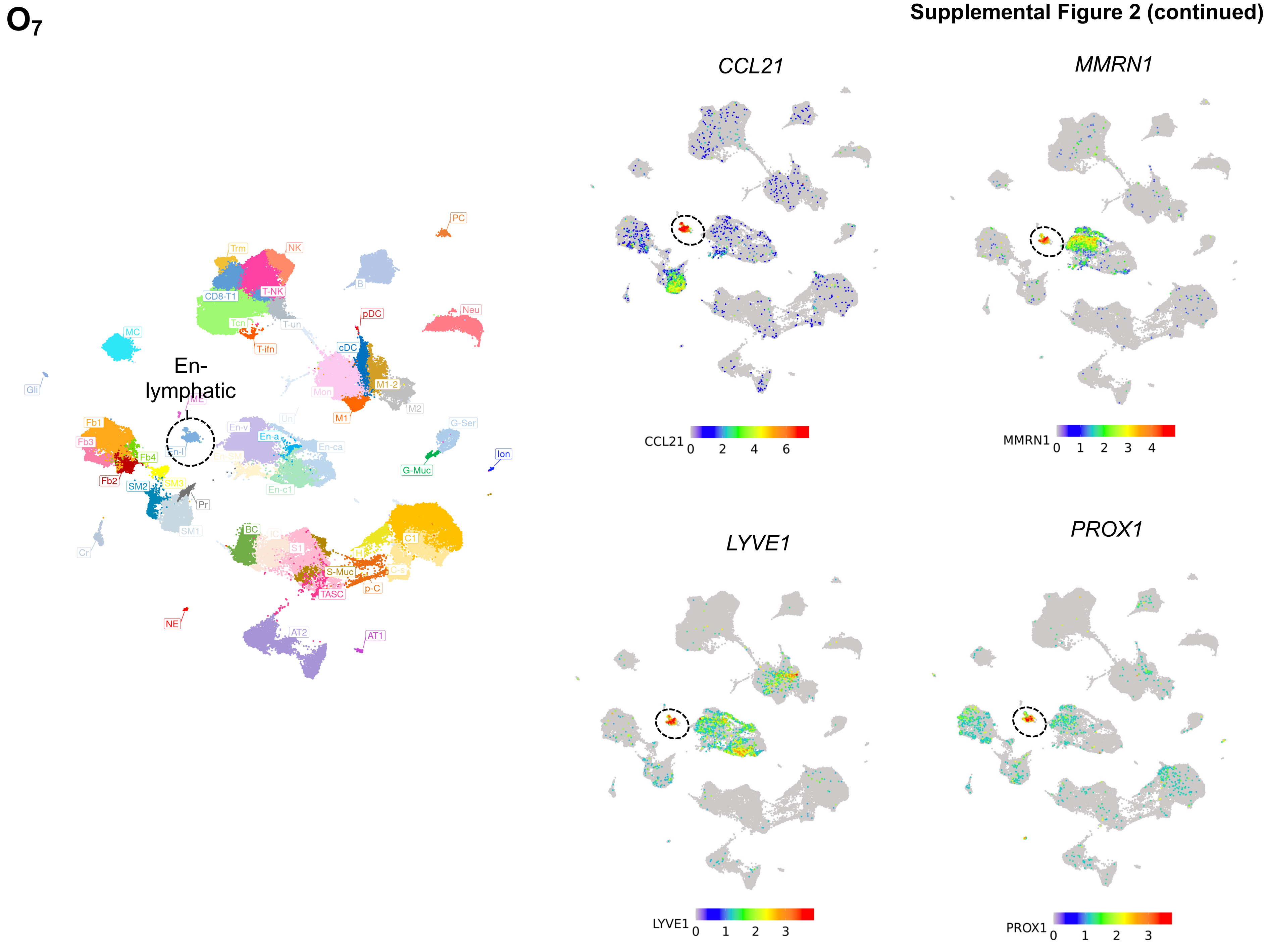

**P**

Supplemental Figure 2 (continued)

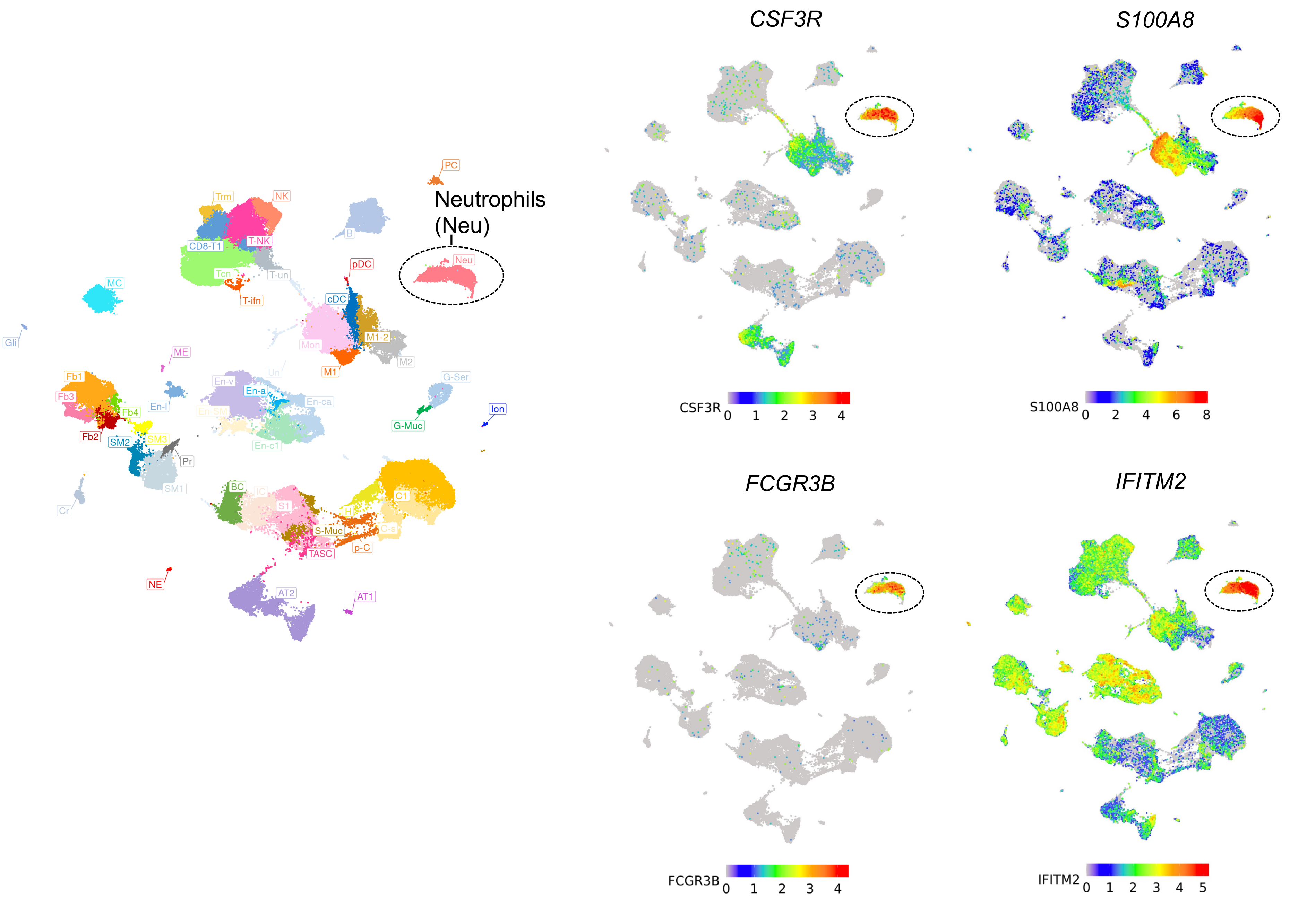

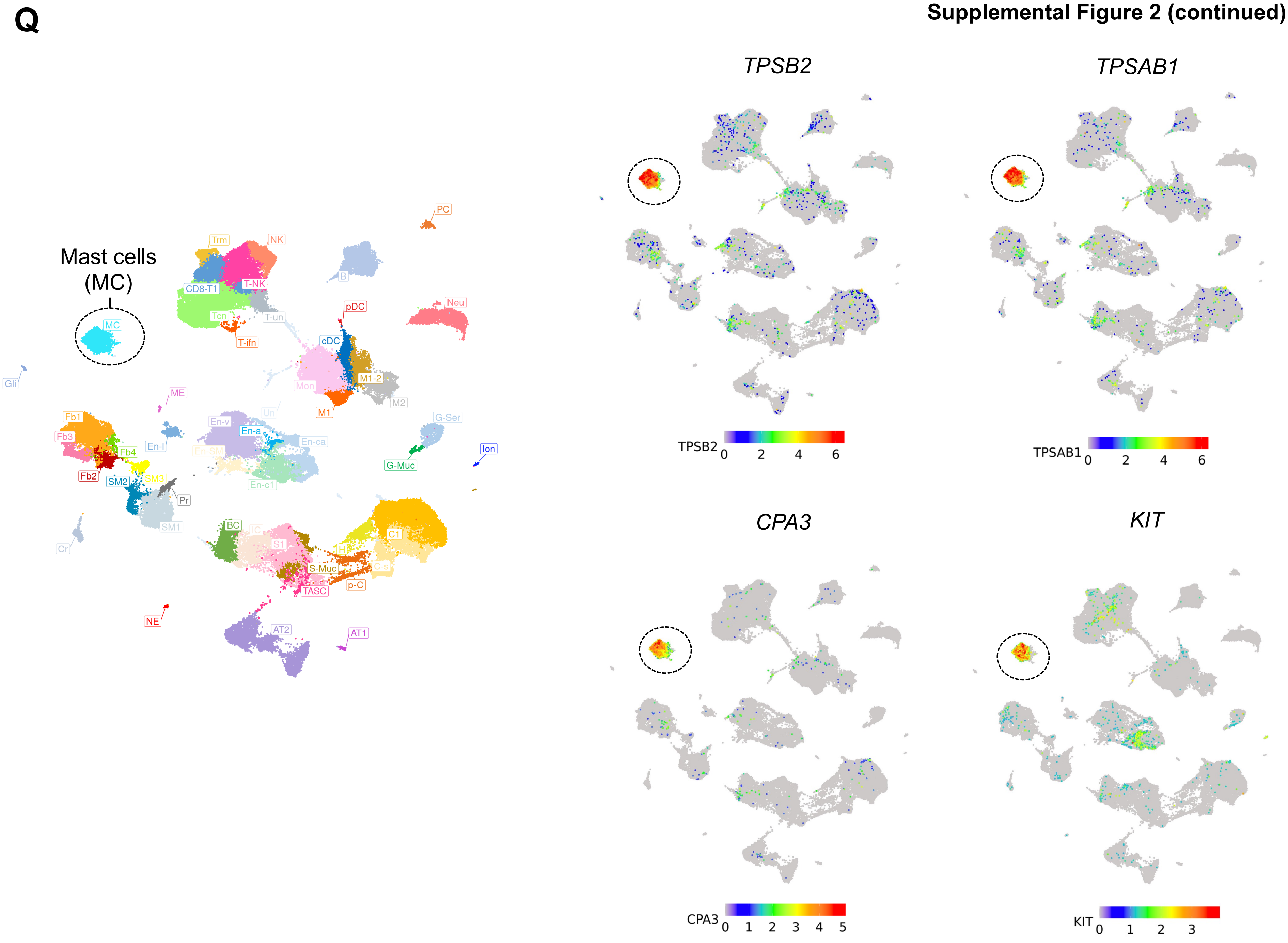
Q
Mast cells
(MC)
MC
Trm
NK
CD8-T1
T-NK
Tcn
T-un
T-ifn
PC
B
pDC
Neu
cDC
Mon
M1-2
M2
M1
Gli
ME
Fb1
Fb3
Fb4
Fb2
En-l
En-v
En-a
En-ca
En-SM
En-c1
SM2
SM3
Pr
SM1
Cr
G-Ser
G-Muc
Ion
BC
IC
S1
C1
H
S-Muc
C-s
p-C
TASC
NE
AT2
AT1
TPSB2
TPSAB1
CPA3
KIT
0 2 4 6
0 1 2 3 4 5
0 1 2 3

**Supplemental Figure 2 (continued)**

**R**

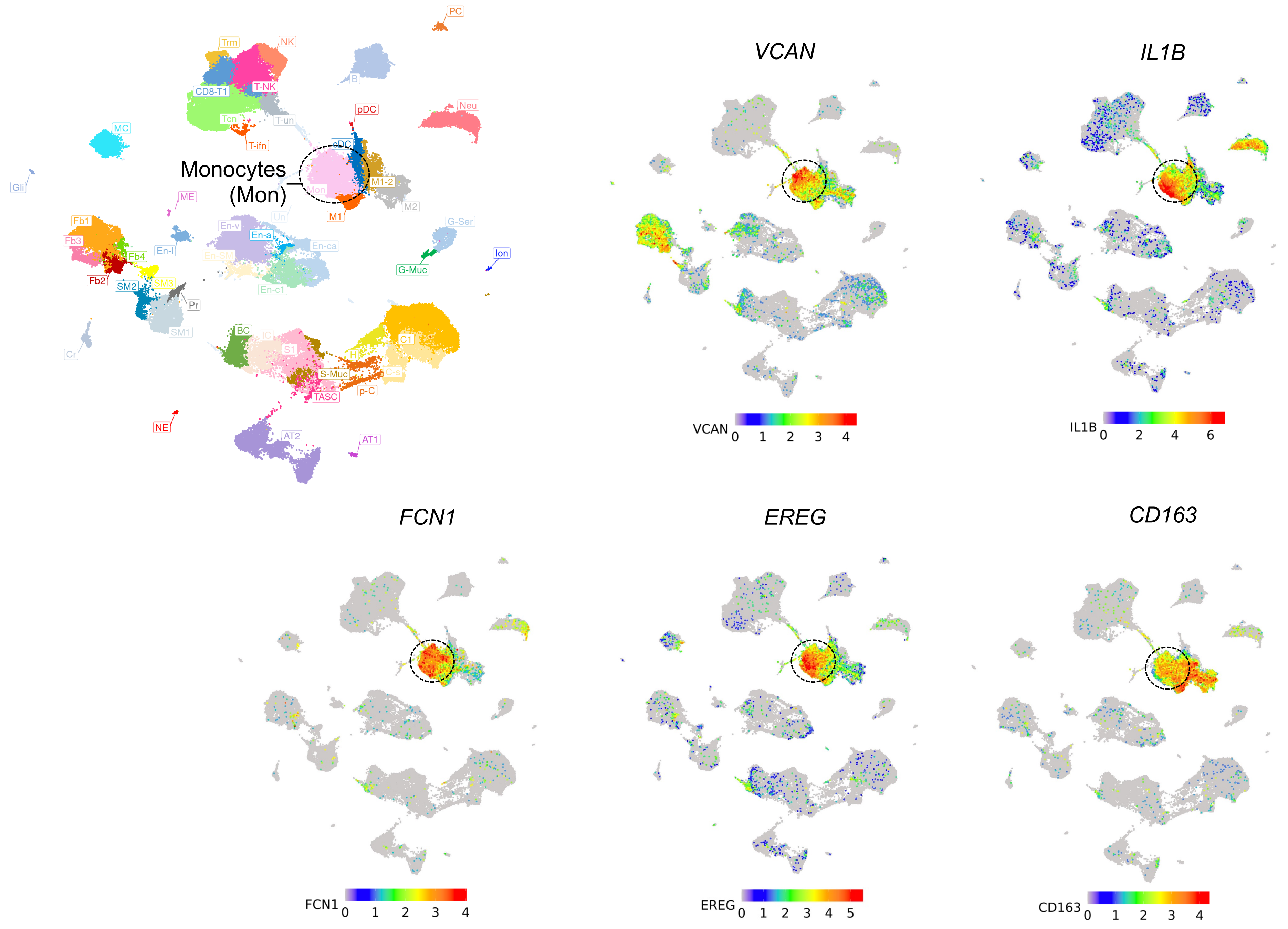

**S1**

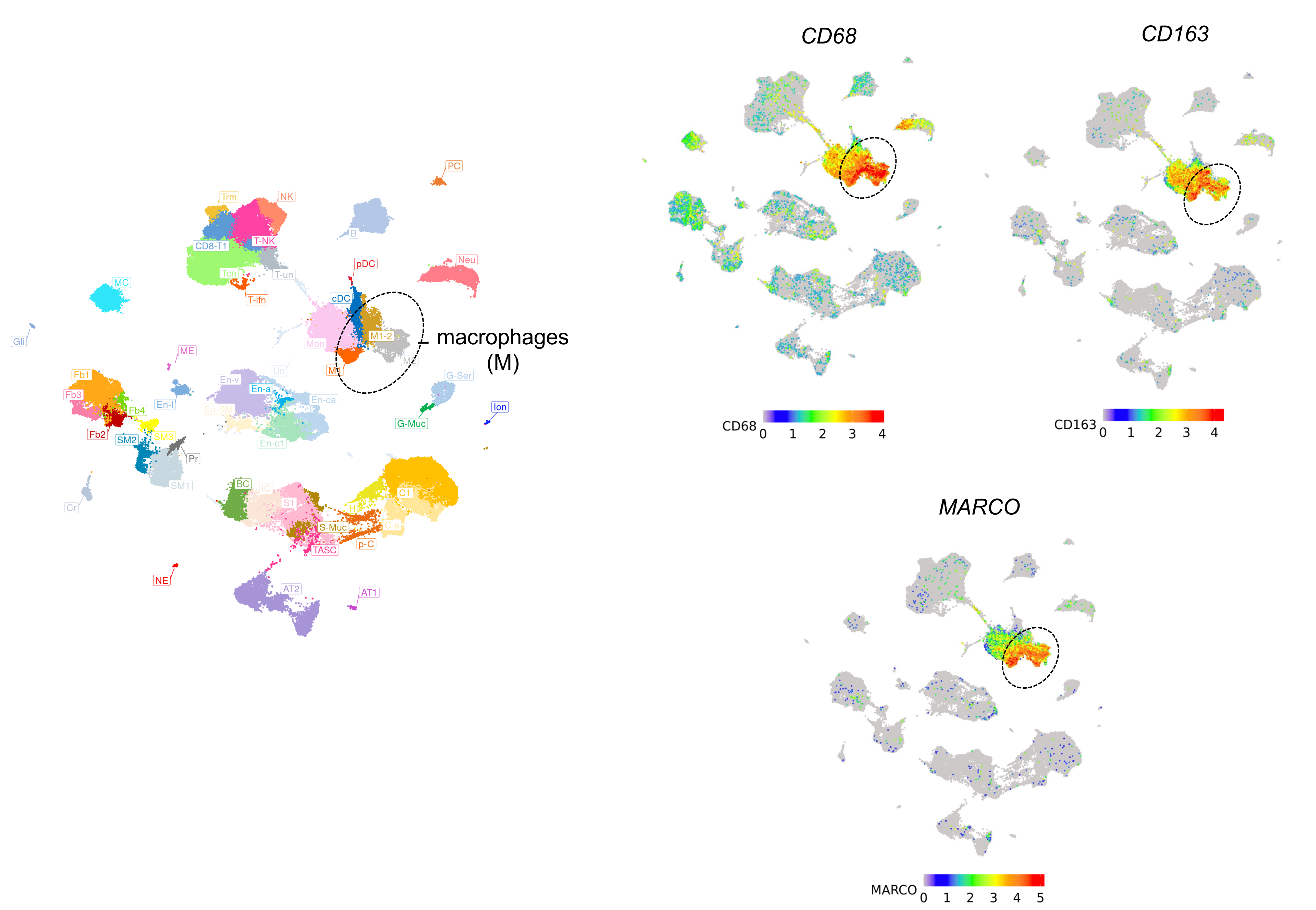
PC
Trm
NK
B
T-NK
CD8-T1
pDC
Neu
Tcn
T-un
MC
T-ifn
cDC
M1-2
macrophages
(M)
Gli
ME
Mon
M1
Un
G-Ser
Fb1
En-v
En-a
Fb3
En-ca
En-l
Ion
Fb4
En-SM
G-Muc
Fb2
SM2
SM3
En-c1
Pr
SM1
BC
IC
C1
Cr
S1
H
S-Muc
C-s
p-C
TASC
NE
AT2
AT1
CD68
CD68 0 1 2 3 4
CD163
CD163 0 1 2 3 4
MARCO
MARCO 0 1 2 3 4 5

S2



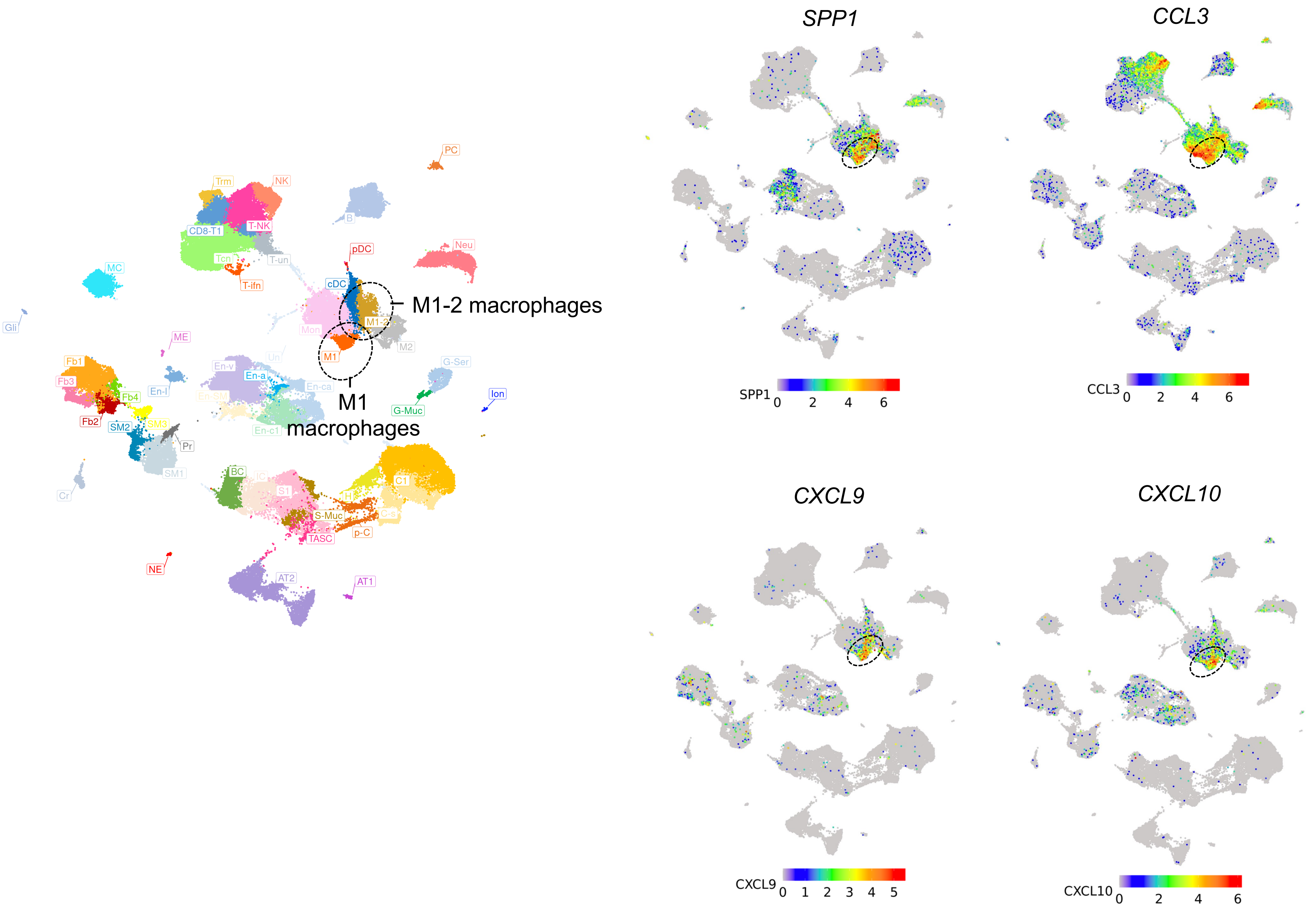

M1-2 macrophages
M1 macrophages
PC
Trm
NK
T-NK
CD8-T1
B
Neu
pDC
Tcn
T-un
T-ifn
MC
cDC
M1-2
Mon
M1
M2
Gli
ME
Un
Fb1
Fb3
Fb4
Fb2
En-v
En-a
En-ca
En-l
En-SM
En-c1
SM2
SM3
SM1
Pr
G-Ser
G-Muc
Ion
Cr
BC
IC
S1
H
C1
C-s
S-Muc
p-C
TASC
NE
AT2
AT1
SPP1
CCL3
CXCL9
CXCL10
SPP1 0 2 4 6
CCL3 0 2 4 6
CXCL9 0 1 2 3 4 5
CXCL10 0 2 4 6

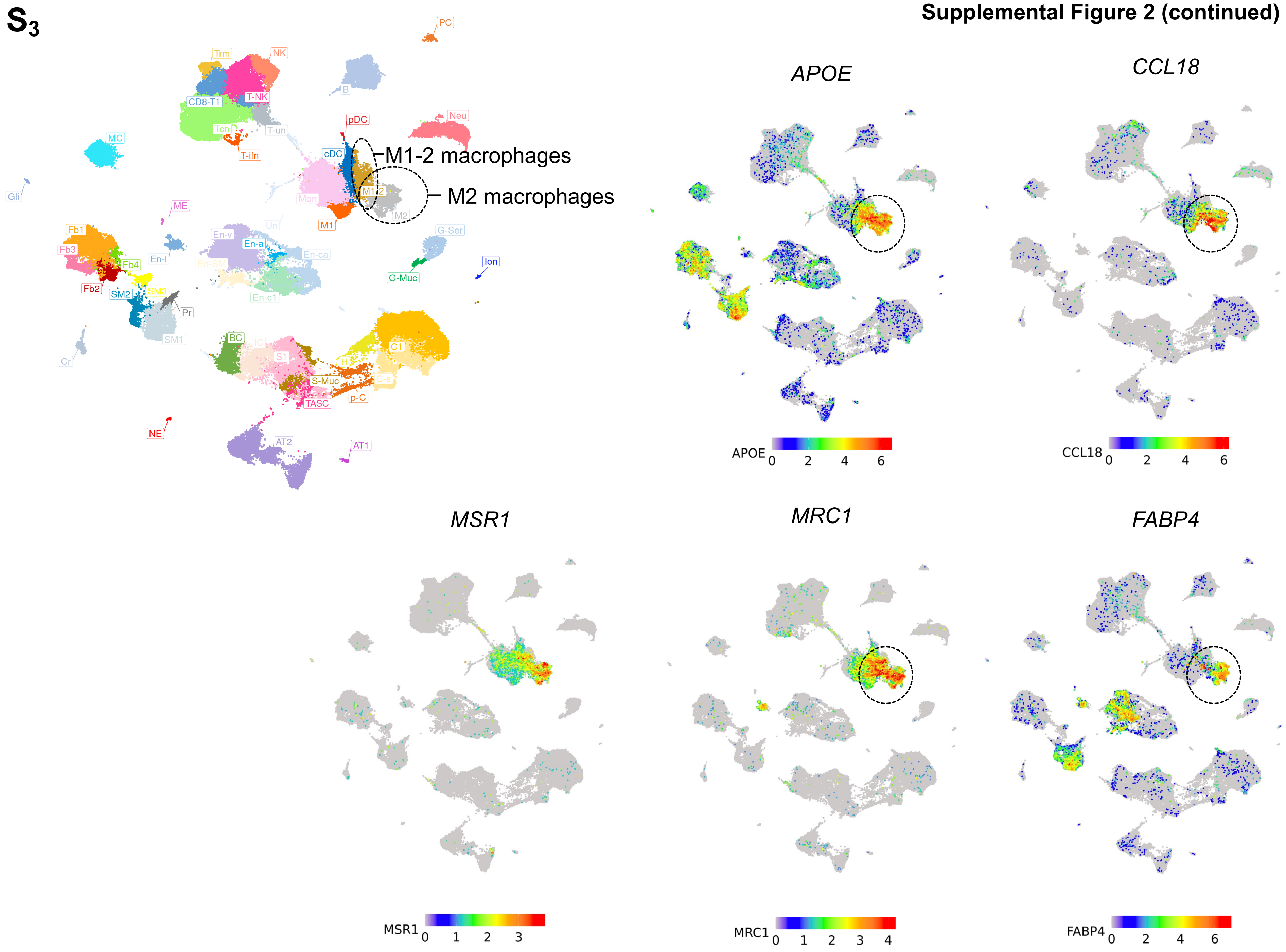
S3
PC
Trm
NK
B
CD8-T1
T-NK
Tcn
T-un
T-ifn
pDC
Neu
MC
cDC
M1-2 macrophages
M1-2
M2 macrophages
Mon
M2
M1
Gli
ME
Un
Fb1
En-v
En-a
En-ca
G-Ser
Fb3
En-l
Fb4
En-SM
Ion
G-Muc
Fb2
SM2
SM3
En-c1
Pr
SM1
BC
IC
C1
Cr
S1
H
S-Muc
C-s
TASC
p-C
NE
AT2
AT1
APOE
CCL18
MSR1
MRC1
FABP4

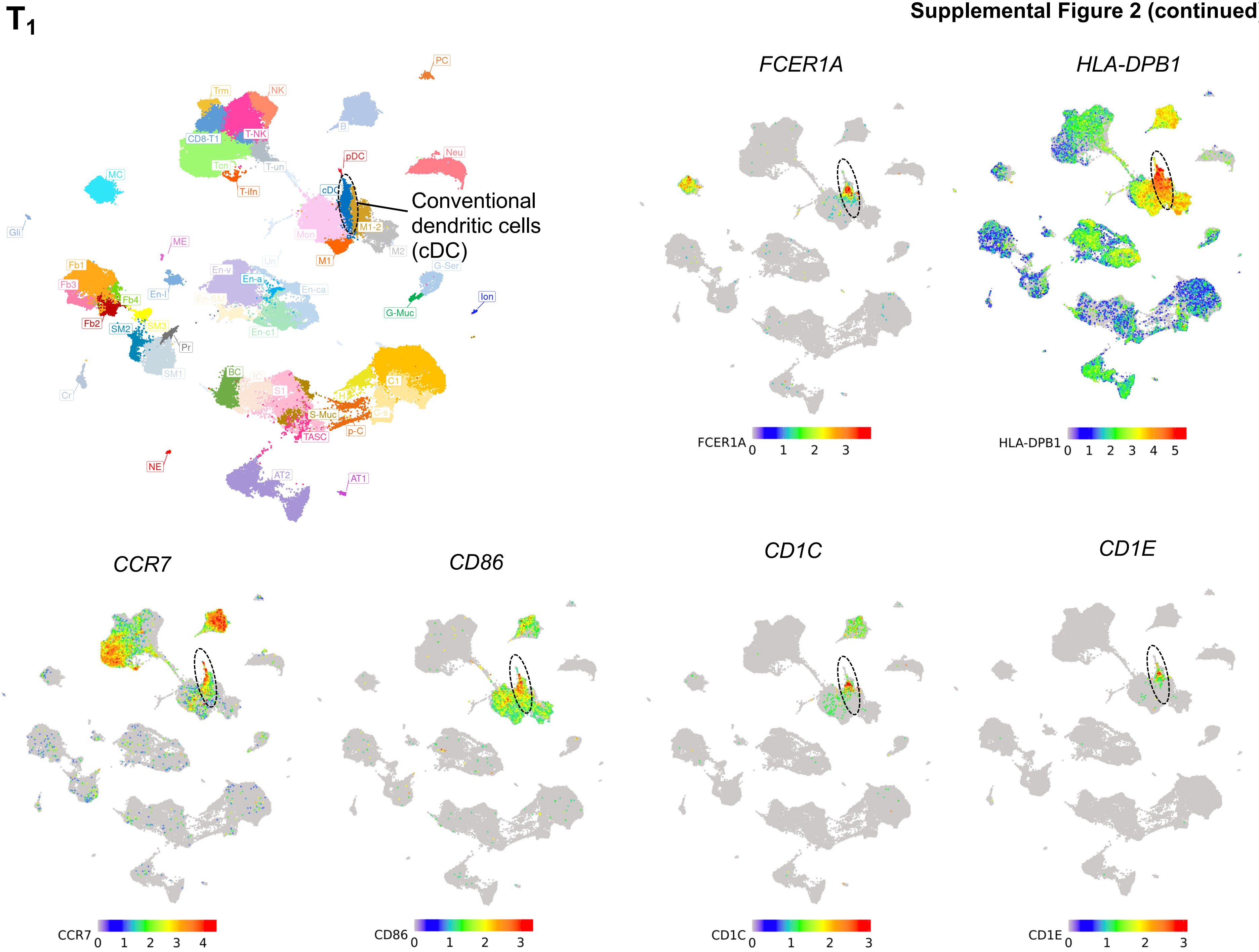

$T_1$
PC
Trm
NK
B
CD8-T1
T-NK
Neu
pDC
Tcn
T-un
MC
T-ifn
cDC
Conventional dendritic cells (cDC)
Mon
M1-2
Gli
M2
ME
M1
G-Ser
Fb1
En-v
Un
Fb3
En-a
En-ca
En-l
Fb4
Ion
G-Muc
Fb2
SM2
SM3
En-c1
Pr
SM1
BC
S1
C1
Cr
H
S-Muc
C-s
TASC
p-C
NE
AT2
AT1
FCER1A
HLA-DPB1
FCER1A 0 1 2 3
HLA-DPB1 0 1 2 3 4 5
CCR7
CD86
CD1C
CD1E
CCR7 0 1 2 3 4
CD86 0 1 2 3
CD1C 0 1 2 3
CD1E 0 1 2 3

**Supplemental Figure 2 (continued)**

**$T_2$**

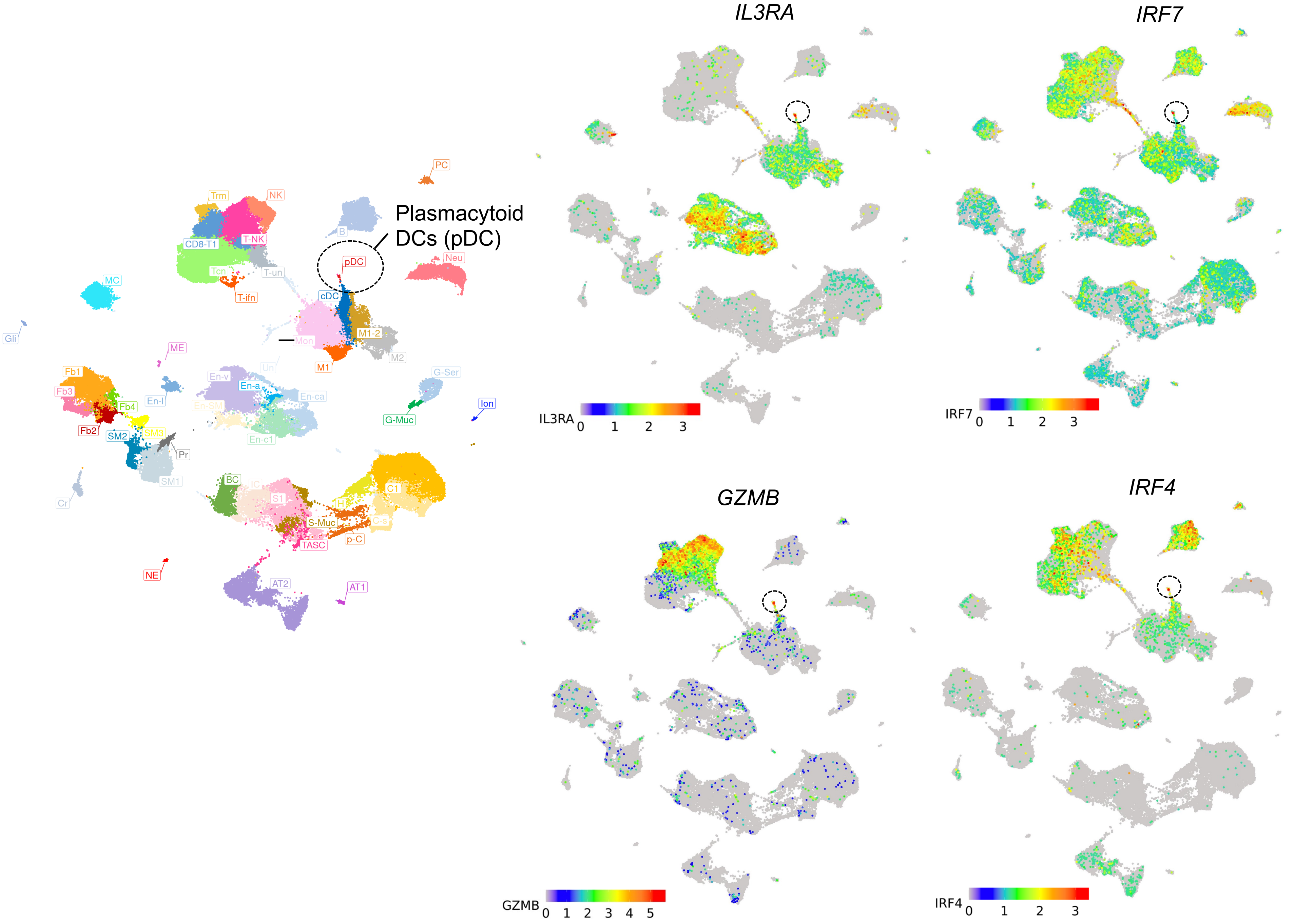

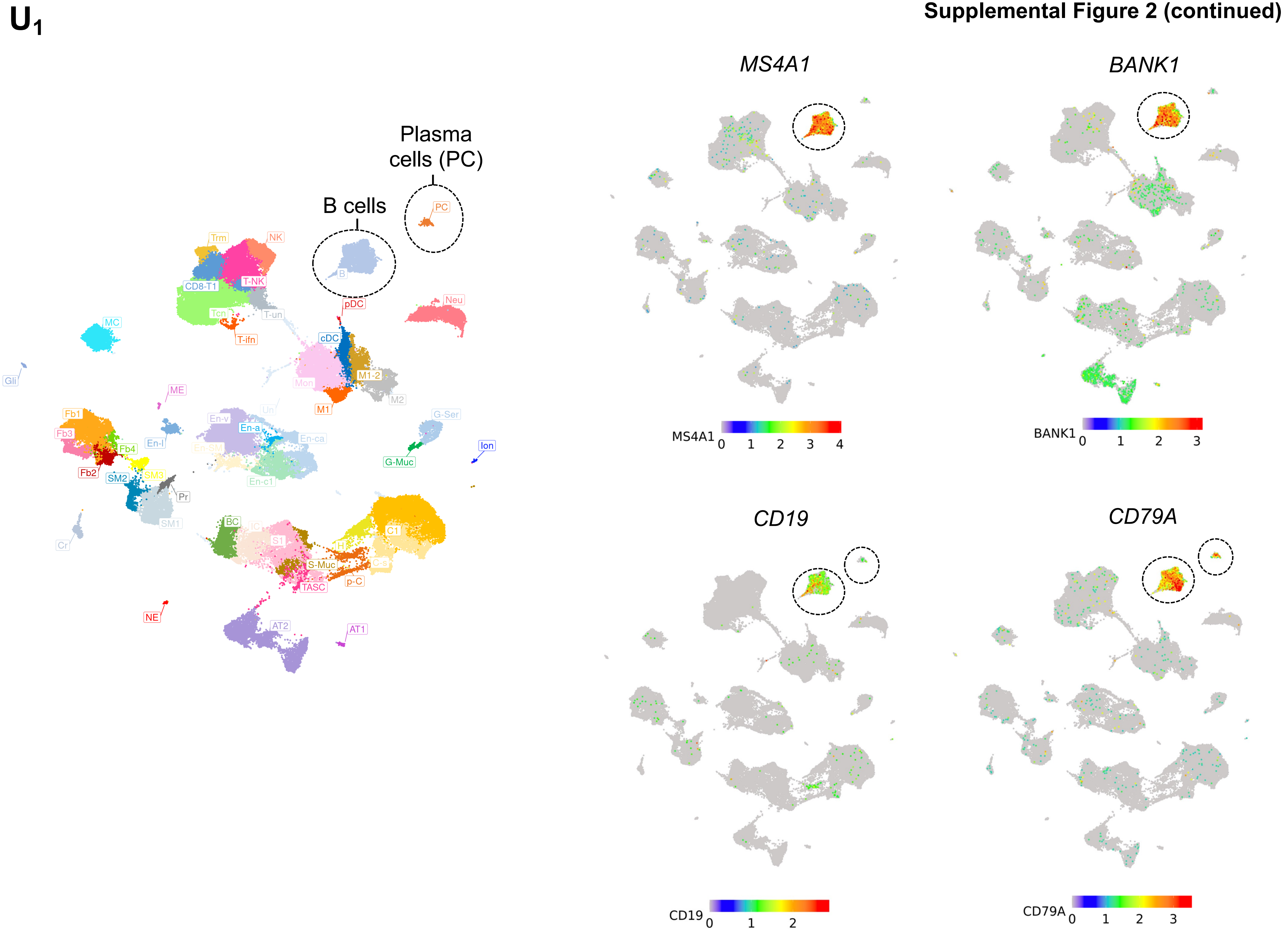
U1
Plasma cells (PC)
B cells
PC
B
Trm
NK
CD8-T1
T-NK
Tcn
T-un
T-ifn
Neu
pDC
MC
cDC
Gli
Mon
M1-2
M2
ME
M1
Fb1
Fb3
En-v
En-a
Un
G-Ser
En-ca
En-l
Fb4
En-SM
Ion
G-Muc
Fb2
SM2
SM3
En-c1
Pr
SM1
BC
IC
S1
C1
Cr
H
C-s
S-Muc
p-C
TASC
NE
AT2
AT1
MS4A1
BANK1
CD19
CD79A
0 1 2 3 4
0 1 2 3
0 1 2
0 1 2 3

Supplemental Figure 2 (continued)

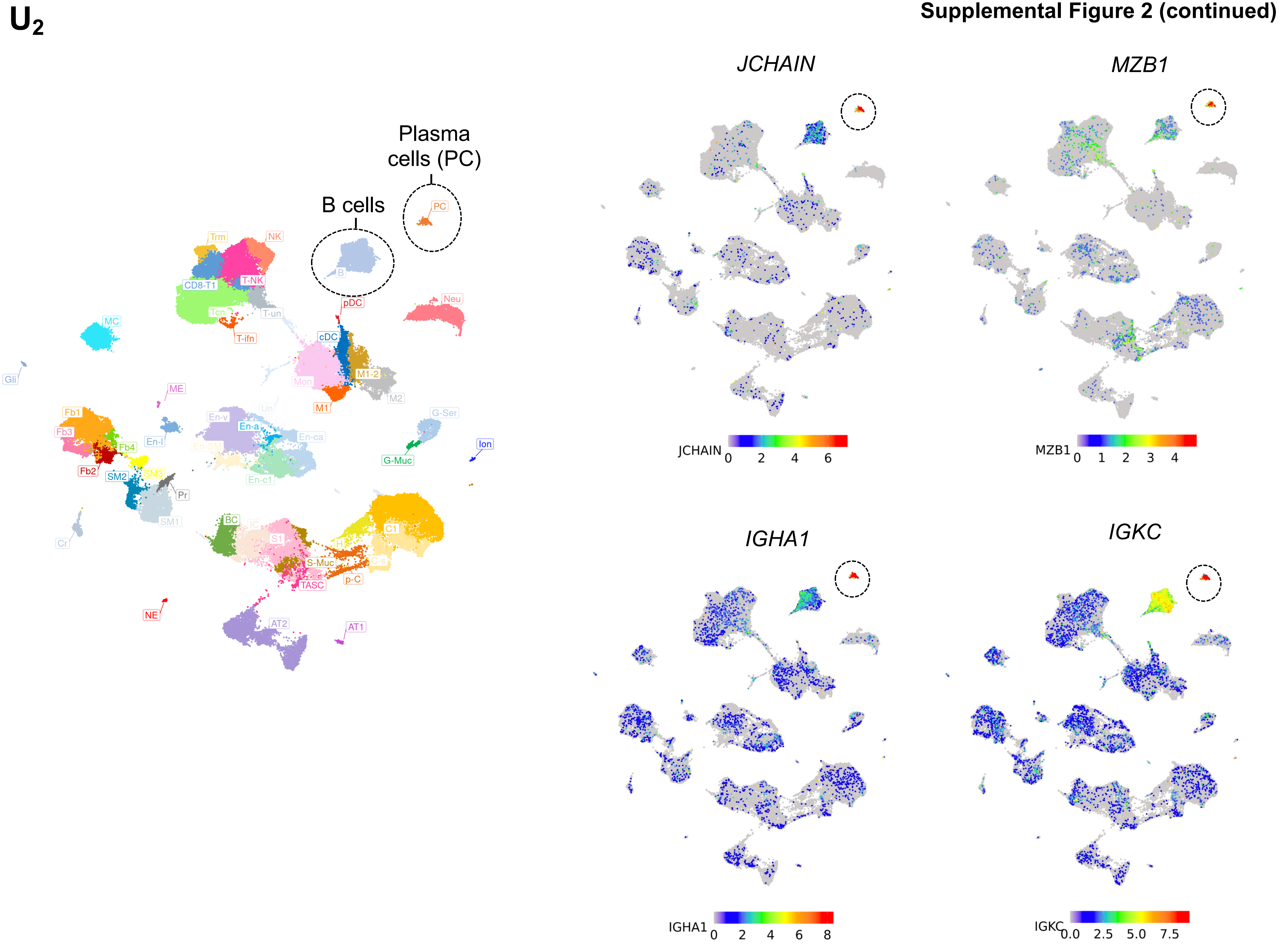

Supplemental Figure 2 (continued)

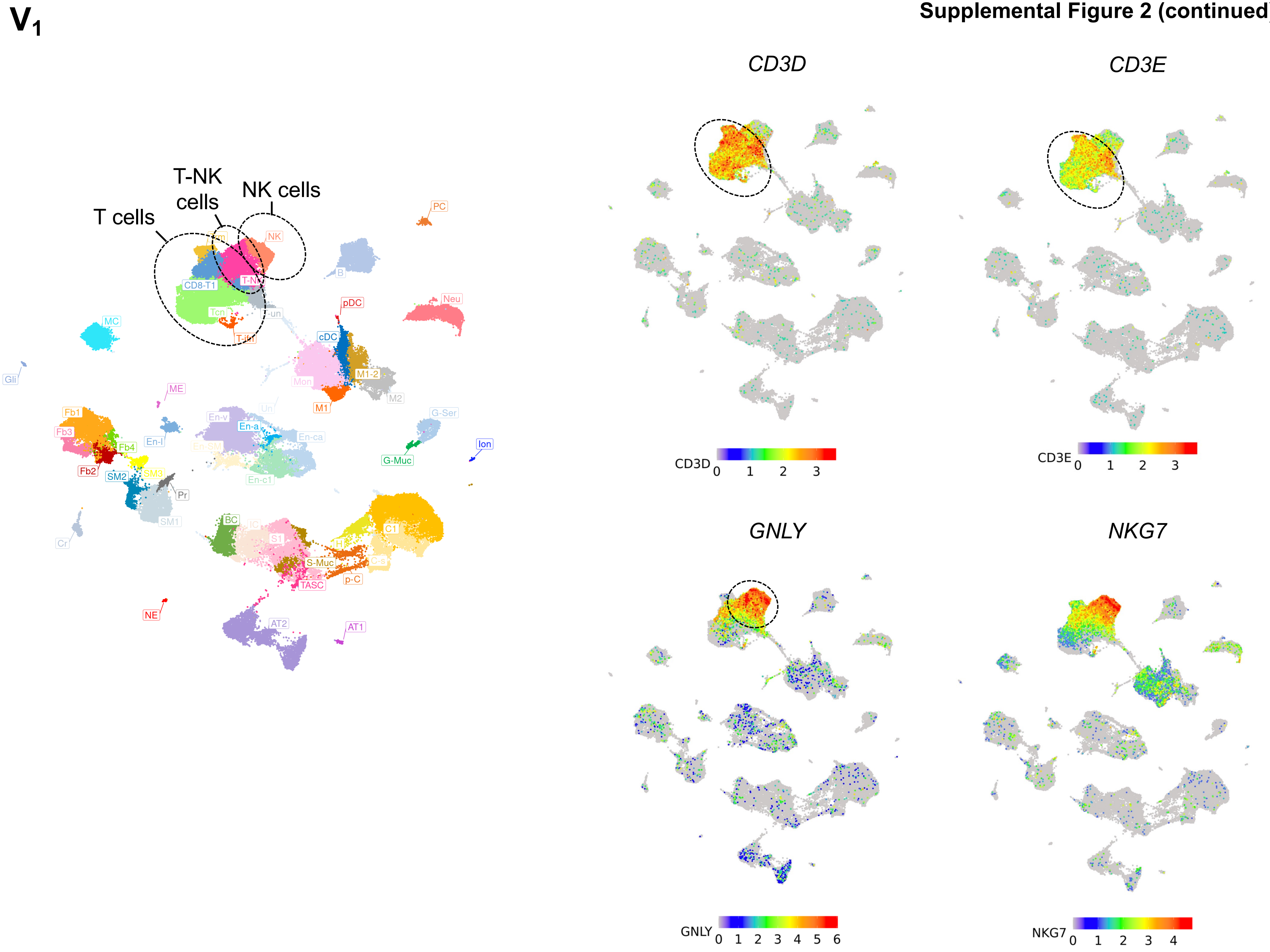

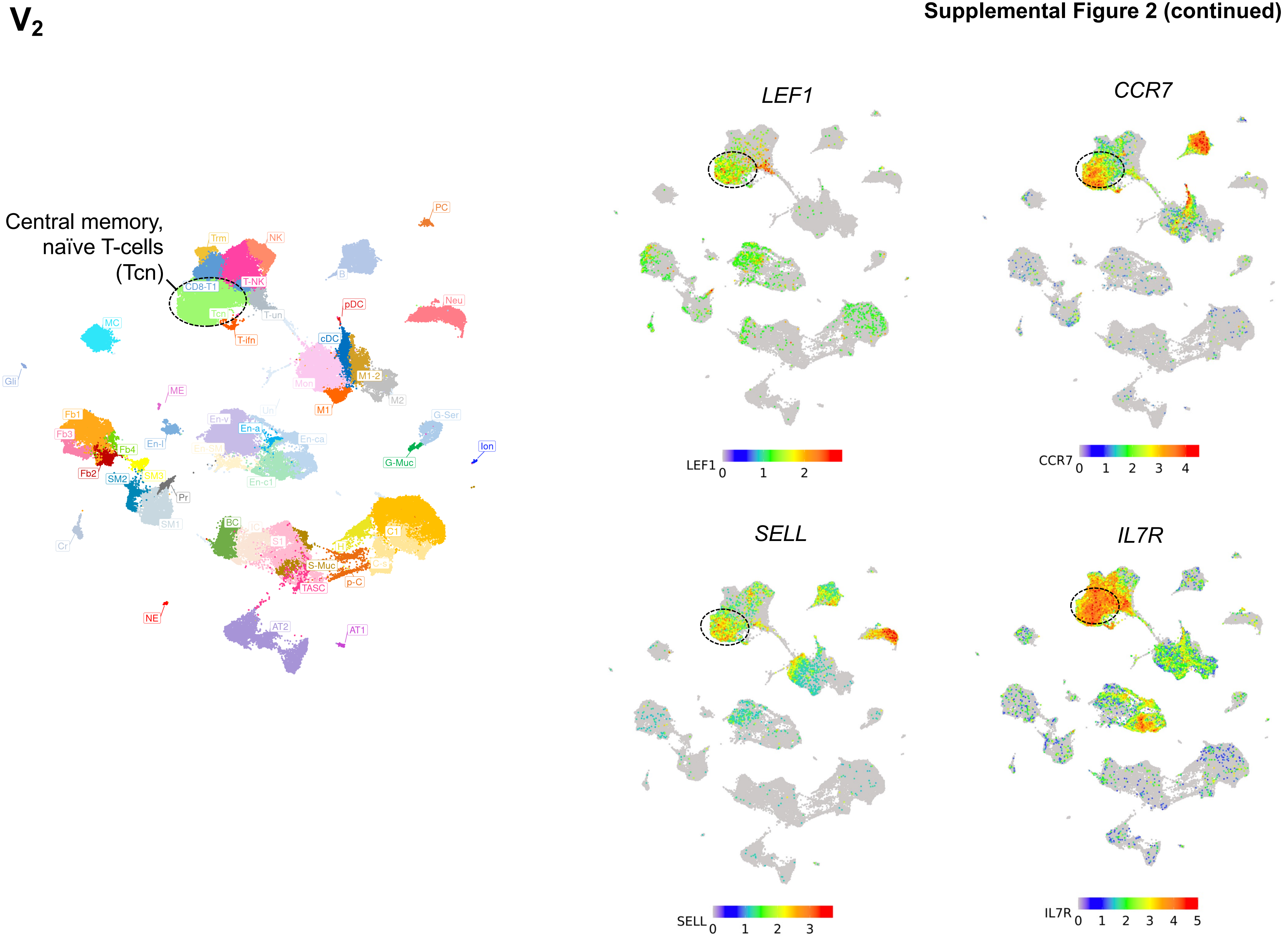

V2
Central memory, naïve T-cells (Tcn)
PC
Trm
NK
B
CD8-T1
T-NK
Tcn
T-un
pDC
Neu
MC
T-ifn
cDC
Gli
M1-2
Mon
ME
M2
M1
Un
G-Ser
Fb1
En-v
En-a
Fb3
En-ca
Ion
En-l
En-SM
Fb4
G-Muc
SM3
Fb2
SM2
En-c1
Pr
SM1
Cr
BC
IC
S1
C1
H
C-s
S-Muc
p-C
TASC
NE
AT2
AT1
LEF1
CCR7
SELL
IL7R

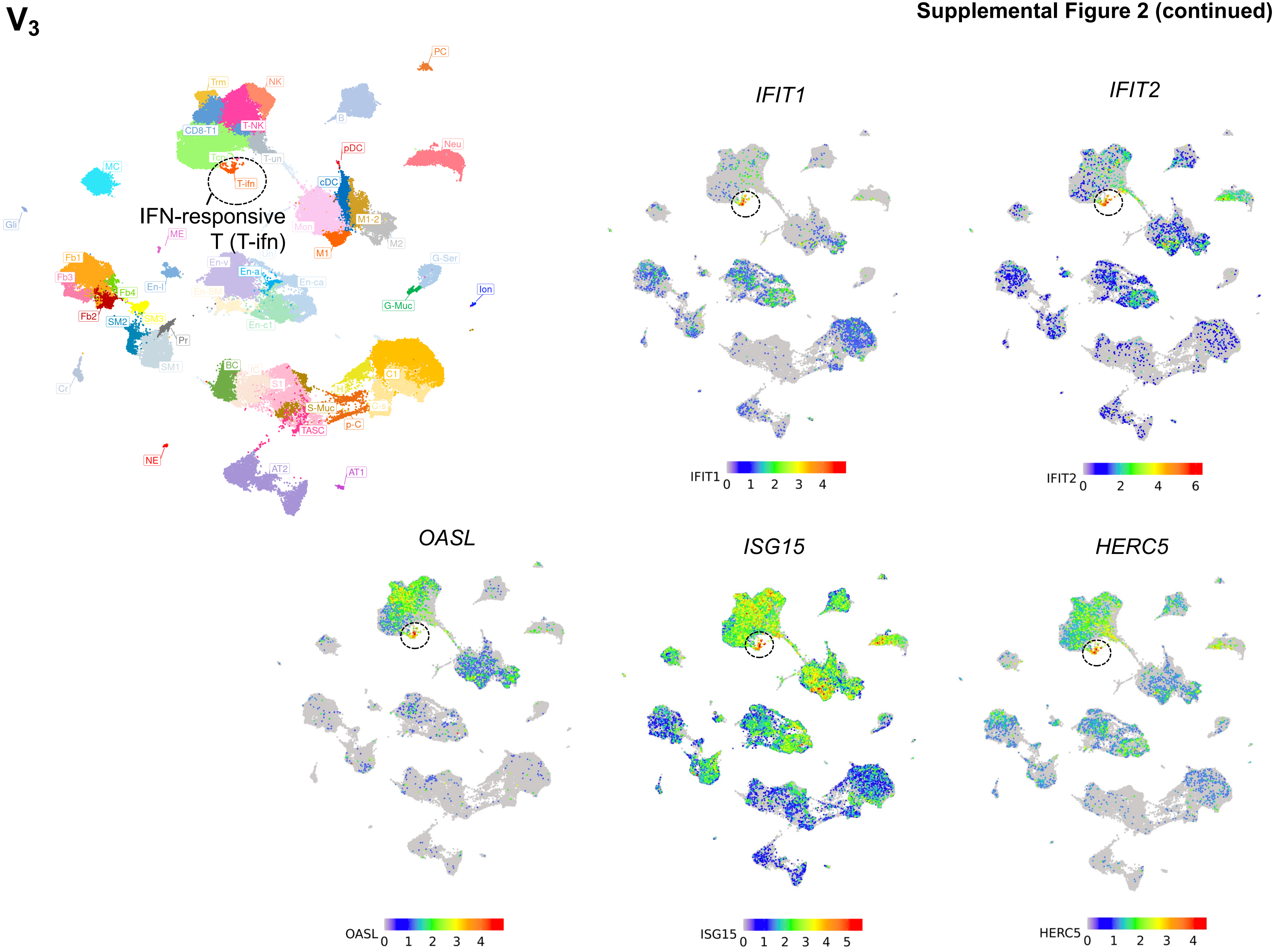
V3
PC
Trm
NK
B
CD8-T1
T-NK
Neu
pDC
T-un
MC
T-ifn
cDC
Gli
IFN-responsive
T (T-ifn)
Mon
M1-2
ME
M2
M1
G-Ser
En-v
Fb1
En-a
En-ca
Fb3
En-l
Ion
Fb4
G-Muc
Fb2
SM2
SM3
En-c1
Pr
SM1
Cr
BC
S1
C1
S-Muc
TASC
p-C
NE
AT2
AT1
IFIT1
IFIT2
IFIT1 0 1 2 3 4
IFIT2 0 2 4 6
OASL
ISG15
HERC5
OASL 0 1 2 3 4
ISG15 0 1 2 3 4 5
HERC5 0 1 2 3 4

**Supplemental Figure 2 (continued)**

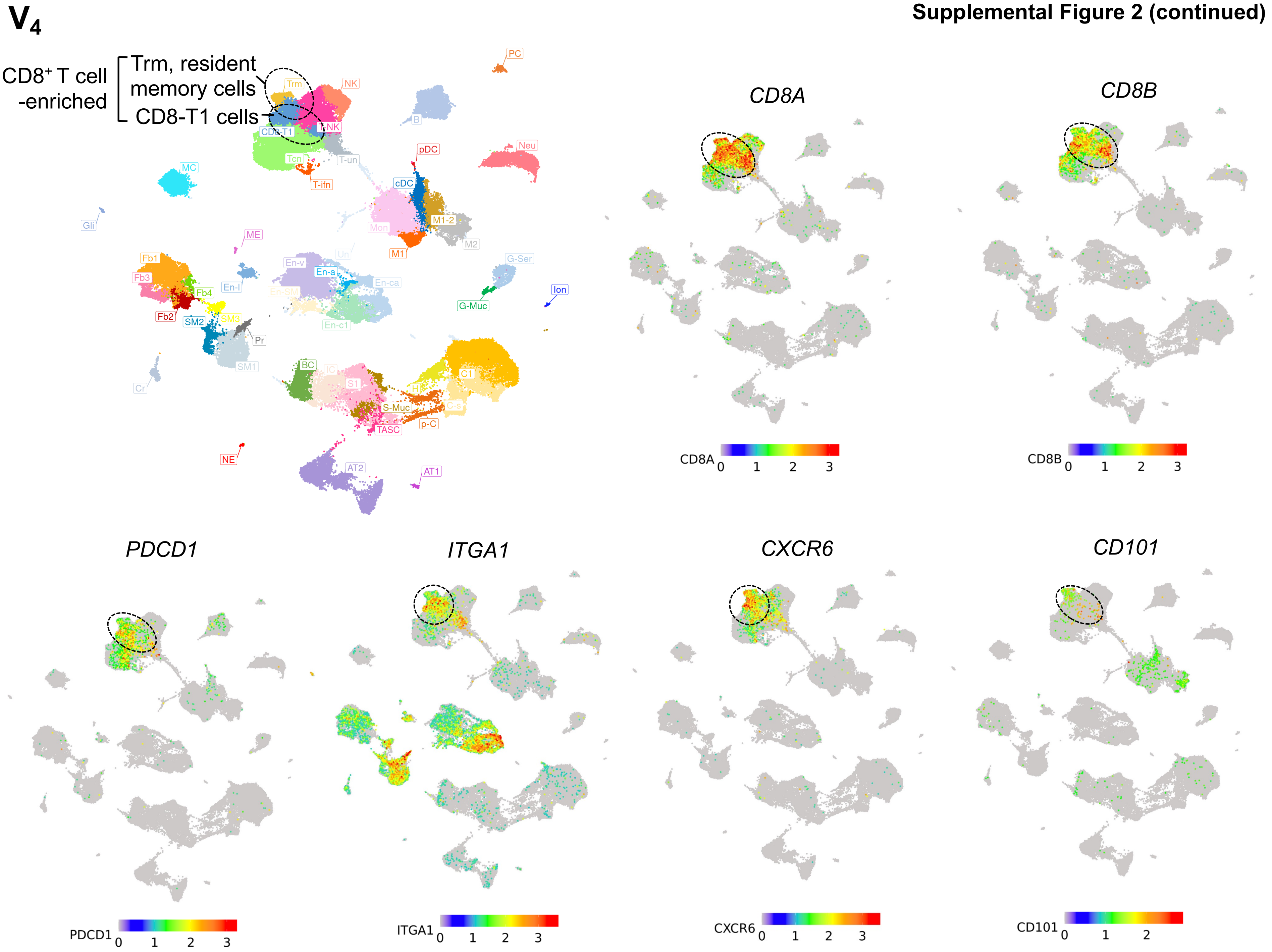

Supplemental Figure 2 (continued)

V5

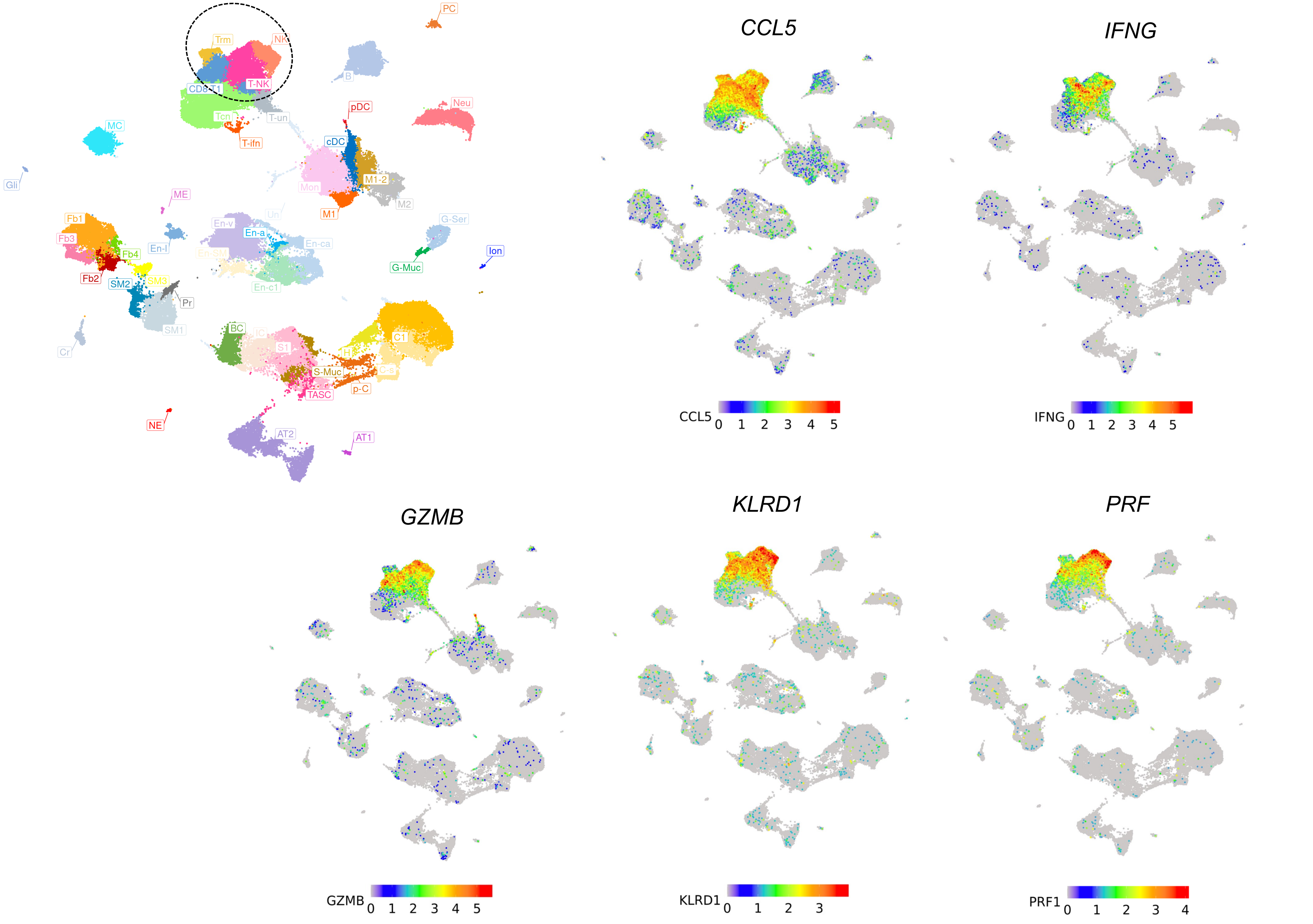

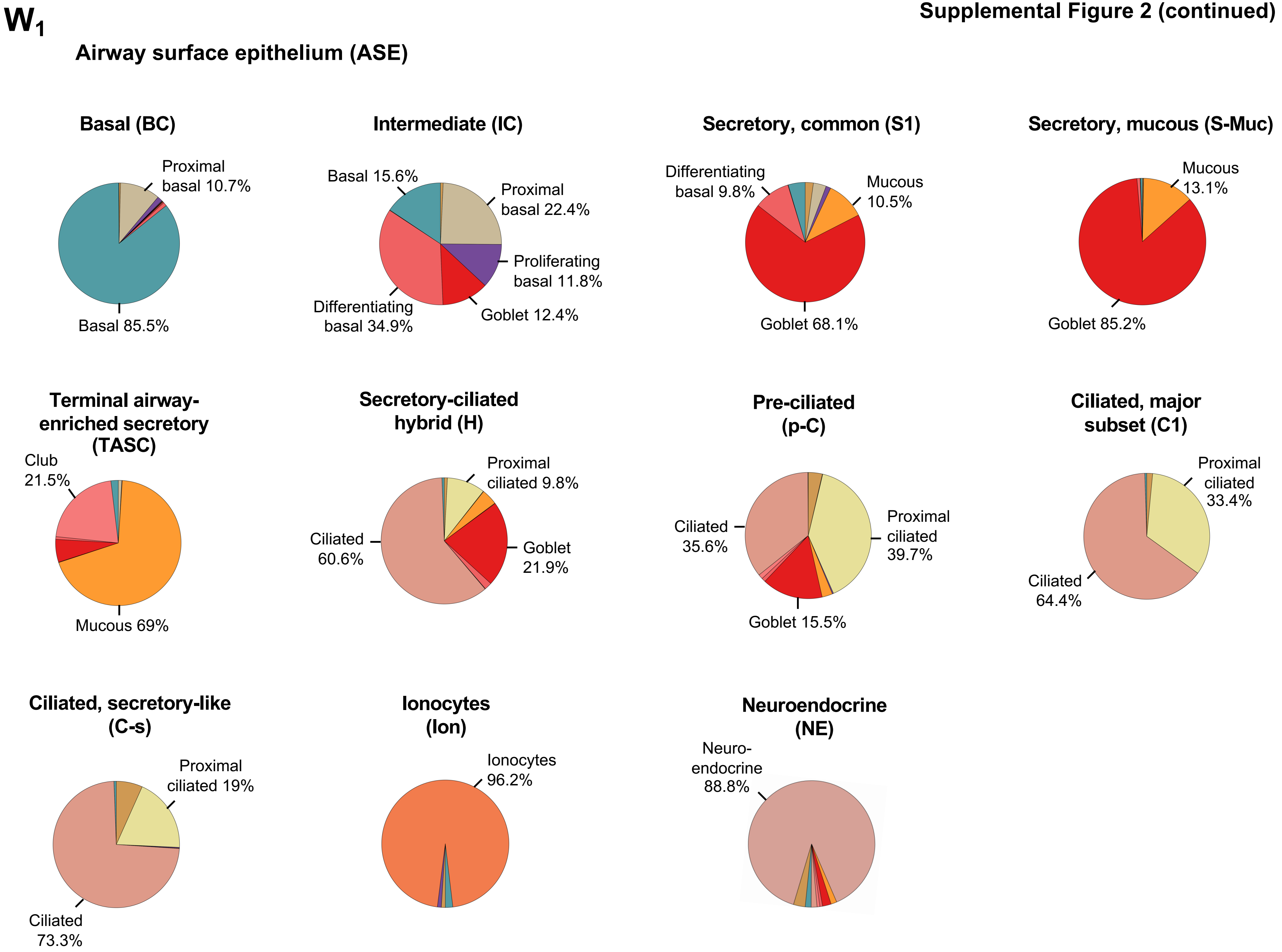
W1
Airway surface epithelium (ASE)
Basal (BC)
Proximal basal 10.7%
Basal 85.5%
Intermediate (IC)
Basal 15.6%
Proximal basal 22.4%
Proliferating basal 11.8%
Goblet 12.4%
Differentiating basal 34.9%
Secretory, common (S1)
Differentiating basal 9.8%
Mucous 10.5%
Goblet 68.1%
Secretory, mucous (S-Muc)
Mucous 13.1%
Goblet 85.2%
Terminal airway-enriched secretory (TASC)
Club 21.5%
Mucous 69%
Secretory-ciliated hybrid (H)
Proximal ciliated 9.8%
Ciliated 60.6%
Goblet 21.9%
Pre-ciliated (p-C)
Ciliated 35.6%
Proximal ciliated 39.7%
Goblet 15.5%
Ciliated, major subset (C1)
Proximal ciliated 33.4%
Ciliated 64.4%
Ciliated, secretory-like (C-s)
Proximal ciliated 19%
Ciliated 73.3%
Ionocytes (Ion)
Ionocytes 96.2%
Neuroendocrine (NE)
Neuro-endocrine 88.8%

W$_2$



#### Submucosal gland (SMG)

##### Myoepithelial (ME)

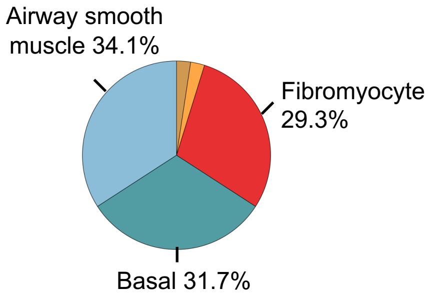


##### SMG Mucous (G-Muc)

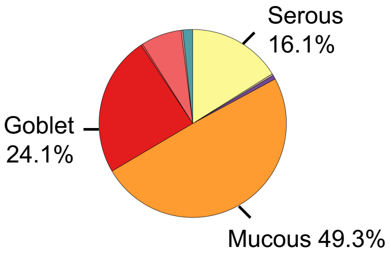


##### Secretory, common (S1)

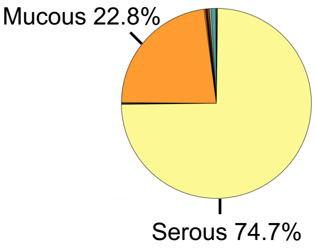


#### Alveolar epithelium (AT)

##### Alveolar, type 1 (AT1)

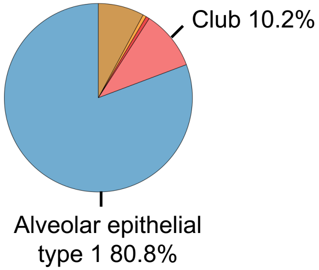


##### Alveolar, type 2 (AT2)

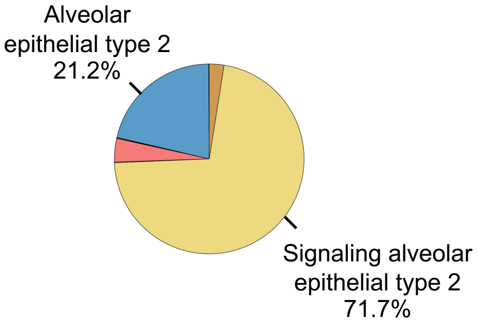

Supplemental Figure 2 (continued)

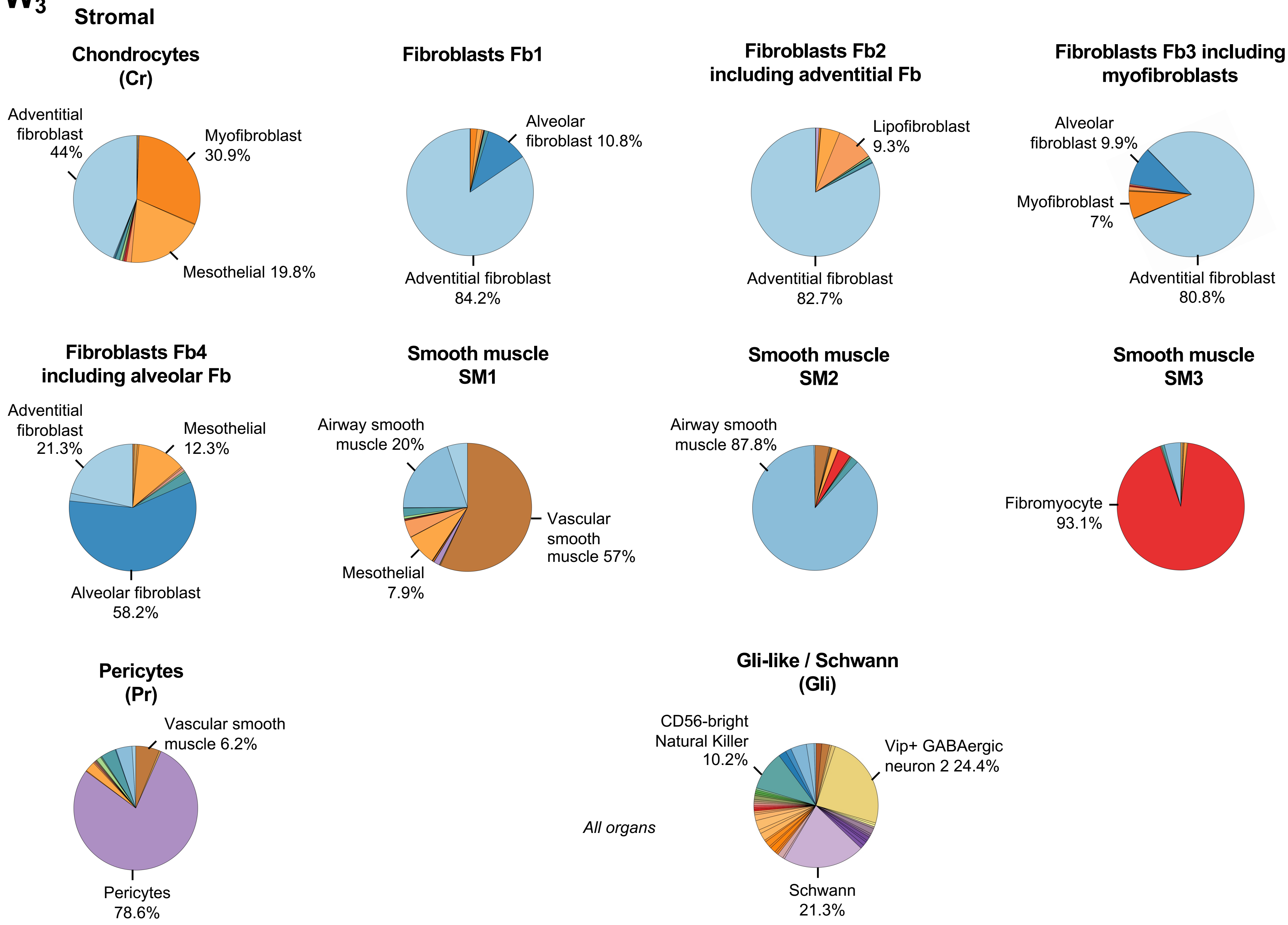

Supplemental Figure 2 (continued)

$W_4$

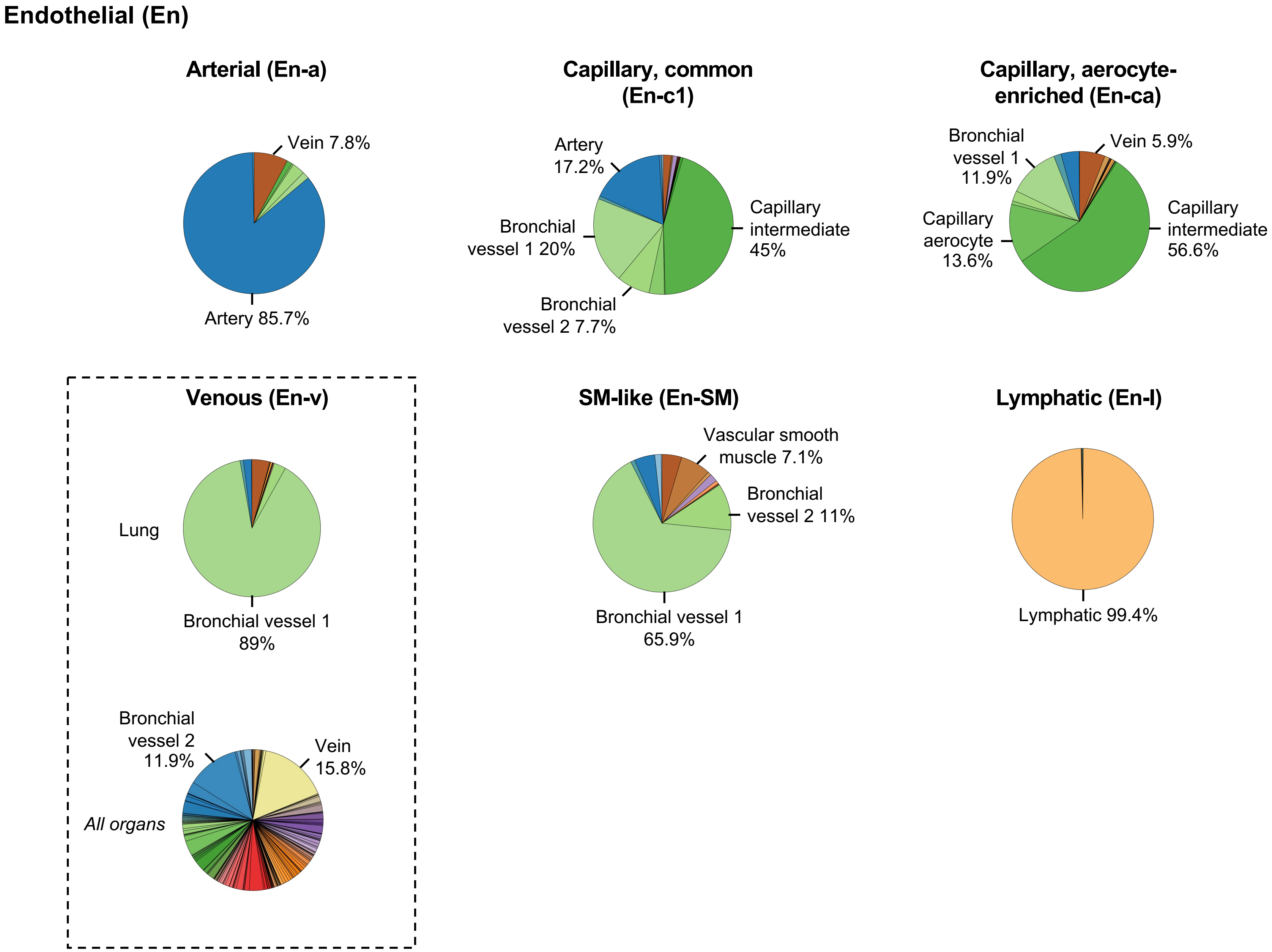

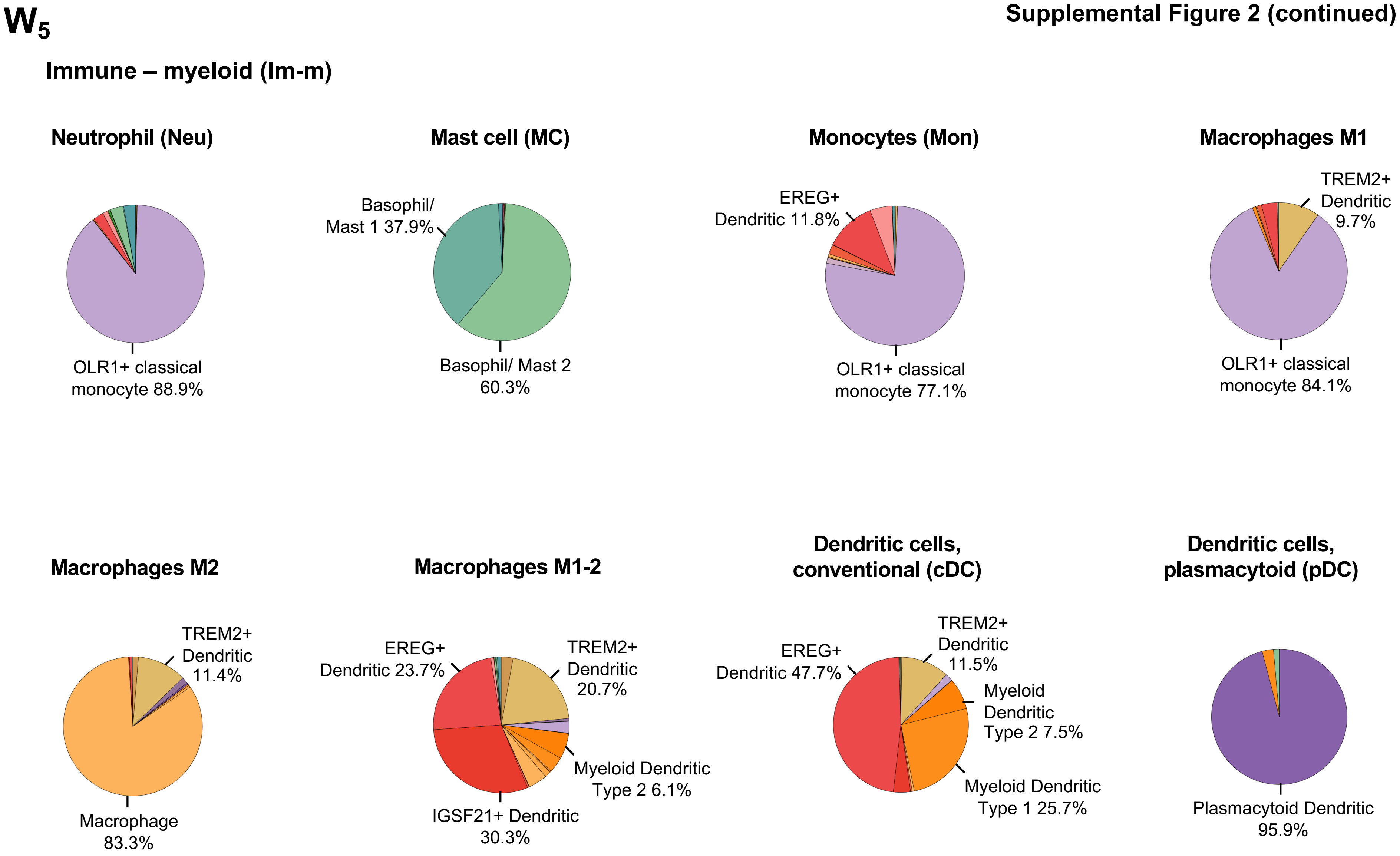
W5
Immune – myeloid (Im-m)
Neutrophil (Neu)
OLR1+ classical monocyte 88.9%
Mast cell (MC)
Basophil/ Mast 1 37.9%
Basophil/ Mast 2 60.3%
Monocytes (Mon)
EREG+ Dendritic 11.8%
OLR1+ classical monocyte 77.1%
Macrophages M1
TREM2+ Dendritic 9.7%
OLR1+ classical monocyte 84.1%
Macrophages M2
TREM2+ Dendritic 11.4%
Macrophage 83.3%
Macrophages M1-2
EREG+ Dendritic 23.7%
TREM2+ Dendritic 20.7%
Myeloid Dendritic Type 2 6.1%
IGSF21+ Dendritic 30.3%
Dendritic cells, conventional (cDC)
EREG+ Dendritic 47.7%
TREM2+ Dendritic 11.5%
Myeloid Dendritic Type 2 7.5%
Myeloid Dendritic Type 1 25.7%
Dendritic cells, plasmacytoid (pDC)
Plasmacytoid Dendritic 95.9%

**Supplemental Figure 2 (continued)**

**$W_6$**

#### Immune – lymphoid (Im-l)

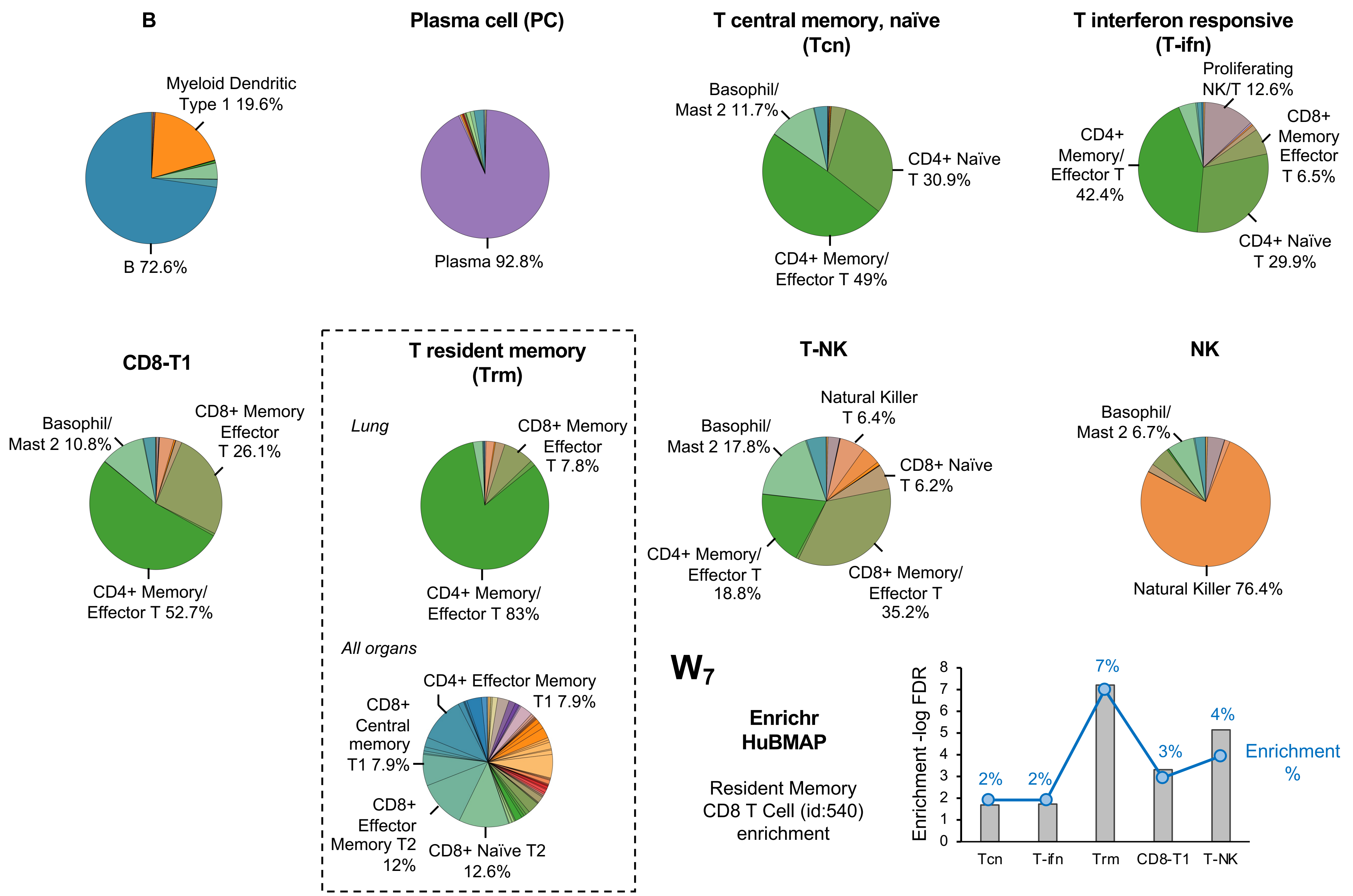

**X₁**

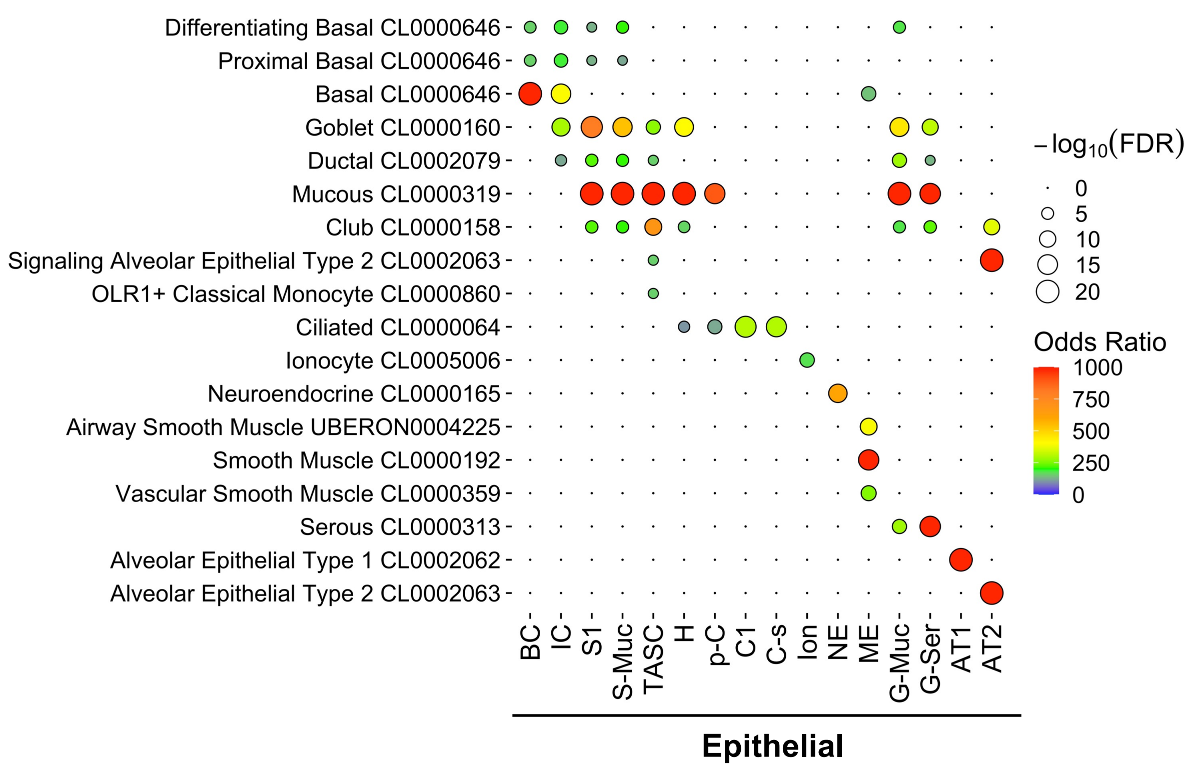
Differentiating Basal CL0000646
Proximal Basal CL0000646
Basal CL0000646
Goblet CL0000160
Ductal CL0002079
Mucous CL0000319
Club CL0000158
Signaling Alveolar Epithelial Type 2 CL0002063
OLR1+ Classical Monocyte CL0000860
Ciliated CL0000064
Ionocyte CL0005006
Neuroendocrine CL0000165
Airway Smooth Muscle UBERON0004225
Smooth Muscle CL0000192
Vascular Smooth Muscle CL0000359
Serous CL0000313
Alveolar Epithelial Type 1 CL0002062
Alveolar Epithelial Type 2 CL0002063
BC
IC
S1
S-Muc
TASC
H
p-C
C1
C-s
Ion
NE
ME
G-Muc
G-Ser
AT1
AT2
Epithelial
$-\log_{10}(\mathrm{FDR})$
0
5
10
15
20
Odds Ratio
1000
750
500
250
0

Supplemental Figure 2 (continued)

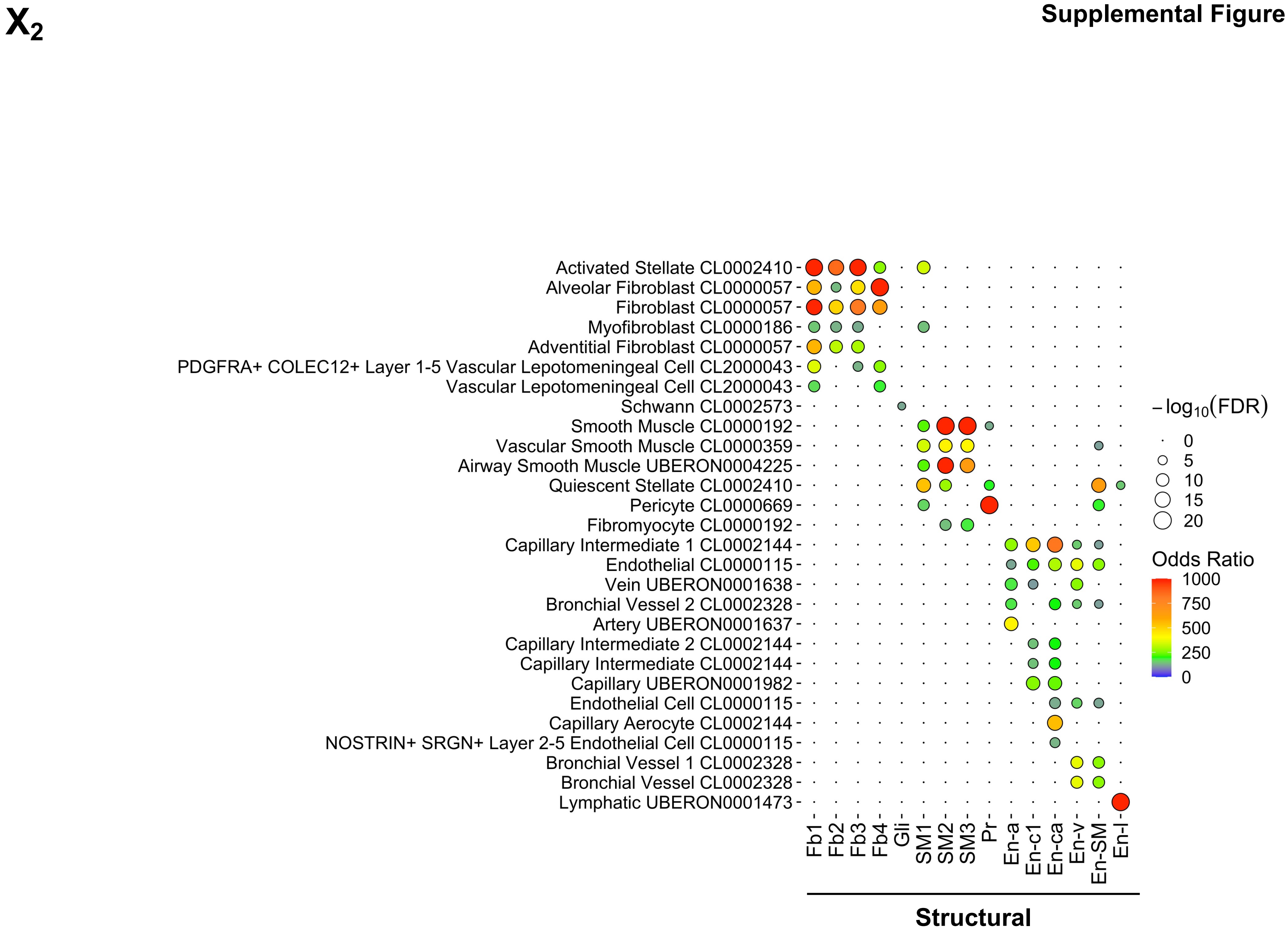

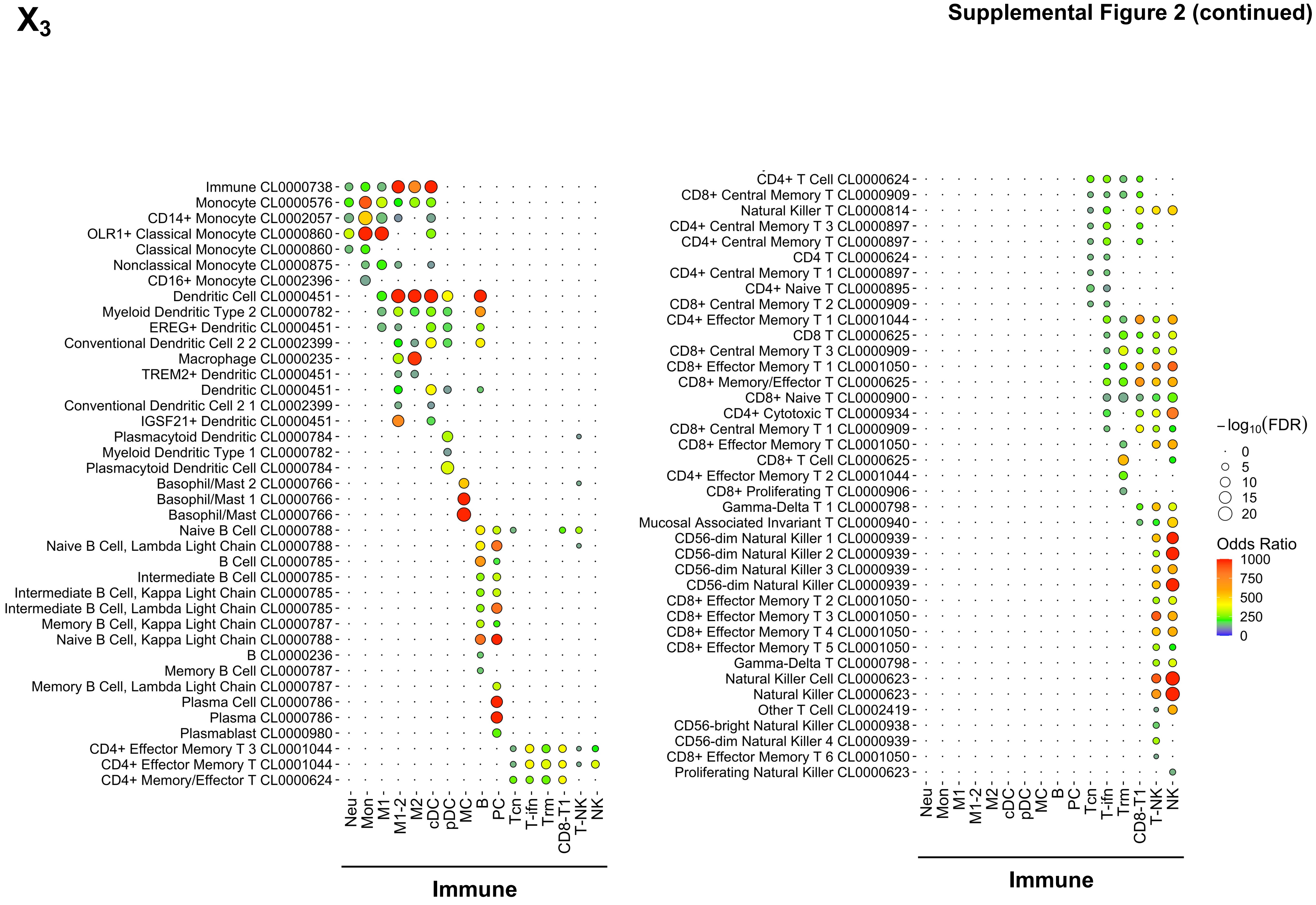
X3
Immune CL0000738
Monocyte CL0000576
CD14+ Monocyte CL0002057
OLR1+ Classical Monocyte CL0000860
Classical Monocyte CL0000860
Nonclassical Monocyte CL0000875
CD16+ Monocyte CL0002396
Dendritic Cell CL0000451
Myeloid Dendritic Type 2 CL0000782
EREG+ Dendritic CL0000451
Conventional Dendritic Cell 2 2 CL0002399
Macrophage CL0000235
TREM2+ Dendritic CL0000451
Dendritic CL0000451
Conventional Dendritic Cell 2 1 CL0002399
IGSF21+ Dendritic CL0000451
Plasmacytoid Dendritic CL0000784
Myeloid Dendritic Type 1 CL0000782
Plasmacytoid Dendritic Cell CL0000784
Basophil/Mast 2 CL0000766
Basophil/Mast 1 CL0000766
Basophil/Mast CL0000766
Naive B Cell CL0000788
Naive B Cell, Lambda Light Chain CL0000788
B Cell CL0000785
Intermediate B Cell CL0000785
Intermediate B Cell, Kappa Light Chain CL0000785
Intermediate B Cell, Lambda Light Chain CL0000785
Memory B Cell, Kappa Light Chain CL0000787
Naive B Cell, Kappa Light Chain CL0000788
B CL0000236
Memory B Cell CL0000787
Memory B Cell, Lambda Light Chain CL0000787
Plasma Cell CL0000786
Plasma CL0000786
Plasmablast CL0000980
CD4+ Effector Memory T 3 CL0001044
CD4+ Effector Memory T CL0001044
CD4+ Memory/Effector T CL0000624
Neu
Mon
M1
M1-2
M2
cDC
pDC
MC
B
PC
Tcn
T-ifn
Trm
CD8-T1
T-NK
NK
Immune
CD4+ T Cell CL0000624
CD8+ Central Memory T CL0000909
Natural Killer T CL0000814
CD4+ Central Memory T 3 CL0000897
CD4+ Central Memory T CL0000897
CD4 T CL0000624
CD4+ Central Memory T 1 CL0000897
CD4+ Naive T CL0000895
CD8+ Central Memory T 2 CL0000909
CD4+ Effector Memory T 1 CL0001044
CD8 T CL0000625
CD8+ Central Memory T 3 CL0000909
CD8+ Effector Memory T 1 CL0001050
CD8+ Memory/Effector T CL0000625
CD8+ Naive T CL0000900
CD4+ Cytotoxic T CL0000934
CD8+ Central Memory T 1 CL0000909
CD8+ Effector Memory T CL0001050
CD8+ T Cell CL0000625
CD4+ Effector Memory T 2 CL0001044
CD8+ Proliferating T CL0000906
Gamma-Delta T 1 CL0000798
Mucosal Associated Invariant T CL0000940
CD56-dim Natural Killer 1 CL0000939
CD56-dim Natural Killer 2 CL0000939
CD56-dim Natural Killer 3 CL0000939
CD56-dim Natural Killer CL0000939
CD8+ Effector Memory T 2 CL0001050
CD8+ Effector Memory T 3 CL0001050
CD8+ Effector Memory T 4 CL0001050
CD8+ Effector Memory T 5 CL0001050
Gamma-Delta T CL0000798
Natural Killer Cell CL0000623
Natural Killer CL0000623
Other T Cell CL0002419
CD56-bright Natural Killer CL0000938
CD56-dim Natural Killer 4 CL0000939
CD8+ Effector Memory T 6 CL0001050
Proliferating Natural Killer CL0000623
Neu
Mon
M1
M1-2
M2
cDC
pDC
MC
B
PC
Tcn
T-ifn
Trm
CD8-T1
T-NK
NK
Immune
−log10(FDR)
0
5
10
15
20
Odds Ratio
1000
750
500
250
0
