## Supplementary material for "A Unique Cellular Organization of Human Distal Airways and Its Disarray in Chronic Obstructive Pulmonary Disease": Fig. S3

**A1**

Proximal (bronchus)

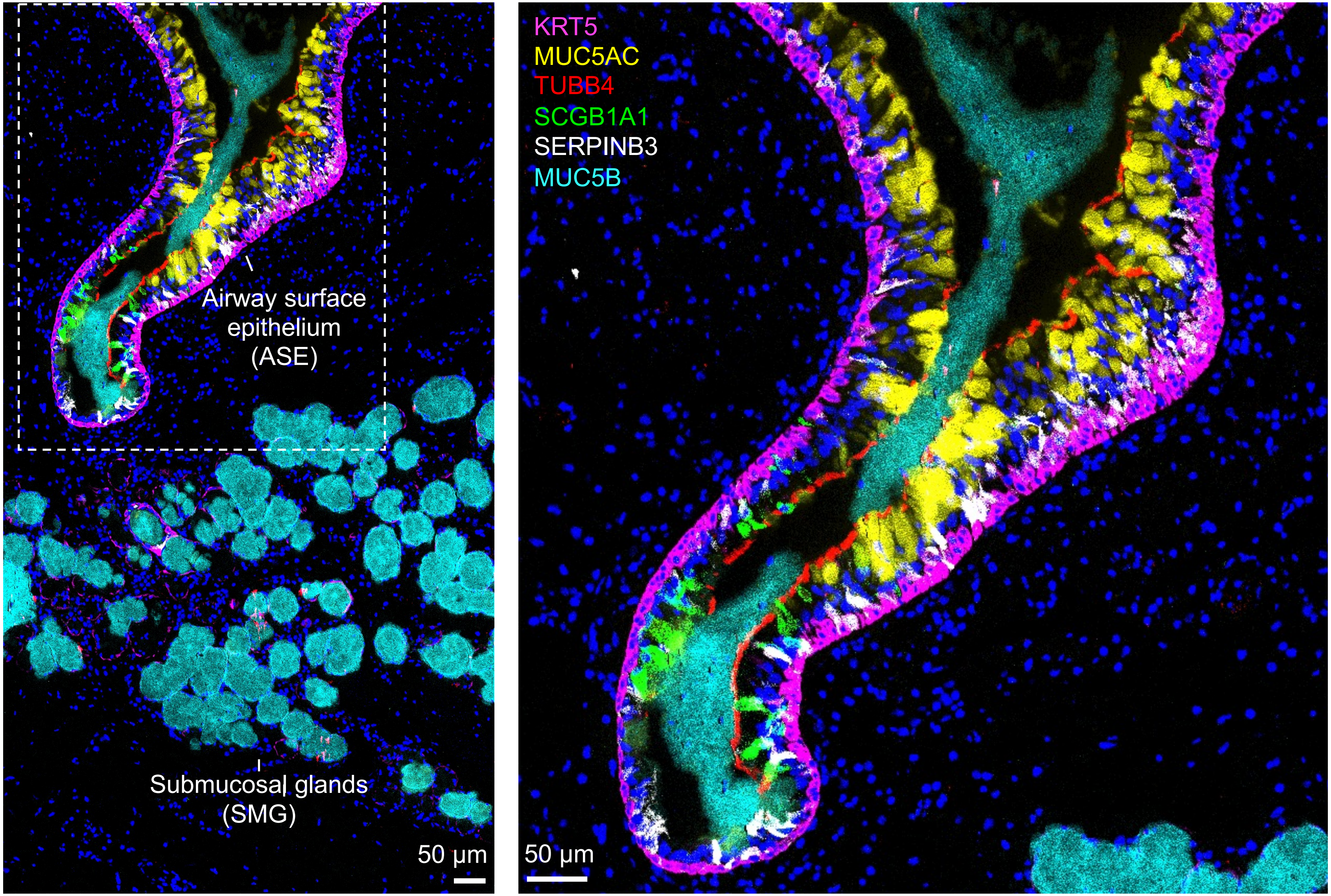

**A2**

Proximal (bronchus)

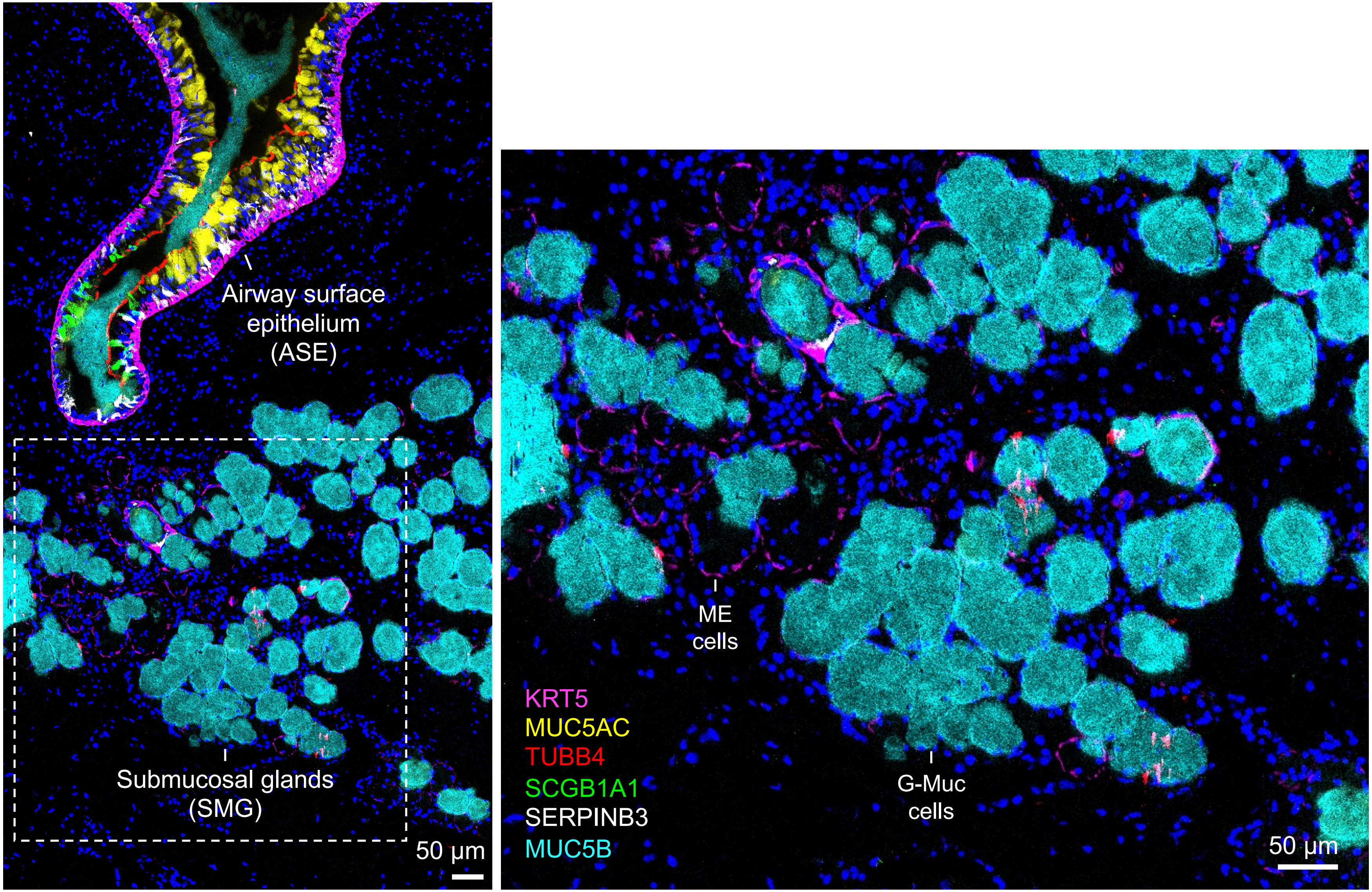

**A3**

Proximal (bronchus)

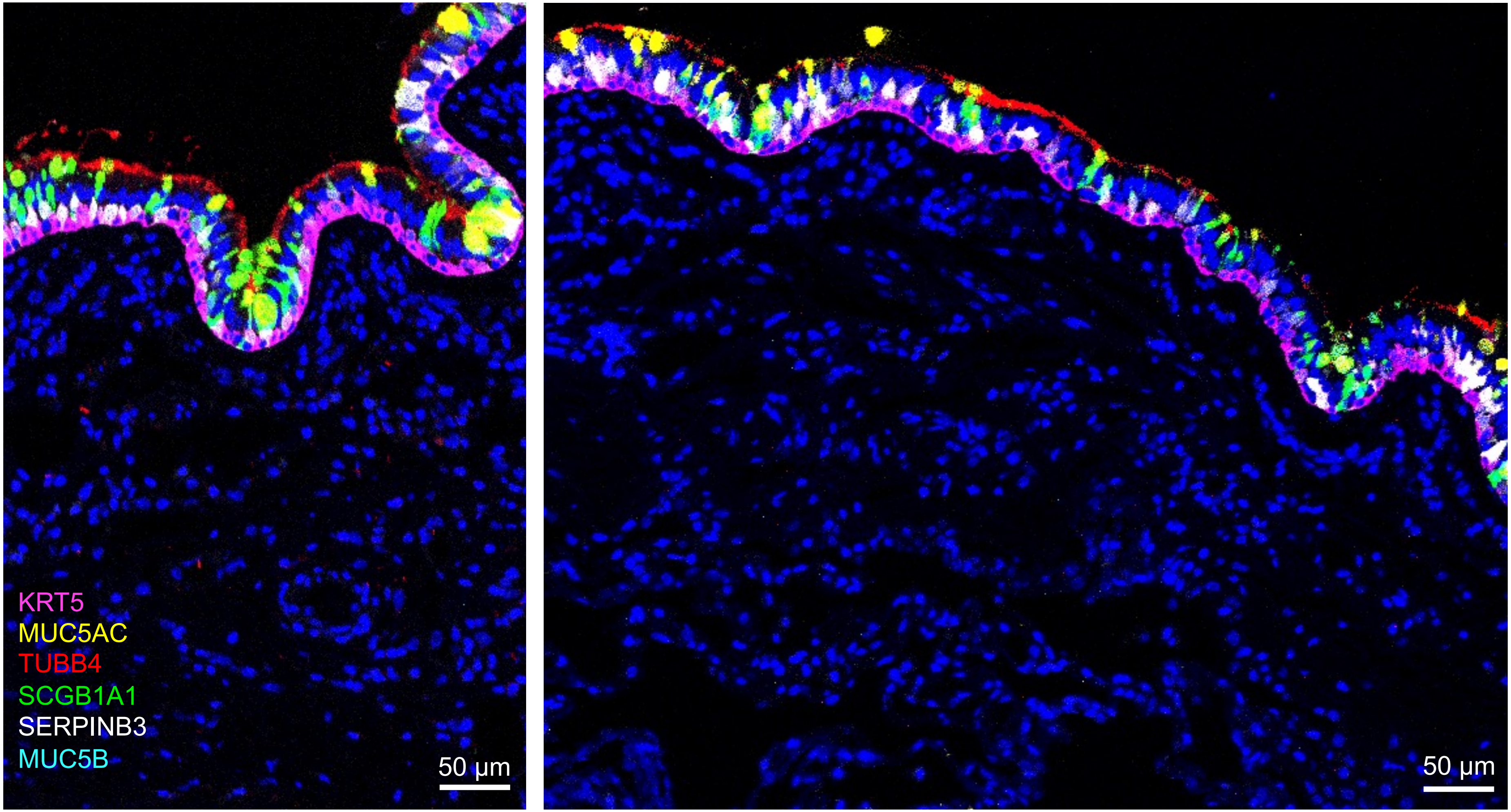

**A$_4$**

Proximal (bronchus)

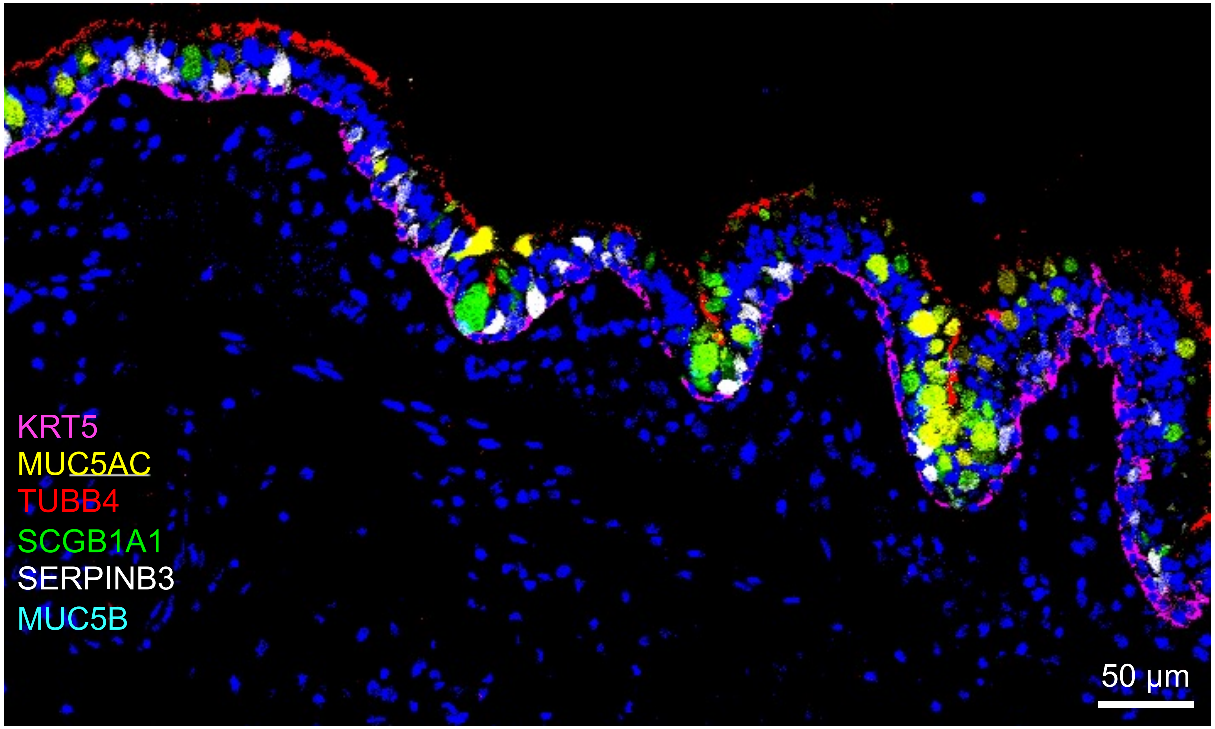

A$_4$

Distal

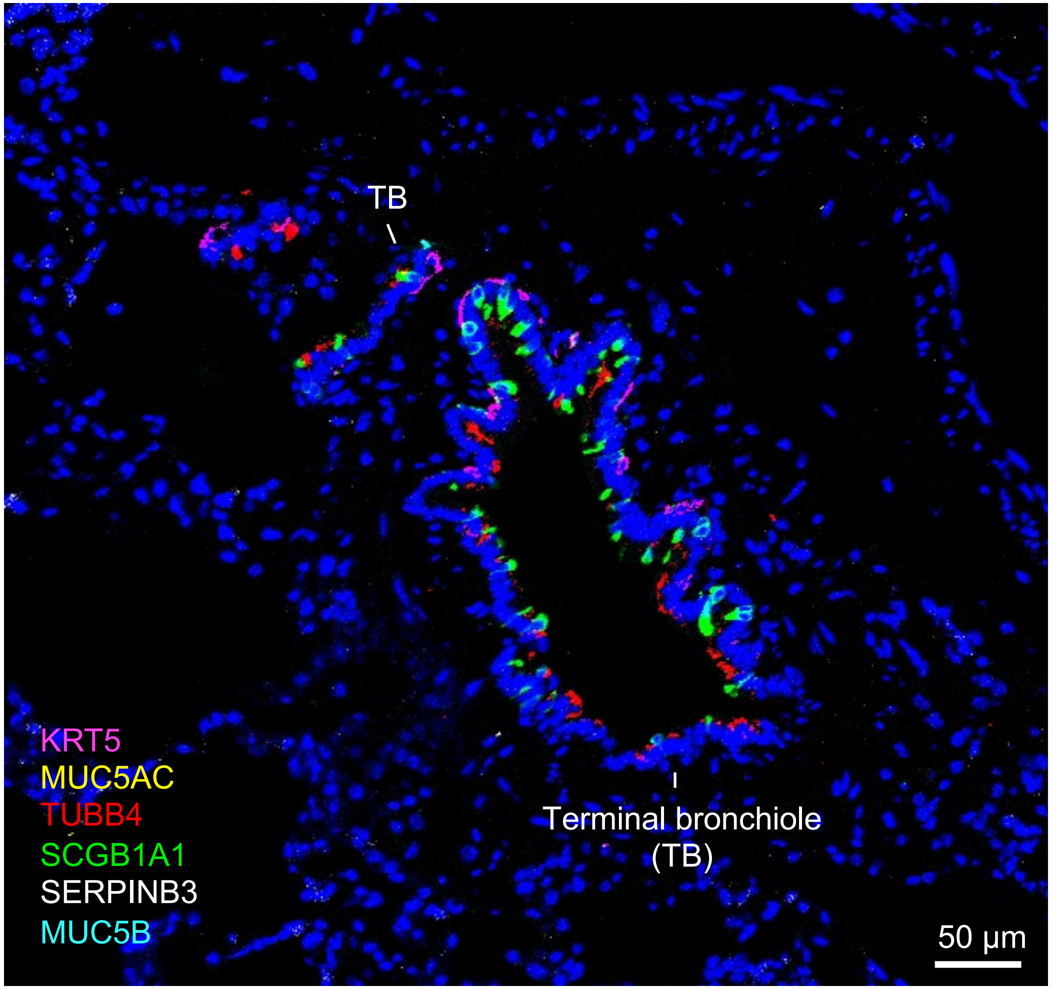
TB
Terminal bronchiole
(TB)
KRT5
MUC5AC
TUBB4
SCGB1A1
SERPINB3
MUC5B
50 µm

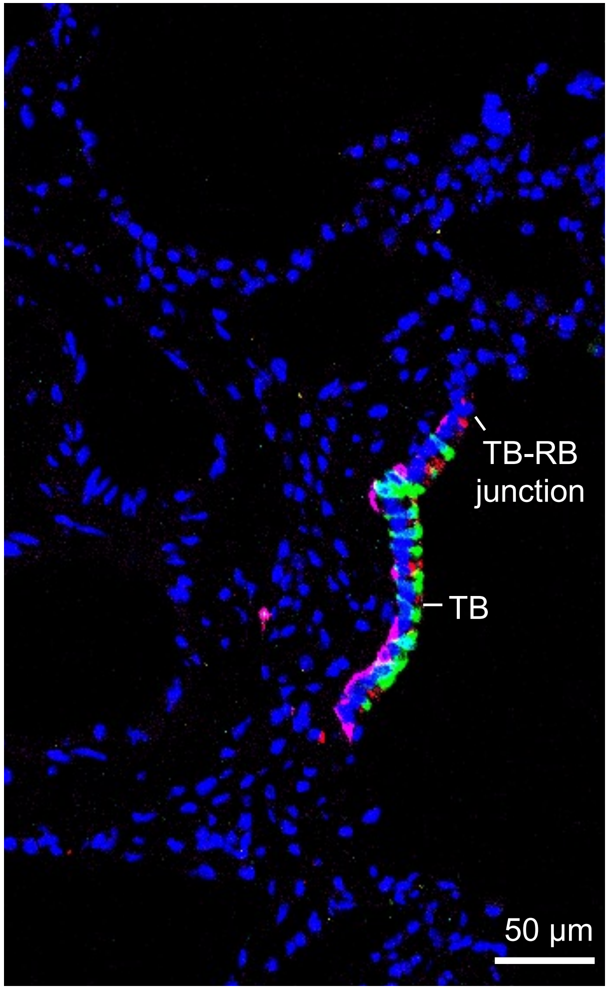
TB-RB
junction
TB
50 µm

**A$_5$**

Distal

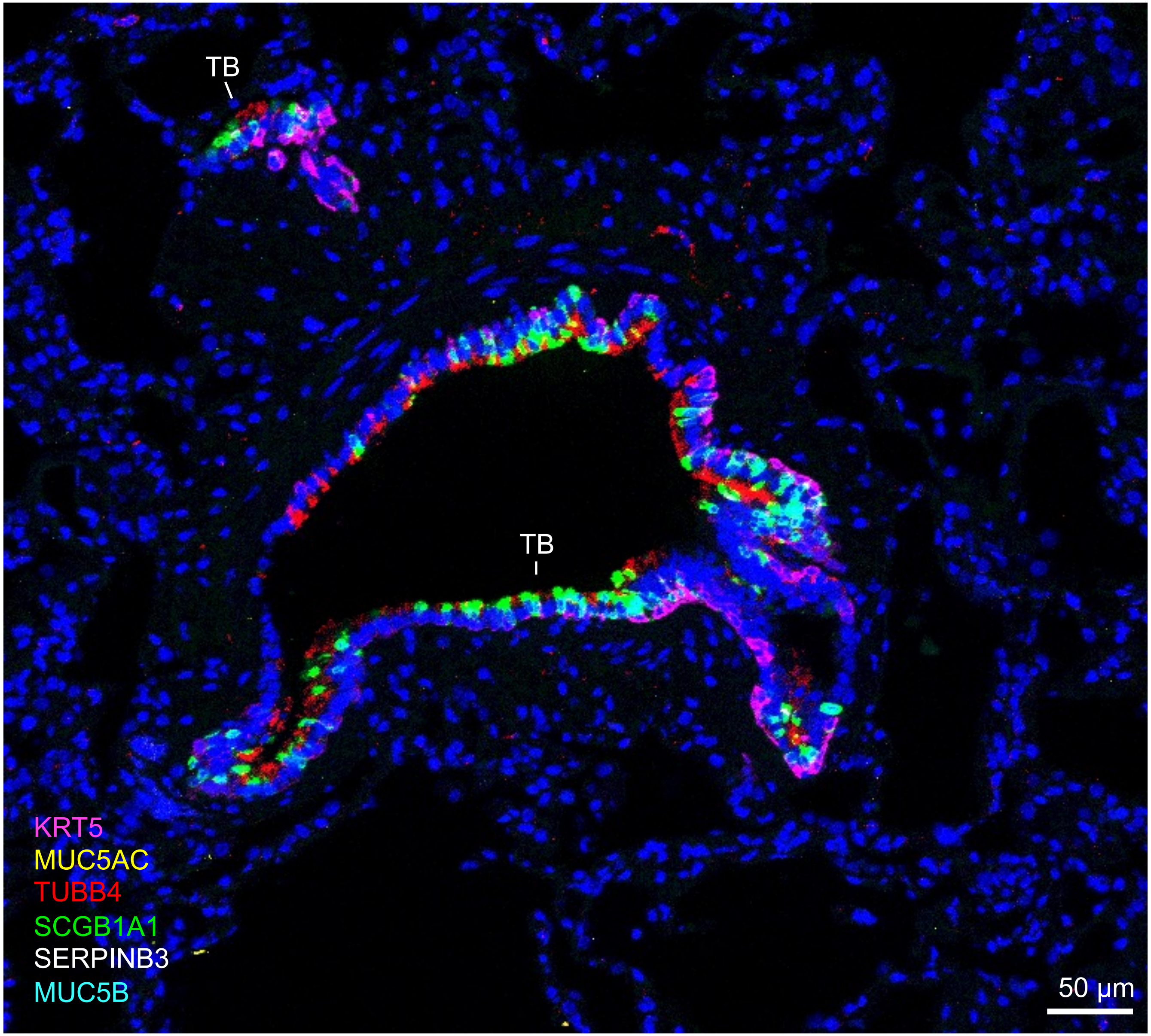

**A$_6$**

Distal

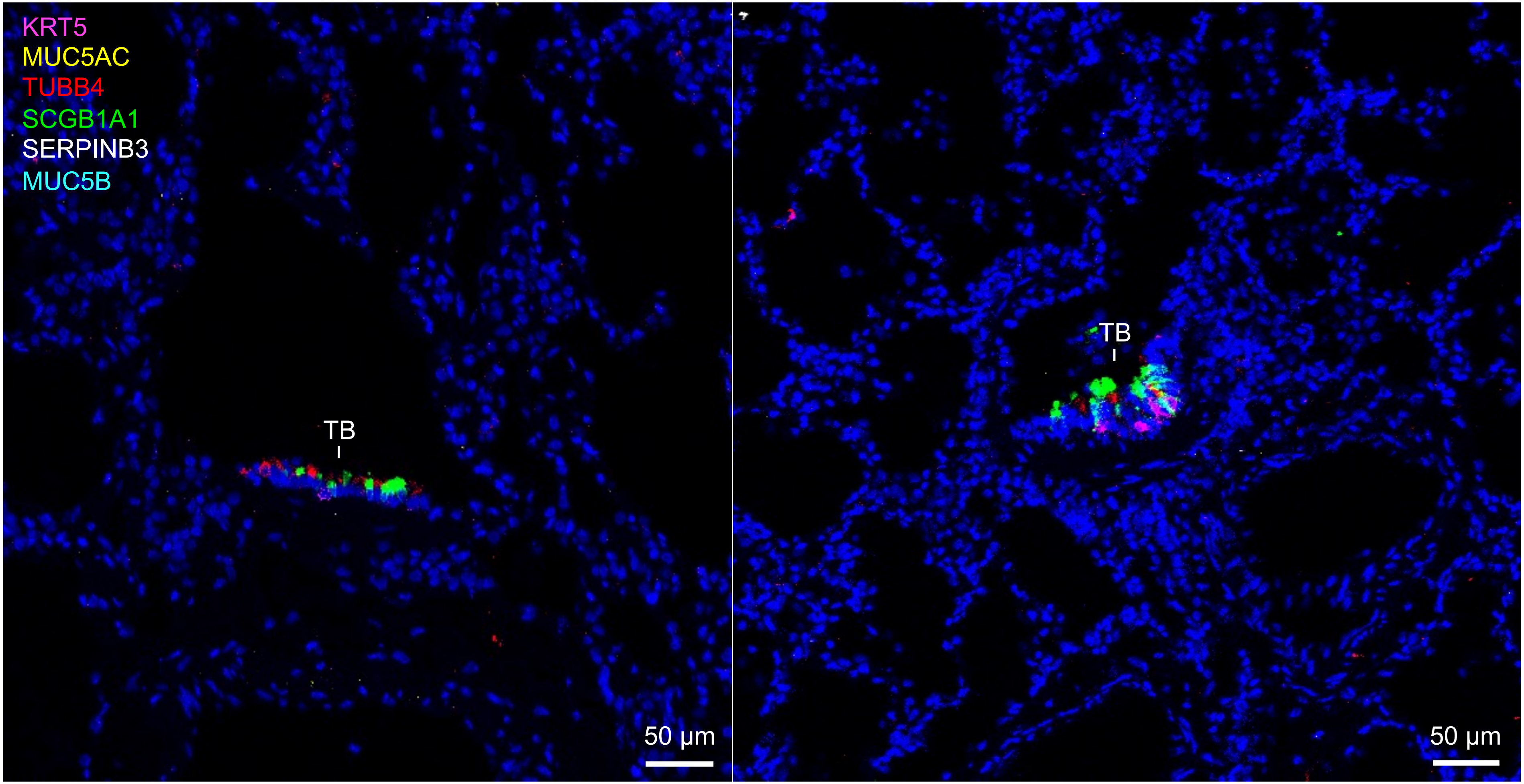

**B$_1$**

Proximal (bronchus)

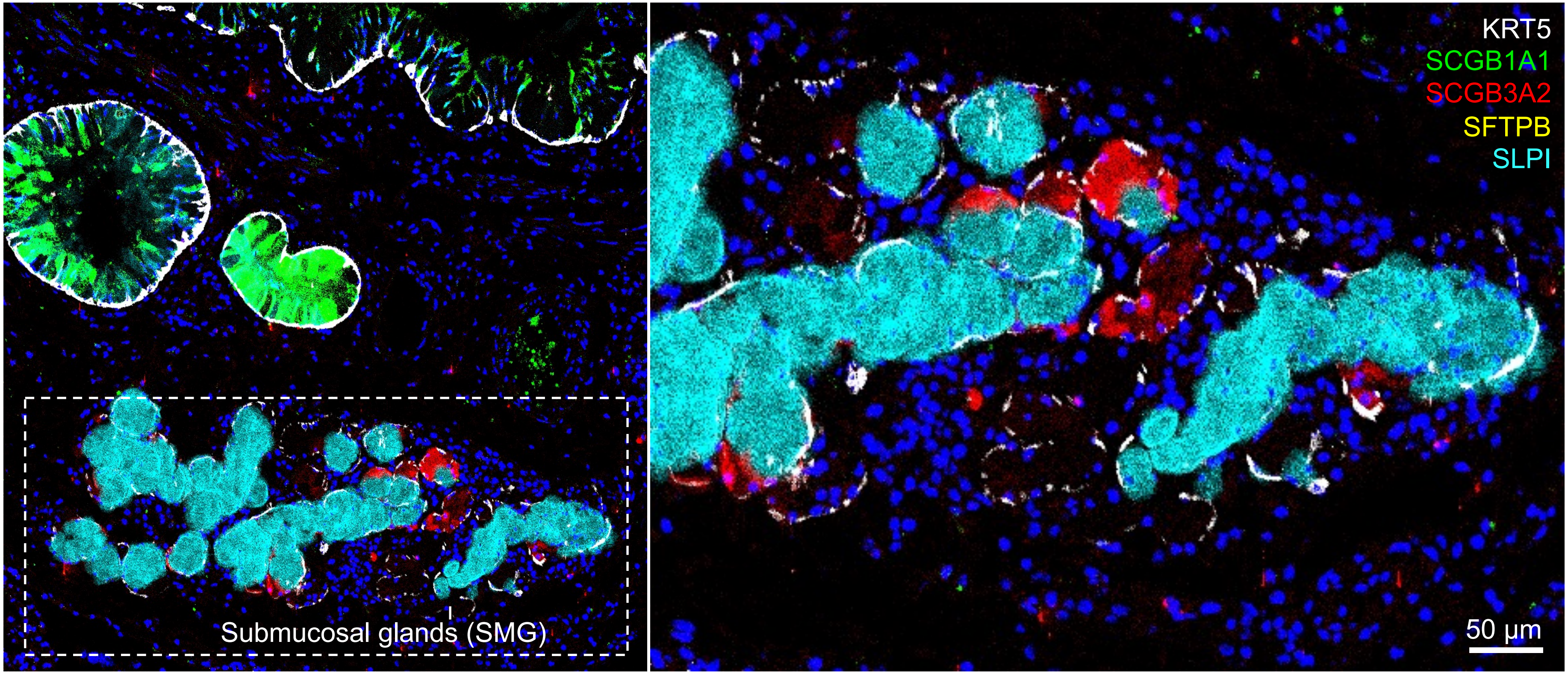

**$B_2$**

Proximal (bronchus)

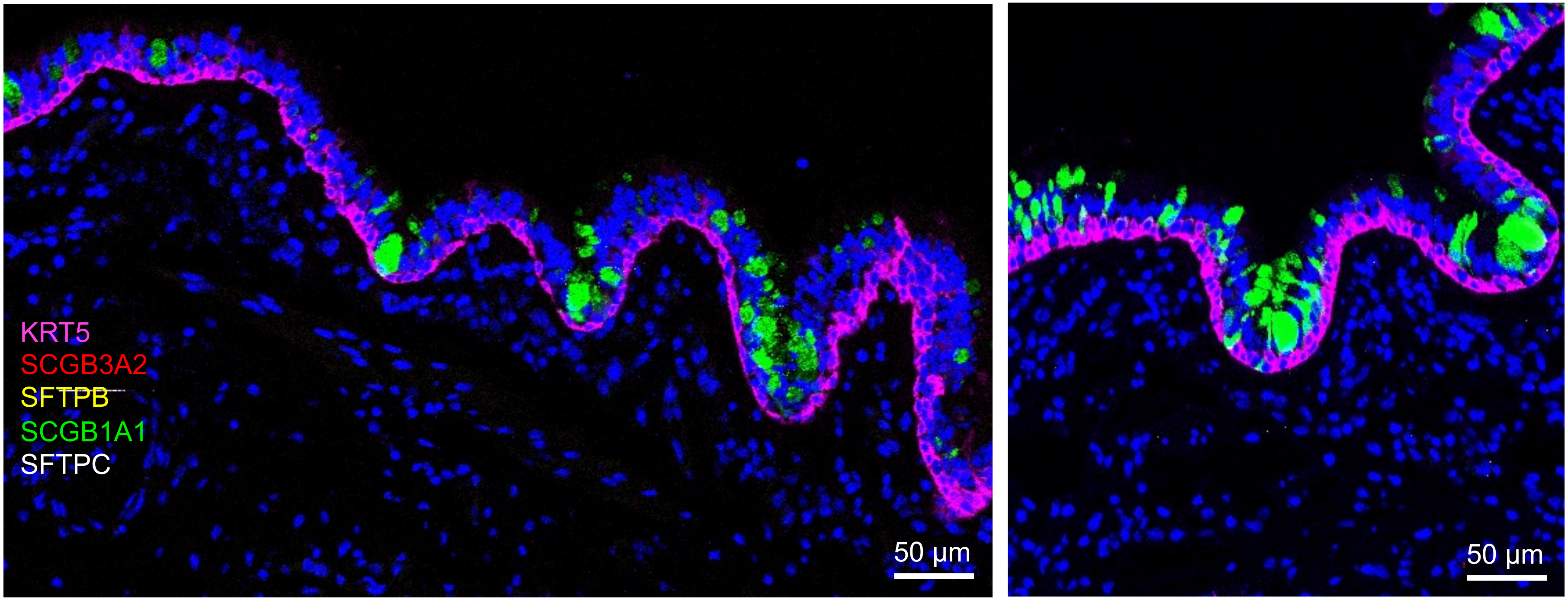

**B$_3$**

Distal

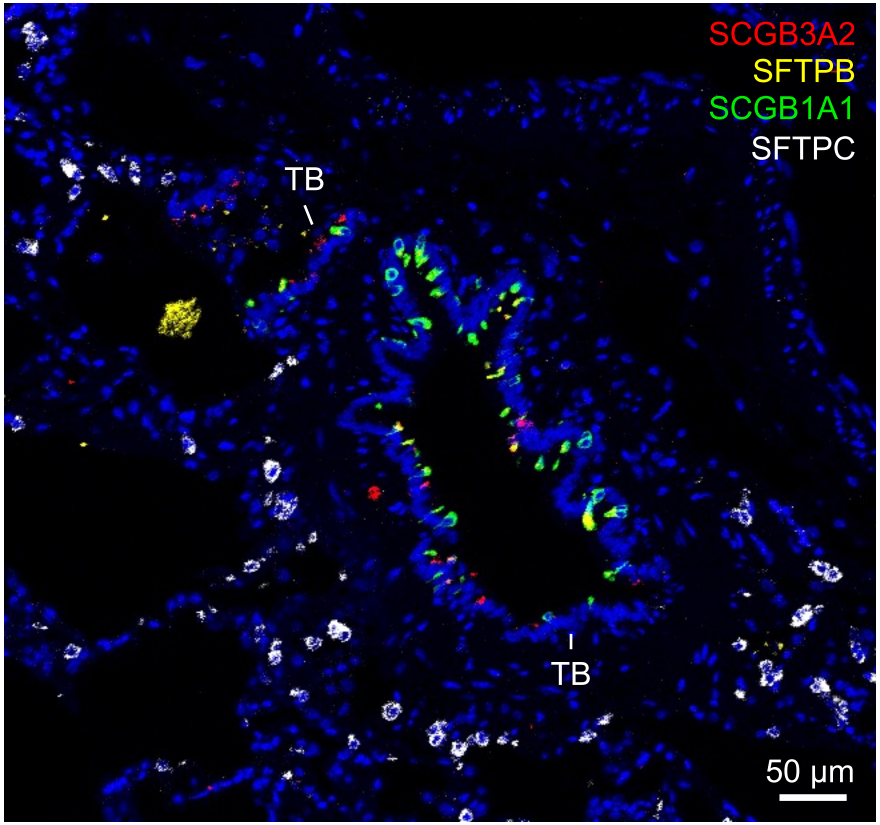
SCGB3A2
SFTPB
SCGB1A1
SFTPC
TB
TB
50 µm

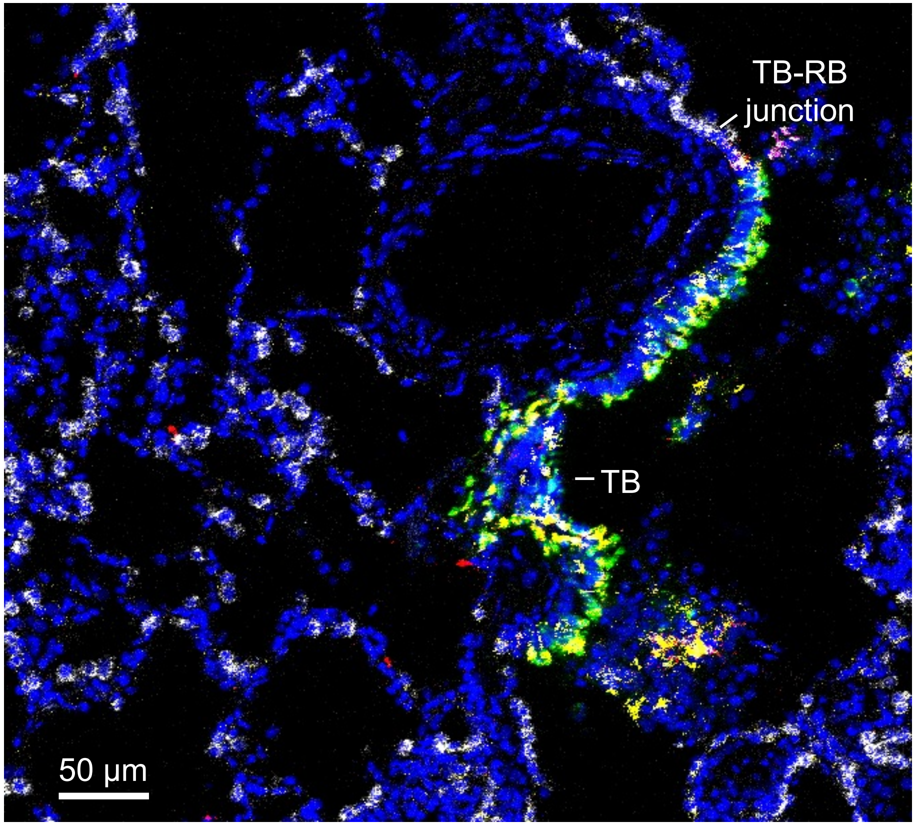
TB-RB
junction
TB
50 µm

**B$_4$**

Distal

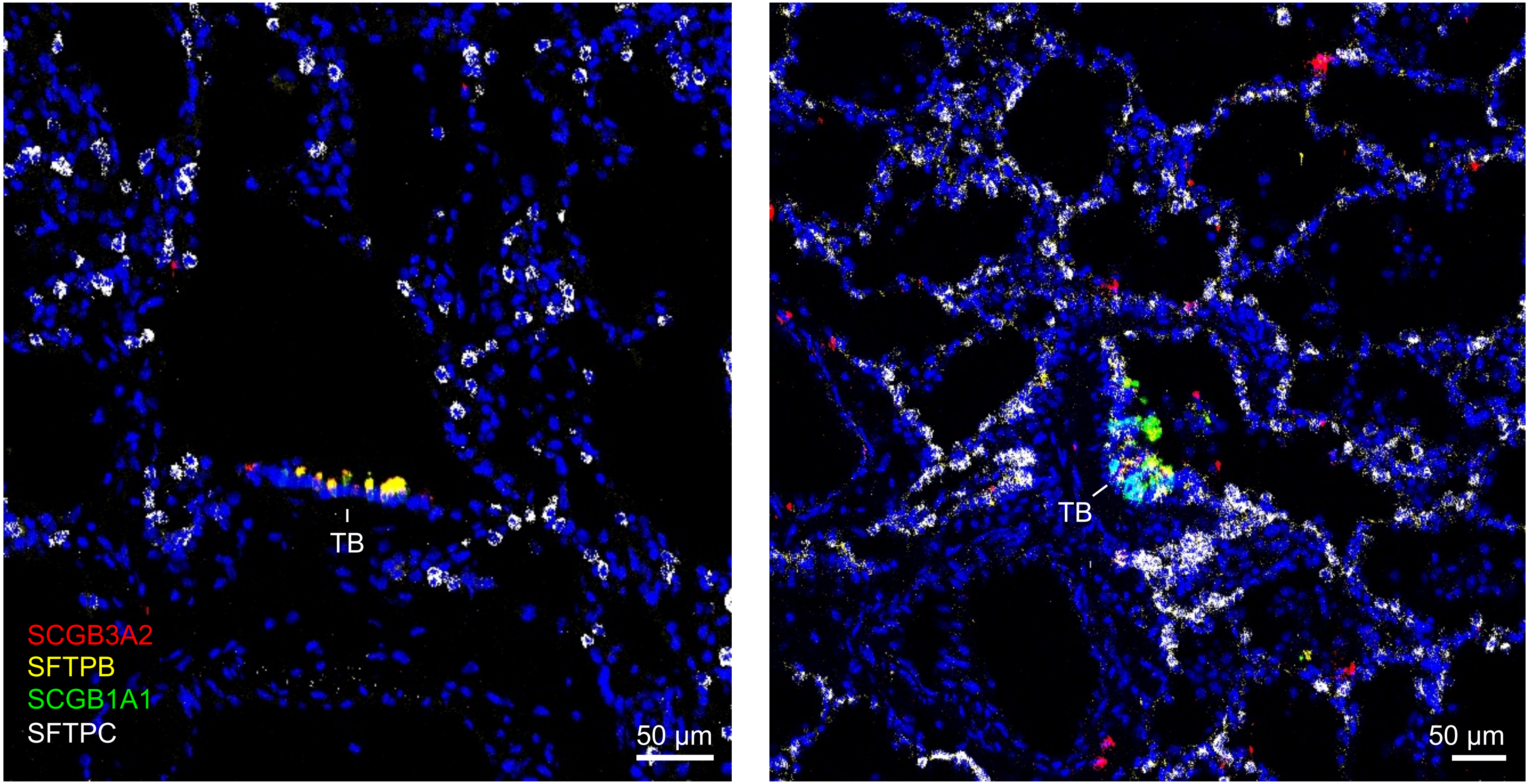

**B$_5$**

Distal

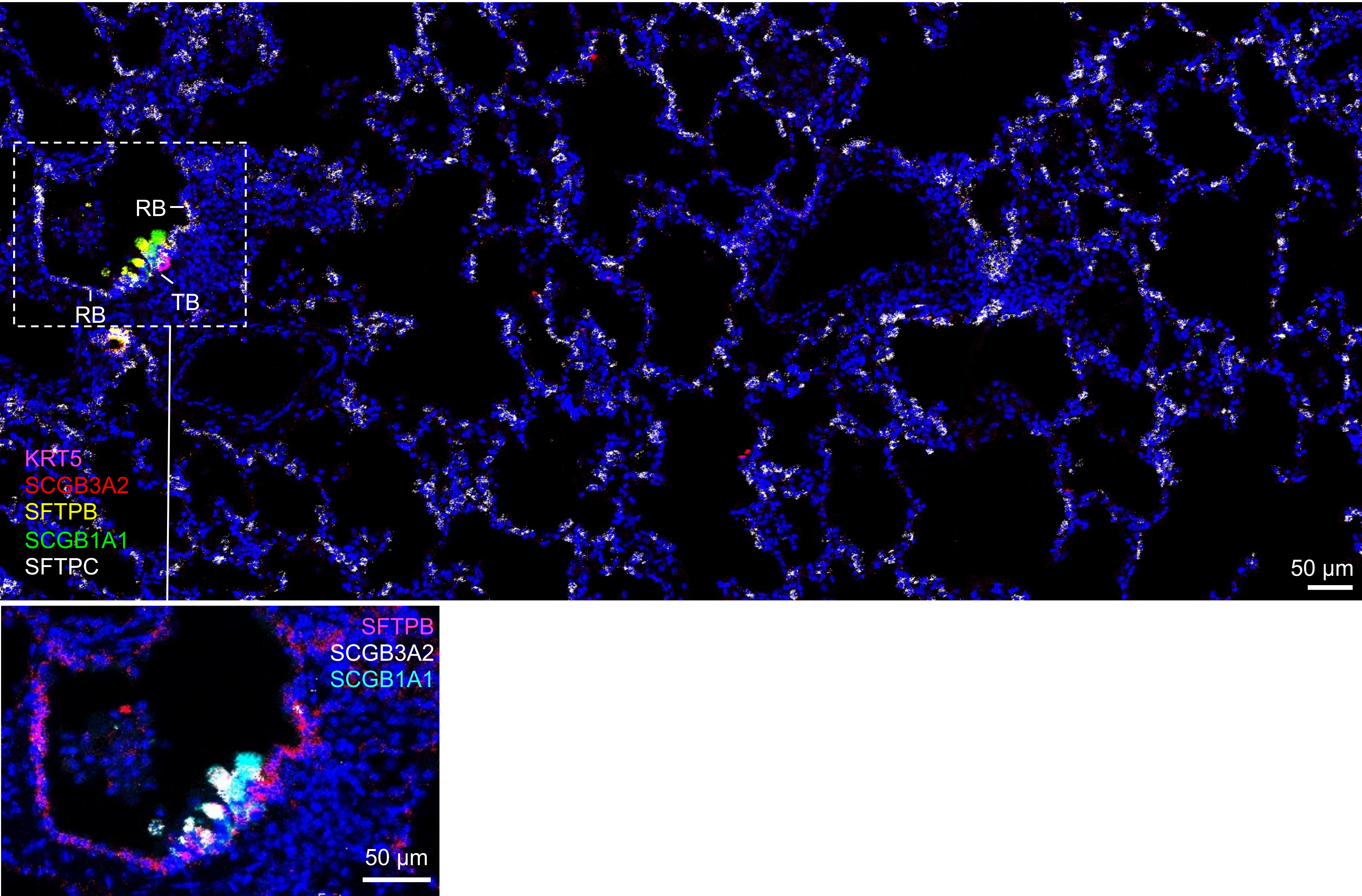
RB
RB
TB
KRT5
SCGB3A2
SFTPB
SCGB1A1
SFTPC
50 µm
SFTPB
SCGB3A2
SCGB1A1
50 µm

**$B_6$**

Distal

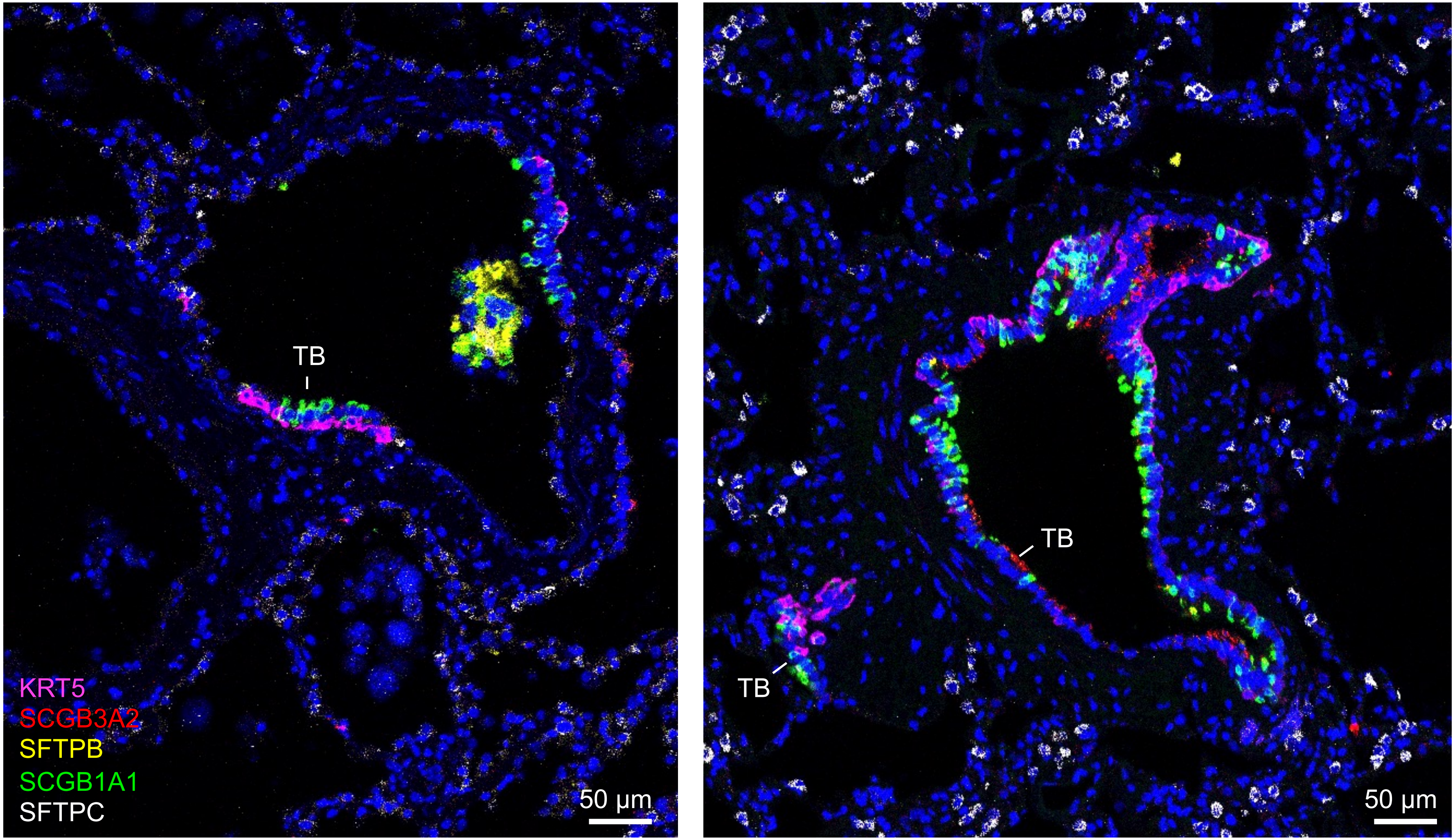

**$B_7$**

Distal

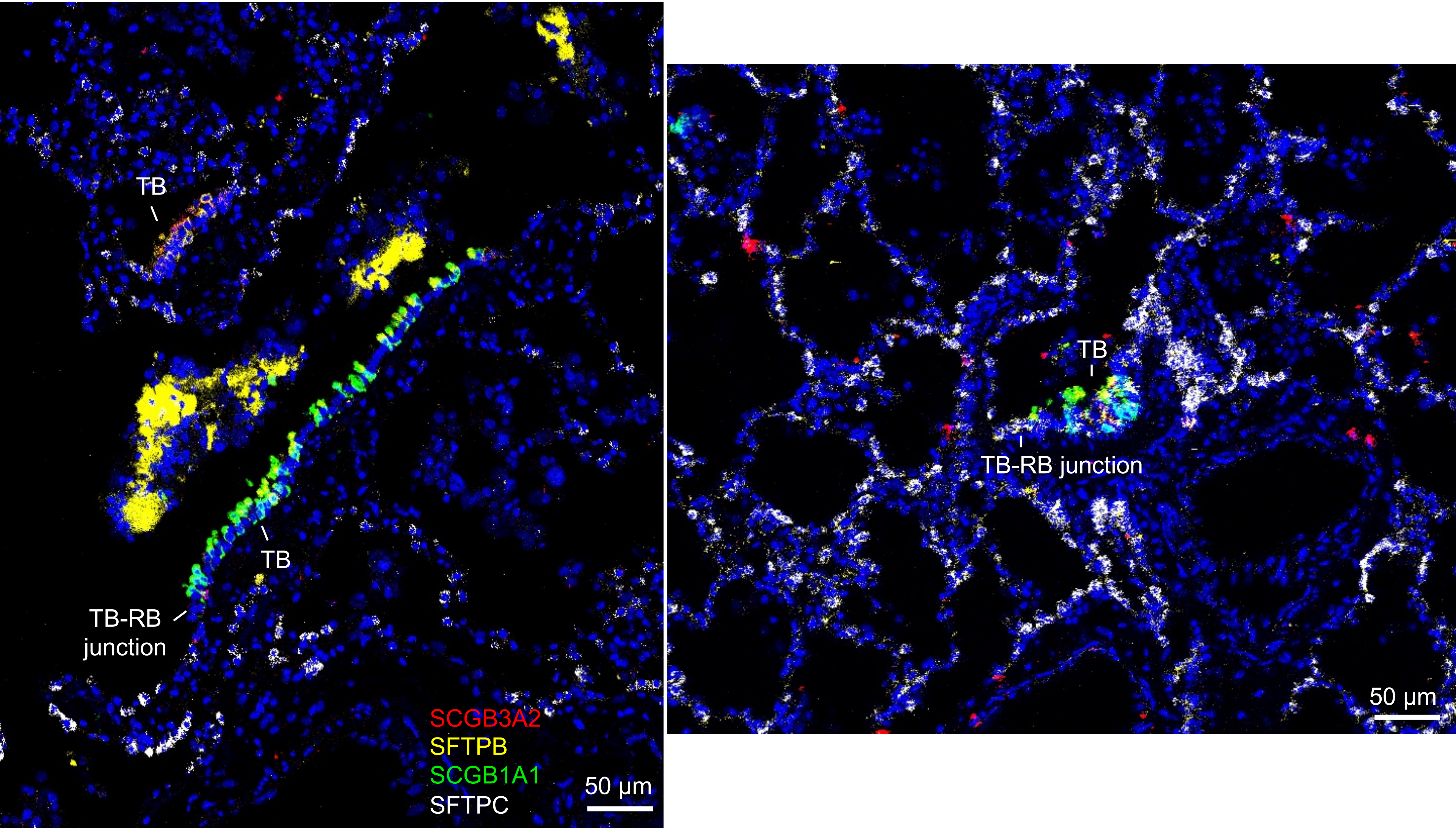
TB
TB
TB-RB junction
SCGB3A2
SFTPB
SCGB1A1
SFTPC
50 µm
TB
TB-RB junction
50 µm
