## Supplementary material for "A Unique Cellular Organization of Human Distal Airways and Its Disarray in Chronic Obstructive Pulmonary Disease": Fig. S4

**A1**

Proximal airway

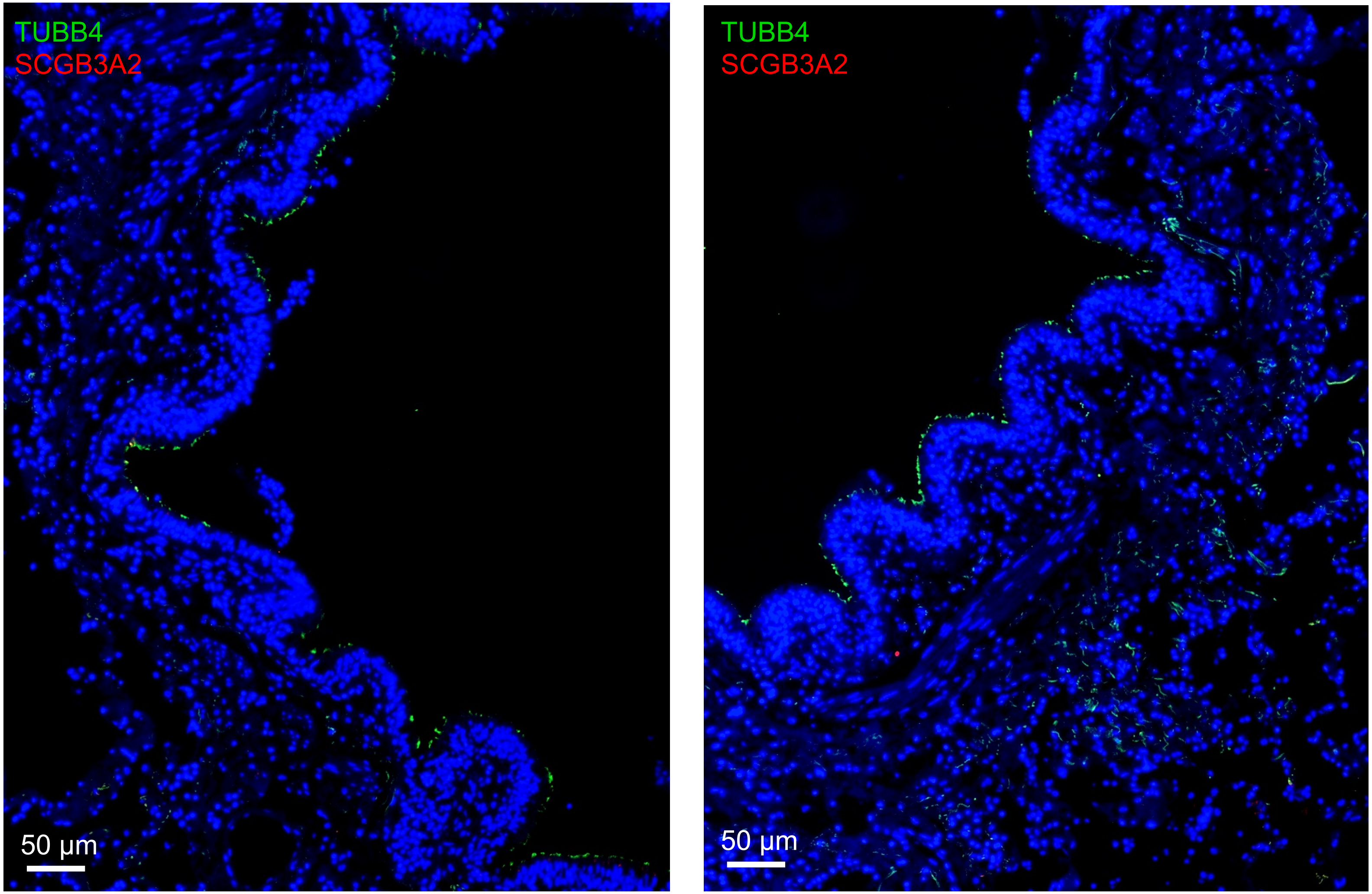

**A₂**

Proximal airway

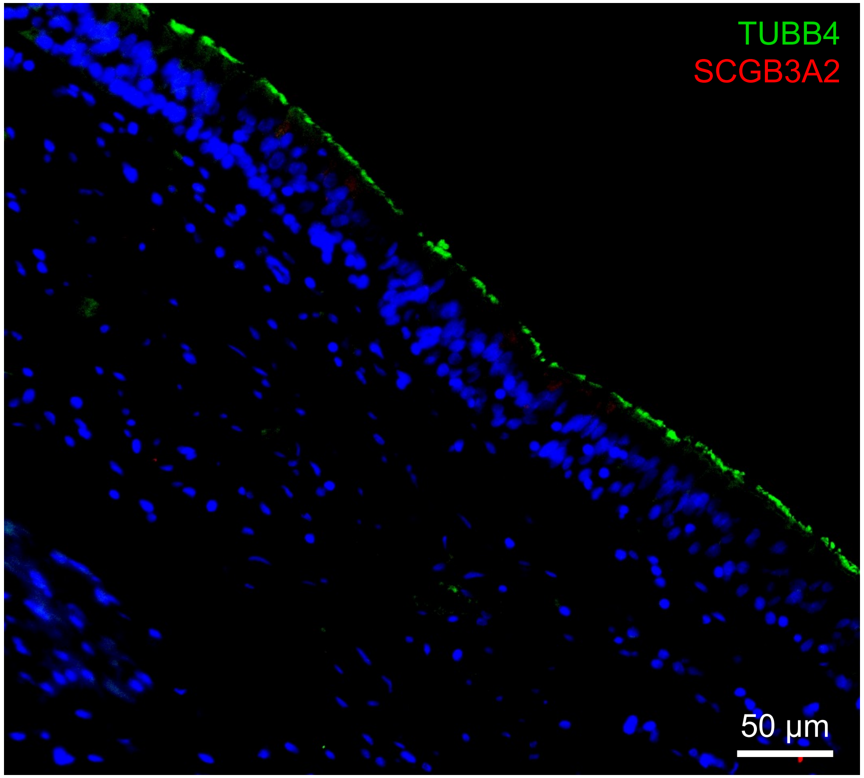


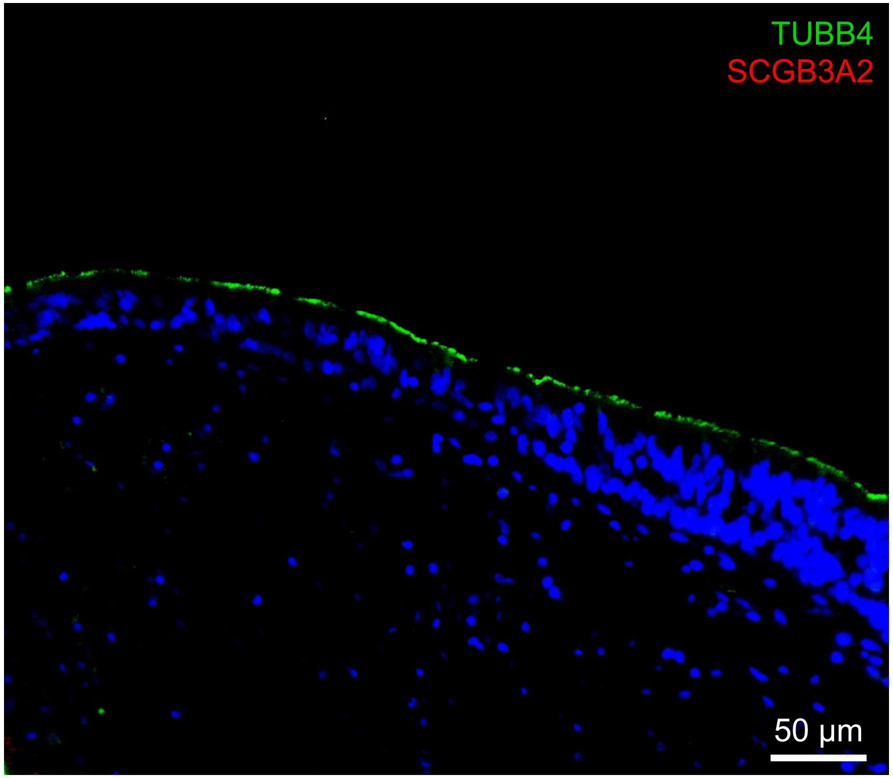

$A_3$

Proximal airway

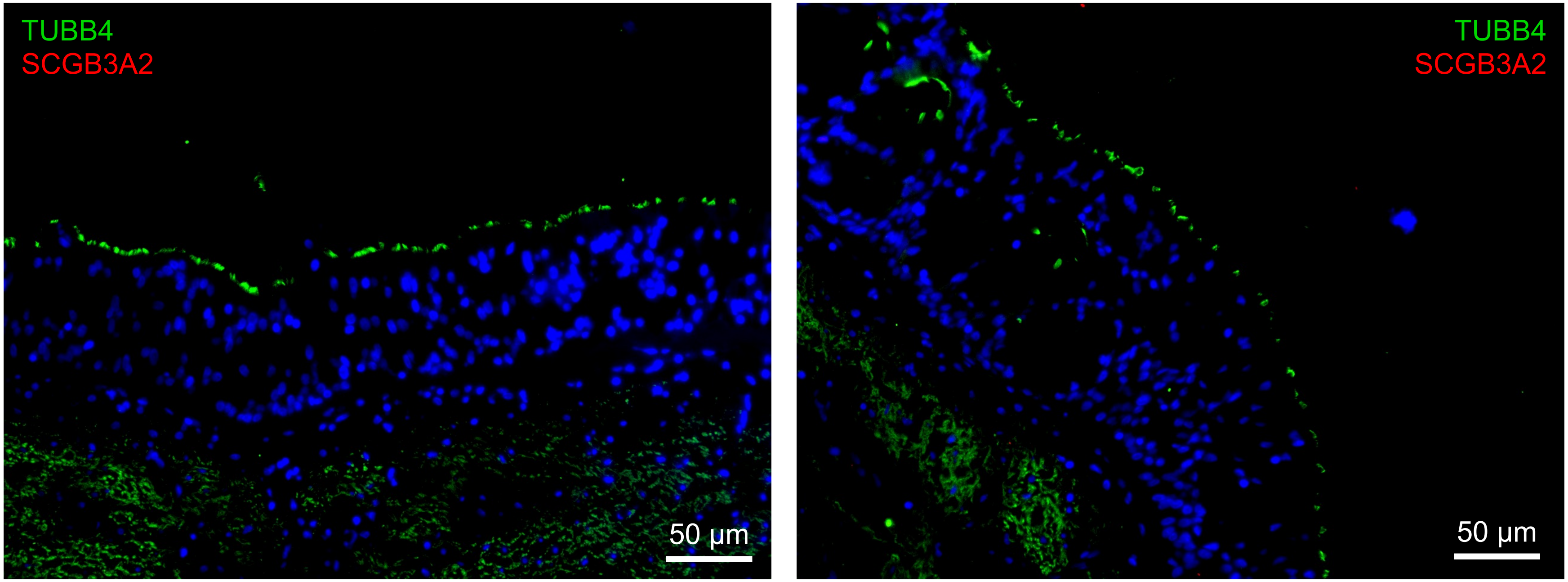

**A₄**

Proximal airway

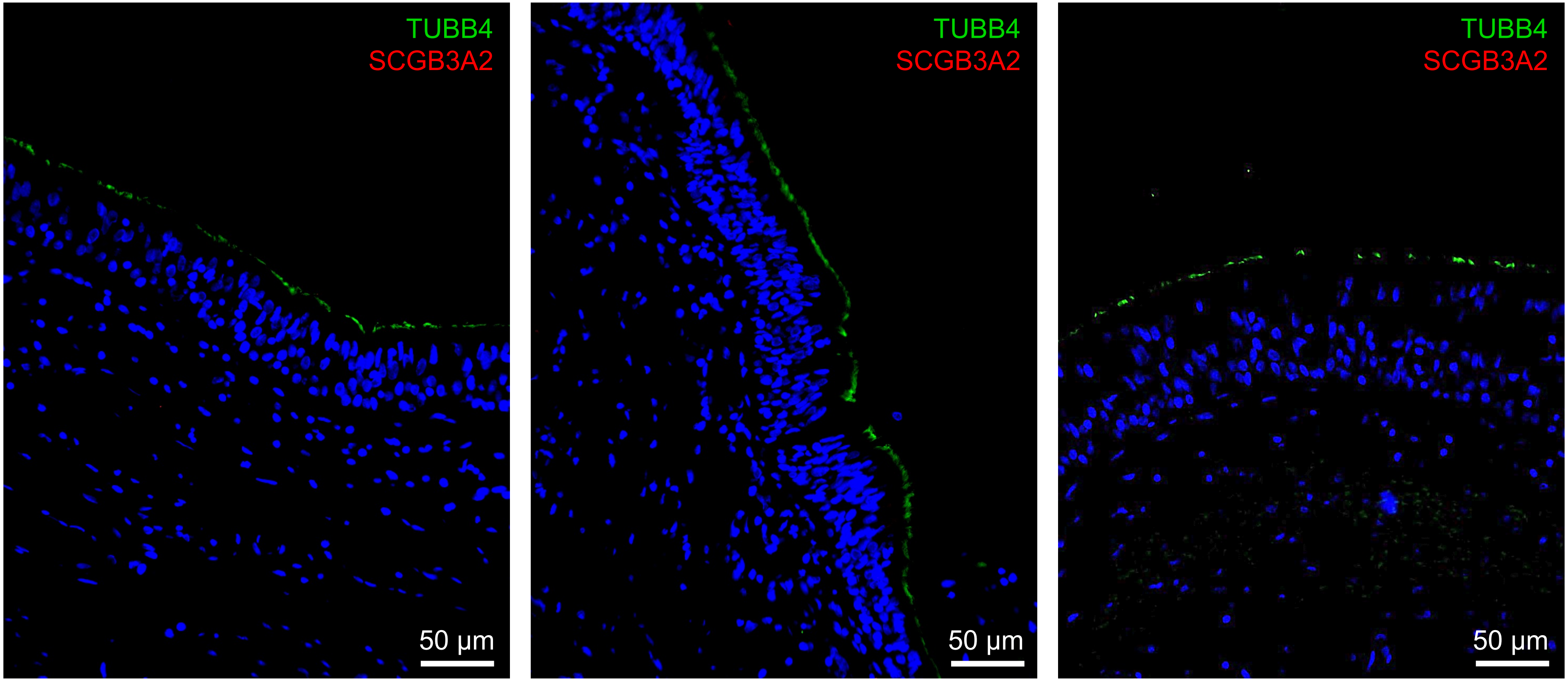

**A5**

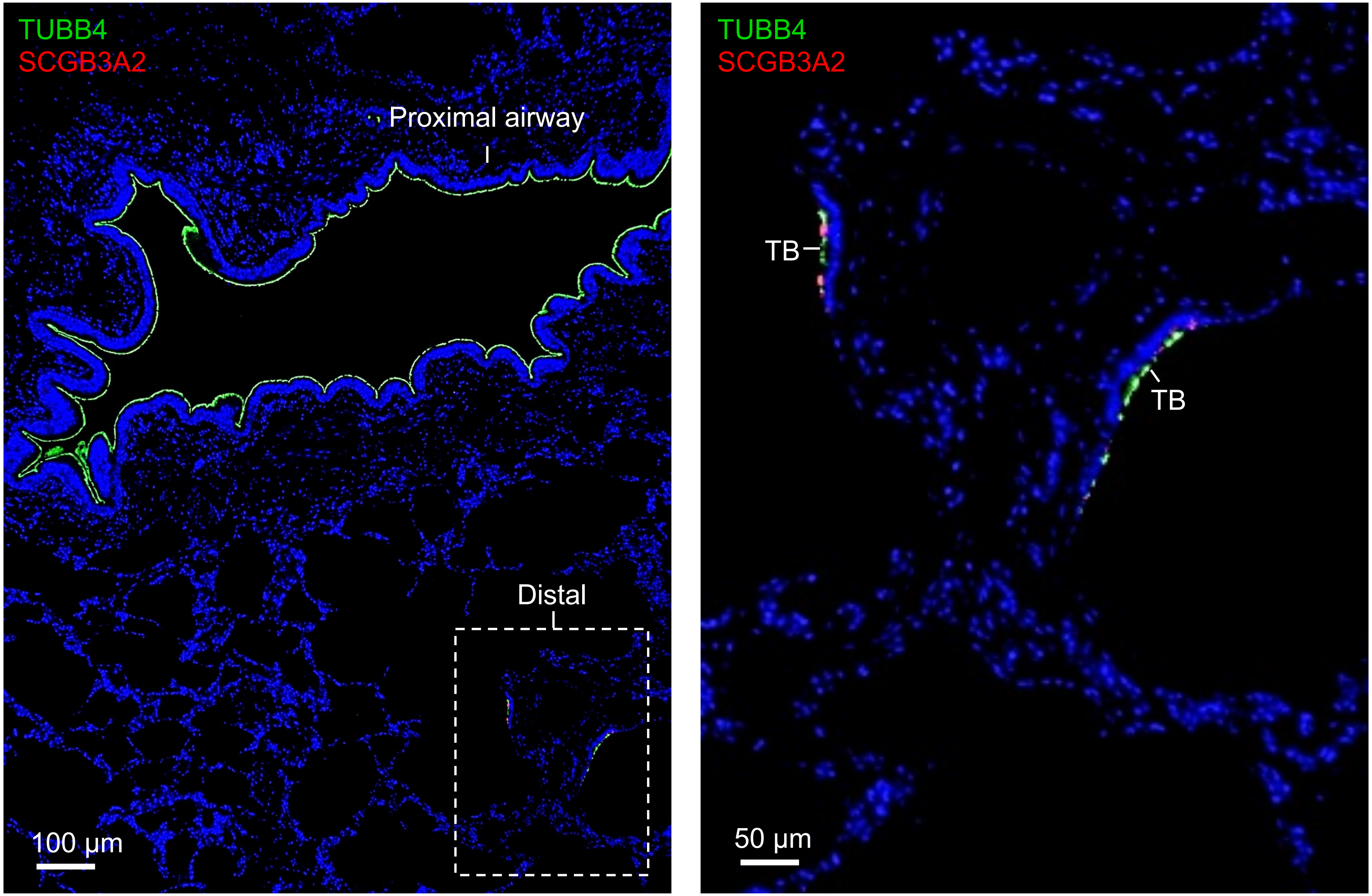
TUBB4
SCGB3A2
Proximal airway
Distal
100 µm
TUBB4
SCGB3A2
TB
TB
50 µm

**A6**

Distal airways

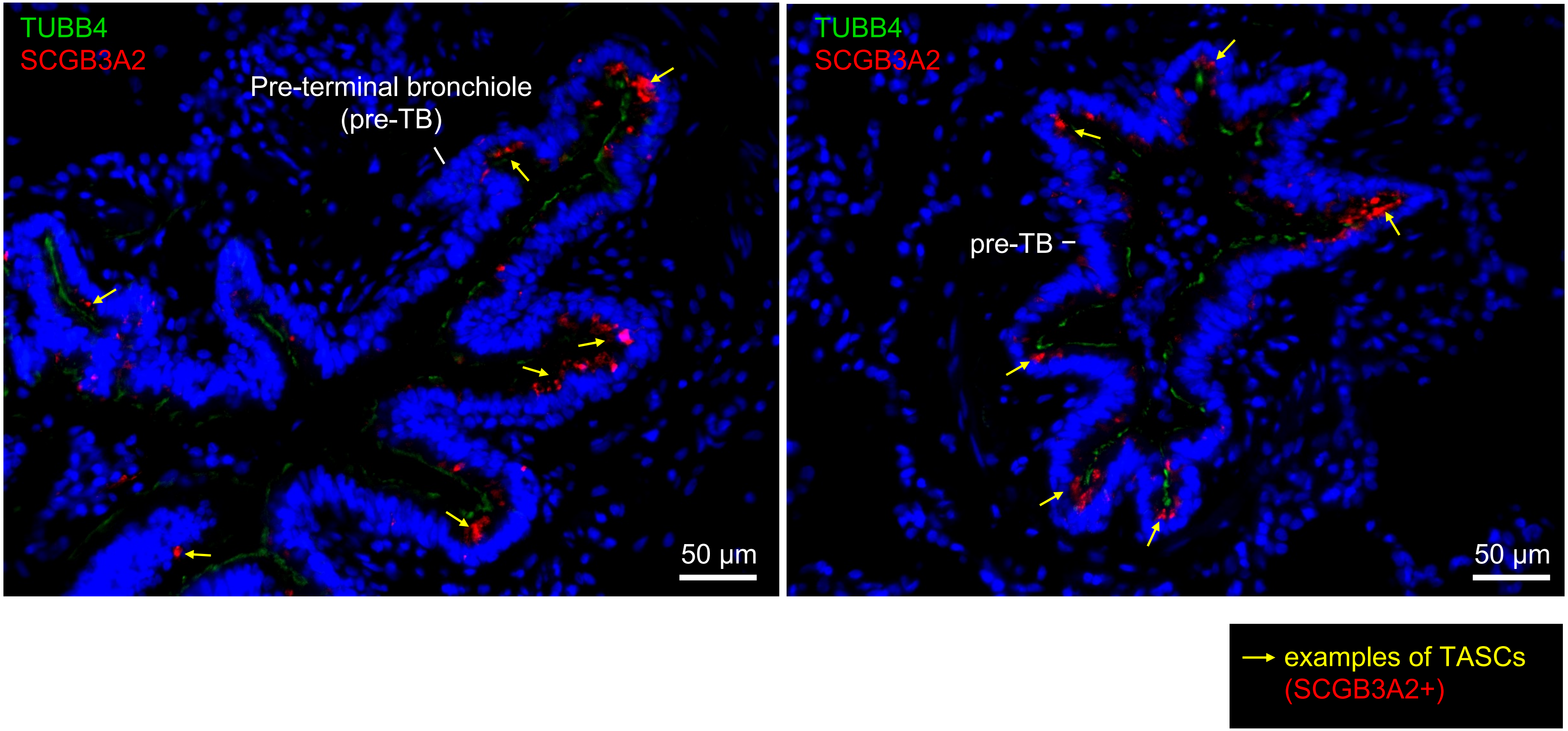

**A7**

Distal airways

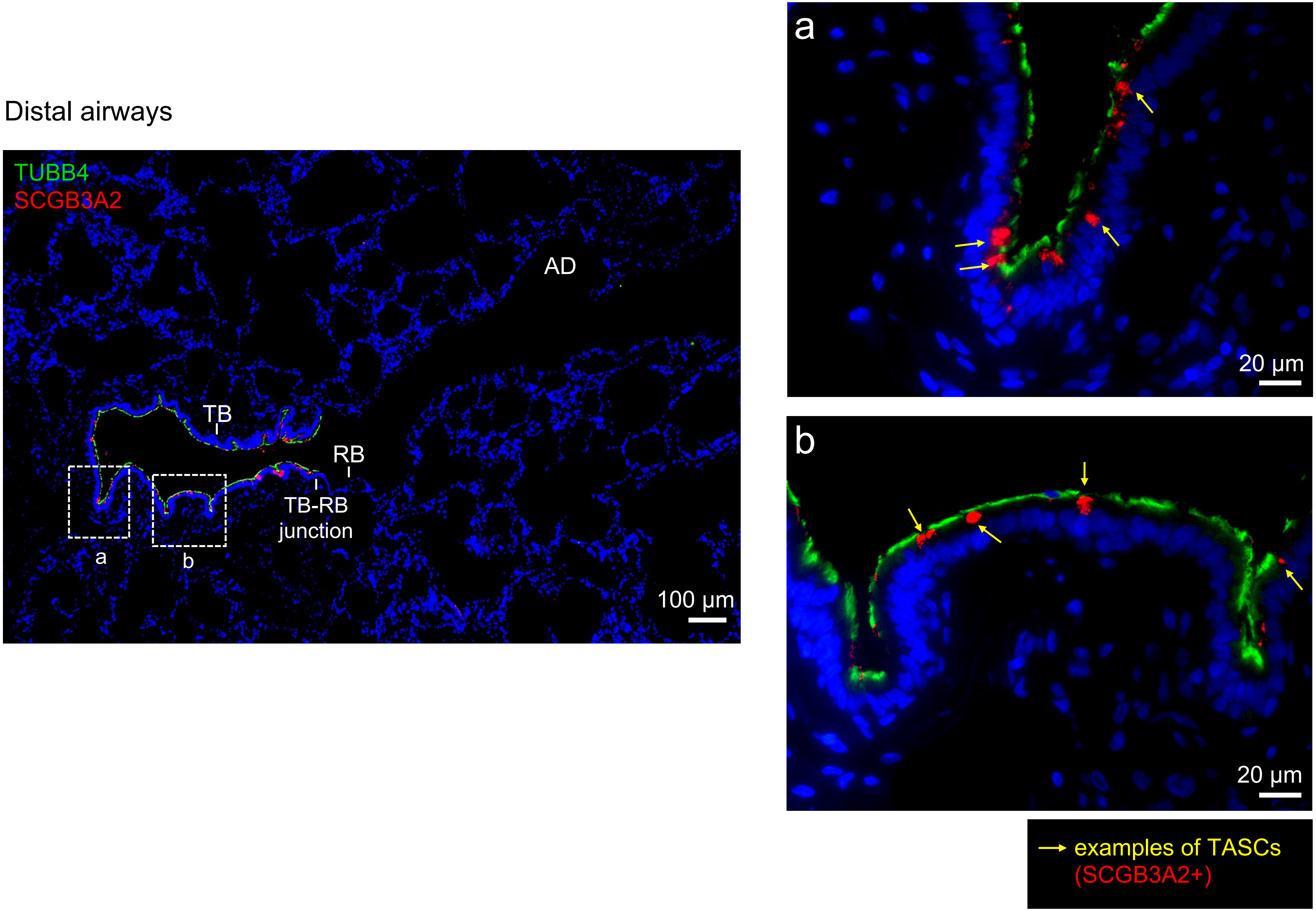

**A8**

Distal airways

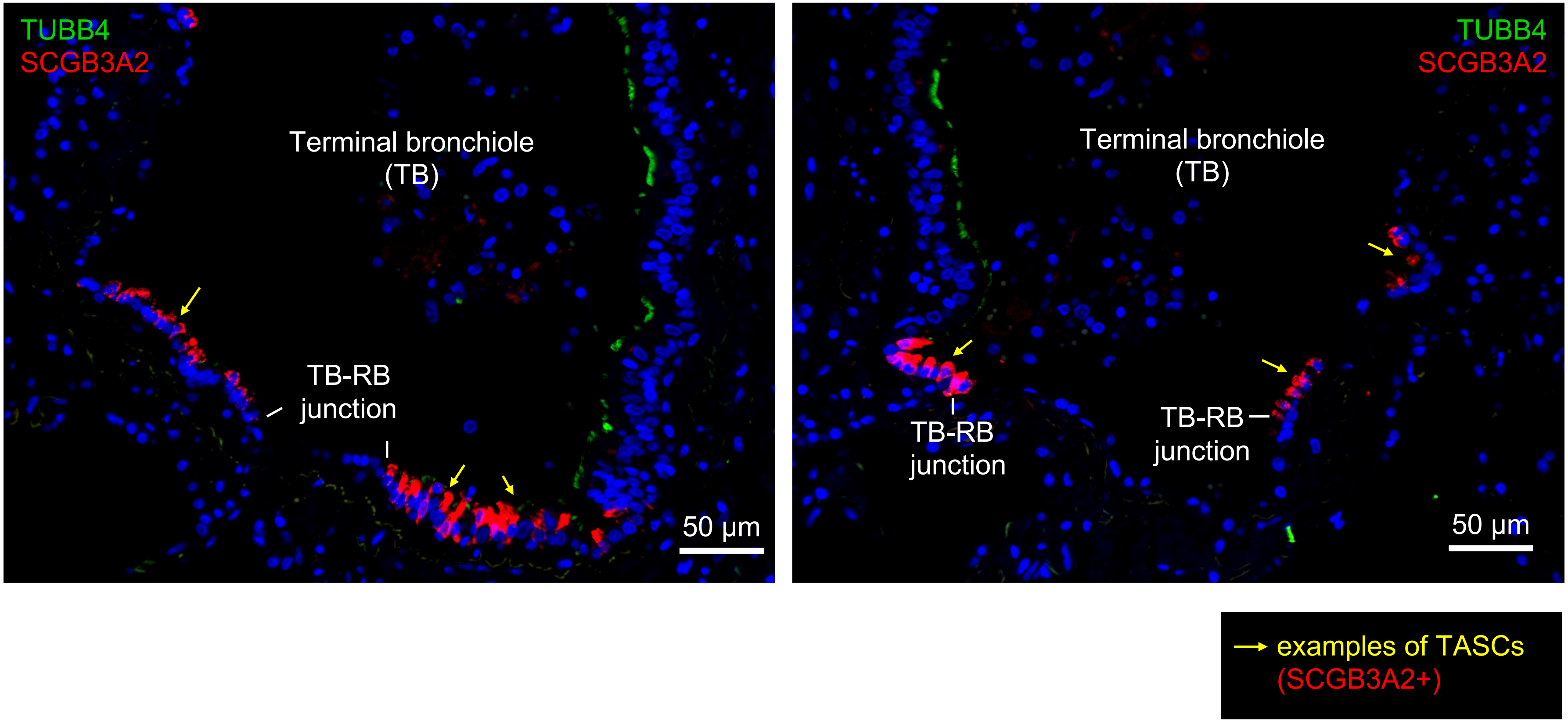

**A9**

Distal airways

Terminal bronchiole (TB)

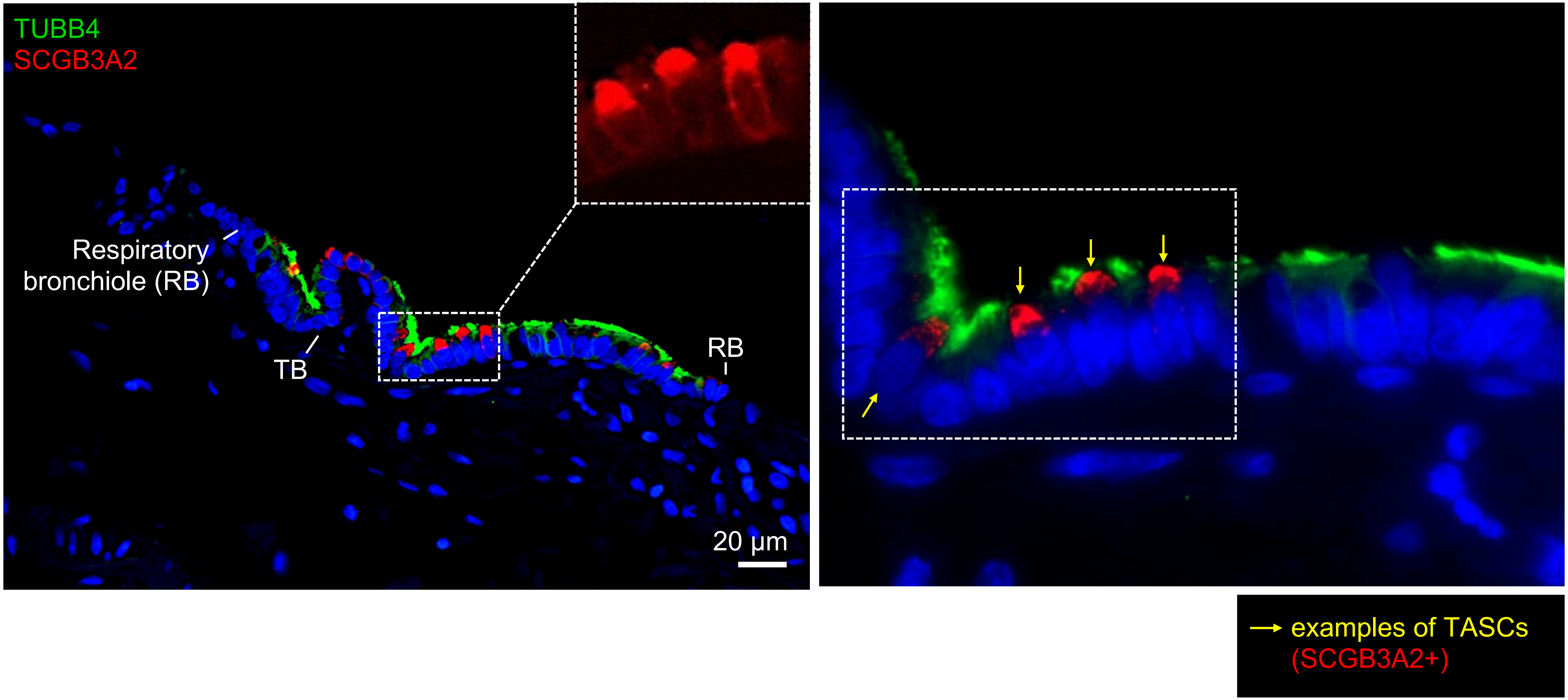

**A10**

Distal airways

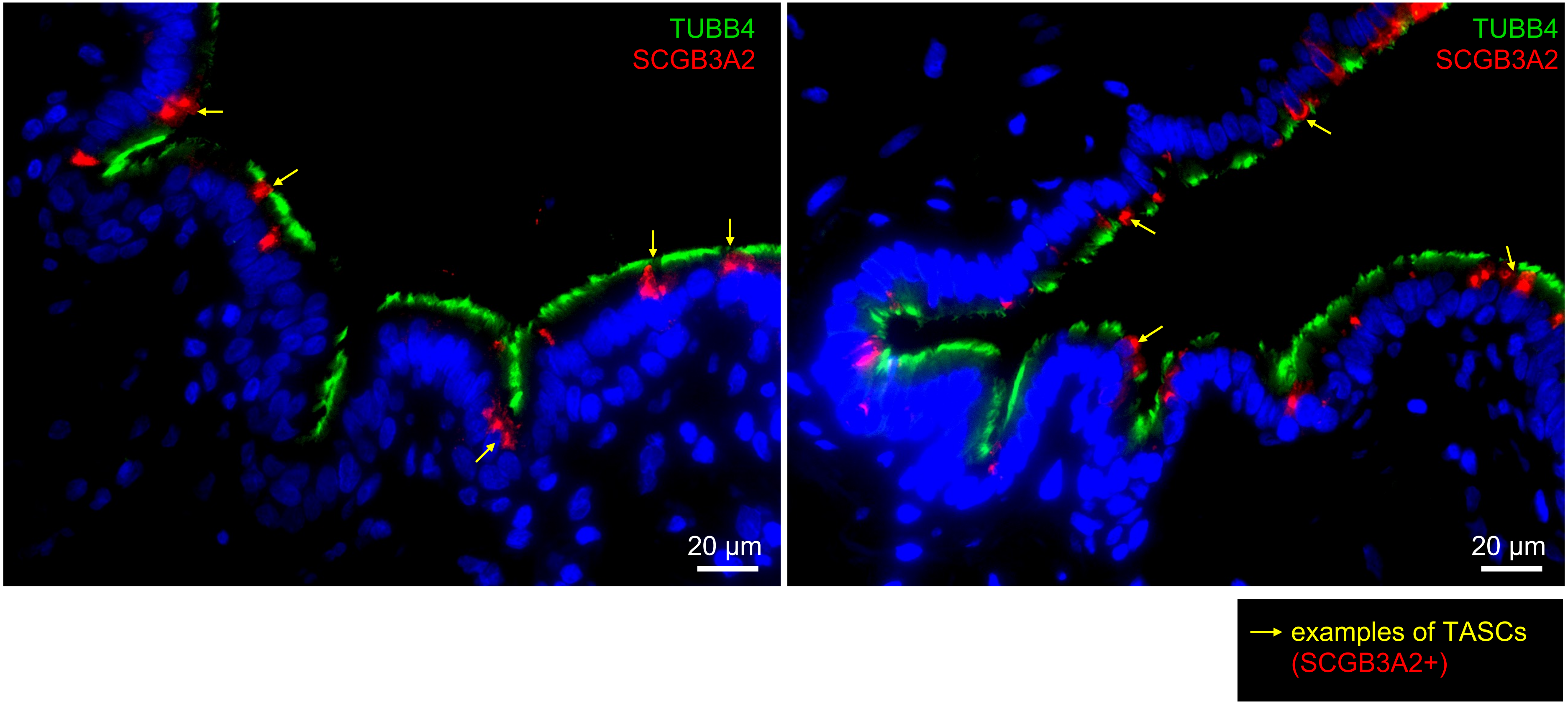

**A**11

Distal airways

Terminal bronchioles (TBs)

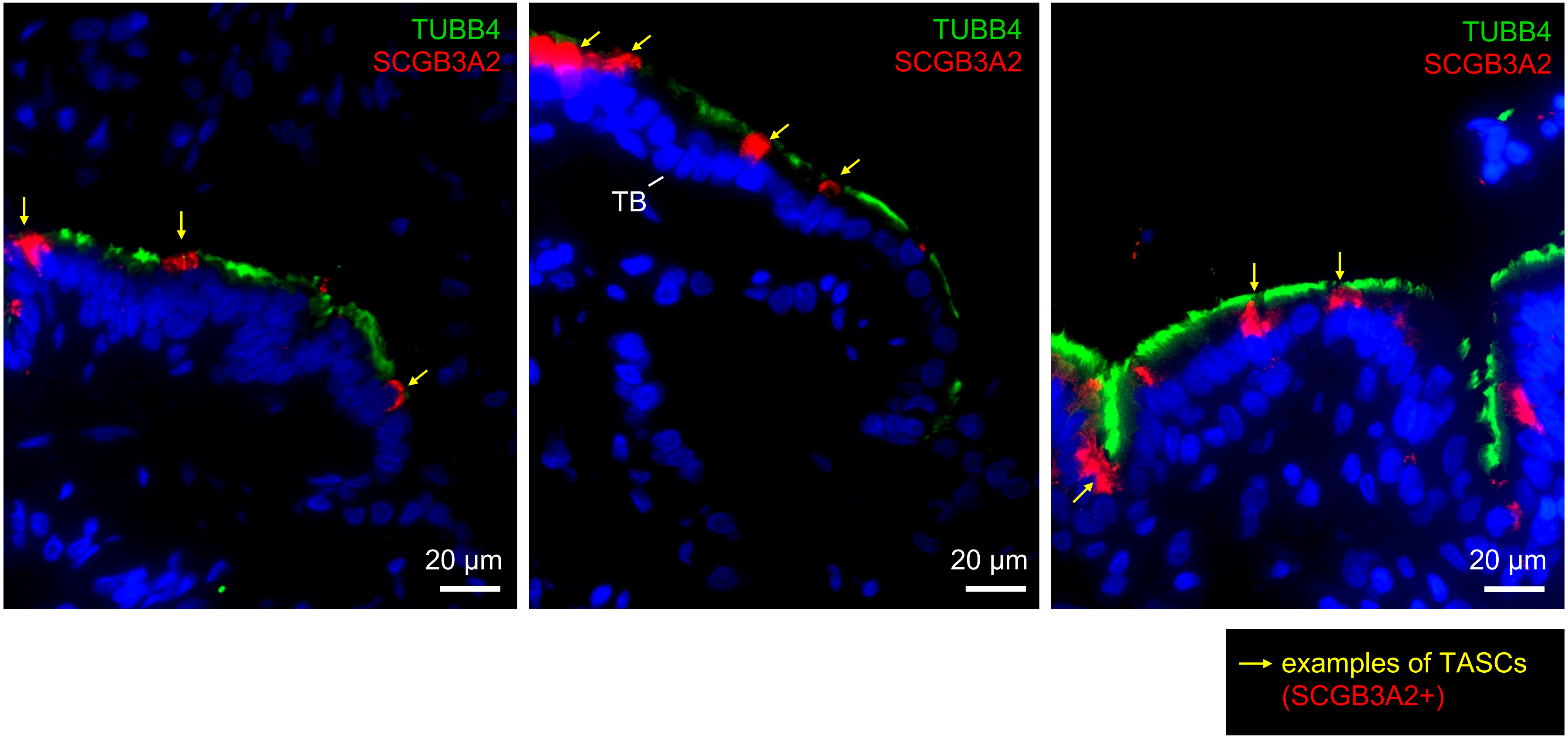

**Supplemental Figure 4**

# A12

Distal airways

Terminal bronchiole (TB)

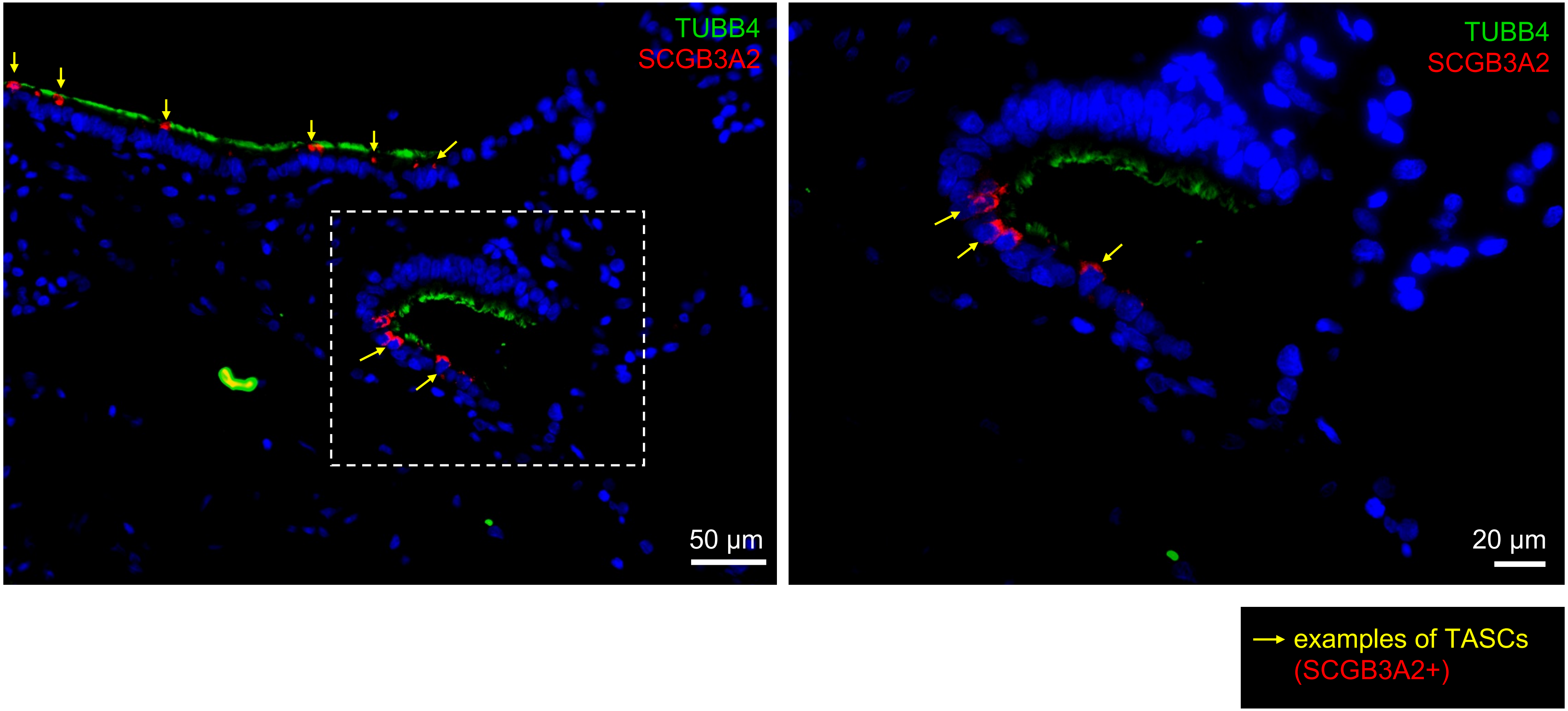

# A13

Distal airways

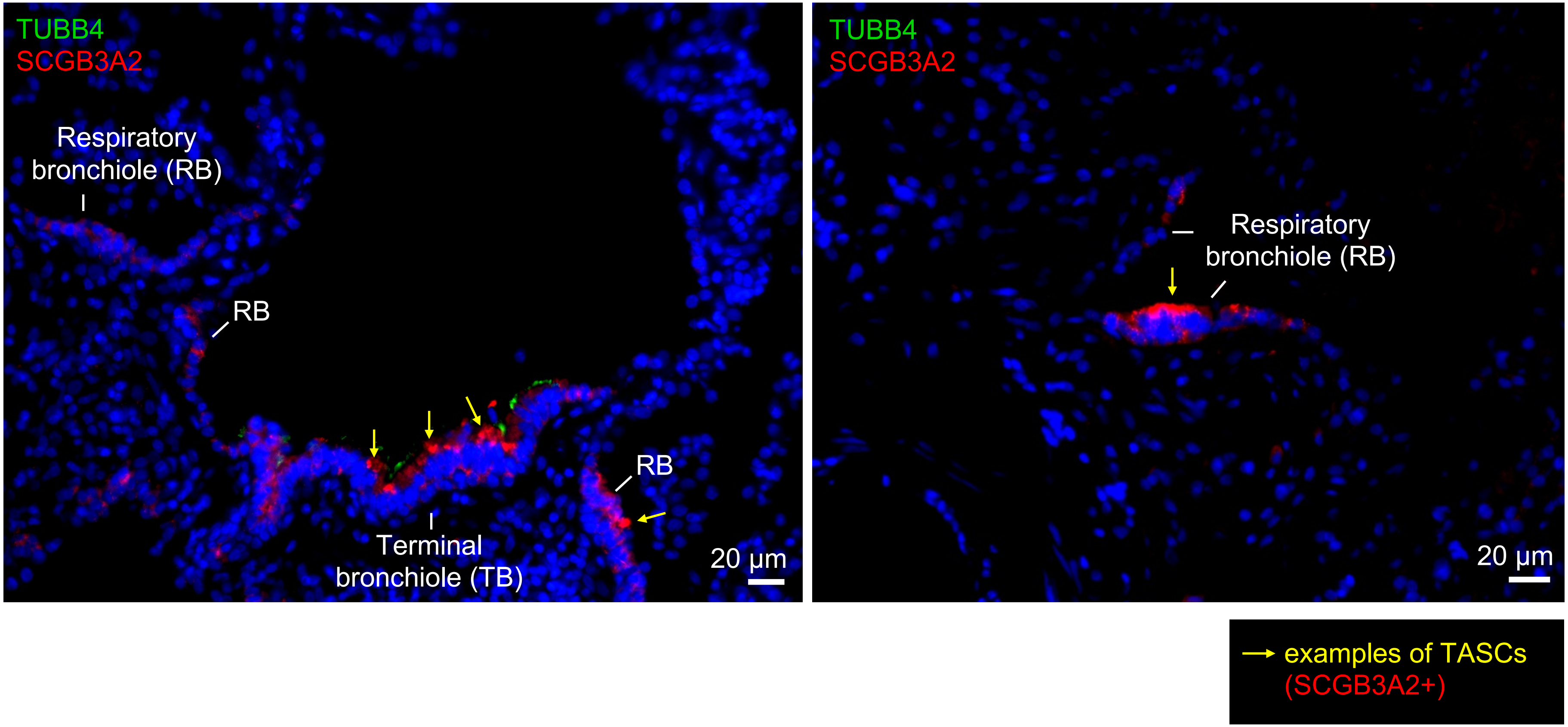

**$B_1$**

Proximal airways

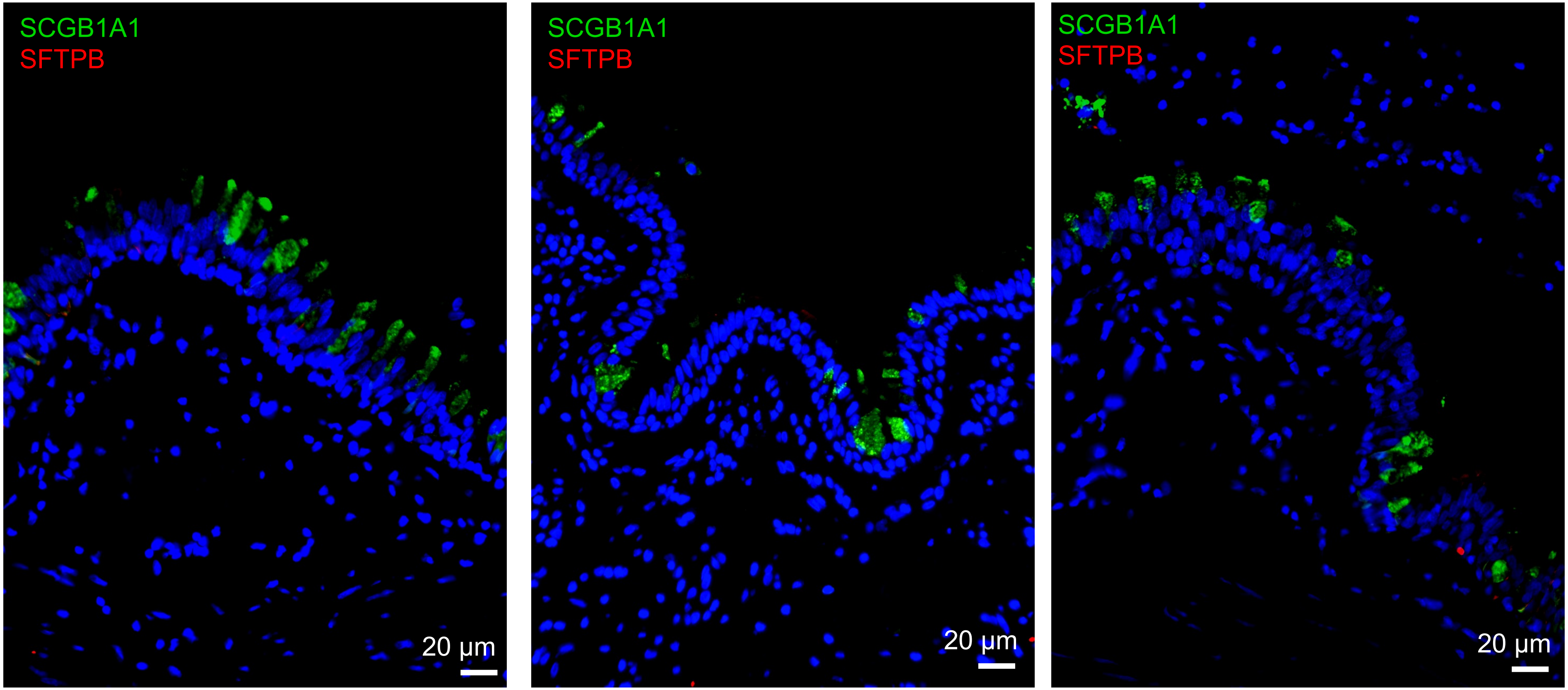

**Supplemental Figure 4**

**$B_2$**

Proximal airways

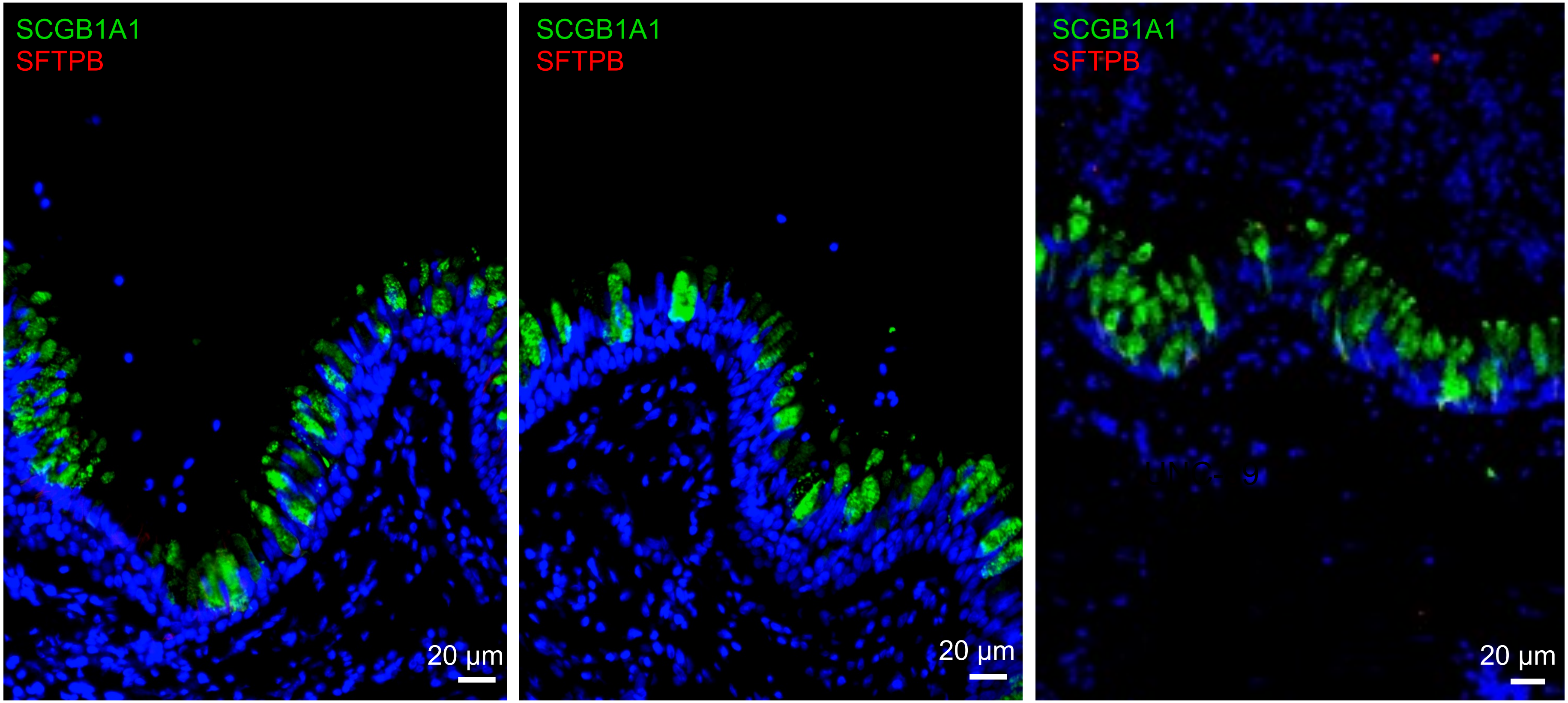

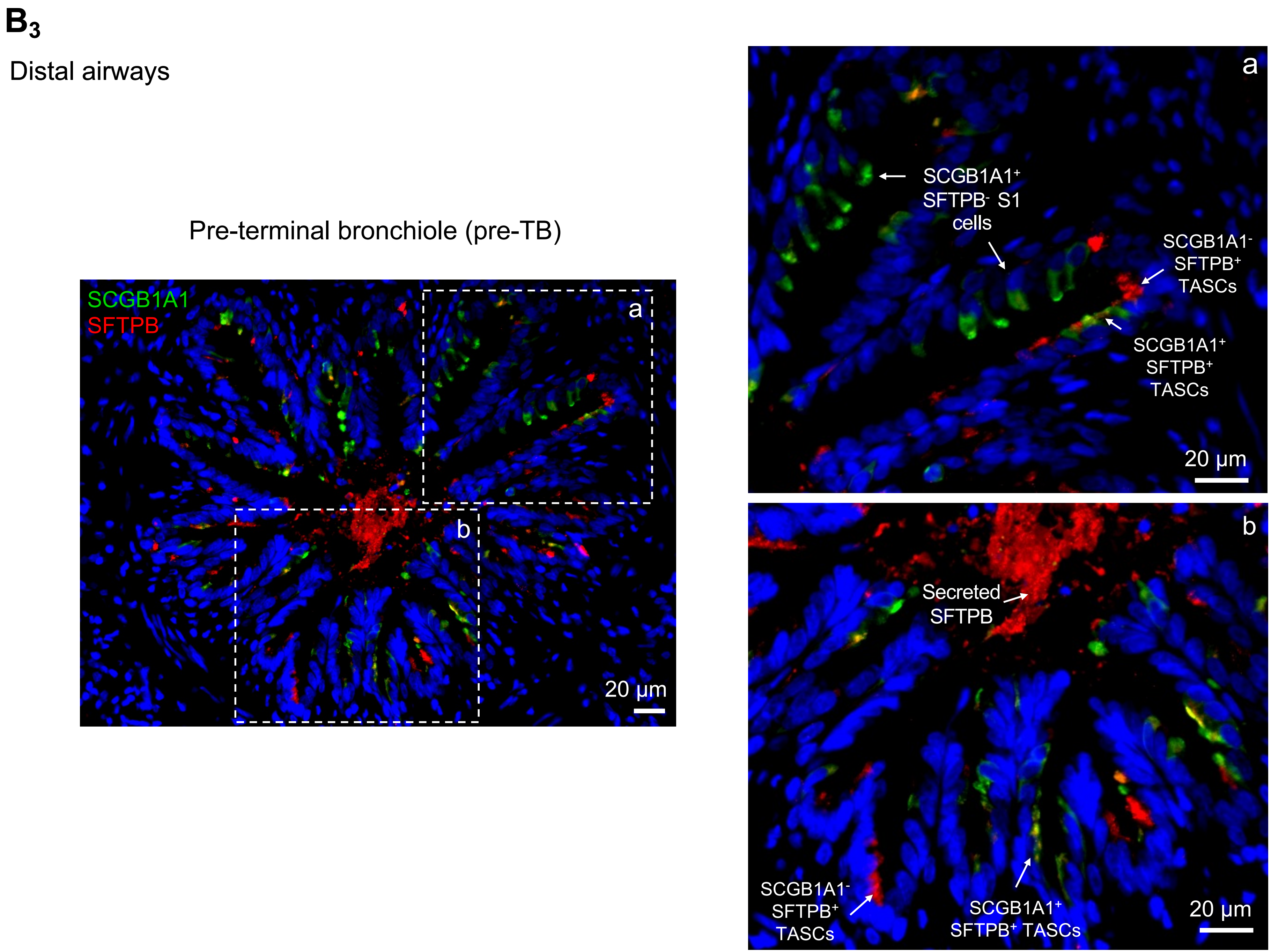
B3
Distal airways
Pre-terminal bronchiole (pre-TB)
SCGB1A1
SFTPB
a
b
20 µm
SCGB1A1+
SFTPB- S1
cells
SCGB1A1-
SFTPB+
TASCs
SCGB1A1+
SFTPB+
TASCs
Secreted
SFTPB
SCGB1A1-
SFTPB+
TASCs
SCGB1A1+
SFTPB+ TASCs

**Supplemental Figure 4**

## $B_4$

Distal airways

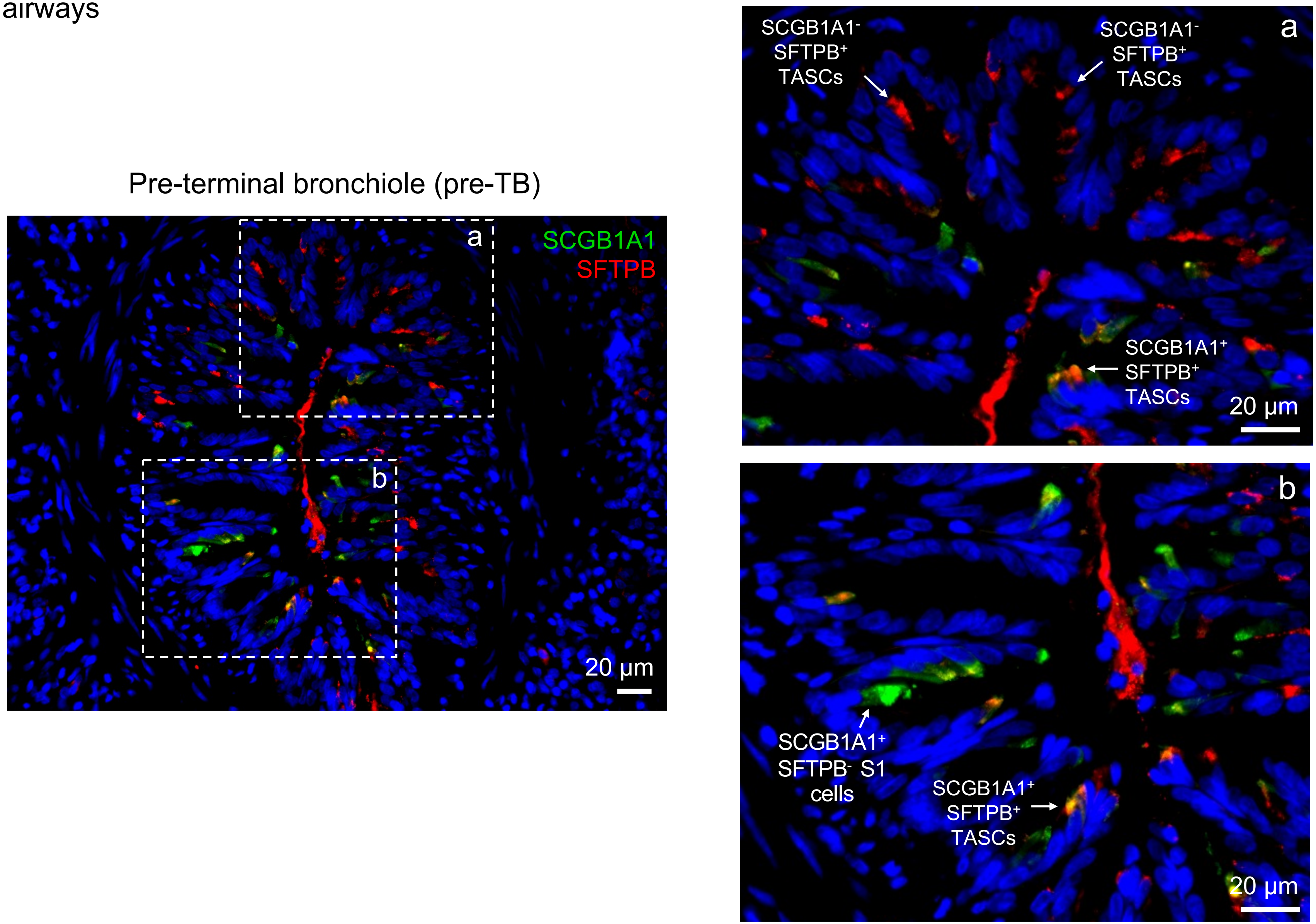

# $B_5$

Distal airways

Terminal bronchioles (TB) and respiratory bronchioles (RB)

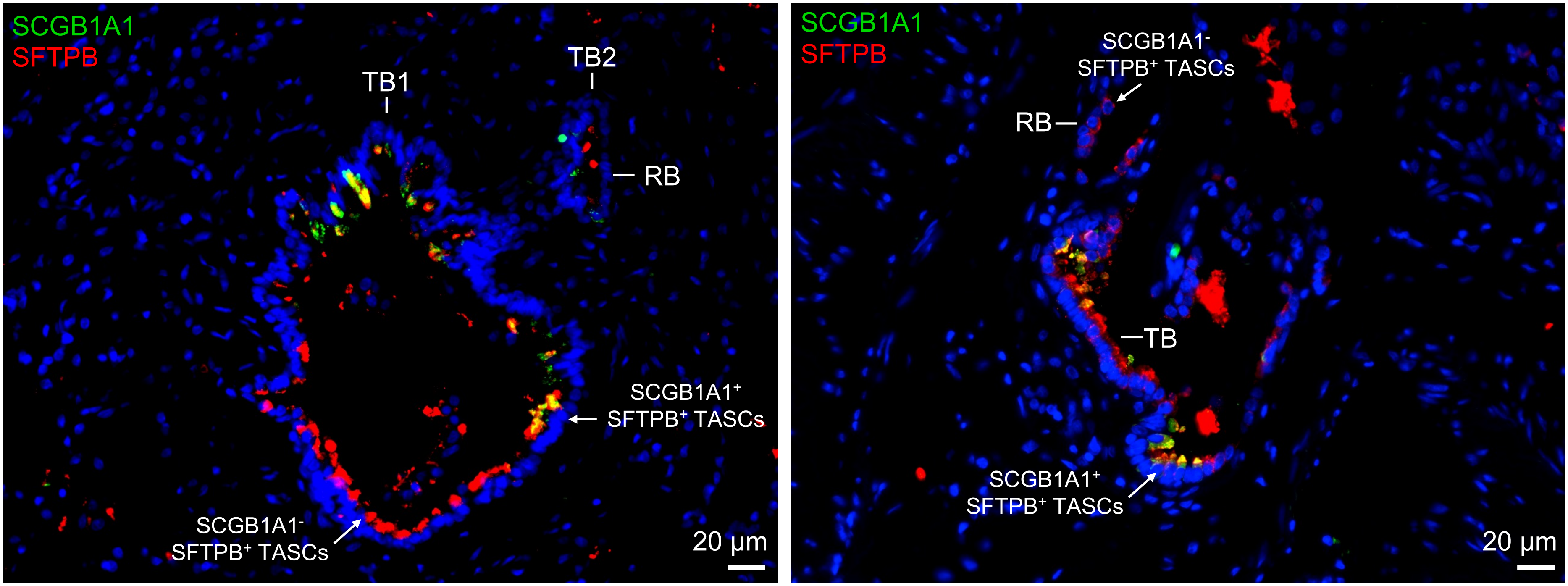

# $B_6$

Distal airways

Terminal bronchioles (TBs)

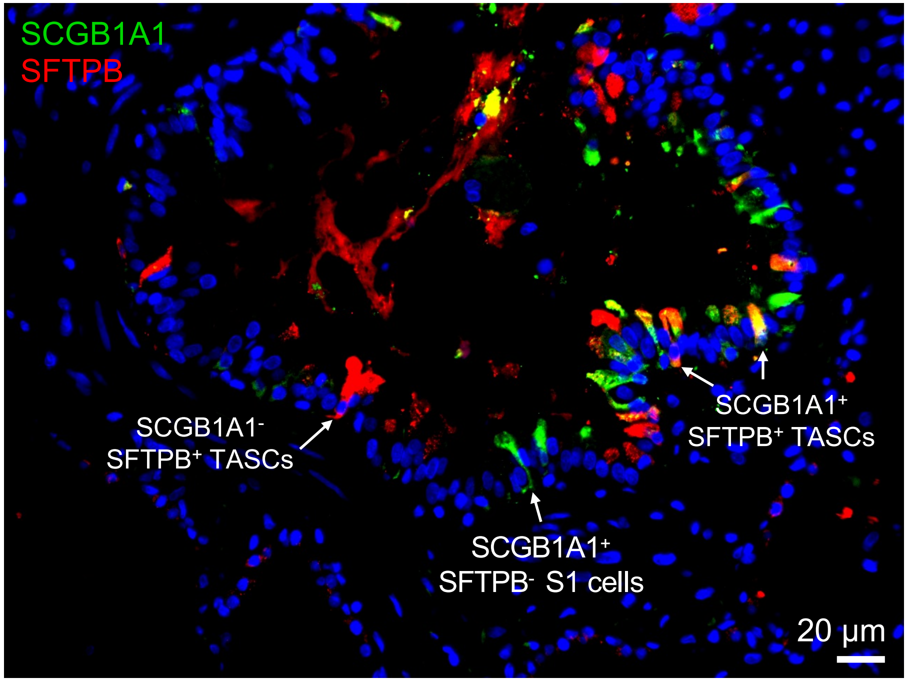


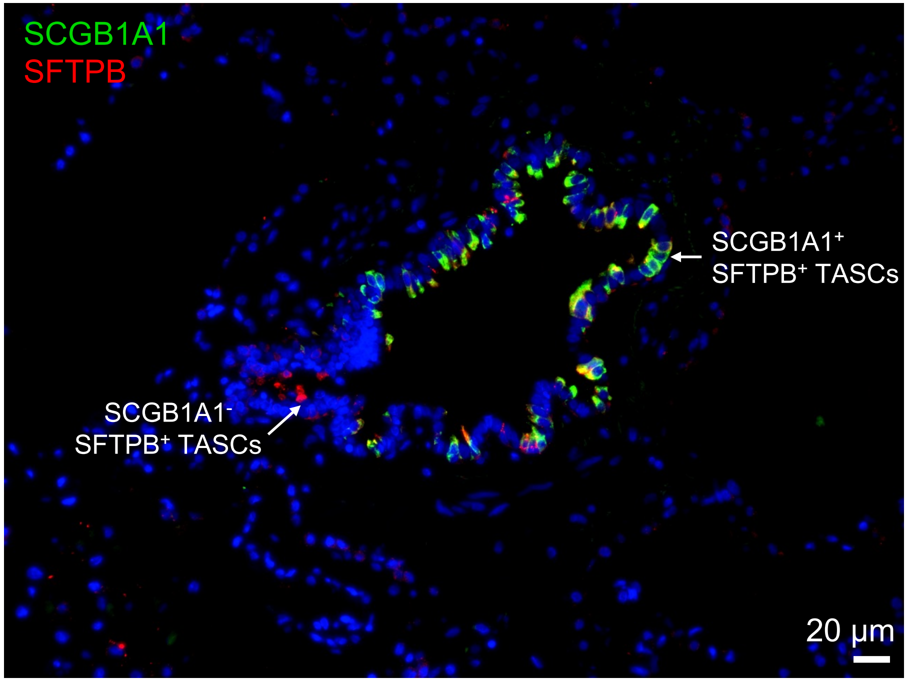

# $B_7$

Distal airways

Terminal bronchioles (TBs)

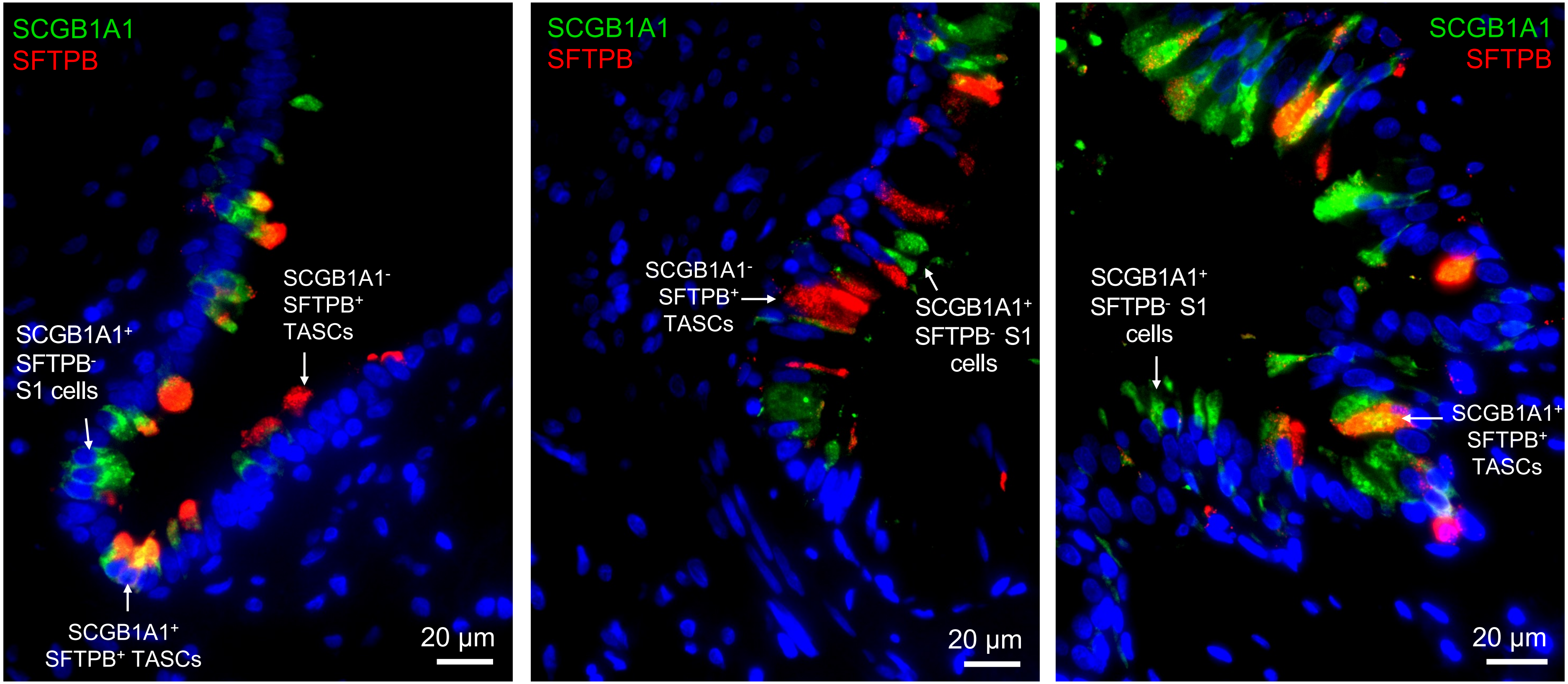

**B₈** Distal airways

Terminal bronchioles (TBs)

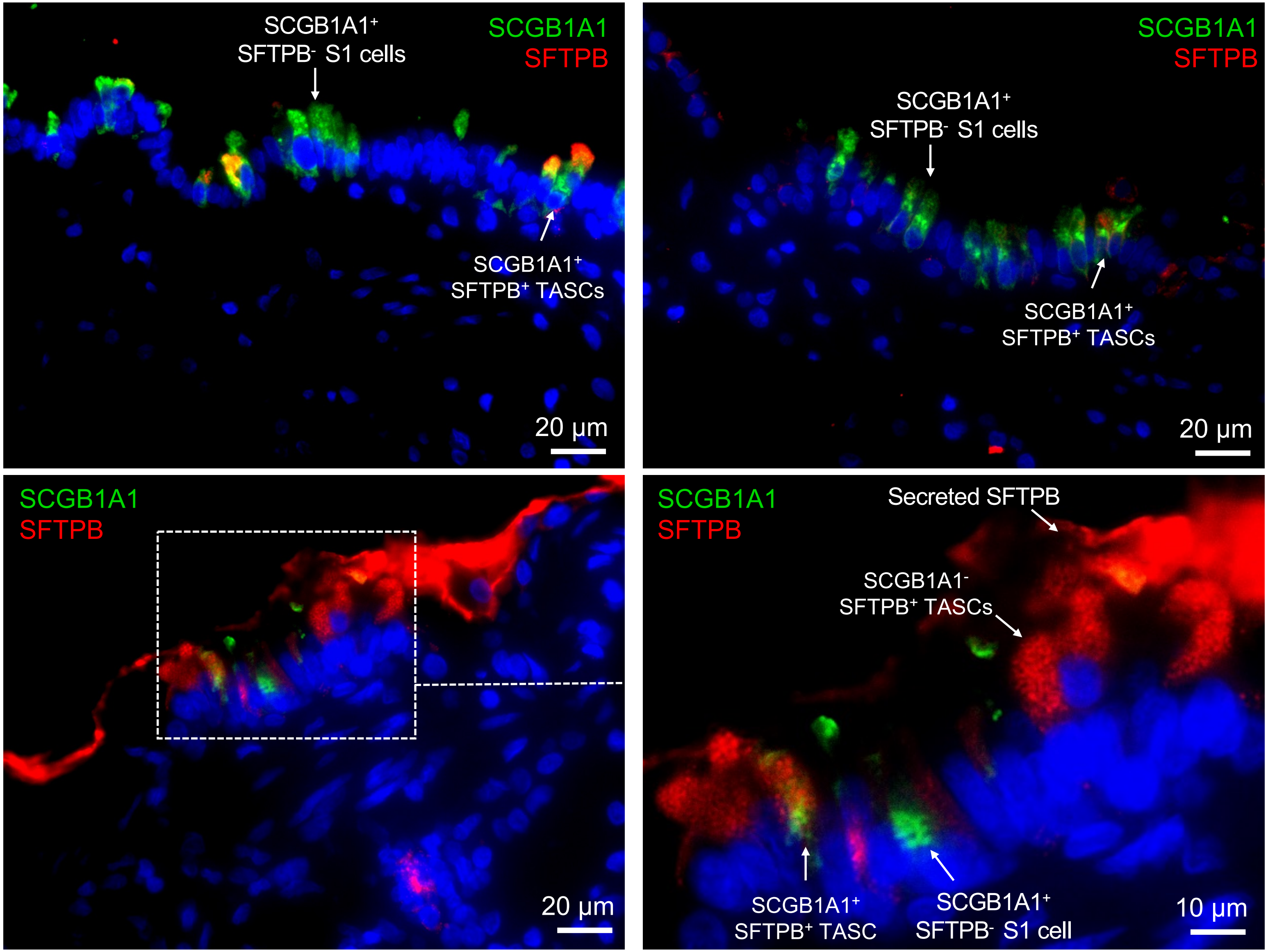

Supplemental Figure 4

**B9**

Distal airways:
Terminal bronchioles
(TBs)

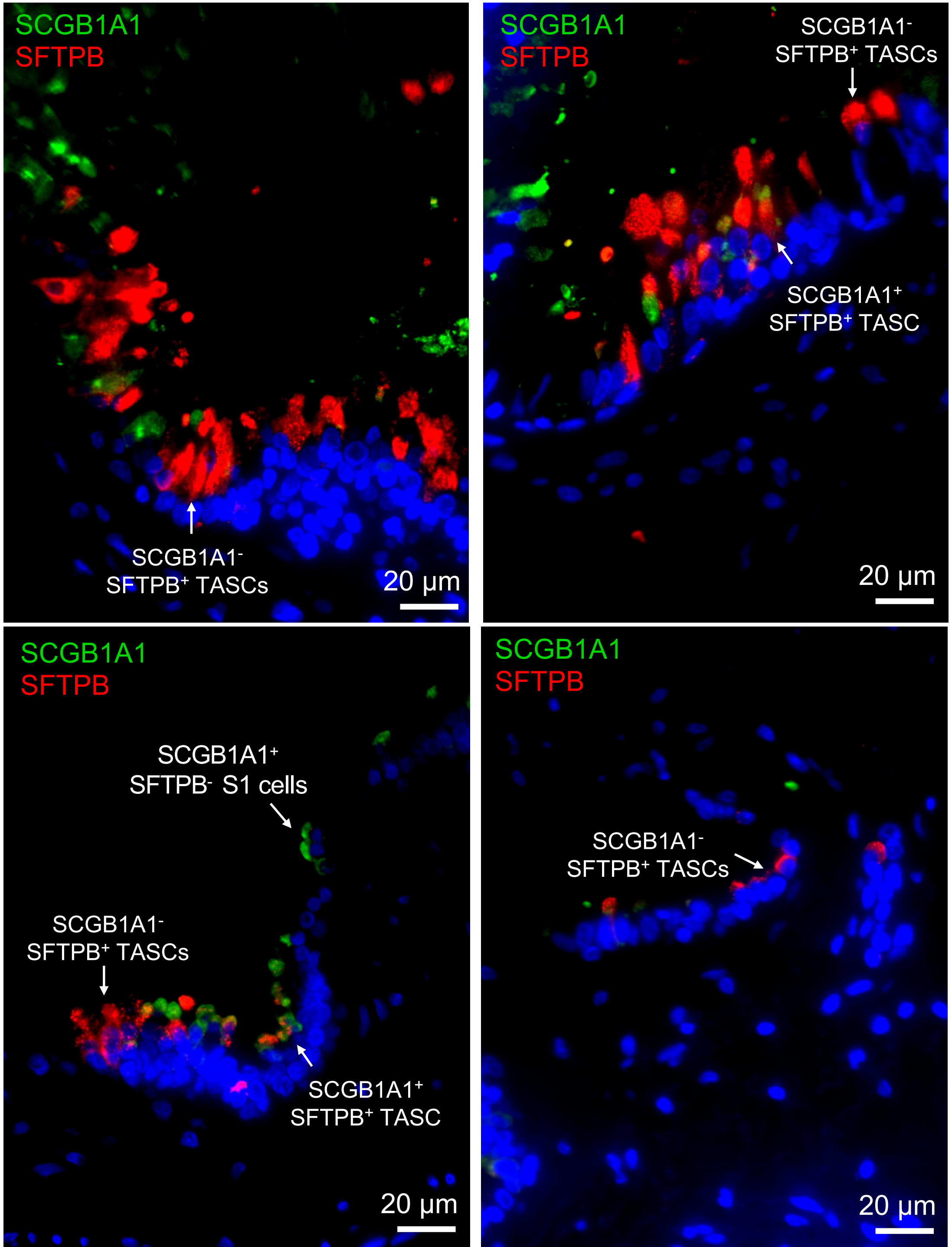

**B10**

Distal airways

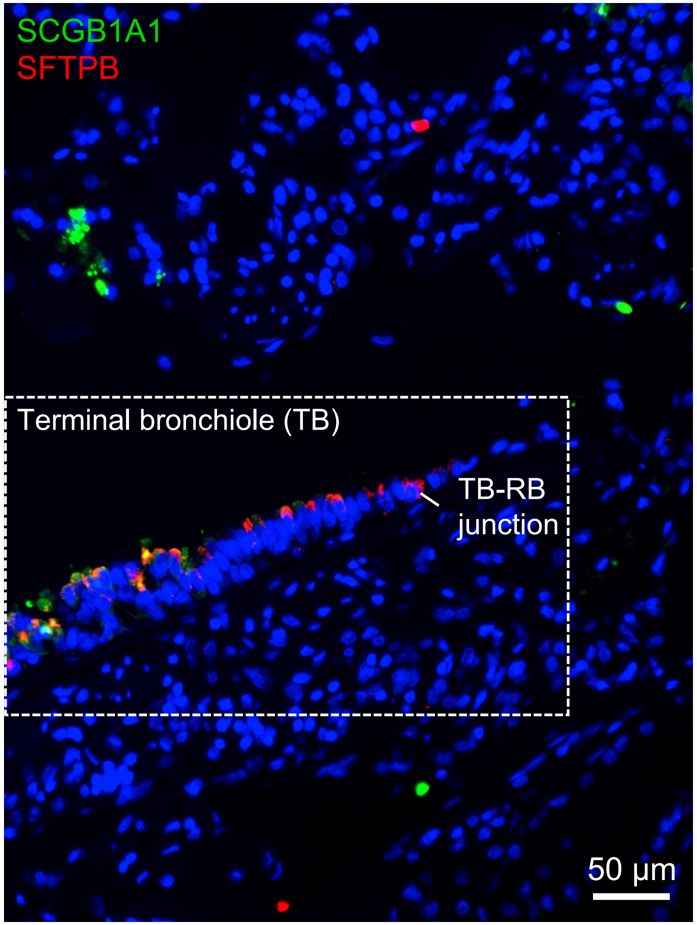


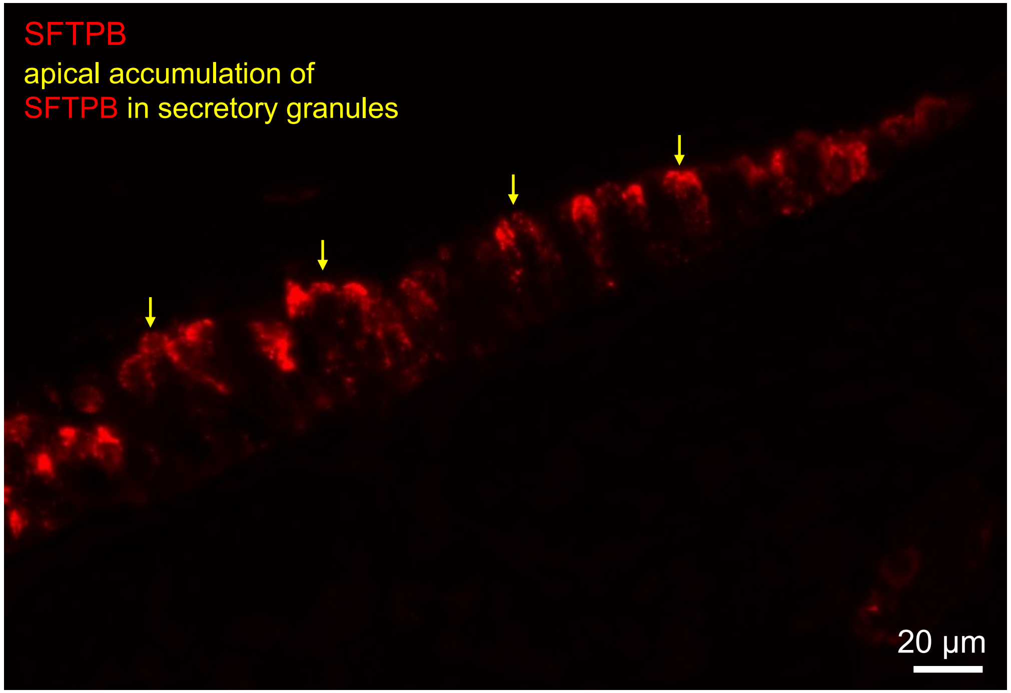

**B11**

Distal airways

Terminal bronchioles (TBs) and respiratory bronchioles (RBs)

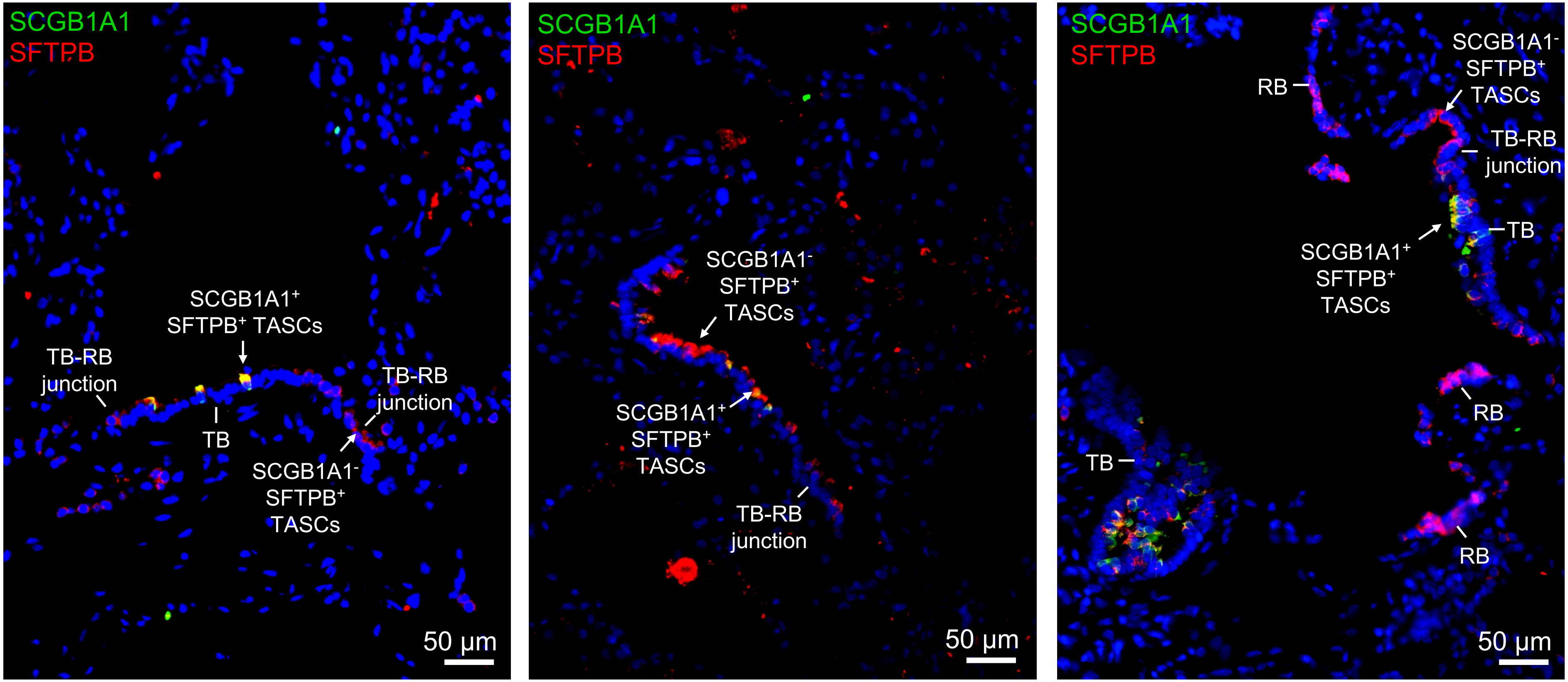

B12

Distal airways

Terminal bronchiole (TB) and respiratory bronchioles (RBs)

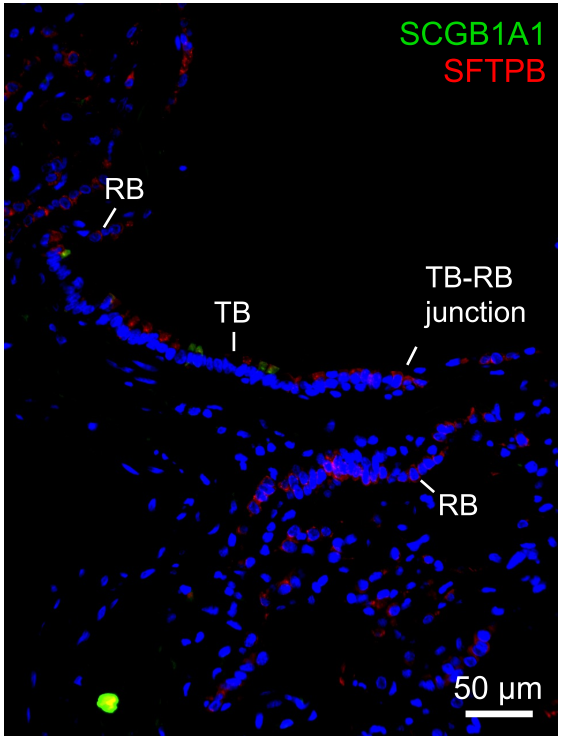


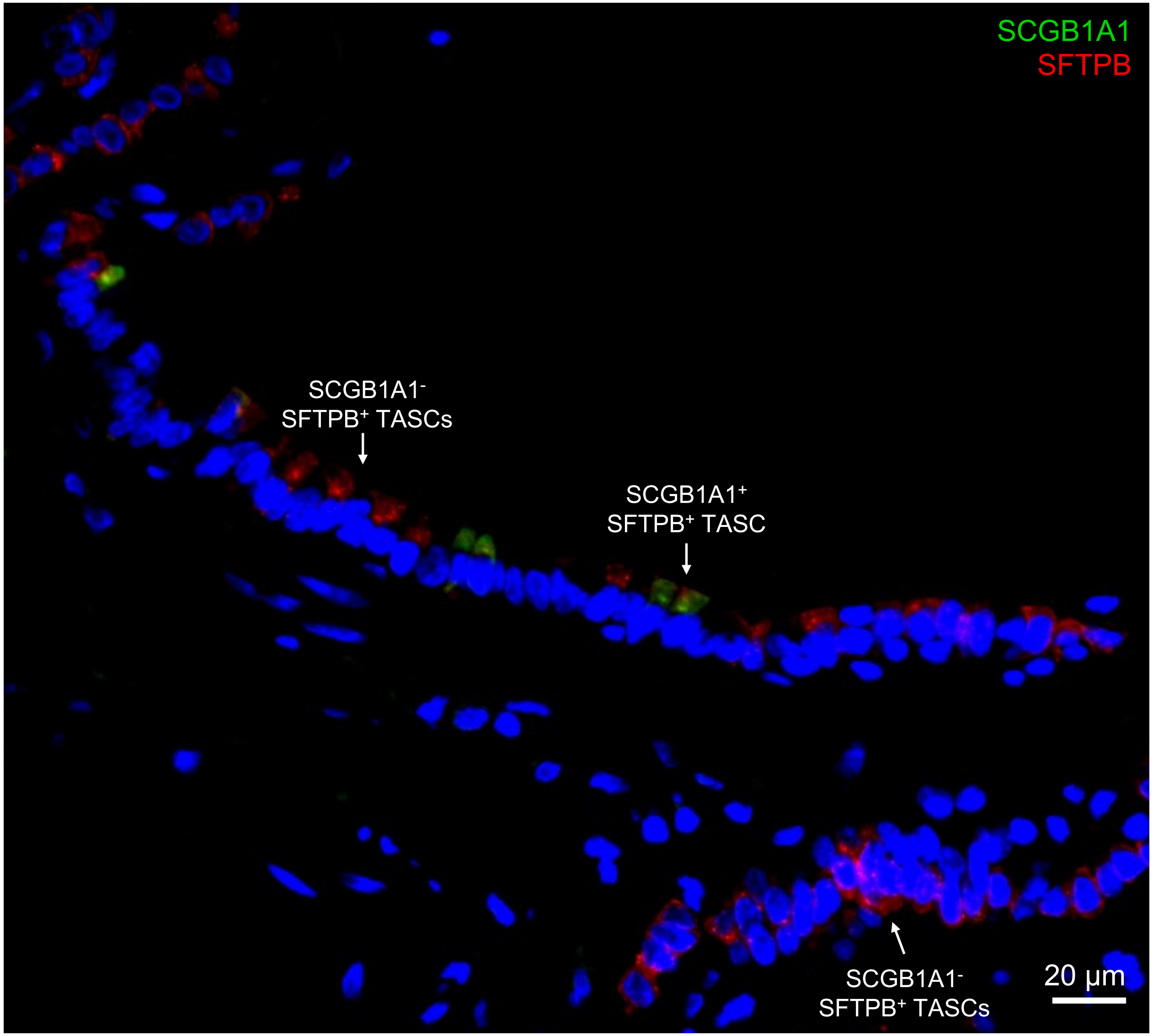
