## Supplementary figures and images for "A Unique Cellular Organization of Human Distal Airways and Its Disarray in Chronic Obstructive Pulmonary Disease"

### Fig. S5

Supplemental Figure 5

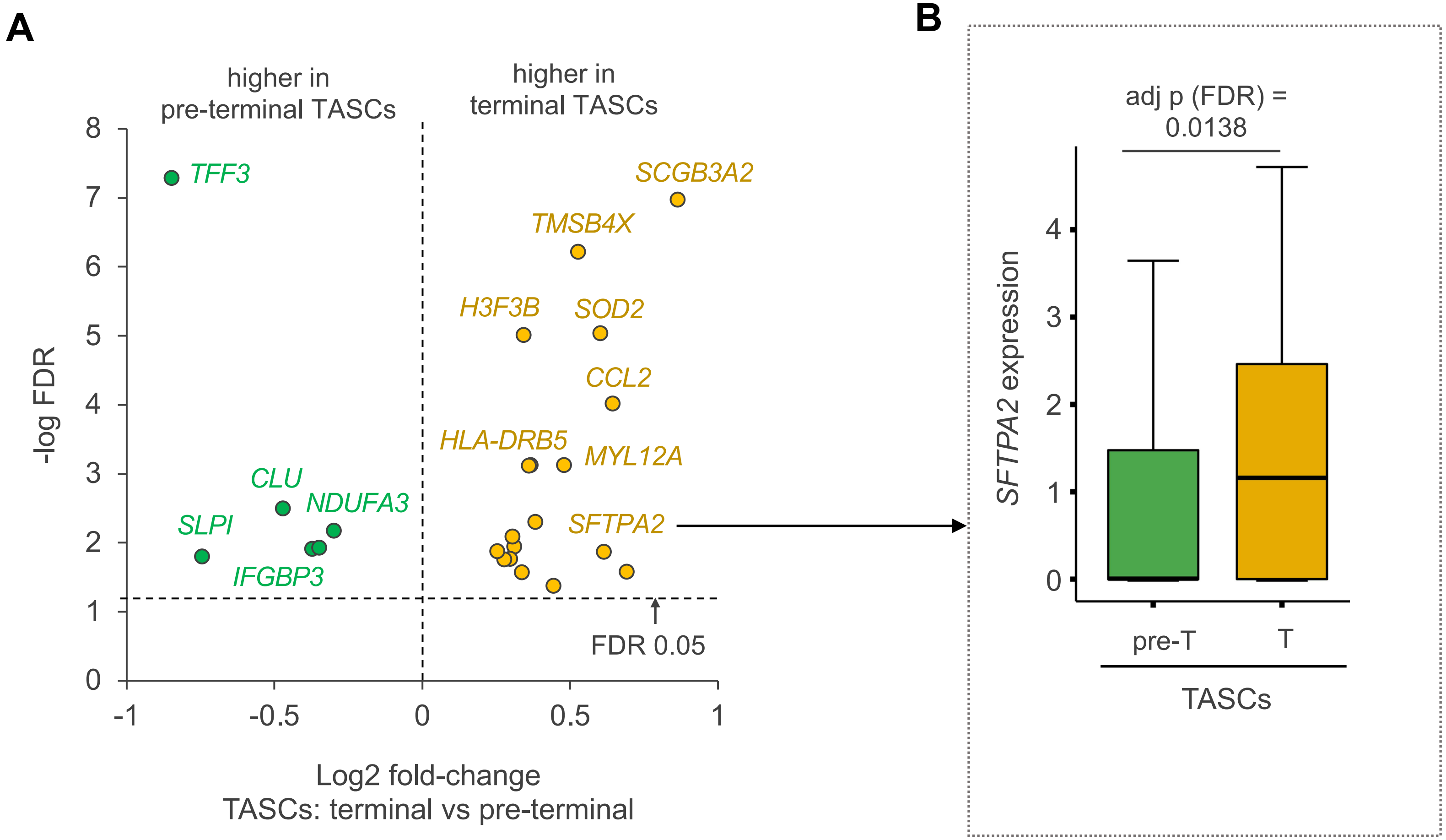

### Fig. S7

**Supplemental Figure 7**

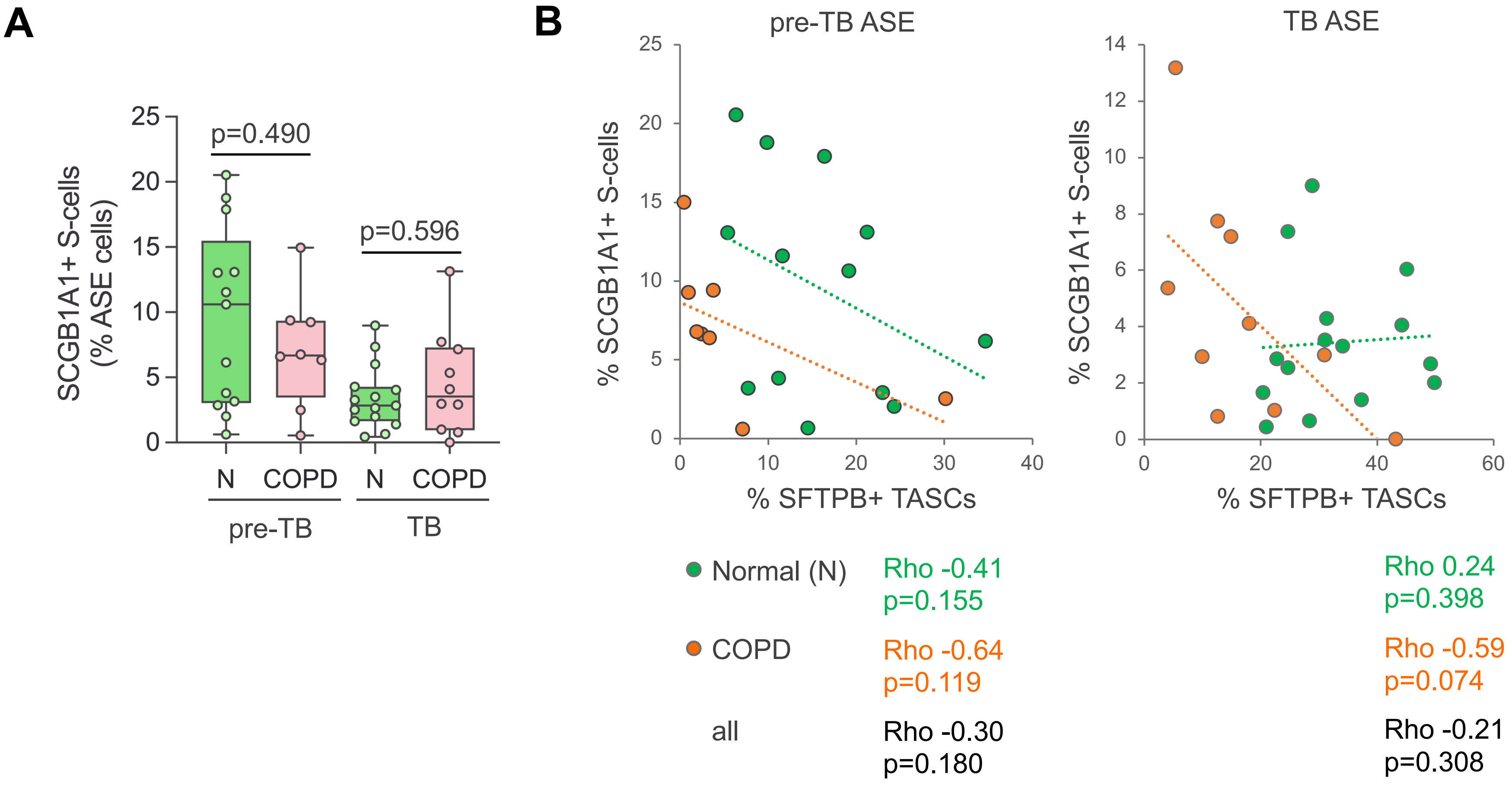

### Fig. S8

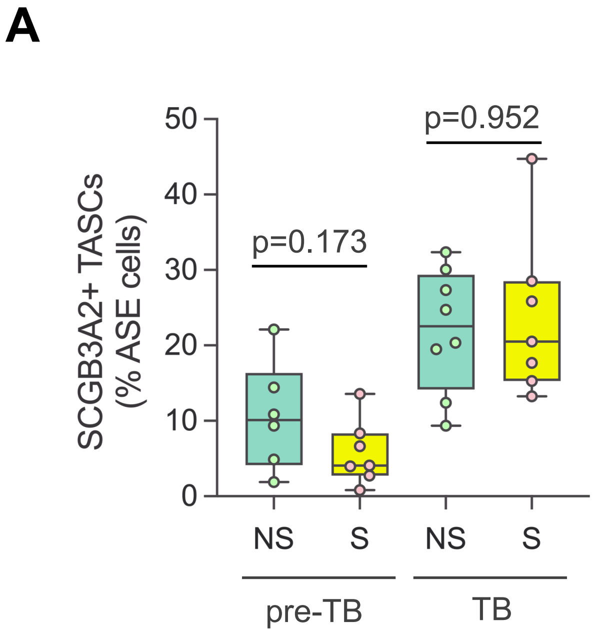

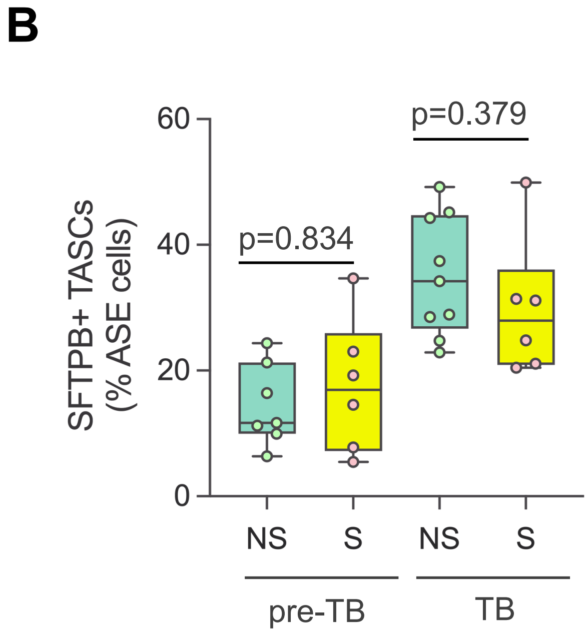
