## Supplementary material for "A Unique Cellular Organization of Human Distal Airways and Its Disarray in Chronic Obstructive Pulmonary Disease": Fig. S6

A$_1$

Supplemental Figure 6

**Pre-TB - SCGB3A2⁺ TASCs - non-diseased lung examples**

red: SCGB3A2 in all images

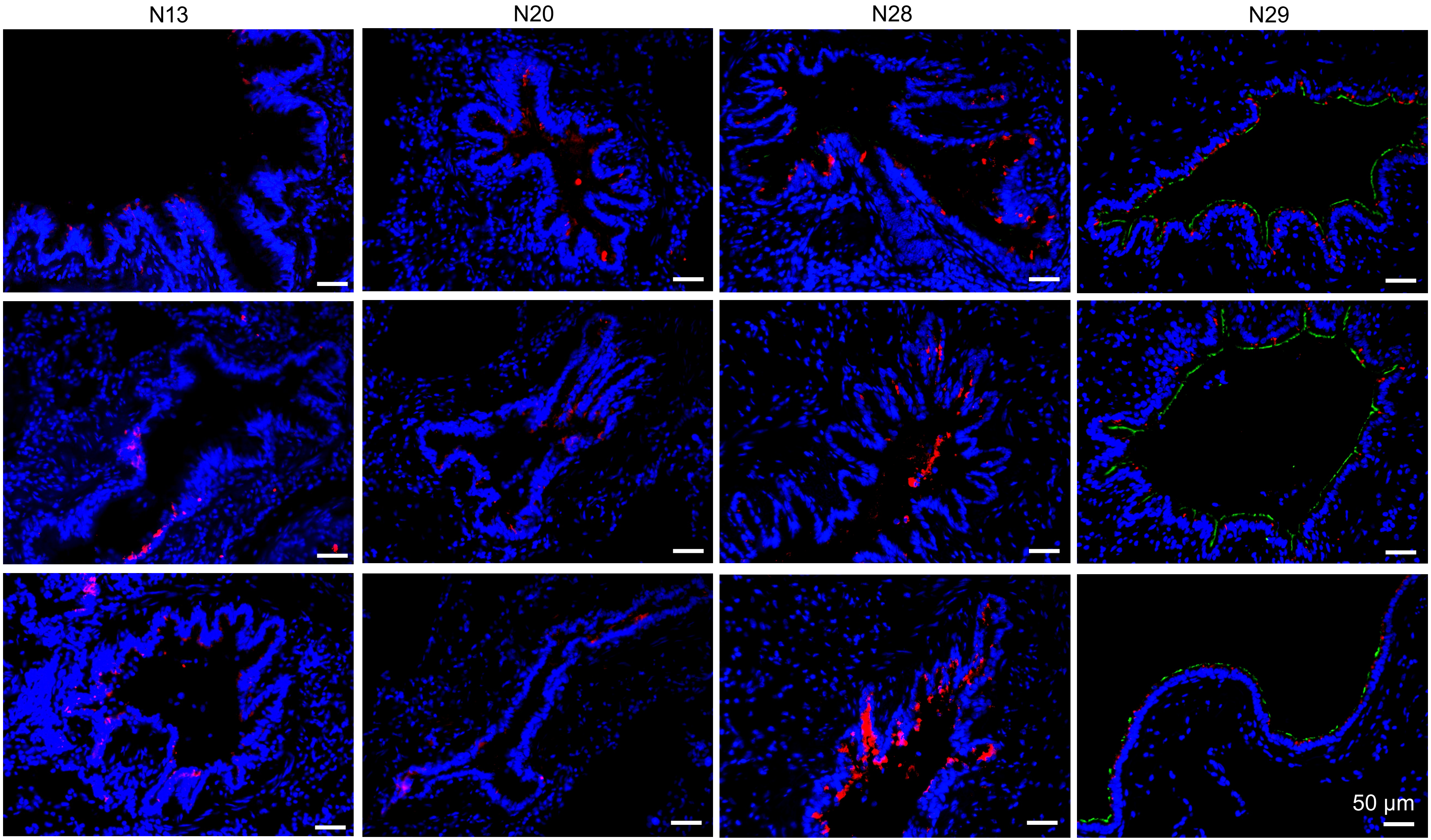


green: TUBB4 (cilia)
additional stain for sample N29

**A$_2$**

**Pre-TB - SCGB3A2$^+$ TASCs – COPD lung examples**

red: SCGB3A2 in all images

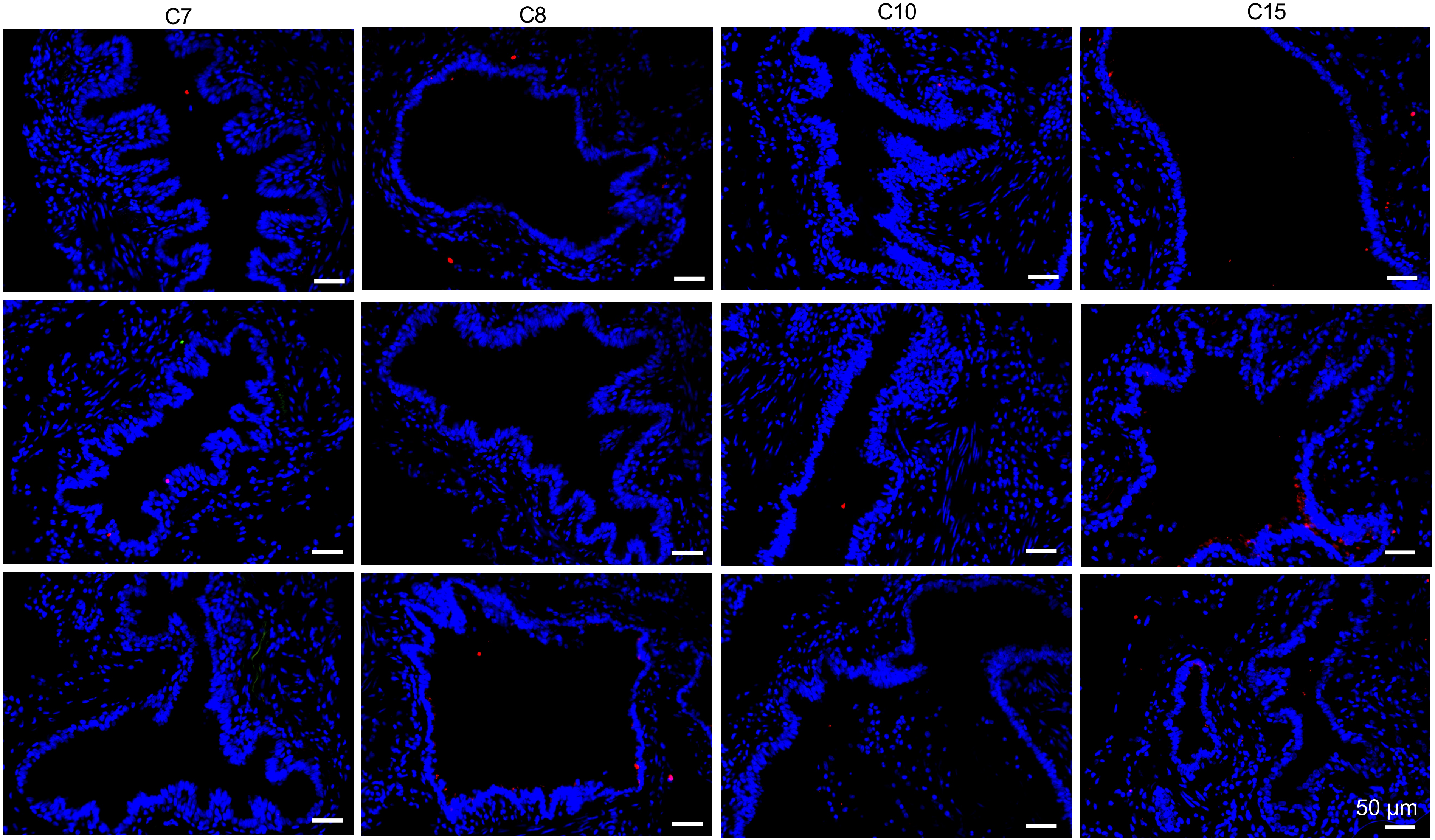

# A$_3$

**TB - SCGB3A2$^+$ TASCs - non-diseased lung examples**

red: SCGB3A2 in all images

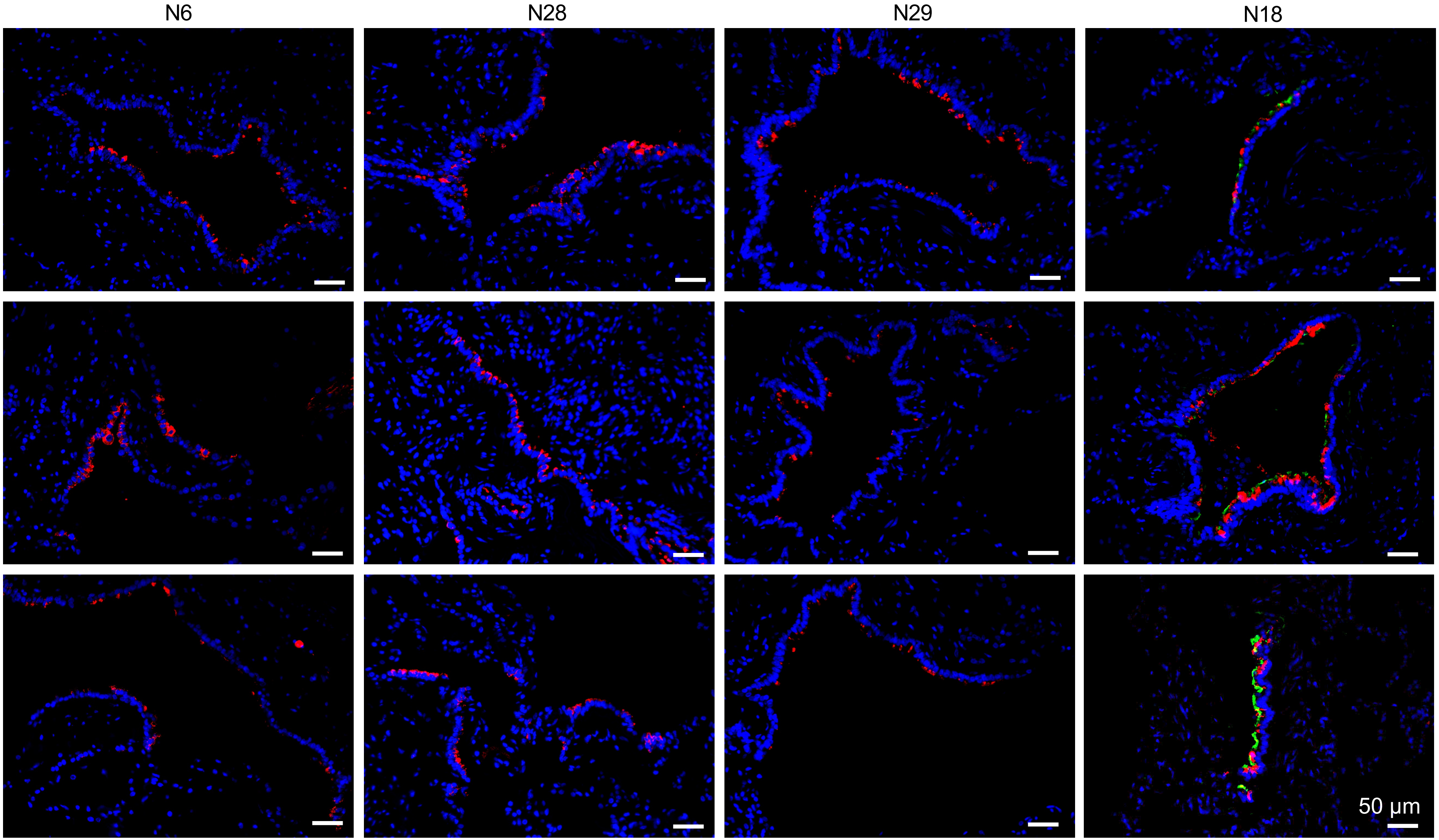


green: TUBB4 (cilia)
additional stain for sample N29

**A$_4$**

**TB - SCGB3A2$^+$ TASCs - COPD lung examples**

red: SCGB3A2 in all images

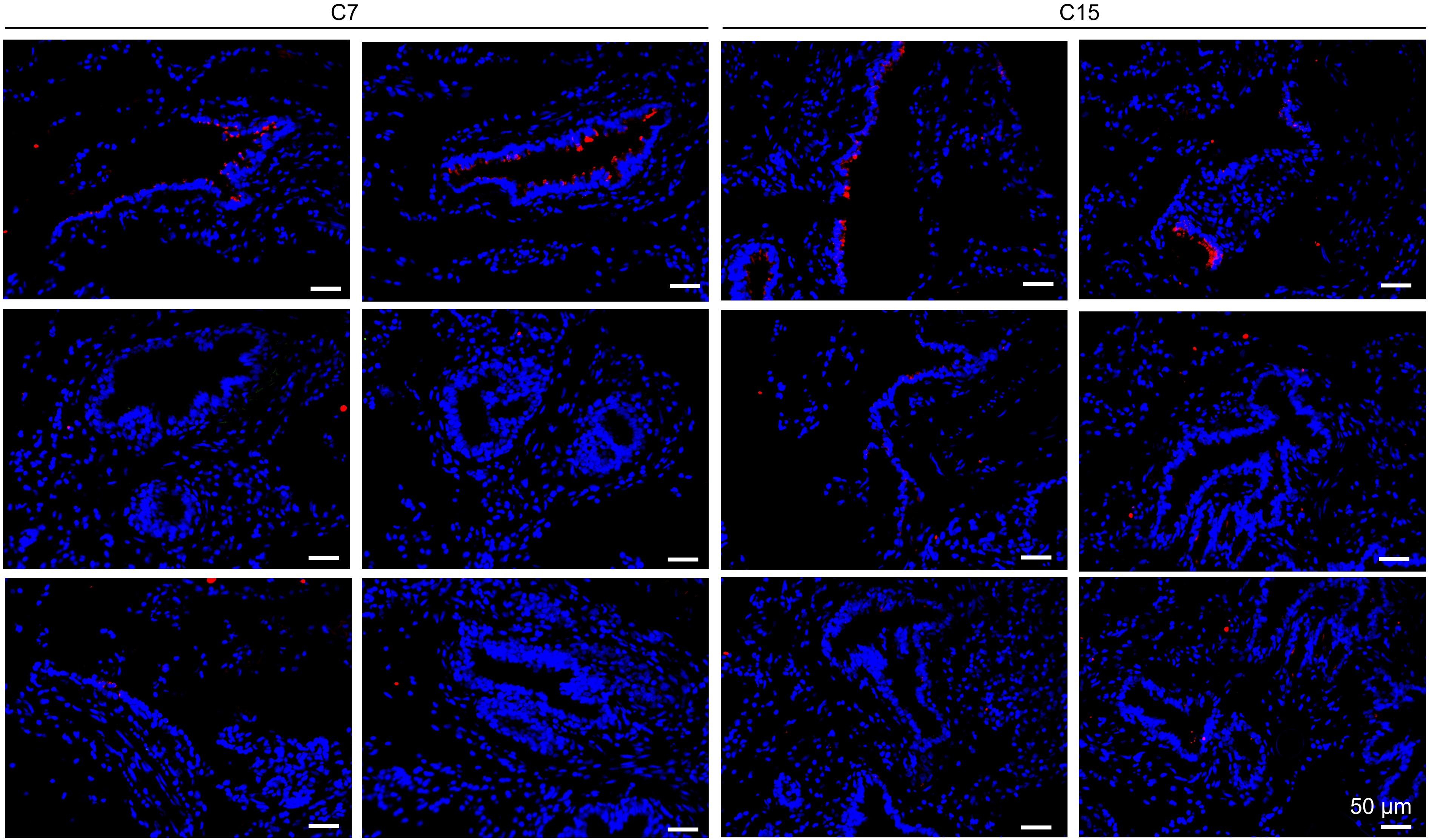

$A_4$ (continued)

**TB - SCGB3A2⁺ TASCs - COPD lung examples**

red: SCGB3A2 in all images

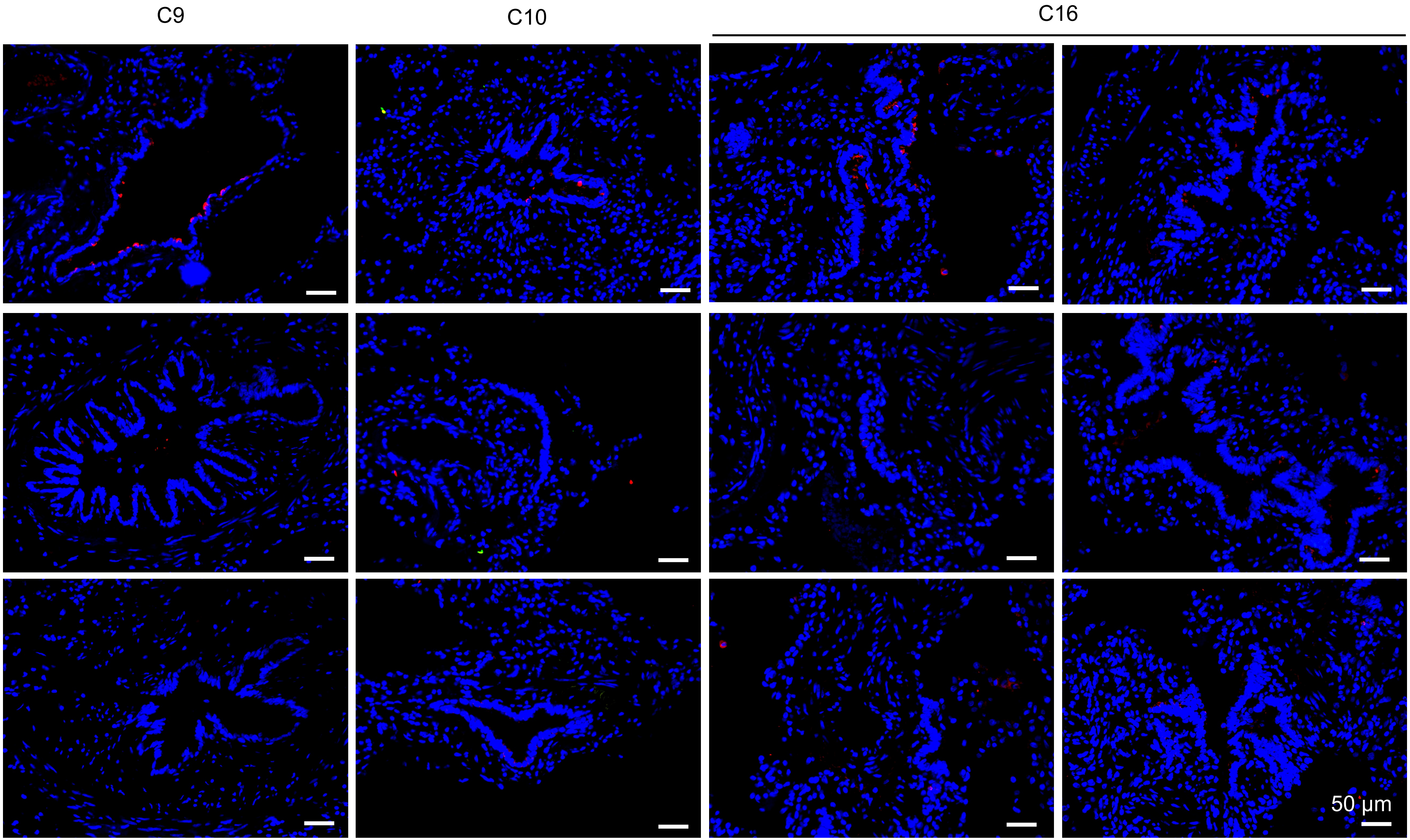

**B$_1$**

**pre-TB - SFTPB$^+$ TASCs – normal lung examples**

red: SFTPB green: SCGB1A1 in all images

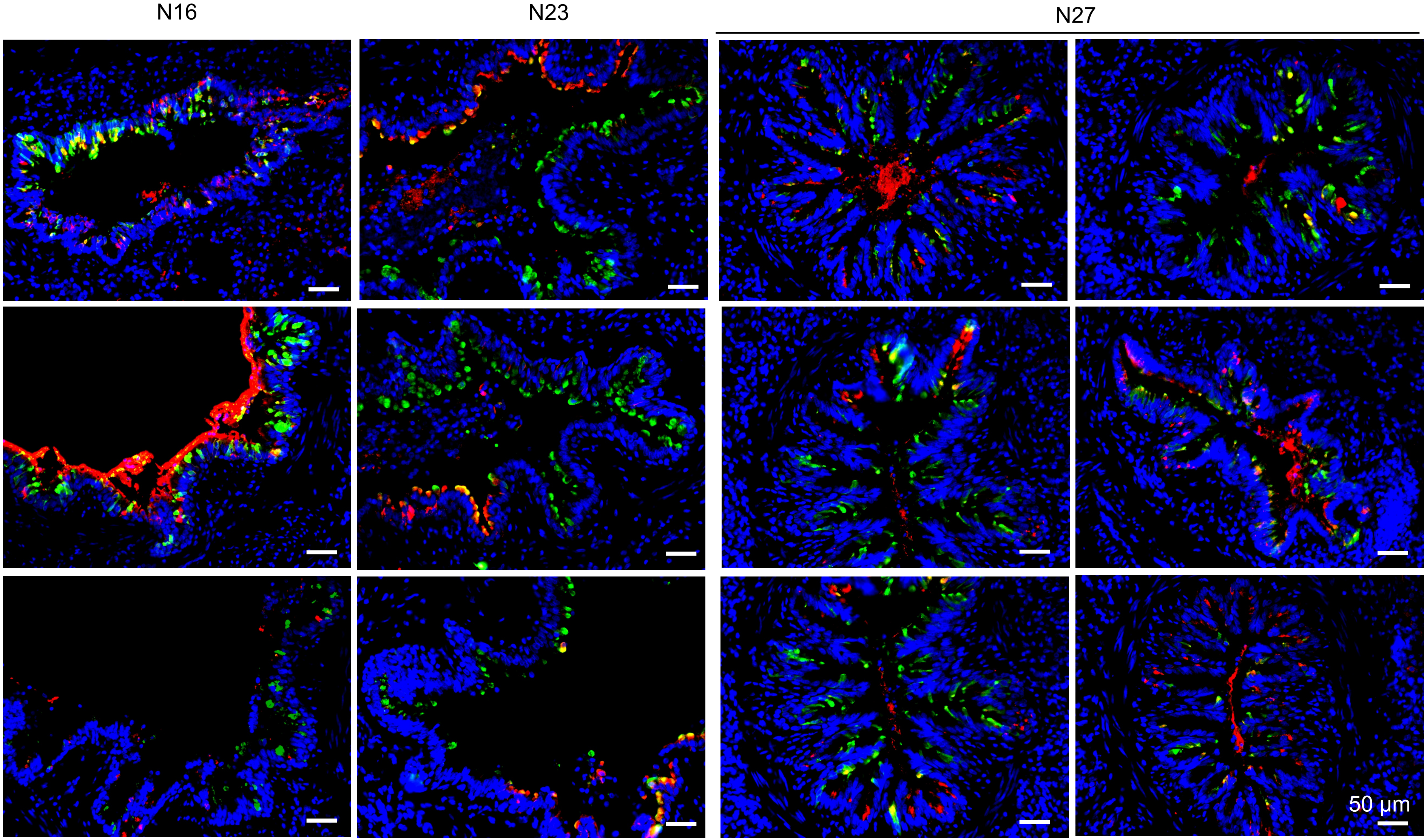

**B$_2$**



**pre-TB - SFTPB$^+$ TASCs – COPD lung examples**

red: SFTPB green: SCGB1A1 in all images

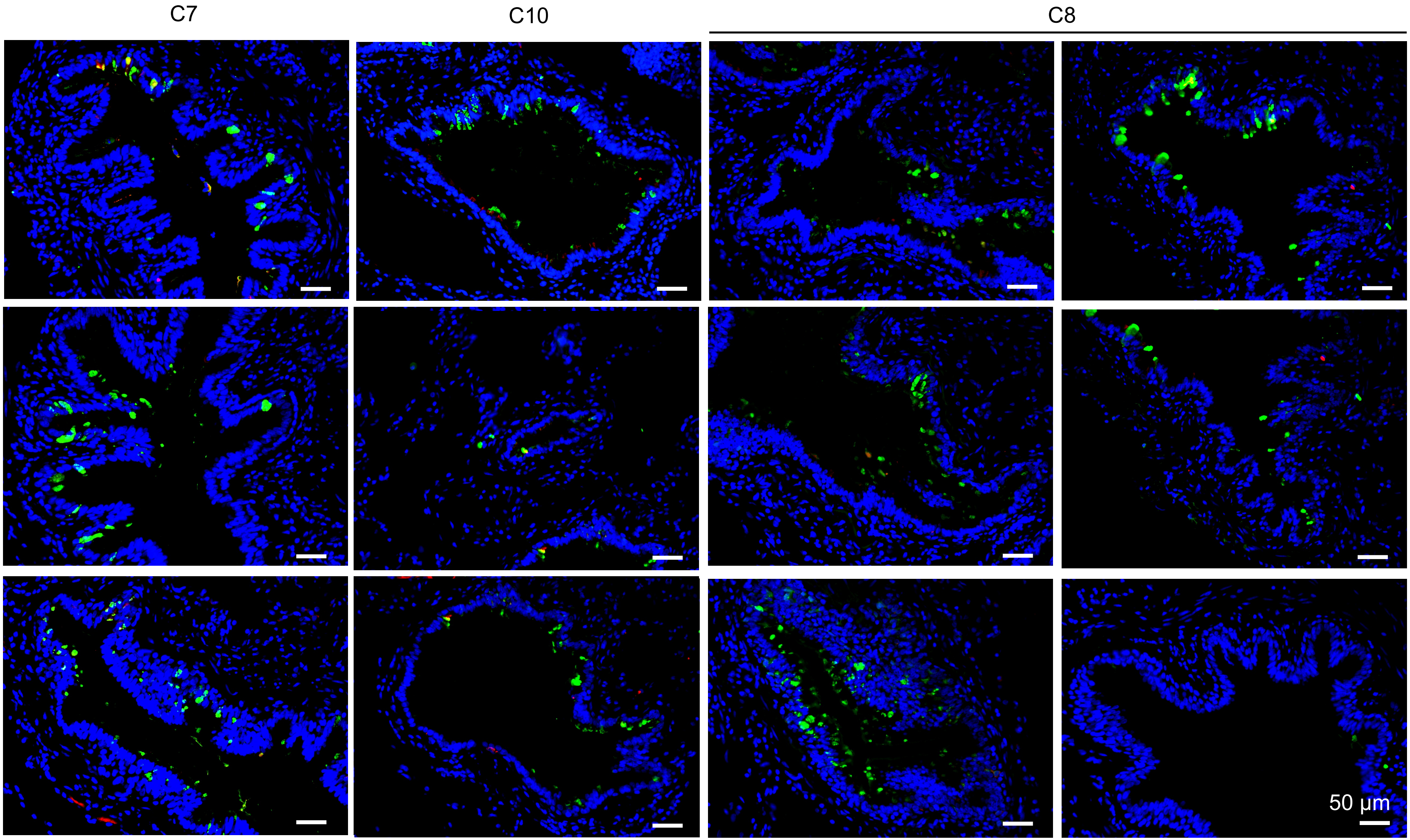

B3

**TB - SFTPB⁺ TASCs – normal lung examples**

red: SFTPB green: SCGB1A1 in all images

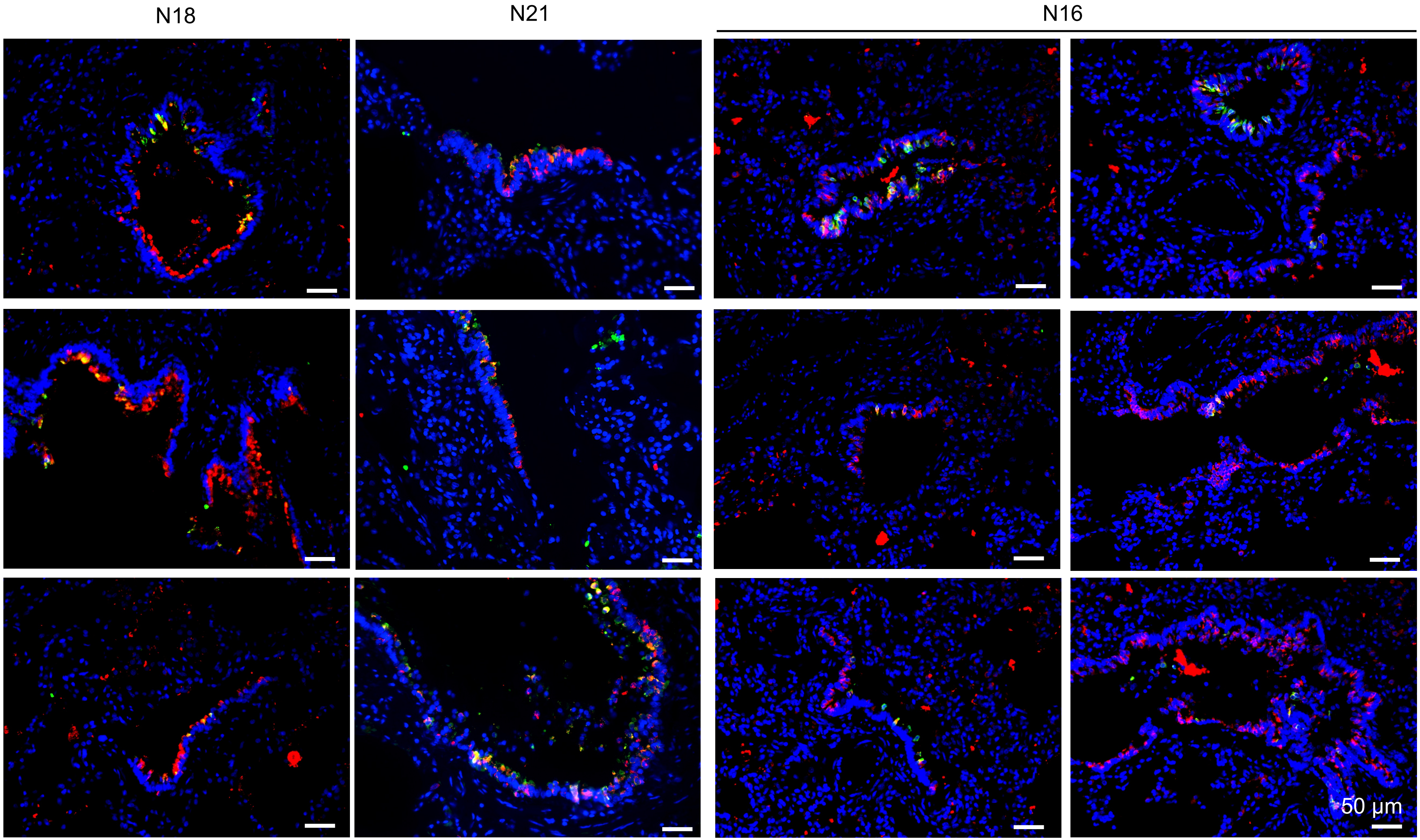

**B$_4$**

**TB - SFTPB$^+$ TASCs – COPD lung examples**

red: SFTPB green: SCGB1A1 in all images

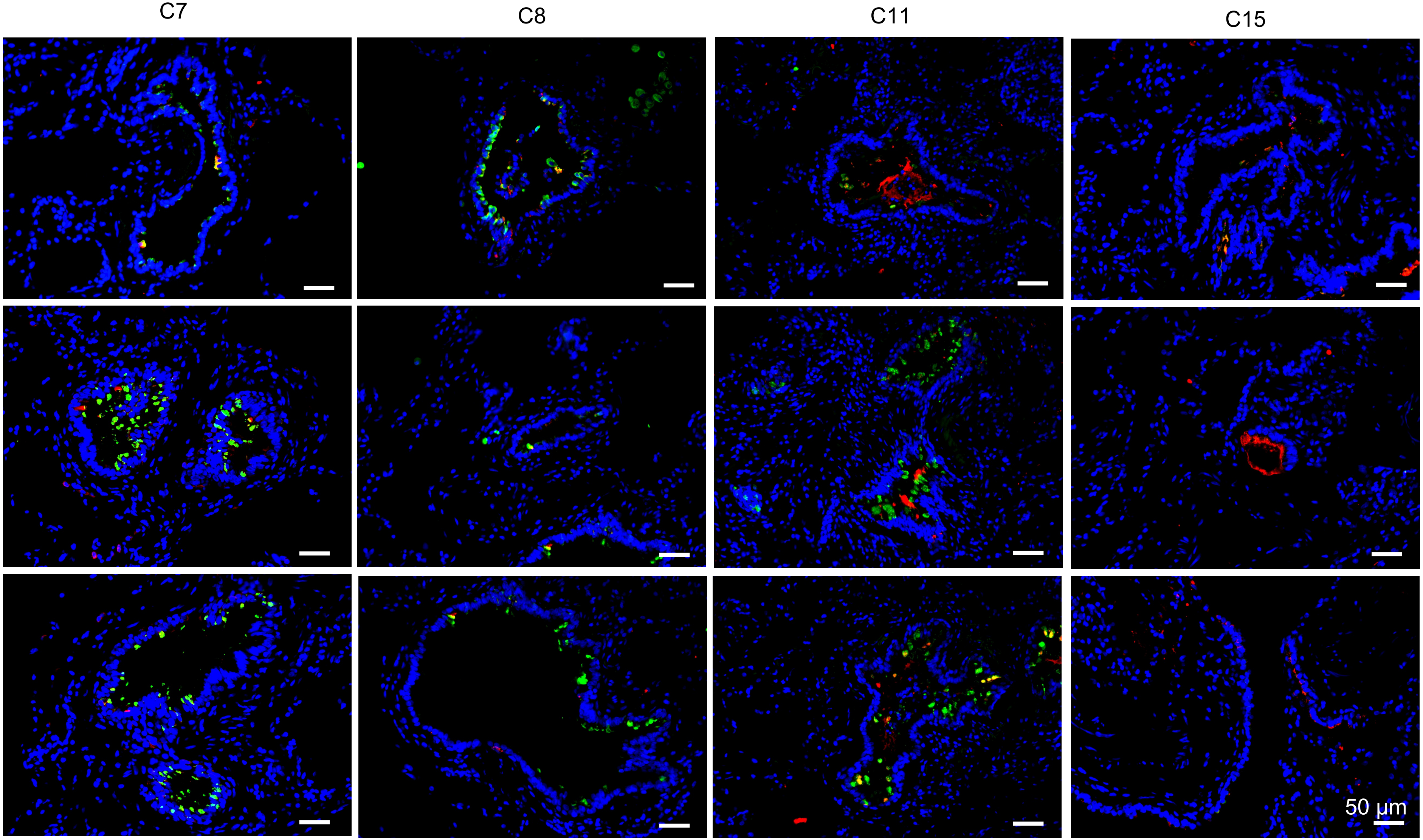
