## Supplementary material for "A Unique Cellular Organization of Human Distal Airways and Its Disarray in Chronic Obstructive Pulmonary Disease": Fig. S9

**Supplemental Figure 9**

**A$_1$**

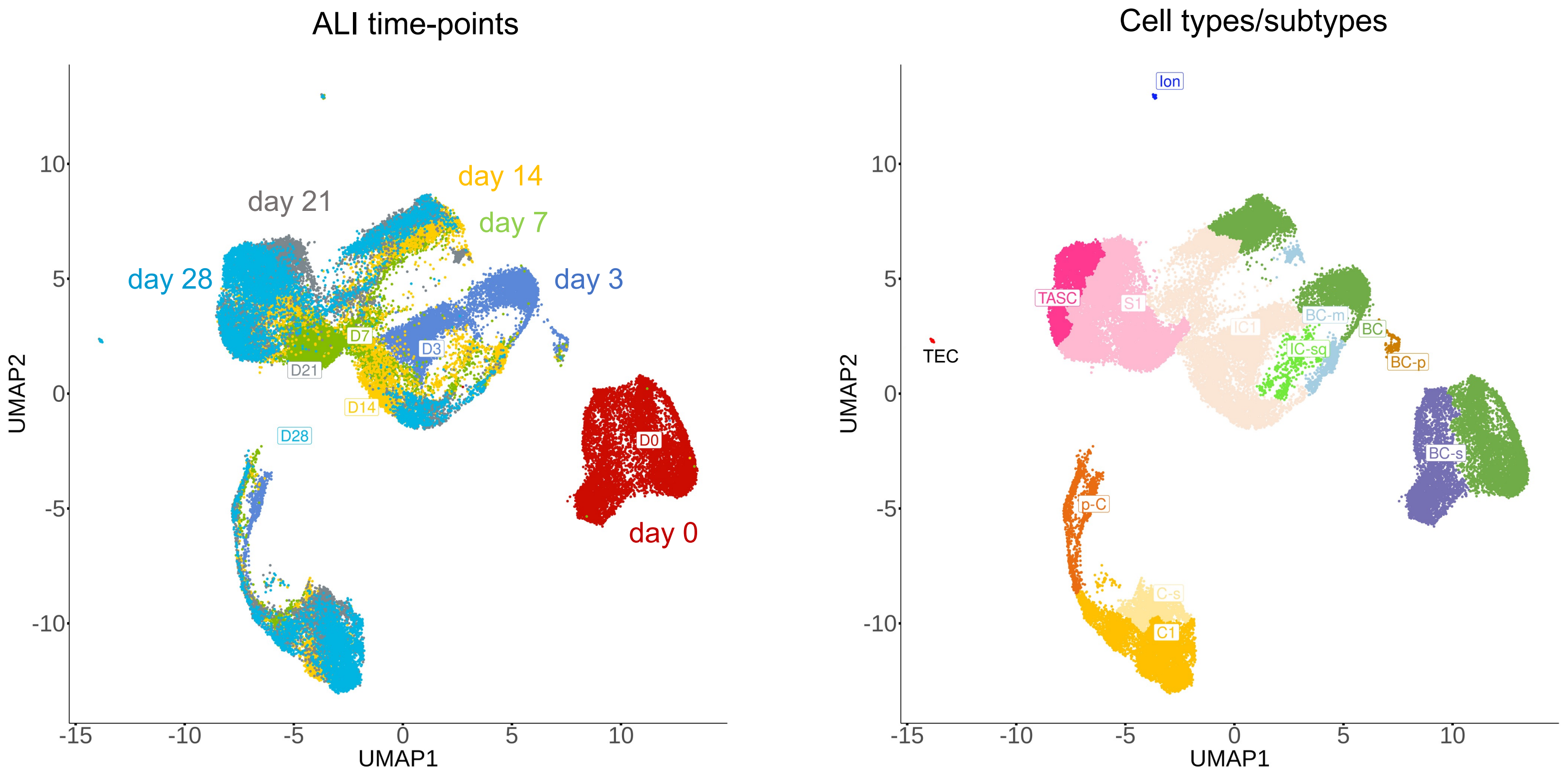

**A$_2$**

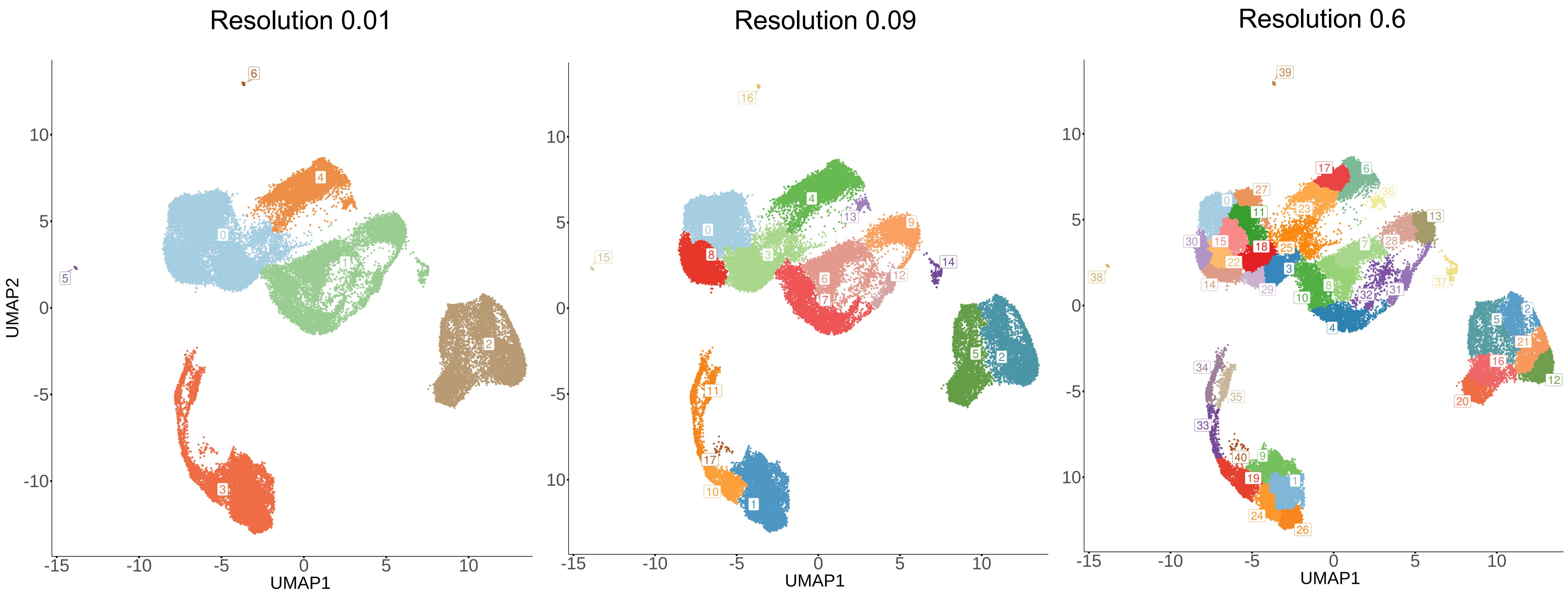

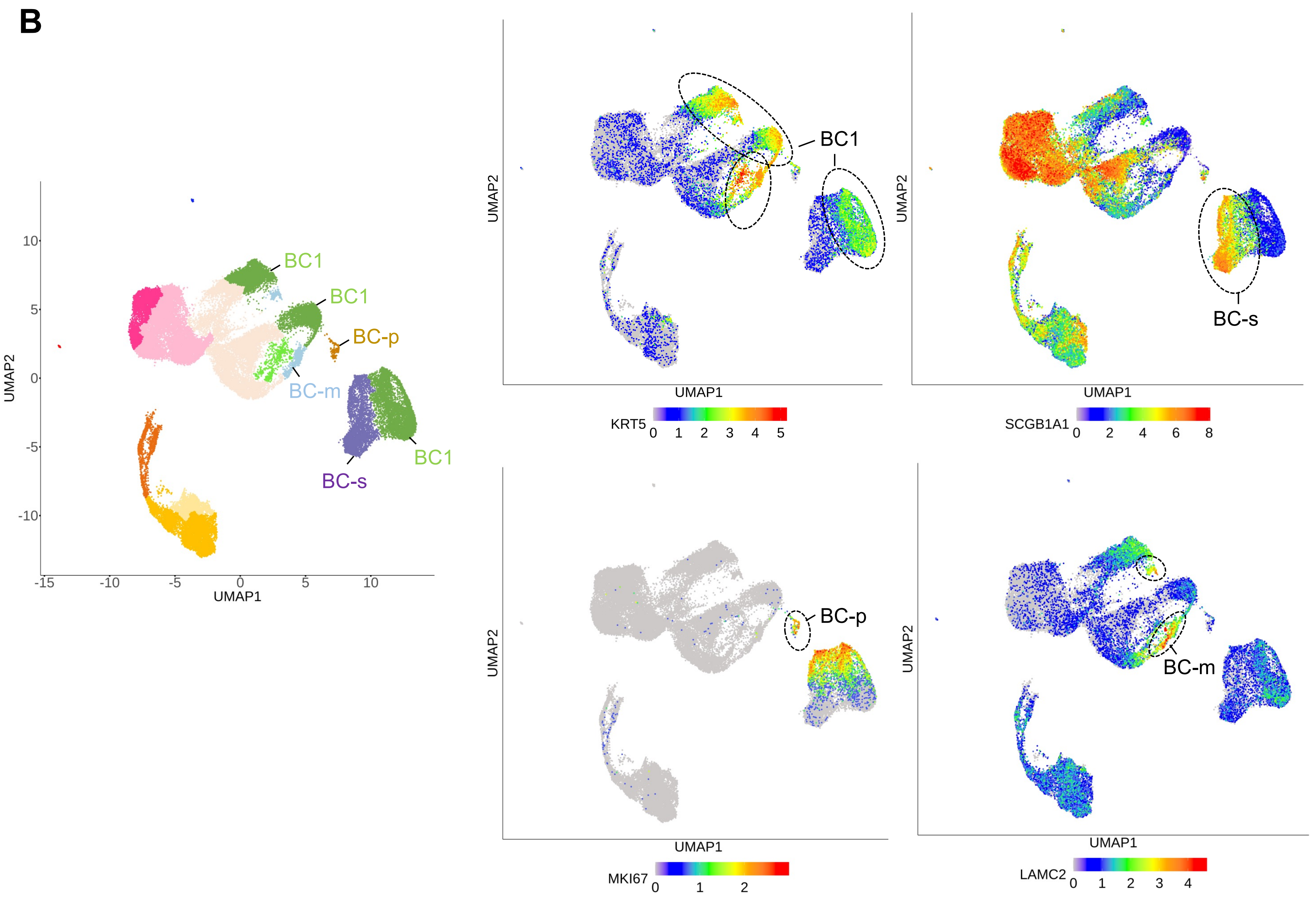
B
UMAP2
UMAP1
10
5
0
-5
-10
-15
-10
-5
0
5
10
BC1
BC1
BC-p
BC-m
BC1
BC-s
KRT5
0 1 2 3 4 5
SCGB1A1
0 2 4 6 8
BC-s
MKI67
0 1 2
BC-p
LAMC2
0 1 2 3 4
BC-m

**Supplemental Figure 9**

**C**

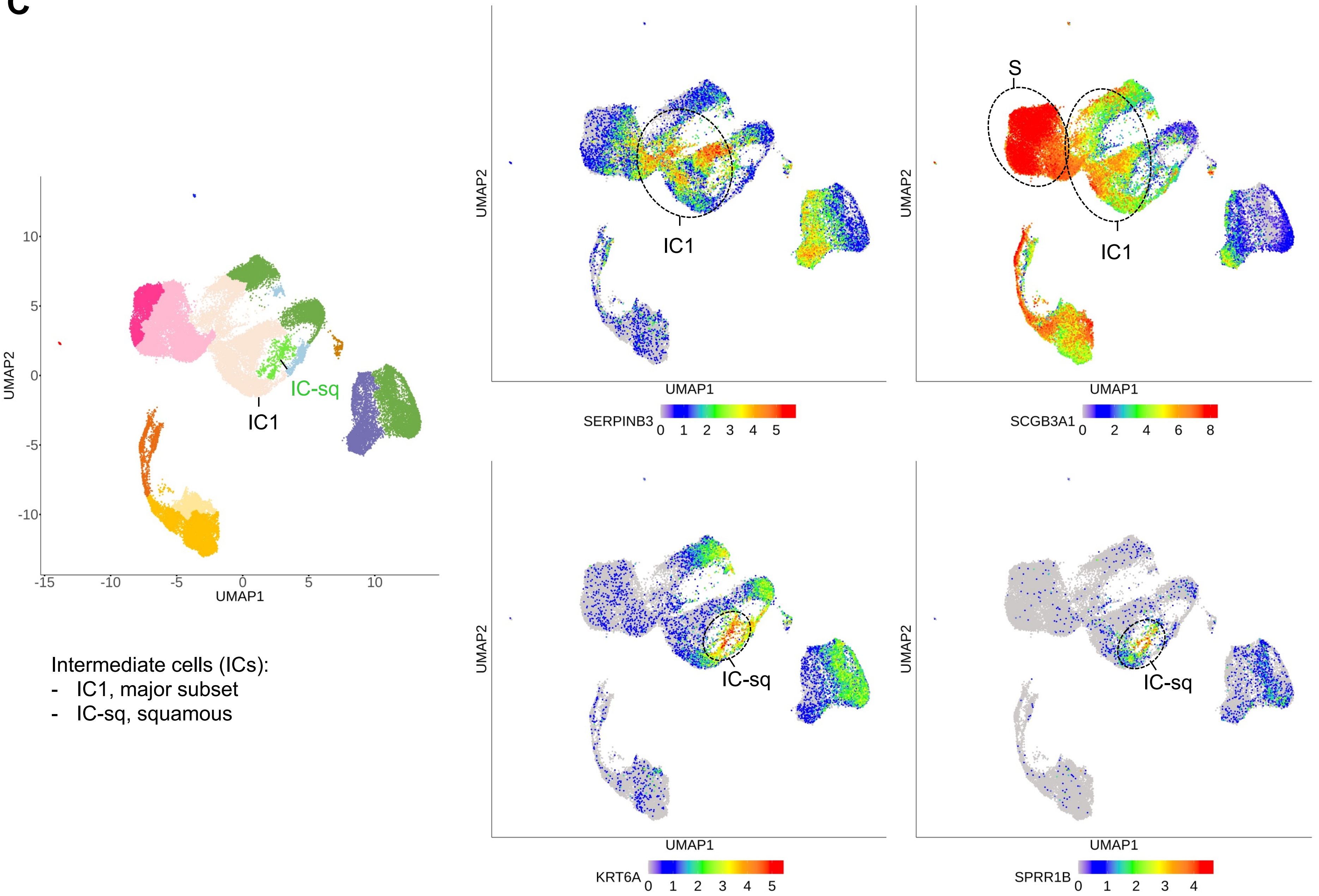

### Supplemental Figure 9

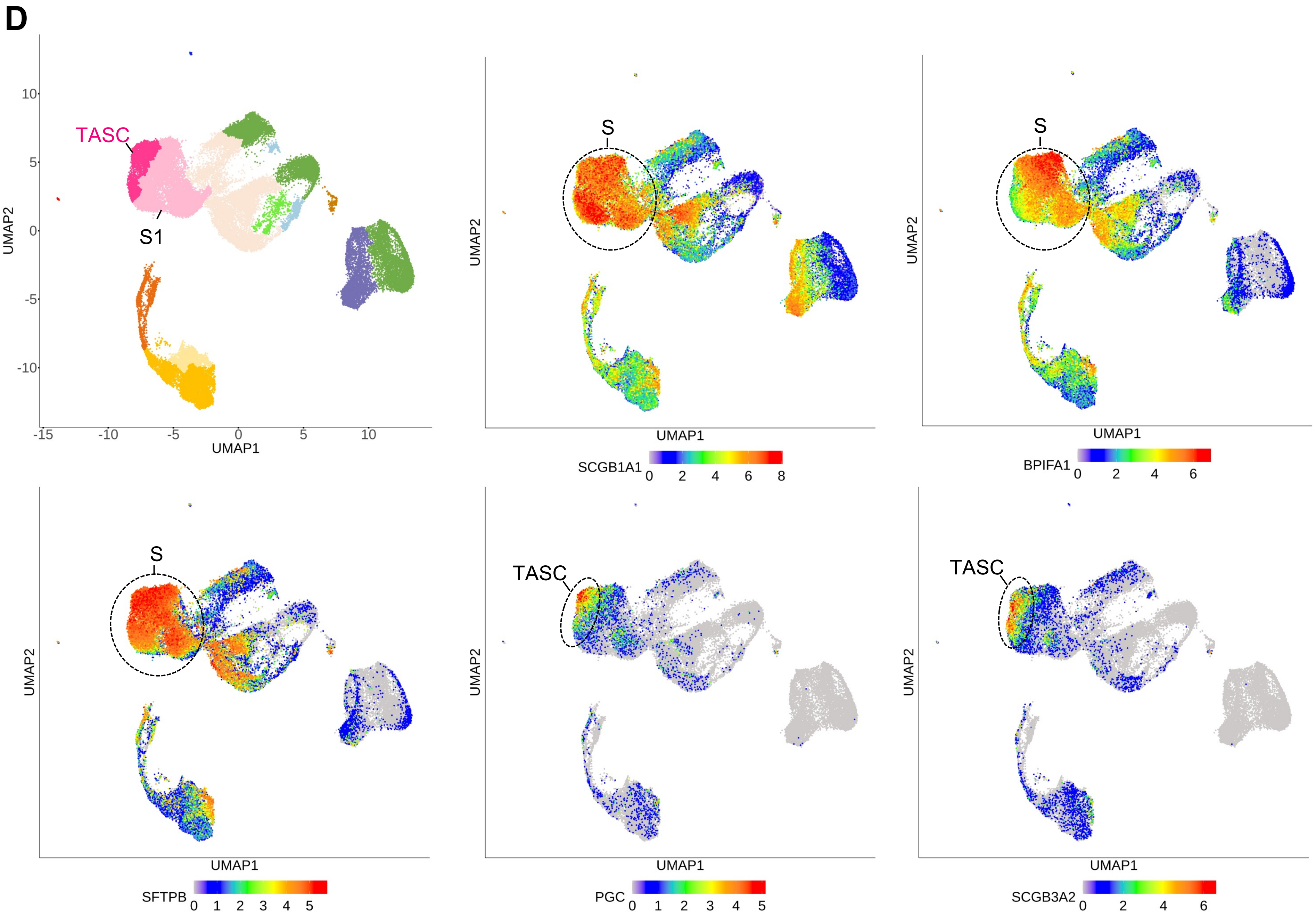

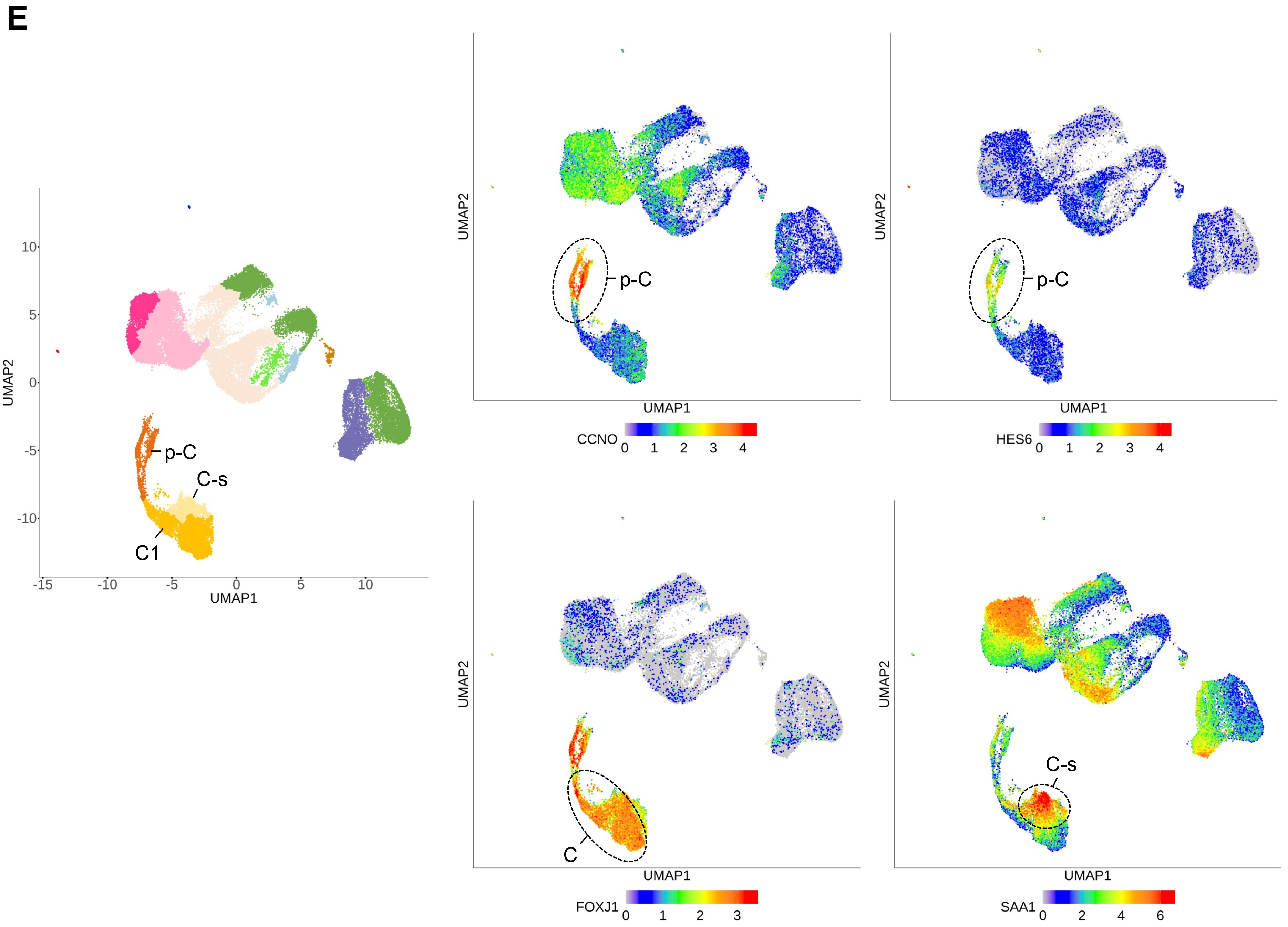
E
UMAP2
UMAP1
10
5
0
-5
-10
-15
-10
-5
0
5
10
p-C
C-s
C1
UMAP2
UMAP1
p-C
CCNO
0
1
2
3
4
UMAP2
UMAP1
p-C
HES6
0
1
2
3
4
UMAP2
UMAP1
C
FOXJ1
0
1
2
3
UMAP2
UMAP1
C-s
SAA1
0
2
4
6

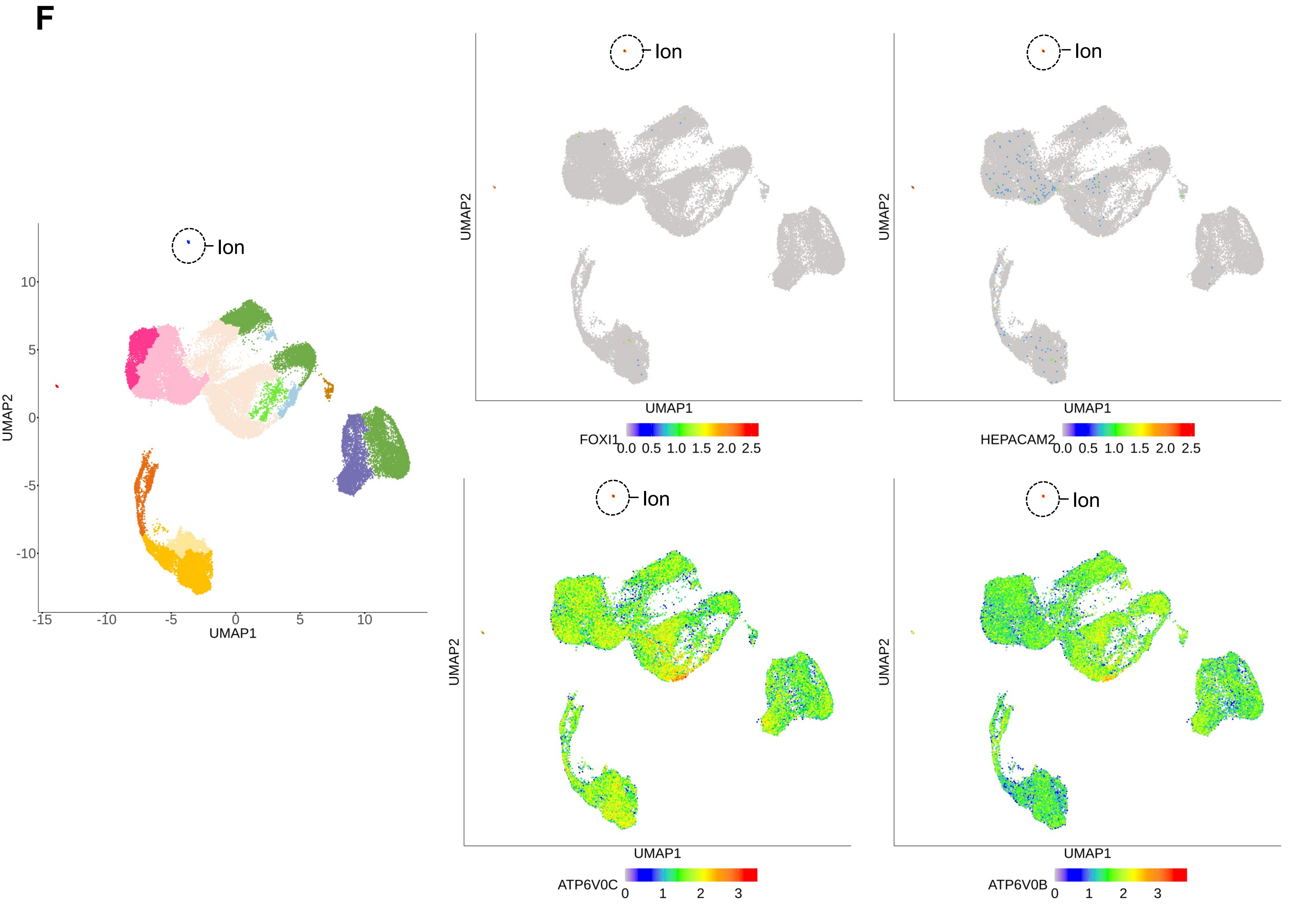
F
Ion
UMAP2
UMAP1
10
5
0
-5
-10
-15
-10
-5
0
5
10
FOXI1
HEPACAM2
0.0 0.5 1.0 1.5 2.0 2.5
ATP6V0C
ATP6V0B
0 1 2 3

**G**

(continued on next page)

TLC, tuft-like cells

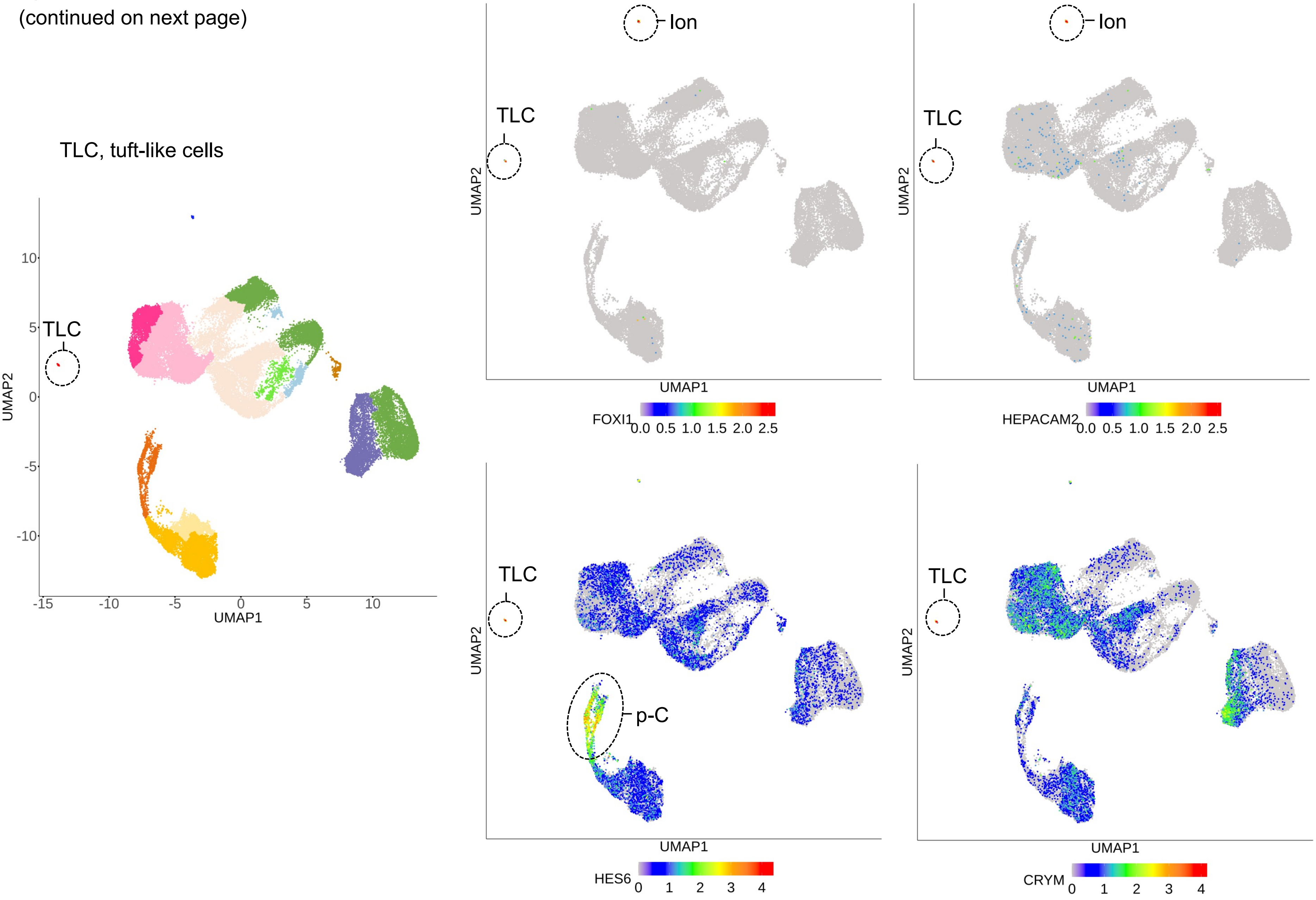

**G**
(continued)

TLC, tuft-like cells

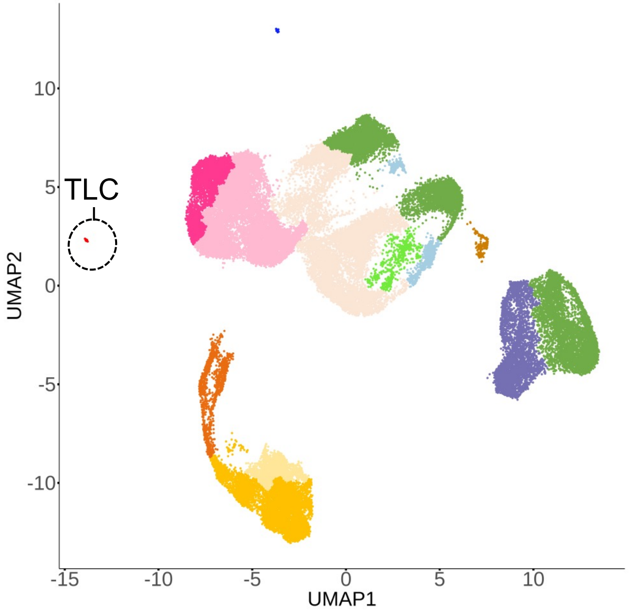


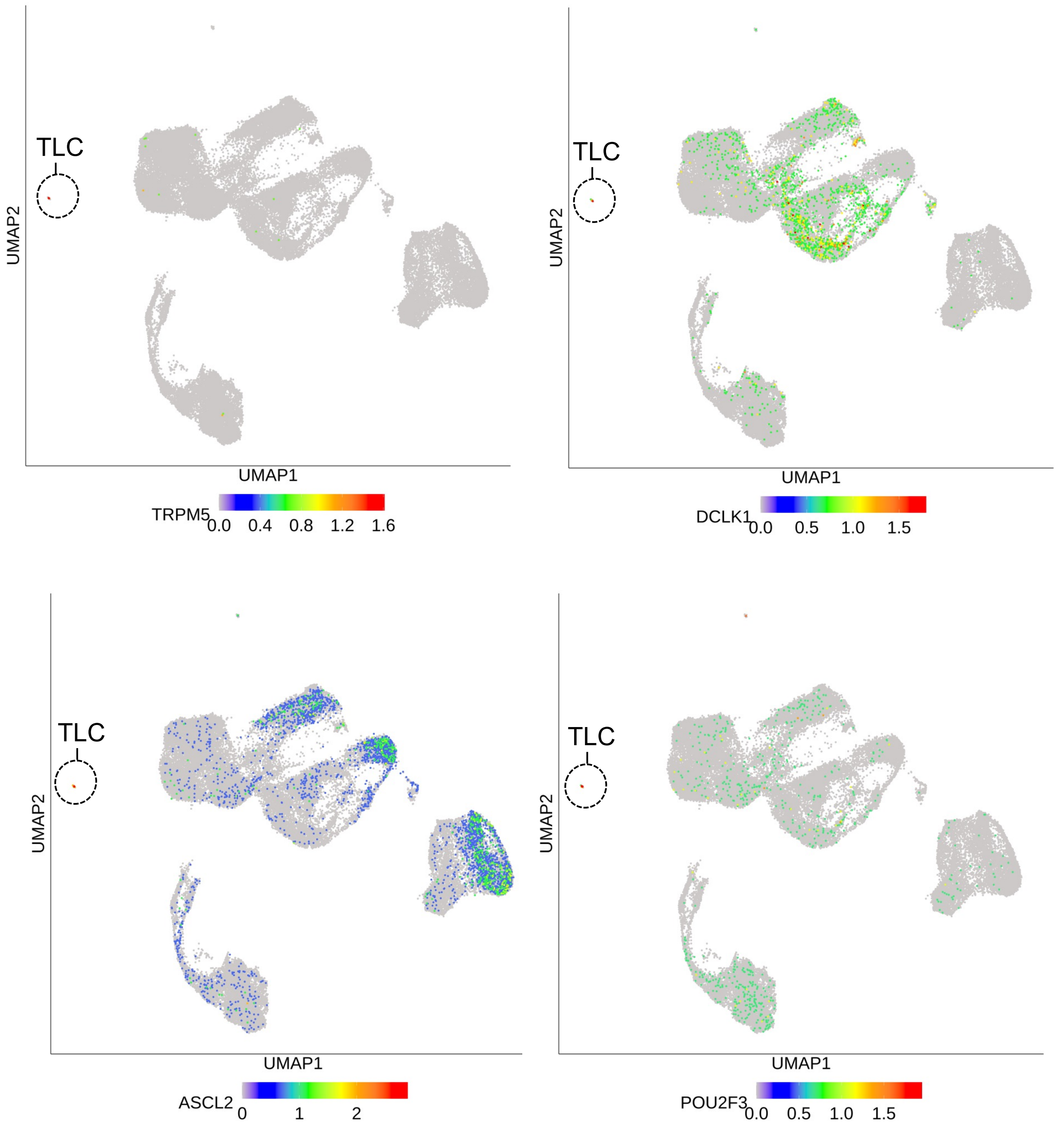

**G**
(continued)

TLC, tuft-like cells

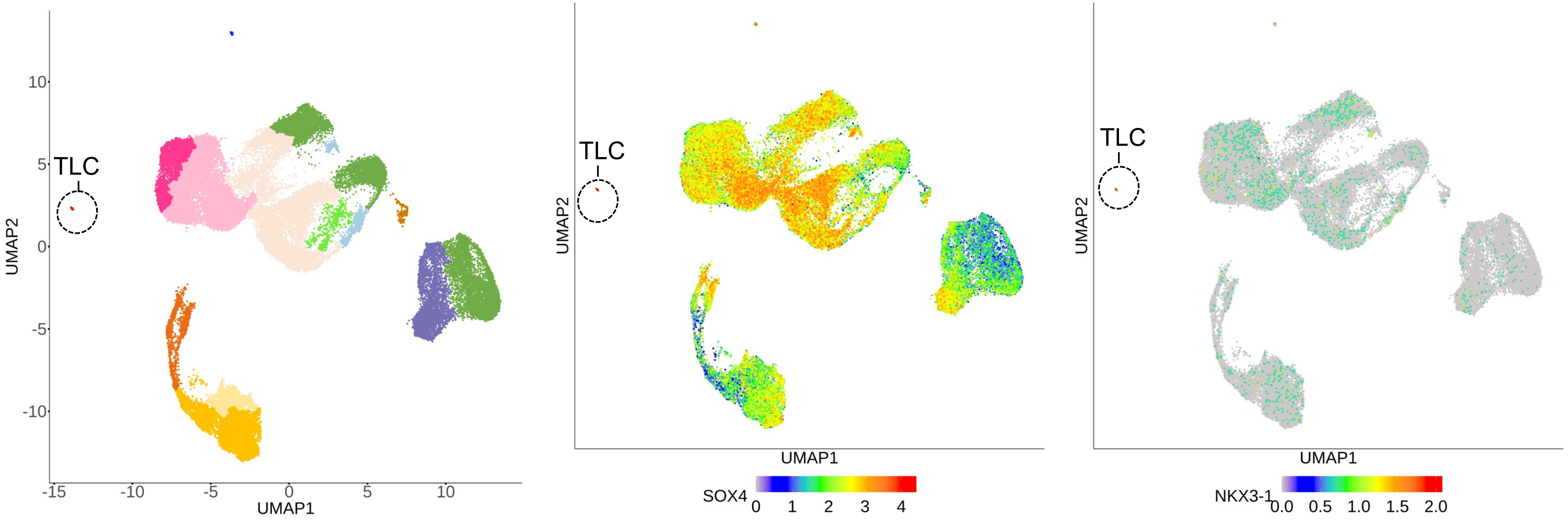
