## Supplementary material for "A Unique Cellular Organization of Human Distal Airways and Its Disarray in Chronic Obstructive Pulmonary Disease": Fig. S10

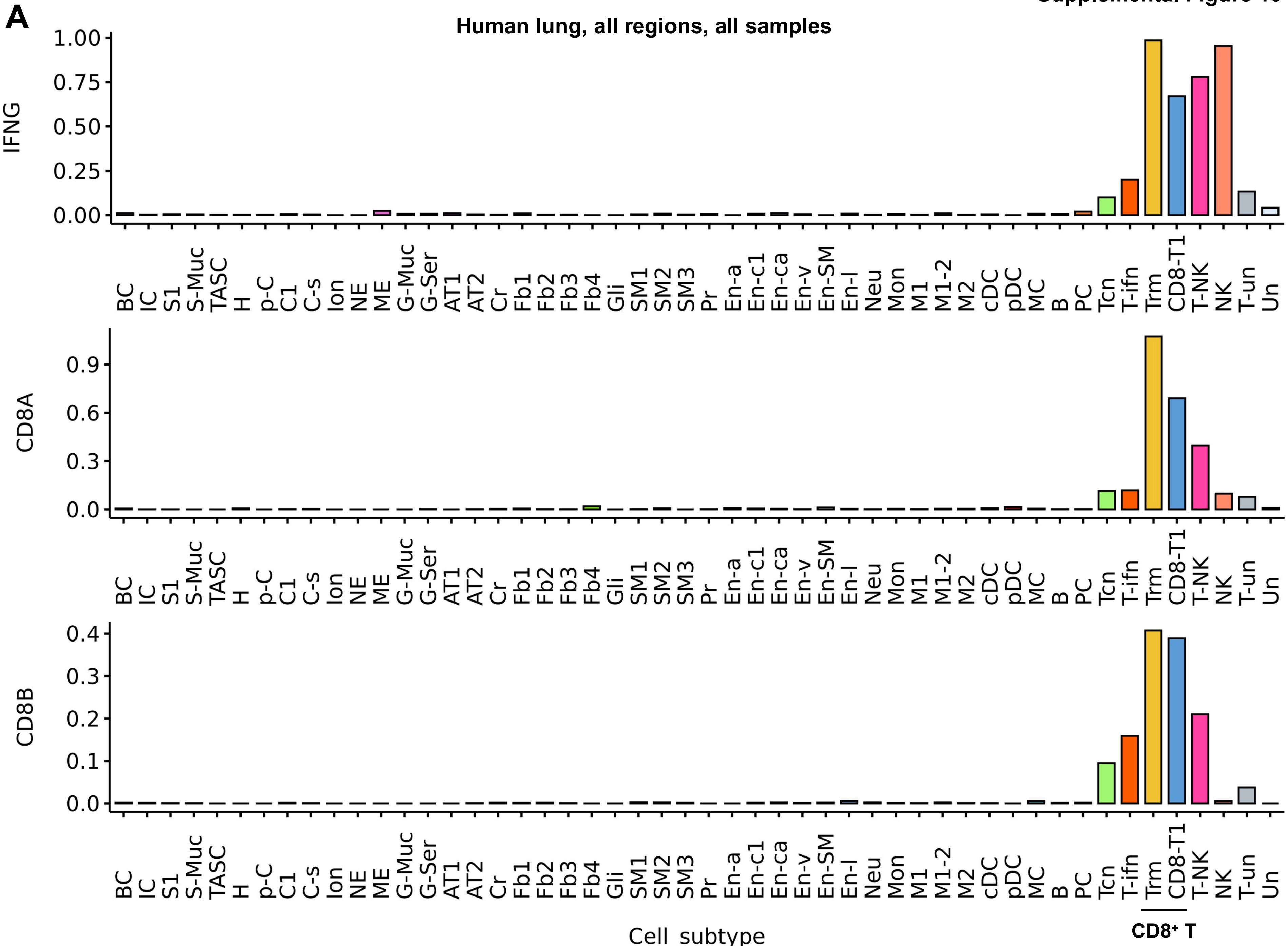
A
Human lung, all regions, all samples
IFNG
CD8A
CD8B
1.00
0.75
0.50
0.25
0.00
0.9
0.6
0.3
0.0
0.4
0.3
0.2
0.1
0.0
BC IC S1 S-Muc TASC H p-C C1 C-s Ion NE ME G-Muc G-Ser AT1 AT2 Cr Fb1 Fb2 Fb3 Fb4 Gli SM1 SM2 SM3 Pr En-a En-c1 En-ca En-v En-SM En-l Neu Mon M1 M1-2 M2 cDC pDC MC B PC Tcn T-ifn Trm CD8-T1 T-NK NK T-un Un
Cell_subtype
CD8+ T

**B$_1$**

**COPD lung tissue samples**

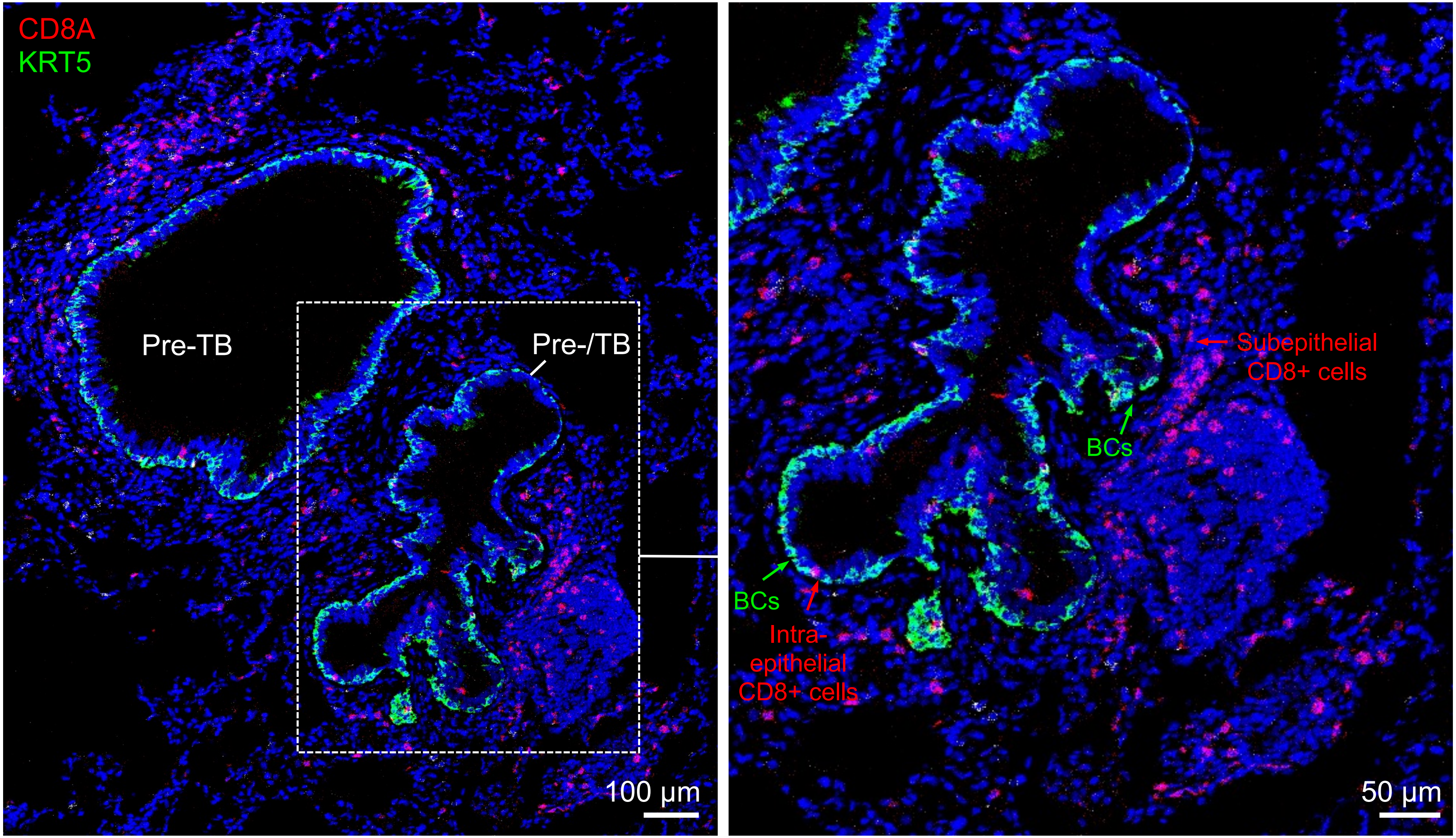

**B$_2$**

**COPD lung tissue samples**

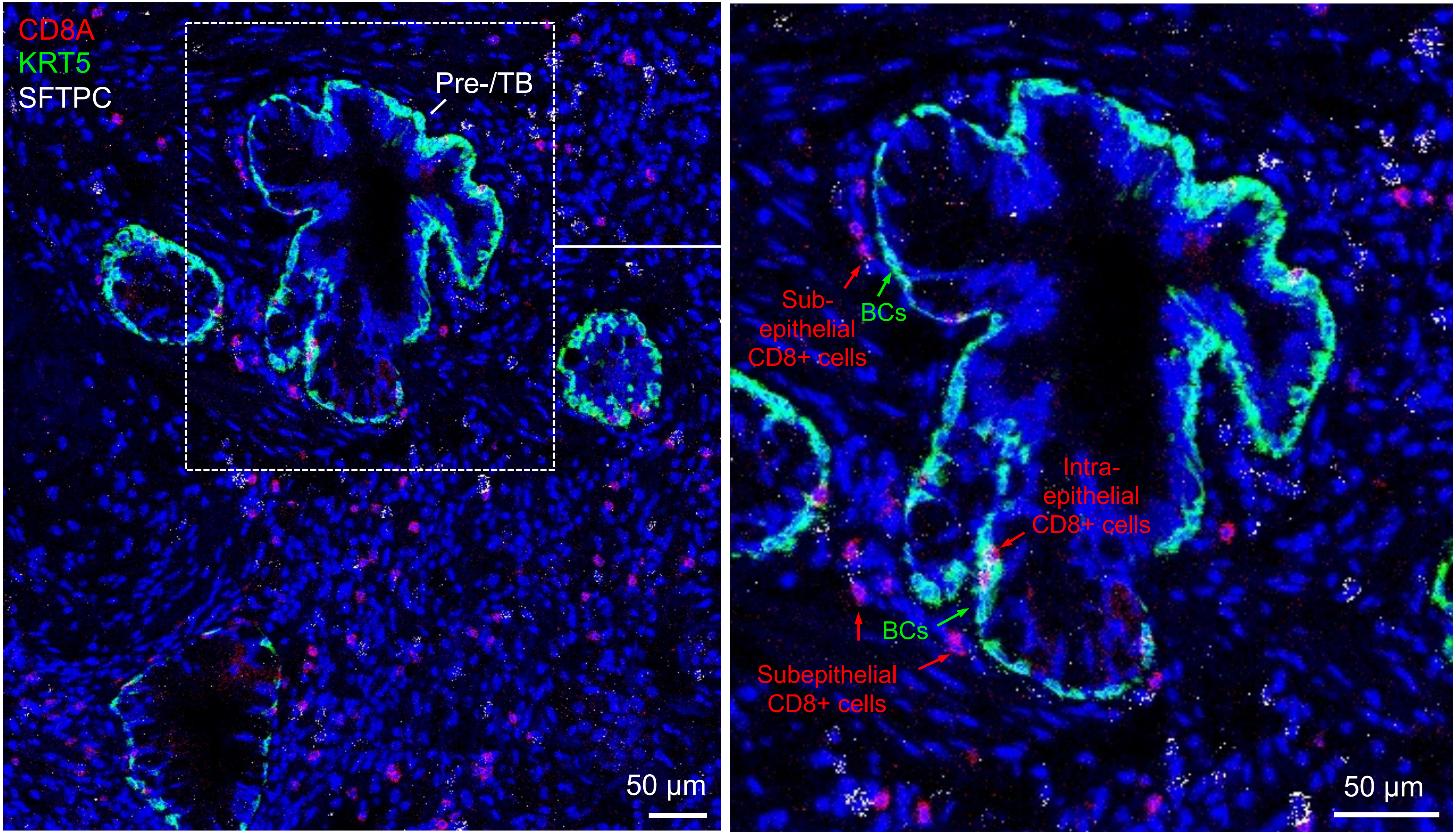

**$B_3$**

**COPD lung tissue samples**

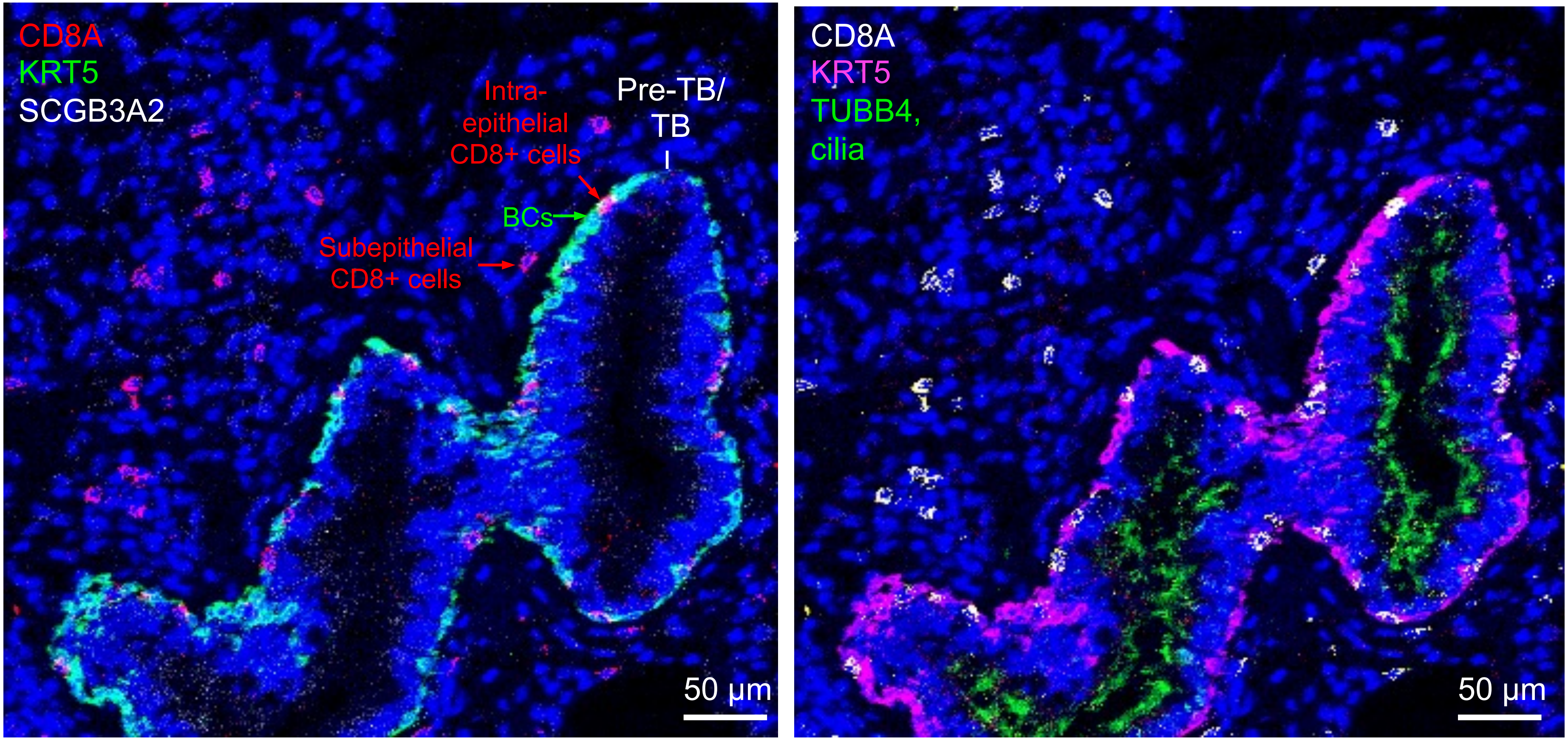

**B$_4$**

### Normal lung tissue samples

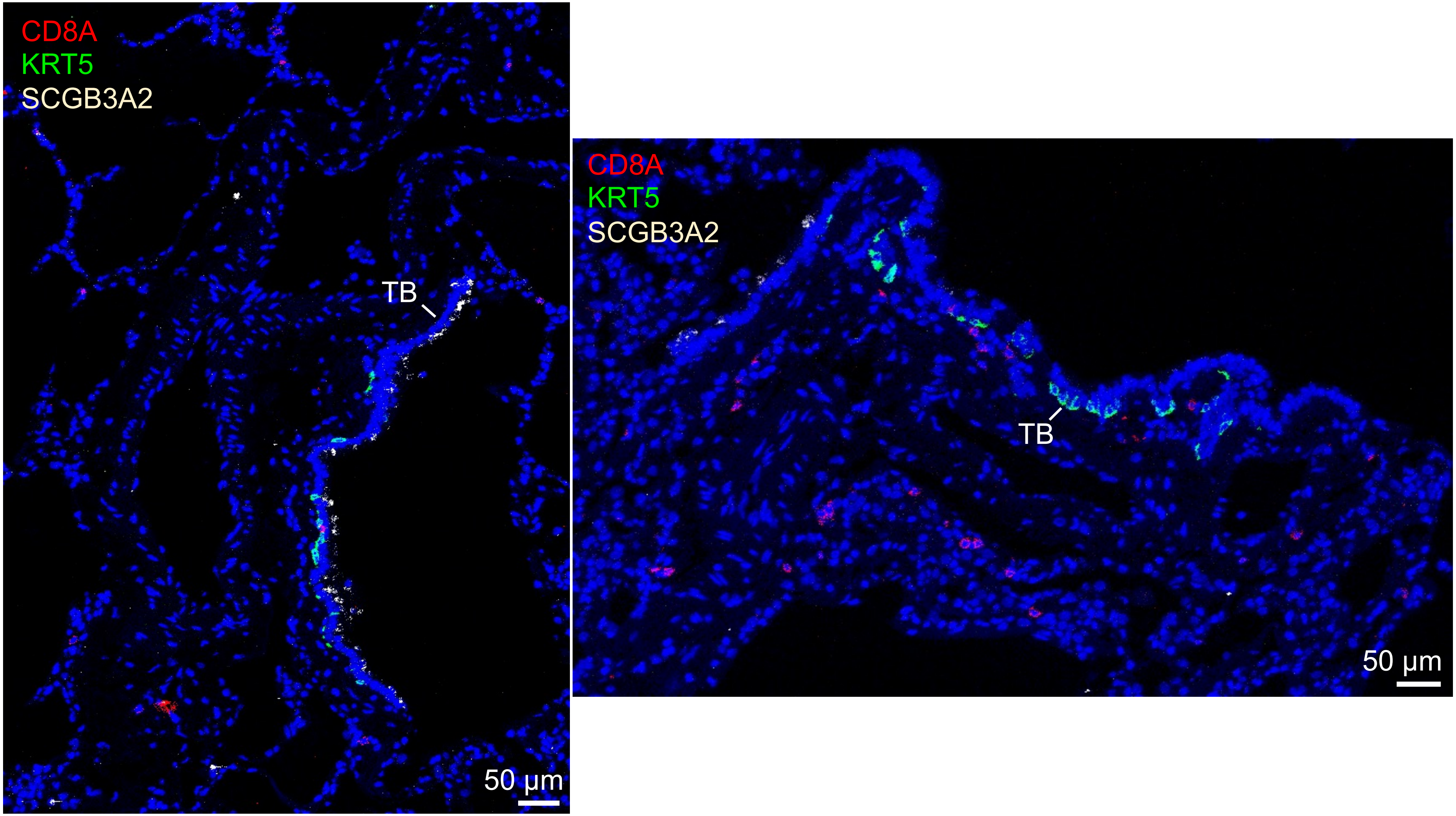

**B$_5$**

### Normal lung tissue samples

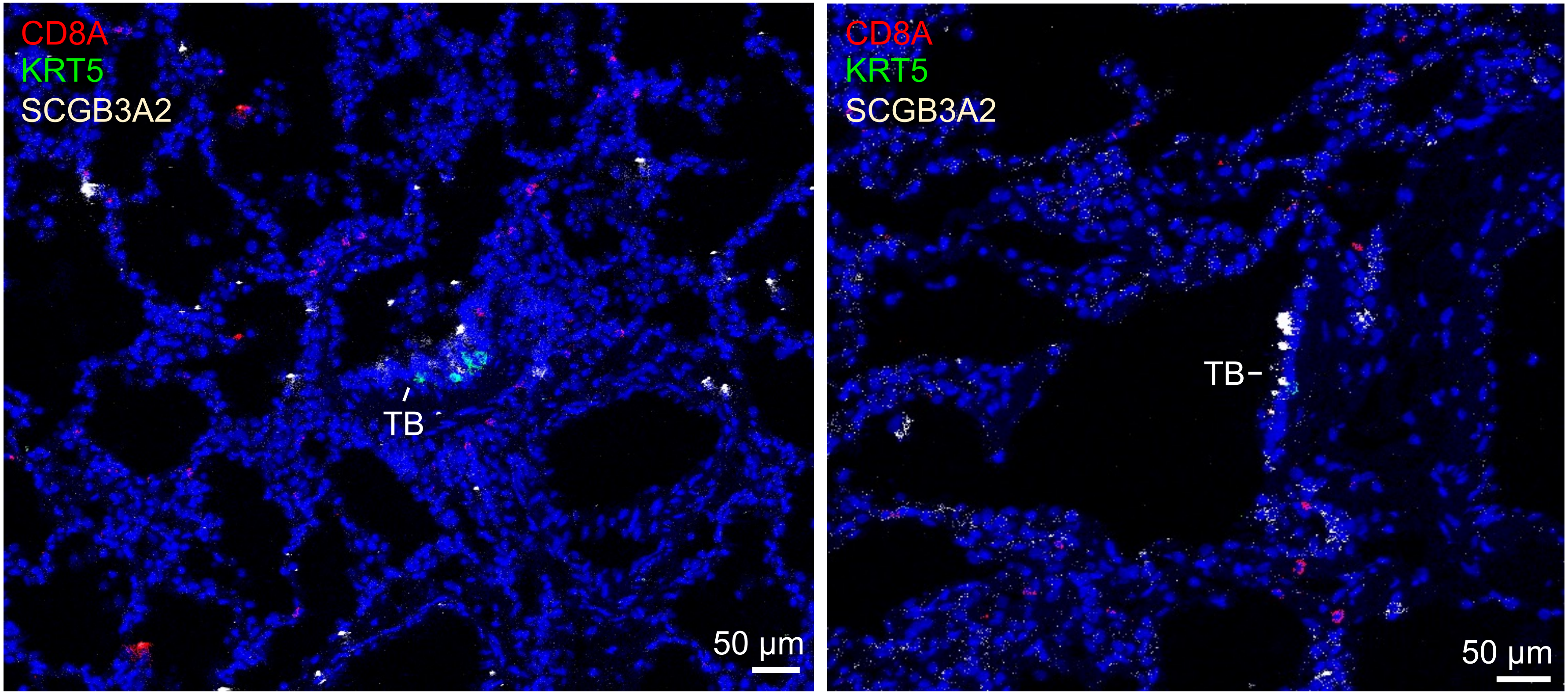

B$_6$

Normal lung tissue samples

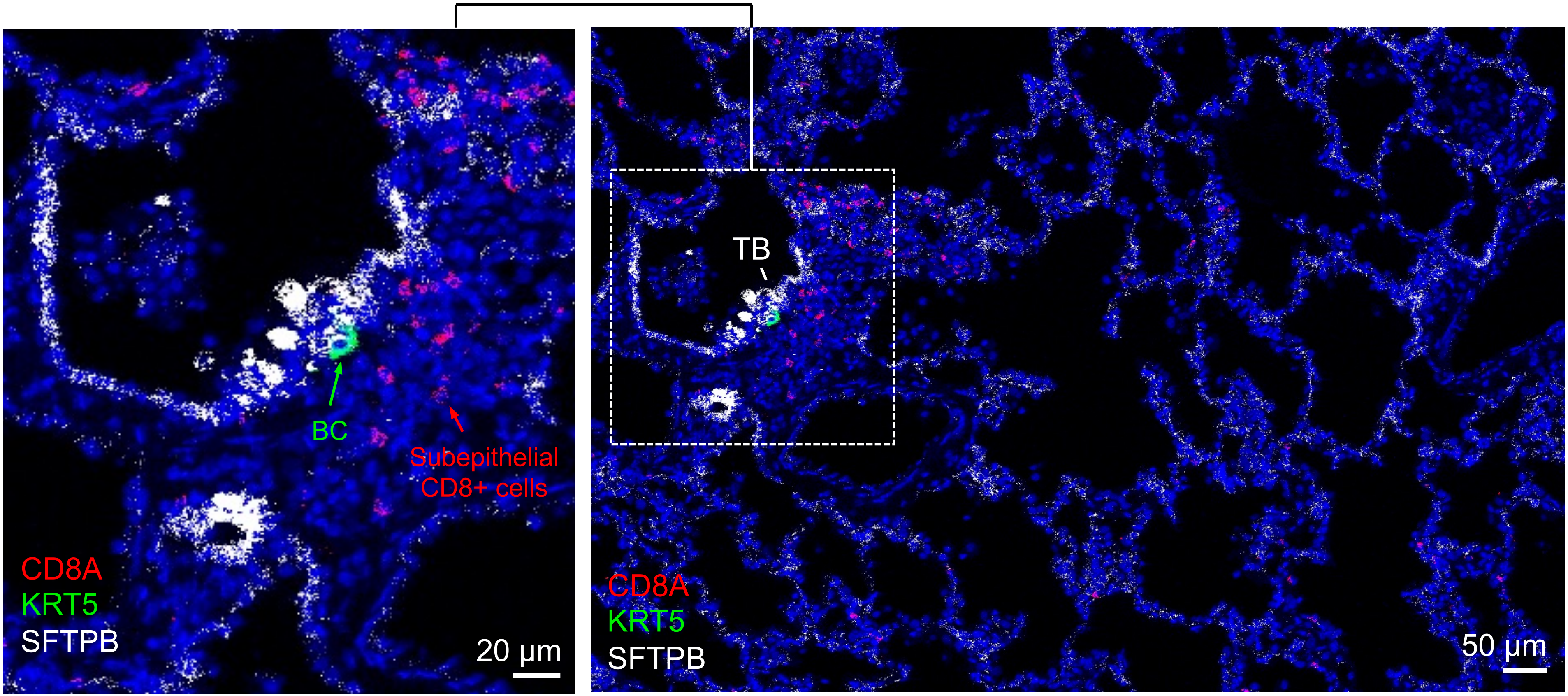

**Supplemental Figure 10**

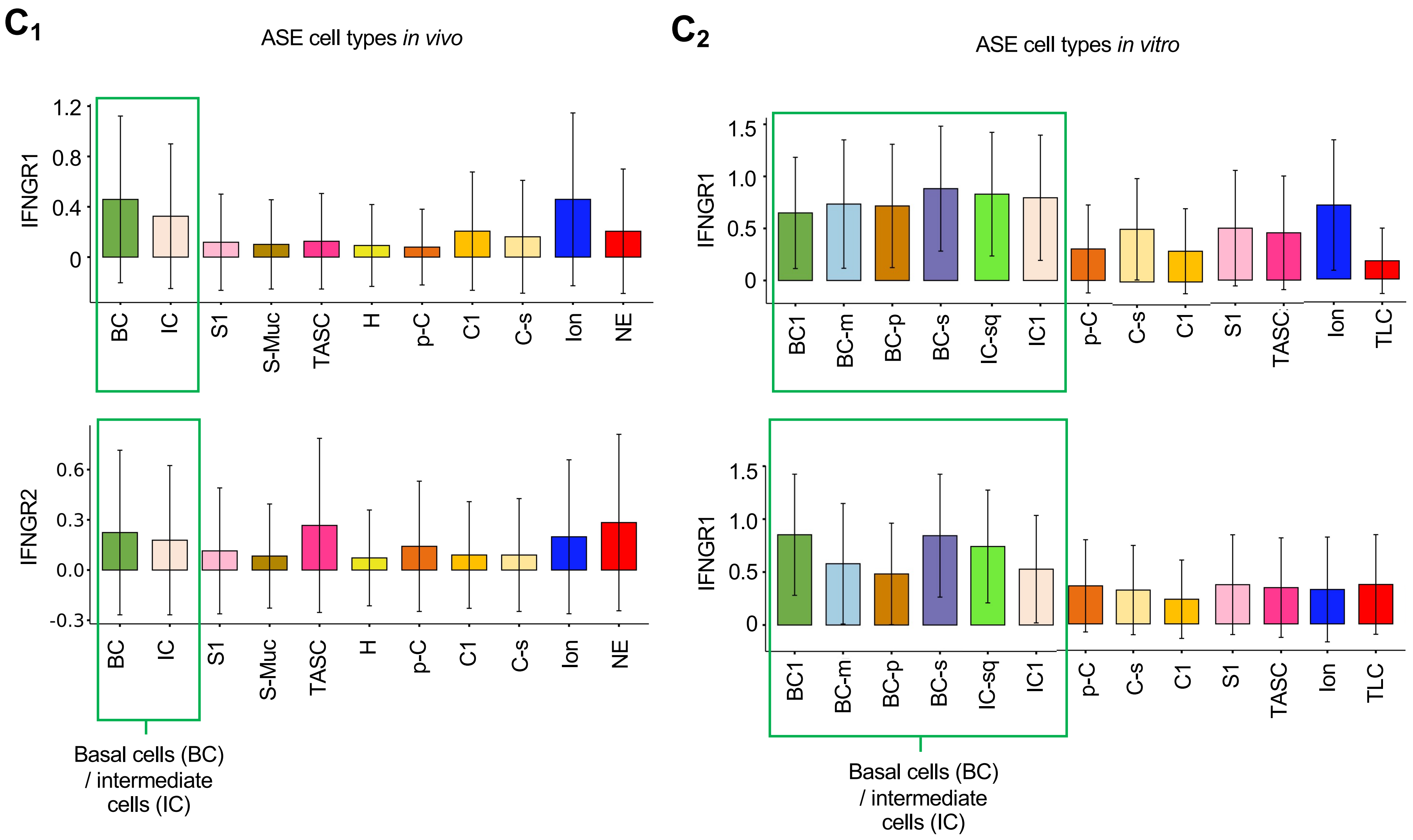

### Supplemental Figure 10

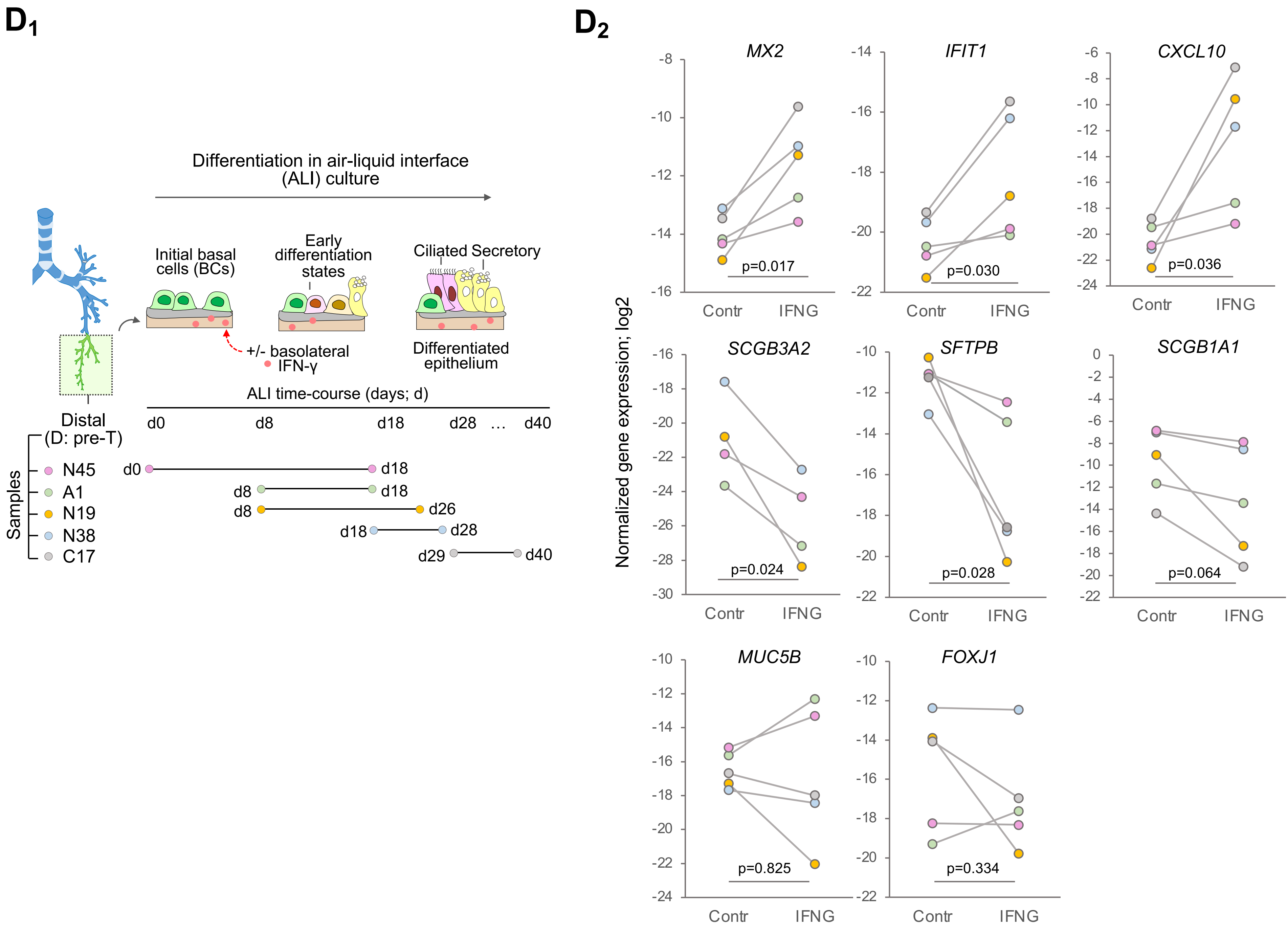
